# A dual-phase enhancer couples progenitor maintenance and pancreatic lineage stability

**DOI:** 10.64898/2026.02.11.705016

**Authors:** Marta Duque, João Amorim, Joana Teixeira, Beatriz Custódio, Mafalda Galhardo, Francisco Camões Magalhães, Joana Marques, Ana Paula Pêgo, José Bessa

**Author notes:** These authors contributed equally to this work (Marta Duque, João Amorim).

## Abstract

Enhancers orchestrate transcriptional programs that control organ development and maintain differentiated cell states, yet how individual enhancers integrate developmental and long-term tissue maintenance logic remains poorly understood. Here, we identify a distal enhancer downstream of *ptf1a* (z3’-DpE) as a regulatory node coupling pancreatic development with acinar cell homeostasis in zebrafish. Deletion of z3’-DpE reduces *ptf1a* expression in pancreatic multipotent progenitor cells (MPCs), leading to depletion of the progenitor pool, altered morphogenesis, and premature exocrine differentiation. Transcriptomic analysis reveals broad repression of proliferation- and morphogenesis-related genes, including Notch pathway components essential for progenitor maintenance. After differentiation, loss of z3’-DpE contributes to acinar cell loss, expansion of ductal and endocrine compartments, and disrupted pancreatic architecture. Chromatin-accessibility profiling of purified acinar cells reveals that reduced *ptf1a* activity leads to widespread remodeling of the acinar chromatin landscape, with decreased accessibility at loci associated with acinar identity and developmental programs, and increased accessibility at sites linked to inflammation, epithelial plasticity, and pancreatic cancer susceptibility. Histopathological analysis shows disorganized acinar tissue with increased duct-like structures and mucinous lesions reminiscent of early pancreatic neoplasia. Thus, z3’-DpE safeguards acinar identity by sustaining *ptf1a* expression and a chromatin landscape that restricts fate instability and pathological plasticity. Our findings demonstrate the mechanistic sufficiency of a single enhancer to coordinate progenitor expansion and long-term lineage stabilization, providing a paradigm for how a developmental regulatory element is redeployed to preserve tissue integrity and suppress disease-associated plasticity.

## Introduction

The pancreas plays a crucial role in human health and disease. It is composed of acinar cells, which secrete digestive enzymes necessary for effective nutrient absorption alongside an extensive ductal network that transports these enzymes. In addition, endocrine cells are organized in islets embedded within the exocrine tissues, responsible for the secretion of hormones essential for maintaining blood sugar balance^1–5^. Dysfunction of the pancreas is linked to major human diseases, including diabetes, pancreatitis, and pancreatic cancer^6,7^. Central to these functions is the pancreas-associated transcription factor 1a (*Ptf1a*) gene, which encodes a basic helix-loop-helix (bHLH) transcription factor (TF). *Ptf1a* is required for pancreatic specification^8^, exocrine versus endocrine fate decision^9–11^, and maintenance of acinar cell identity^8,12,13^, in addition to neurodevelopmental roles^7,14–17^. These multiple functions of *Ptf1a* are achieved by multiple enhancers that activate its expression in specific cell types and developmental time points^18^, including very early in pancreatic multipotent progenitor cells (MPC)^6,19–21^ and later in differentiated acinar cells of the exocrine pancreas^19,21^. The early expression of *Ptf1a* in MPC is controlled by a key enhancer which deletion causes pancreatic agenesis in humans^6^, results that have been recapitulated in *in vivo* models as mouse^20^ and zebrafish^22^. Notably, the deletion of the *Ptf1a* MPC enhancer in mice disrupts chromatin states within MPCs, triggering transcriptomic alterations that hinder pancreatic growth and endocrinogenesis, ultimately causing insulin-deficient neonatal diabetes^20^. The late enhancers responsible for driving *Ptf1a* expression in acinar cells remain incompletely characterized in humans and mice. In contrast to mammals, studies in zebrafish have revealed that the MPC enhancer (z3’-DpE) can also activate expression in differentiated acinar cells^22^. Importantly, *Ptf1a* is regarded as a tumor suppressor gene, as evidenced by studies showing that its conditional knockout in differentiated acinar cells leads to alterations in the transcriptomic profile of cells that culminate in the development of pancreatic ductal adenocarcinoma (PDAC)^12,23^. Further supporting this tumor suppressor function, the ectopic expression of *Ptf1a* has been shown to reduce the tumorigenic properties of PDAC cells^24,25^. It remains to be understood whether mutations in a late acinar *Ptf1a* enhancer are sufficient to activate an oncogenic pathway and what early events drive cells toward a pro-oncogenic state.

In this study, we investigated how the deletion of the z3’-DpE enhancer affects cell state in early MPCs and late differentiated acinar cells in zebrafish. First, we performed transcriptomic analysis of z3’-DpE-responsive MPCs with the z3’-DpE enhancer inactivated and compared them to control cells. Differentially transcribed genes highlighted alterations in Notch signaling pathway components, suggesting that Ptf1a may activate an endogenous Notch signaling program important for MPC differentiation, while other differentially expressed genes suggest alterations in morphogenesis and development, phenotypes that have been further validated by microscopy. Importantly, in later stages of development of z3’-DpE mutant juveniles (z3’-DpE-/-), a decrease in the number of acinar cells was observed with a concomitant increase in the number of duct cells. Furthermore, acinar cells from adult z3’-DpE-/- showed a reduced expression of a reporter specific to acinar cell identity, comparing to control cells, suggesting that z3’-DpE-/- acinar cells could be in a different cellular state. To further test this possibility, we have isolated acinar cells from z3’-DpE mutants and controls and performed Assay for Transposase-Accessible Chromatin with sequencing (ATAC-seq) to probe for changes in chromatin accessibility. We observed that regions with decreased accessibility in z3’-DpE-/- were linked to acinar and developmental genes, while increased accessibility was associated with inflammation, proliferation, epithelial-to-mesenchymal transition (EMT), and pancreatic cancer-related TFs. These results suggest that z3’-DpE-/- acinar cells undergo a shift in regulatory state consistent with a pro-pancreatic cancer program, a conclusion further supported by the spontaneous emergence of pancreatic tumors in adult z3’-DpE mutant animals.

Overall, these results identify the z3’-DpE enhancer as a critical regulatory element that supports pancreatic development and, following differentiation, safeguards acinar cell identity. Loss of z3’-DpE destabilizes the acinar regulatory state, promoting cellular plasticity and creating a permissive context for pancreatic tumorigenesis. These findings suggest that PTF1A-associated regulatory elements in humans may be important non-coding genomic regions targeted by mutations that may contribute to pancreatic cancer development, and their genomic profiling has the potential for preventive and therapeutic approaches to this severe disease.

## Results

### z3’-DpE sustains transcriptional networks governing pancreatic progenitor expansion and identity

In our previous work^22^, we combined ATAC-seq and H3K27ac ChIP-seq profiling of whole adult zebrafish pancreas to identify putative cis-regulatory elements active in pancreatic tissue. This approach revealed a distal non-coding region marked by open chromatin and active enhancer histone modifications, termed z3’-DpE (Figure 1A). Using enhancer reporter assays, where the z3’-DpE sequence was cloned upstream of a minimal promoter driving the expression of GFP, we demonstrated that this element functions as an active enhancer in the MPCs of the ventral pancreatic bud (Figure 1B) and in the acinar cells of the differentiated pancreas (Figure 1C), suggesting that z3’-DpE recapitulates *ptf1a* expression within the pancreatic lineage^10,26,27^. To further investigate the cell-type-specific accessibility of z3’-DpE within the endogenous *ptf1a* locus, we performed ATAC-seq on FACS-purified MPCs and acinar cells. We observed that z3’-DpE is accessible in both populations (Figure 1A), supporting its activity across distinct developmental stages and suggesting a potential role in lineage specification and maintenance.

**Figure 1.**
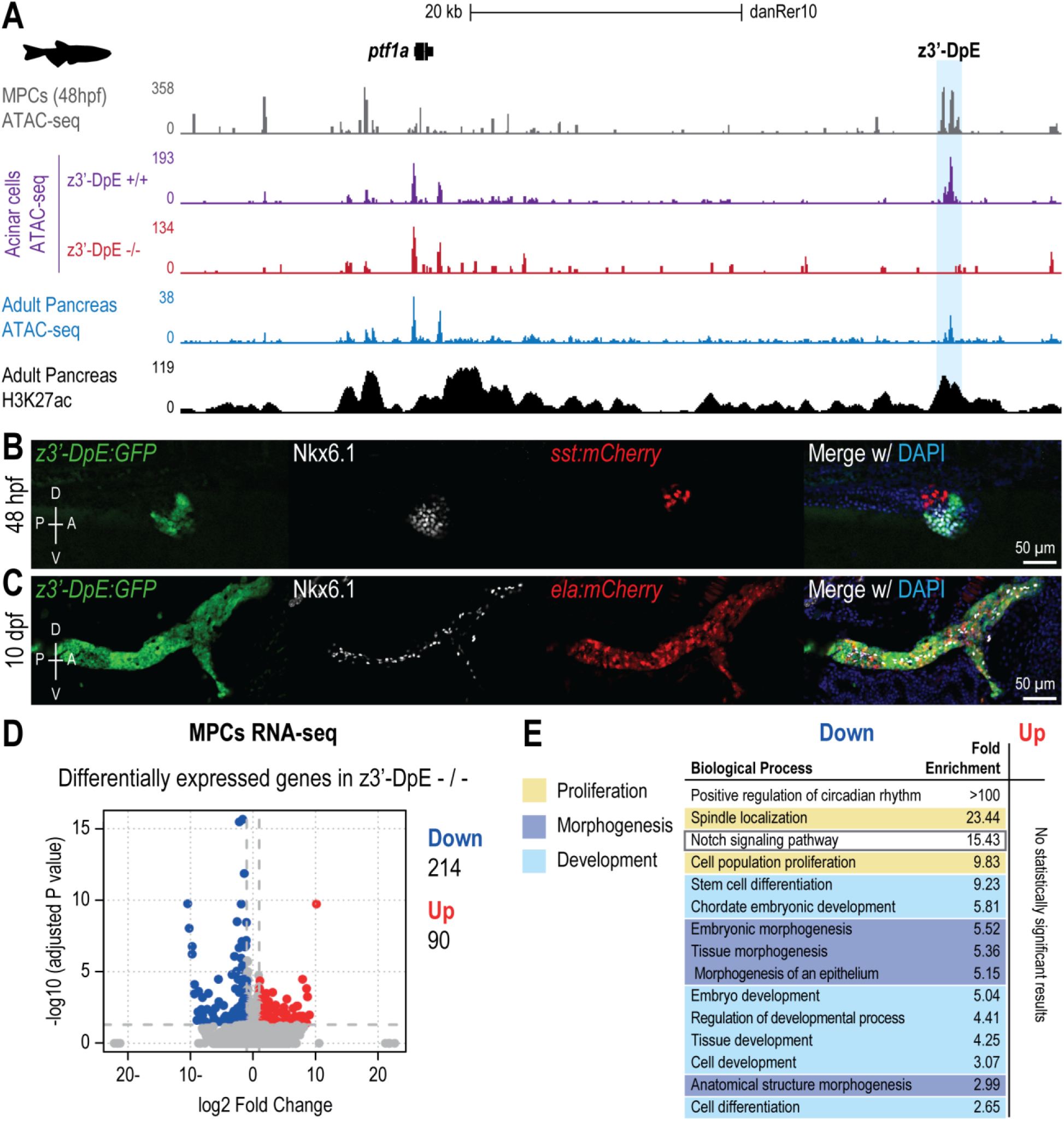
The zebrafish distal *ptf1a* enhancer z3’-DpE is required for transcriptional programs controlling pancreatic progenitor expansion and morphogenesis. **(A)** Genome browser tracks showing ATAC-seq profiles at the endogenous *ptf1a* locus from FACS-purified pancreatic multipotent progenitor cells (MPCs, 48 hpf), adult wild-type and z3’-DpE-/- acinar cells, and adult pancreas, with H3K27ac ChIP-seq density in adult pancreas. The z3’-DpE enhancer (blue) is accessible in wild-type MPCs and mature acinar cells. **(B-C)** Representative confocal maximum intensity projections of the *Tg(z3’-DpE:GFP)* reporter. **(B)** At 48 hpf, GFP (green) colocalizes with Nkx6.1+ MPCs (white) surrounding the principal islet (*sst:mCherry+*, red). **(C)** At 10 dpf, GFP persists in *ela:mCherry+* acinar cells (red) but is excluded from Nkx6.1+ duct cells (white), indicating acinar-restricted enhancer activity. DAPI marks nuclei (blue). D, dorsal; V, ventral; P, posterior; A, anterior. **(D)** Volcano plot of differentially expressed genes in z3’-DpE-/- versus wild-type MPCs (48 hpf). Dashed lines denote adjusted p-value = 0.05 and log_2_ fold change = ±1. Red dots indicate 90 upregulated genes (Up); blue dots indicate 214 downregulated genes (Down). **(E)** GO analysis of downregulated genes shows enrichment for proliferation, Notch signaling, morphogenesis, and developmental processes.

To investigate the functional role of z3’-DpE *in vivo*, we generated a zebrafish line carrying a homozygous deletion of the enhancer (z3’-DpE-/-) using CRISPR-Cas9^22^. We examined the impact of enhancer loss on pancreas development and function, focusing on its early activity in MPCs and its sustained activity in differentiated acinar cells. Previously, we showed that z3’-DpE deletion reduces *ptf1a* expression within the pancreatic progenitor domain, resulting in a smaller pancreatic bud at 48 hpf^22^, closely mimicking the early effects of *ptf1a* gene ablation^10,26,28^. However, unlike *ptf1a* null mutants, which consistently exhibit complete exocrine pancreas loss^10,28^, z3’-DpE-/-mutants display a spectrum of acinar phenotypes, from normal development to mild or severe tissue reduction^22^. This enhancer-driven variability in *ptf1a* expression produces dose-sensitive effects on pancreatic development and offers a unique opportunity to dissect gene regulatory mechanisms, dosage sensitivity, and phenotypic heterogeneity in ways not possible with *ptf1a* null mutants alone. Given the enhancer’s role in modulating *ptf1a* levels and its early activity in MPCs, we next explored its impact on transcriptional programs within these progenitors by performing RNA-seq on FACS-purified MPCs from z3’-DpE-/- and z3’-DpE+/+ embryos. Differential expression analysis revealed 214 downregulated and 90 upregulated genes in z3’-DpE-/- MPCs relative to controls (Figure 1D, Supplementary Figure 1). While upregulated genes showed no significant enrichment for specific biological pathways, downregulated genes were strongly associated with key processes governing progenitor expansion and tissue organization (Figure 1E; Supplementary Table 1). Proliferation-related pathways were among the most affected, with significant enrichment of Gene Ontology (GO) terms such as spindle localization (GO:0051653; 23.44-fold, p = 2.87 × 10^-4^) and cell population proliferation (GO:0008283; 9.83-fold, p = 1.61 × 10^-4^), indicating impaired proliferative capacity in z3’-DpE-/- MPCs. This is further supported by the smaller bud size seen in these mutants, consistent with impaired progenitor proliferation. Notably, genes within the Notch signaling pathway (GO:0007219) were also downregulated (15.43-fold, p = 1.35 × 10^-4^), suggesting that z3’-DpE may regulate progenitor maintenance and differentiation through Notch-dependent mechanisms^18^. Morphogenesis-related terms, including embryonic morphogenesis (GO:0048598; 5.52-fold, p = 5.98 × 10^-7^), tissue morphogenesis (GO:0048729; 5.36-fold, p = 1.72 × 10^-5^), and anatomical structure morphogenesis (GO:0009653; 2.99-fold, p = 1.24 × 10^-6^), were also significantly enriched, pointing to disrupted pancreatic architecture. In addition, developmental programs such as stem cell differentiation (GO:0048863; 9.23-fold, p = 1.12 × 10^-5^), chordate embryonic development (GO:0043009; 5.81-fold, p = 8.54 × 10^-6^), embryo development (GO:0009790; 5.04-fold, p = 4.43×10^-9^), and tissue development (GO:0009888; 4.25-fold, p = 4.85 × 10^-9^) were broadly downregulated, implicating z3’-DpE in regulating transcriptional networks that control progenitor fate and lineage specification.

Together, these findings establish z3’-DpE as a critical enhancer required to sustain the transcriptional networks that govern MPC proliferation, morphogenesis, and fate determination during early pancreatic organogenesis. Loss of z3’-DpE disrupts Notch signaling and destabilizes the transcriptional networks that preserve progenitor identity and developmental potential.

### z3’-DpE loss leads to MPC depletion and premature exocrine differentiation

Loss of z3’-DpE was previously shown to reduce *ptf1a* expression and diminish the Nkx6.1+ pancreatic progenitor domain in z3’-DpE-/- embryos at 48 hpf^22^. Consistent with this, our RNA-seq analysis revealed downregulation of genes essential for pancreatic development (Figure 1E, Supplementary Figure 1), suggesting broader developmental consequences beyond the previously described reduction in Nkx6.1+ cells. At the organismal level, genotypes were recovered at approximately Mendelian ratios from embryogenesis to adulthood (Supplementary Figure 2A), indicating that loss of z3’-DpE does not impair overall viability despite these transcriptomic alterations. To follow the z3’-DpE-/- progenitors throughout pancreas development, we generated zebrafish carrying the *Tg(z3’-DpE:GFP)* transgene on a z3’-DpE-/- background and compared them to wild-type embryos. This strategy enabled visualization of the MPCs that would normally receive z3’-DpE-driven regulatory input (Figure 2A). Immunostaining of Nkx6.1, expressed in progenitor populations during embryonic stages and mature duct cells in larvae and adults^29^, was also performed, as *nkx6.1* expression precedes *ptf1a* expression in MPCs during early pancreas development, making it a robust marker for tracking the full MPC population during development^18,29,30^.

**Figure 2.**
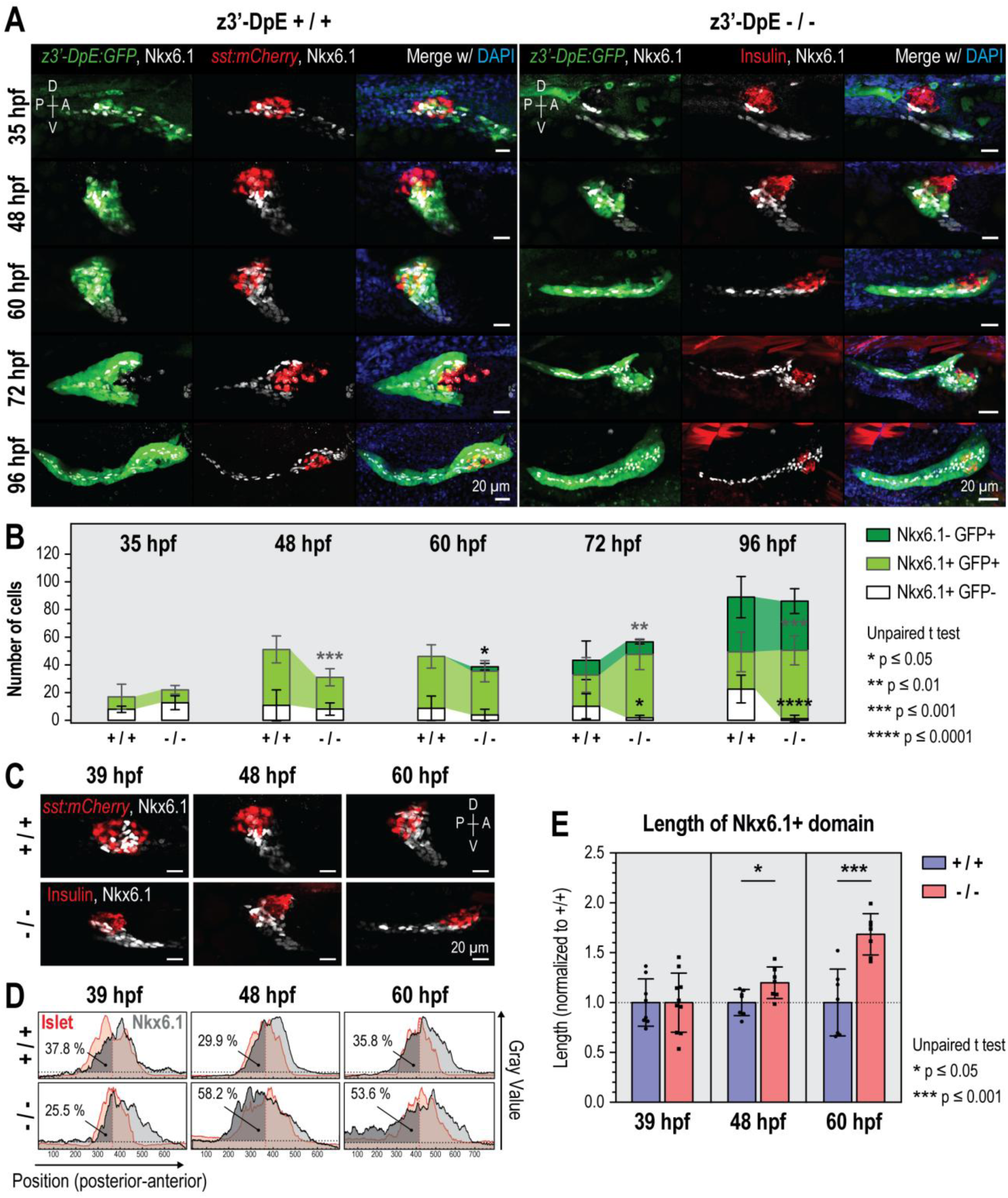
Loss of z3’-DpE impairs MPC expansion and disrupts the pancreatic progenitor domain. **(A)** Representative confocal maximum intensity projections of *Tg(z3’-DpE:GFP); Tg(sst:mCherry)* z3’-DpE+/+ (left) and *Tg(z3’-DpE:GFP)* z3’-DpE-/- embryos (right) from 35 to 96 hpf; Nkx6.1 marks MPCs (white), *sst:mCherry* and Insulin mark islet cells (red), and DAPI marks nuclei (blue). **(B)** Quantification of Nkx6.1+GFP-, Nkx6.1+GFP+, and Nkx6.1-GFP+ cells per embryo (mean ± SD). **(C)** Nkx6.1 (white) and Islet markers (red) at 39, 48, and 60 hpf; the principal islet is marked by *sst:mCherry* expression in z3’-DpE+/+ and Insulin staining in z3’-DpE-/-. **(D)** Distribution profiles of Nkx6.1 (gray) and Islet (red) markers along the posterior-anterior axis. **(E)** Length of the Nkx6.1+ domain normalized to wild type (mean ± SD). D, dorsal; V, ventral; P, posterior; A, anterior.

Using this system, we defined three major cell populations: (1) Nkx6.1+GFP-early progenitors, in which the enhancer is not yet active; (2) Nkx6.1+GFP+ late progenitors, which would normally receive z3’-DpE regulatory input; and (3) Nkx6.1-GFP+ pro-acinar cells, transitioning toward exocrine fate (Figure 2A-B, Supplementary Figure 2B-C). Developmental differences between wild-type and z3’-DpE-/- embryos emerged at 48 hpf, when the total number of MPCs was significantly reduced in mutants (31.1 ± 7.7 vs. 47.9 ± 8.1, p = 9.00 × 10^-4^, Figure 2B), consistent with our previous findings^22^. This reduction was primarily due to a decrease in the pool of Nkx6.1+GFP+ late progenitors (23.0 ± 5.9 vs. 40.6 ± 8.6, p = 3.00 × 10^-4^), whereas the Nkx6.1+GFP- population remained stable across genotypes (8.1 ± 4.2 vs. 7.3 ± 2.1, p = 0.49). Notably, z3’-DpE-/- embryos exhibited premature emergence of pro-acinar cells. In these mutants, Nkx6.1-GFP+ cells were observed by 60 hpf, 12 hours earlier than in wild types, indicating an accelerated transition from progenitor to exocrine fate (Figure 2B). Although final acinar cell numbers were comparable by 96 hpf (35.5 ± 8.5 vs. 41.5 ± 14.8, p = 0.49), the Nkx6.1+GFP- progenitor pool was nearly exhausted in mutants (0.4 ± 0.7 vs. 24.0 ± 8.8, p < 1.00× 10^-4^) (Figure 2B and Supplementary Figure 2B), indicating a premature depletion of MPCs and early commitment to differentiated lineages, consistent with the model that z3’-DpE maintains progenitor multipotency and controls differentiation timing. The accelerated lineage commitment was accompanied by abnormal spatial distribution of MPCs. At 48 and 60 hpf Nkx6.1+ staining was predominantly posterior to the principal islet in mutants, whereas in wild-type embryos the signal remained mostly anteriorly distributed (Figure 2C-D). Additionally, at 60 hpf the Nkx6.1+ staining is markedly more dispersed along the anterior-posterior axis, as evidenced by an increased domain length (Figure 2E), without an associated increase in Nkx6.1+ cell numbers (Figure 2B), pointing to altered morphogenesis and disrupted spatial organization upon enhancer loss.

Collectively, these findings demonstrate that z3’-DpE is essential for expanding and maintaining the pancreatic progenitor pool during organogenesis. Its loss leads to premature differentiation and spatial mispatterning, underscoring the enhancer’s pivotal role in sustaining multipotency and coordinating morphogenesis and the timing of exocrine fate allocation.

### z3’-DpE deletion redistributes pancreatic lineages and disrupts organ architecture

z3’-DpE-/- animals displayed no overt morphological abnormalities compared with z3’-DpE+/+ siblings, aside from pancreas-specific defects^22^. Consistently, morphometric analysis of multiple larval anatomical structures, including eye and retina area, inner ear area, otolith area, somite length, and notochord thickness, revealed no significant differences between genotypes (Supplementary Figure 3A). To quantitatively assess acinar tissue, we utilized the *Tg(ela:mCherry)* transgenic reporter line, which expresses mCherry specifically in pancreatic acinar cells. In z3’-DpE-/- mutant larvae (10-13 dpf), the mCherry-expressing domain was reduced (Figure 3A-B, Supplementary Figure 3B-C). Based on the extent of acinar tissue loss, larvae could be categorized into three severity groups: Severe, Moderate, and No Reduction (Figure 3C, Supplementary Figure 3D), as previously reported^22^. Furthermore, we utilized the *Tg(ins:GFP)* transgene to label differentiated pancreatic beta-cells, while pancreatic duct cells were identified via immunostaining for Nkx6.1. These complementary markers revealed that the overall reduction in acinar tissue in z3’-DpE-/- mutants was accompanied by an increase of pancreatic ducts and endocrine beta-cells (Figure 3B, Supplementary Figure 3C and E), suggesting a shift in pancreatic lineage allocation favoring non-acinar lineages at the expense of acinar cells. Notably, z3’-DpE-/- larvae exhibited reduced *ela:mCherry* fluorescence intensity in the pancreas (Figure 3D), suggesting that while MPCs are still able to differentiate into elastase-producing acinar cells, these cells may differ from wild-type acinar cells in their transcriptional output, maturity, or functional state.

**Figure 3.**
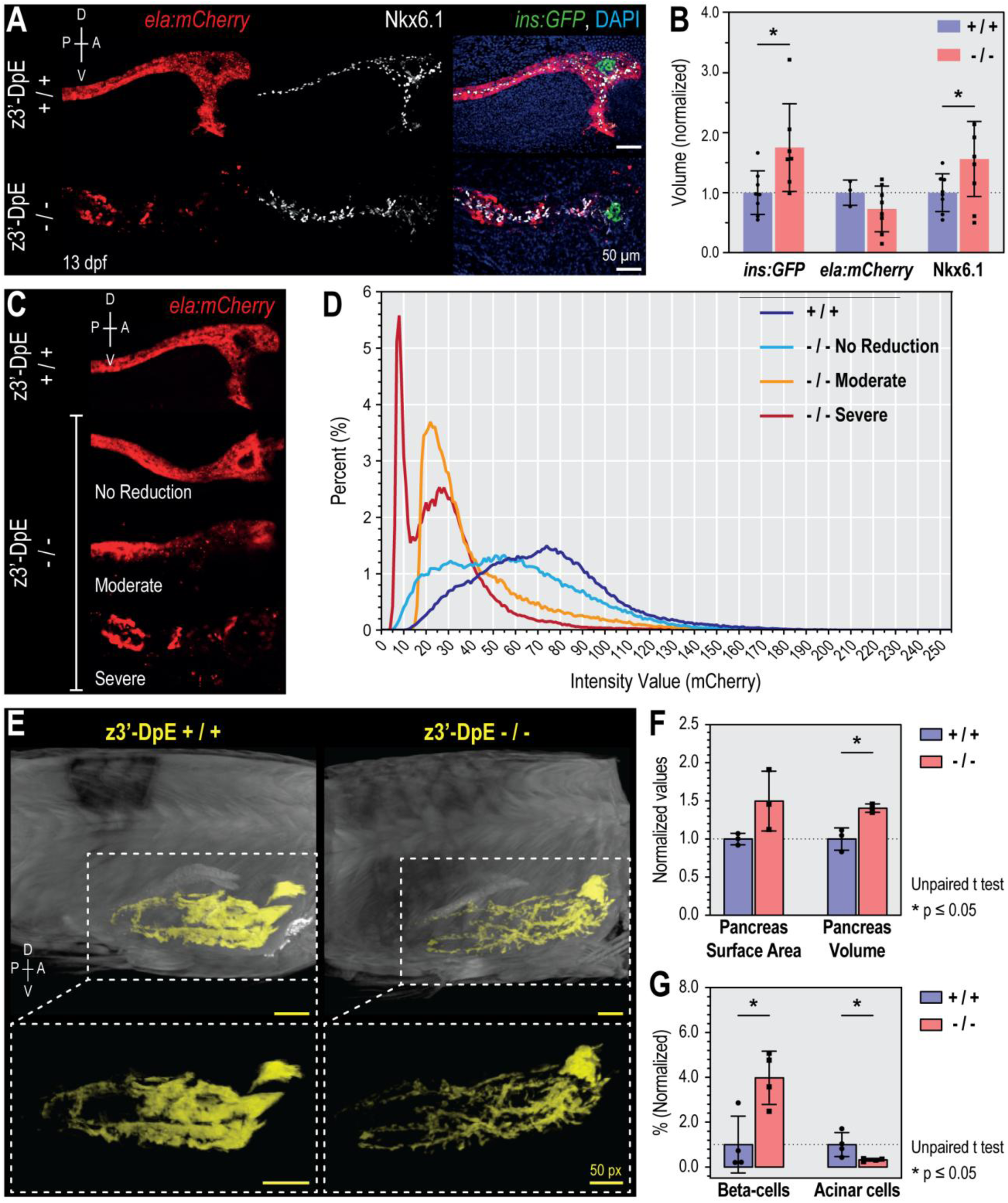
Loss of z3’-DpE leads to acinar cell depletion and altered adult pancreas architecture. **(A)** Representative confocal maximum intensity projections of *Tg(ins:GFP); Tg(ela:mCherry)* larvae at 13 dpf; *ela:mCherry* marks acinar cells (red), Nkx6.1 duct cells (white), *ins:GFP* endocrine β-cells (green), and DAPI nuclei (blue). **(B)** Quantification of marker volumes normalized to wild type (mean ± SD). **(C)** Phenotypic variability in z3’-DpE-/- larvae classified as no reduction, moderate, or severe acinar loss. **(D)** Histogram of mean *ela:mCherry* fluorescence intensity (linearly normalized, 0-255) showing a leftward shift in mutants. **(E)** Three-dimensional micro-CT reconstructions of adult males (sagittal planes); total body volume in grayscale and segmented pancreatic tissue in yellow. **(F)** Quantitative micro-CT analysis of pancreatic surface area and volume normalized to wild type (mean ± SD). **(G)** Flow-cytometry quantification of β-and acinar-cell fractions in adults normalized to wild type (mean ± SD). D, dorsal; V, ventral; P, posterior; A, anterior.

Next, we asked whether the disruption of pancreatic morphology and altered cell composition observed in larvae persisted in adulthood. Micro-CT imaging of adult pancreata revealed that while z3’-DpE+/+ pancreata were compact and well organized, z3’-DpE-/- pancreata displayed fragmented, irregular architecture dominated by duct-like structures (Figure 3E). Additionally, z3’-DpE-/- pancreata exhibited an increase in both surface area and overall volume (Figure 3F), despite the significant decrease in acinar cell numbers (Figure 3G). Under normal physiological conditions, the bulk of pancreas surface area and volume is accounted for by densely packed acinar tissue^22,31^. The depletion of acinar cells in z3’-DpE-/- mutants, coupled with increased organ size, indicates that non-acinar compartments (ductal and endocrine) undergo disproportionate expansion and/or that tissue architecture becomes more dispersed and fragmented. Quantitative analysis confirmed a 0.32-fold reduction in acinar cells (p = 1.39 × 10^-2^) and a 3.98-fold increase in beta-cells (p = 4.49 × 10^-2^), resulting in a dramatic shift in the beta-to-acinar cell ratio (1.33 vs. 0.08 in wild-type, Figure 3G).

Subsequently, we performed single-cell RNA sequencing of whole adult pancreata from z3’-DpE+/+ and z3’-DpE-/- animals. We recovered a total of 10,871 cells across all clusters, of which only 826 cells were assigned to the pancreatic parenchymal cluster (cluster 6, Supplementary Figure 4A-B). Due to this limited representation of acinar, ductal, and endocrine cells within cluster 6, our analysis did not yield robust or biologically interpretable differentially expressed gene signatures for these main pancreatic lineages (Supplementary Table 2), with the single-cell data primarily informing shifts in cellular composition. The data confirmed a shift in cellular composition, revealing an expansion of ductal (12.2% vs. 5.6%) and endocrine (19.1% vs. 6.4%) compartments in mutants compared with controls (p < 0.05; Supplementary Figure 4B). Consistent with these compositional changes, the distribution of differentially expressed genes (DEGs) across clusters (Supplementary Table 3) showed that pancreatic cells contributed the largest fraction of DEGs in z3’-DpE-/- versus z3’-DpE+/+ pancreata, with most of these DEGs being downregulated and restricted to the pancreatic cluster (Supplementary Figure 4C). This pattern supports the notion that z3’-DpE loss preferentially perturbs transcriptional programs within pancreatic parenchymal cells, while having comparatively smaller effects on non-pancreatic stromal and immune populations.

Altogether, these results establish that z3’-DpE is essential for maintaining acinar cell allocation and proper organ architecture, restricting both the expansion of alternative pancreatic lineages and abnormal tissue remodeling during development and into adulthood. This abnormal remodeling, characterized by increased ductal and endocrine tissue at the expense of acinar cells, may predispose the organ to functional deficits or disease, such as chronic pancreatitis or tumorigenesis.

### z3’-DpE deletion remodels acinar chromatin, driving lineage instability and oncogenic-primed states

Given the reduction in *ela:mCherry* fluorescence intensity in acinar cells of z3’-DpE-/- larvae (Figure 3D), we hypothesized that their cellular state was altered. To investigate whether loss of z3’-DpE alters acinar cell state at the epigenomic level, we performed ATAC-seq on FACS-purified acinar cells from adult wild-type and z3’-DpE-/-zebrafish. Peak-calling analysis identified 35,887 accessible regions in wild-type acinar cells and 30,669 in mutant acinar cells, with 20,176 regions exclusive to wild-type (56.2% of wild-type peaks), 14,842 exclusive to mutants (48.4% of mutant peaks), and 15,711 shared between both conditions (Supplementary Figure 5A). These results show that, although genome wide chromatin availability is not being broadly altered in z3’-DpE-/-acinar cells, open regions are being significantly shifted potentially affecting the regulatory landscapes of genes. While 72.2% of chromatin available peaks located in the same genomic positions in wild-type and mutant acinar cells overlap with promoter regions (Common peaks; Supplementary Figure 5A), most genotype-exclusive peaks are mapped to non-promoter genomic regions (Exclusive peaks; 83.5% in wild-type; 82.5% in mutant; Supplementary Figure 5A), suggesting that z3’-DpE primarily regulates acinar chromatin accessibility of cis-regulatory sequences. We next performed differential accessibility analysis to compare ATAC-seq signal at these regions between wild-type and mutant acinar cells, applying a statistical cutoff to identify significantly altered loci. This analysis revealed 5,968 regions with decreased (Down regions) and 4,475 with increased accessibility (Up regions) in z3’-DpE-/- acinar cells (Figure 4A-B). GO enrichment analysis of genes associated with these regions (Supplementary Tables 4 and 5) revealed that loss of z3’-DpE significantly alters pathways that can be broadly grouped into Acinar cell function, Endocrine cell function, Developmental processes, Notch signaling, Proliferation, Regulation of EMT & Tissue Migration, and Inflammatory processes, rather than causing broad, nonspecific chromatin disruption (Figure 4C, Supplementary Figure 5B-C). Down regions were preferentially located near key genes involved with exocrine pancreas activity, such as digestive system process (GO:0022600), digestion (GO:0007586), and digestive tract morphogenesis/development (GO:0048546/GO:0048565), suggesting that chromatin closing at these loci may directly impair transcription of genes necessary for acinar identity, differentiation, and digestive enzyme production (Figure 4C, Supplementary Figure 5B-C). Other terms also include pancreas development (GO:0031016), regulation of endodermal cell fate specification (GO:0042663), and determination of pancreatic left/right asymmetry (GO:0035469) (Figure 4C, Supplementary Figure 5B-C), indicating that the chromatin changes extend to loci that drive organogenesis and tissue patterning, potentially underpinning the early progenitor depletion and morphogenic defects within the pancreatic progenitor domain observed in mutant embryos (Figure 2). Notably, Down regions were also associated with genes linked to insulin secretion (GO:0030073), hormone secretion (GO:0046879), and response to glucose (GO:0009749), which indicates that z3’-DpE not only safeguards exocrine fate, but may also intersect with regulatory programs affecting endocrine competence (Figure 4C, Supplementary Figure 5B-C). This suggests that the cells are in a regulatory state of plasticity or dedifferentiation, characterized by broad suppression of mature lineage programs (both acinar and endocrine) and potential emergence of new, non-lineage-specific, stress/adaptation pathways. In line with this idea, Up regions highlight a restructuring of the acinar cell regulatory landscape toward pathways that are not typically active in mature acinar cells. GO terms enriched in Up regions include regulation of leukocyte differentiation (GO:1902105), regulation of T cell differentiation (GO:0045580), and macrophage activation involved in immune response (GO:0002281), suggesting that the acinar cells in mutants may acquire features reminiscent of tissue stress or inflammation (Figure 4C, Supplementary Figure 5B-C). Up regions also included those linked to cell division (spindle localization, GO:0051653), cellular proliferation (endothelial cell proliferation, GO:0001935; epithelial cell proliferation, GO:0050673), and tissue repair (wound healing, GO:0044319), further suggesting the activation of pathways typical of regenerative and injury-response programs and underlining a shift from stable, mature acinar identity to a plastic, repair-prone, and potentially transformation-permissive state (Figure 4C, Supplementary Figure 5B-C). Such broad chromatin remodeling is consistent with cellular dedifferentiation and a microenvironment conducive to metaplasia or early neoplastic development^32^. In agreement with this regulation of epithelial to mesenchymal transition (GO:0010717), another well-established marker of increased cellular plasticity and indicator of both metaplastic and tumorigenic potential^33–36^, also displayed significant enrichment in the Up regions (Figure 4C, Supplementary Figure 5B-C). These changes included loci such as *crim1*, a negative modulator of BMP/TGF-β signaling^37,38^, a pathway that is downregulated in later stages of pancreatic cancer progression^39^. Additionally, *cxcl20,* a teleost specific inducible chemokine with the potential to shape local inflammatory conditions^40^, also showed increased chromatin accessibility in mutant acinar cells and upregulation in z3’-DpE-/- pancreatic cells (cluster 6; Supplementary Figure 5D). Upregulation of *cxcl20* is consistent with the acquisition of an inflammatory microenvironment which may initially support tumor survival and progression^41^. These two cases illustrate how z3’-DpE loss not only remodels broad pathway-level chromatin accessibility but also leads to locus-specific changes at genes potentially linked to acinar stress and inflammation.

**Figure 4.**
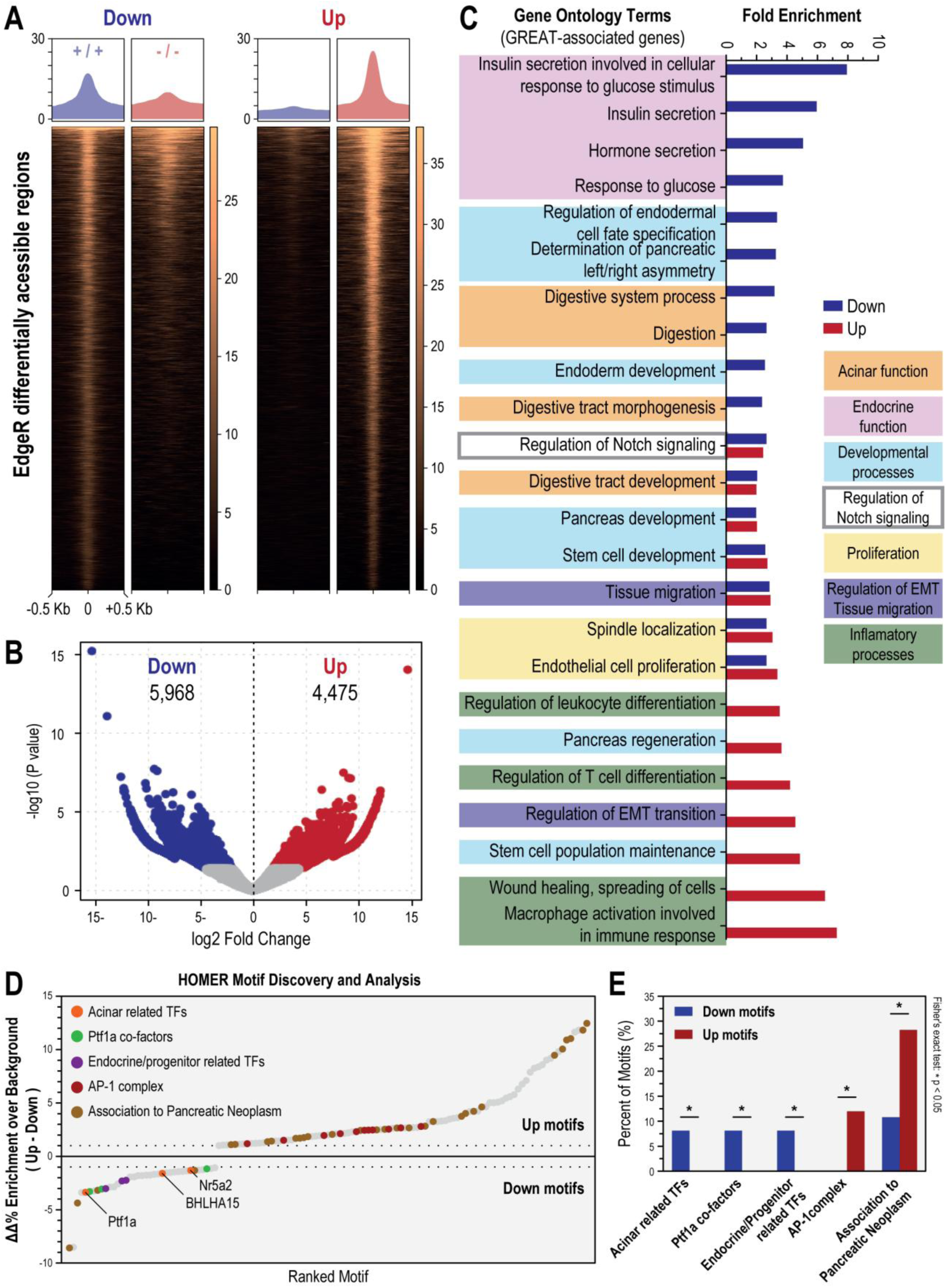
Loss of z3’-DpE leads to genome-wide chromatin remodeling and activation of inflammatory and regenerative programs in acinar cells. **(A)** Average profiles and heatmaps of differentially accessible chromatin regions (DARs) between z3’-DpE+/+ and z3’-DpE-/- acinar cells (adjusted p≤0.05, |log_2_ fold change|≥1); ATAC-seq read density shown ±5 kb around peak summits. **(B)** Volcano plot of DARs in z3’-DpE-/-versus z3’-DpE+/+. **(C)** GO enrichment of genes associated with DARs (GREAT analysis). **(D)** HOMER motif enrichment analysis (ΔΔ% = [ΔUp - ΔDown] over background) showing transcription factor (TF) motifs preferentially enriched in regions gaining or losing accessibility; colored circles denote functionally related TF groups manually curated from the literature. **(E)** Percent of TF motifs within functional groups enriched in Down or Up regions.

Overall, these results suggest that chromatin accessibility changes are locus-specific and reflect a selective rewiring of gene regulatory networks. In this scenario, certain genes within these pathways may acquire a higher regulatory potential while others a decreased potential, resulting in disrupted coordination and functional instability within major developmental and differentiation modules. This pattern is consistent with widespread lineage instability in mutant acinar cells, rather than targeted fate change and supports the idea that acinar cells in z3’-DpE mutants undergo a reconfiguration of differentiation programs, losing identity and adopting heterogeneous regulatory states, a hallmark of metaplasia, regeneration, and preneoplastic processes^35,42–46^.

To further investigate how z3’-DpE loss alters the transcriptional regulatory landscape in acinar cells, we performed motif analysis on differentially accessible regions in z3’-DpE mutants and wild-type acinar cells. Motifs for acinar identity-related TFs, including Ptf1a, Bhlha15, and Nr5a2, were preferentially associated with regions that lost chromatin accessibility (Down TFs), alongside Ptf1a cofactors (e.g., Tcf3, Tcf12, and Tcf4) and endocrine/progenitor-related TFs (e.g., Ascl1, Ascl2, and NeuroD1) (Figure 4D-E). This suggests that z3’-DpE and its regulatory network help sustain a chromatin environment that supports both mature acinar function and residual accessibility at developmental or alternative lineage loci. Loss of z3’-DpE disrupts this coordinated network, weakening regulatory domains required for acinar identity maintenance and reducing accessibility at loci normally poised for lineage flexibility, collectively predisposing cells to identity instability. Conversely, TFs associated with pancreatic neoplasm, such as AP-1 complex components^47–51^, ETS1^52–56^, and FLI1^57,58^, were enriched in Up regions (Up TFs) (Figure 4D-E), suggesting a shift toward increased chromatin accessibility at oncogenic TF binding sites and greater potential responsiveness to pro-tumorigenic signals.

One possible mechanism underlying reduced chromatin accessibility at regions enriched for motifs of Down TFs in the z3’-DpE mutant background is altered chromatin accessibility at the loci encoding these TFs themselves. Such locus-specific accessibility changes may lead to transcriptional dysregulation of these TF genes, reducing TF availability at their binding sites genome wide and thereby contributing to local chromatin closure. Consistent with this model, although no global differences in accessibility were detected across TF-encoding loci, a subset of Down TF genes critical for acinar function, including Ptf1a co-factors, exhibited reduced chromatin accessibility across their loci in mutant acinar cells, including *ptf1a*, *bhlha15*, and *rbpjb* (Supplementary Figure 6A-B). Similarly, some UP TFs encoding genes were observed to contain predominantly open regions in their genomic landscapes (Supplementary Figure 6A-B).

Another possible mechanism underlying the widespread chromatin remodeling observed in z3’-DpE mutant acinar cells could derive indirectly from the depletion of Ptf1a, if Ptf1a co-binds cooperatively with other TFs to establish and maintain acinar-specific regulatory landscapes. To test whether Down regions containing predicted Ptf1a binding sites are preferentially enriched for additional TF motifs, we performed a motif co-occurrence analysis. Down regions were stratified into Ptf1a-positive and Ptf1a-negative subsets based on predicted Ptf1a sites, and motifs were assessed for enrichment in Ptf1a-positive regions relative to Ptf1a-negative regions. To avoid false co-occurrence due to similarity with the Ptf1a motif, candidate motifs were filtered based on overlap with predicted Ptf1a sites (Supplementary Figure 7A-C). We identified seven motifs that were significantly enriched in Ptf1a-positive Down regions and exhibited ≥ 2-fold enrichment relative to Ptf1a-negative Down regions (Supplementary Figure 7B); Nr5a2 (also identified among Down TFs), Pbx3, MafB, ZNF416, GATA:SCL, MITF, and Usf2. Among these, Nr5a2 (Nuclear Receptor Subfamily 5 Group A Member 2), like Ptf1a, is an established regulator of acinar identity^59^ is considered a pancreatic cancer susceptibility gene in humans^60^. Pbx3 is PBX family TALE homeodomain factor. Members of this family have documented roles in exocrine transcriptional regulation. Notably, related TALE family member PBX1b, form heterotrimeric complexes with PDX1 and MEIS2 at the ELA1 (elastase I) enhancer to activate exocrine gene expression^61,62^.

Though the contributions to acinar homeostasis and lineage stability of the five additional co-enriched motifs (MafB, ZNF416, GATA:SCL, MITF, and Usf2) remain to be elucidated, these findings suggest that reduced Ptf1a dosage may destabilize a subset of Down regions that integrate multiple acinar TF inputs, potentially contributing to coordinated chromatin closure at acinar regulatory modules.

Consistent with a potential transition toward a tumor-permissive state, prior studies in mice have shown that Ptf1a loss induces acinar-to-ductal metaplasia (ADM) and a KRAS-permissive, pre-neoplastic transcriptional profile^12,23^. To assess whether the chromatin remodeling in z3’-DpE-/- acinar cells reflects conserved pathways associated with PDAC susceptibility, we compared the genes associated with differentially accessible regions in z3’-DpE-/-acinar cells to transcriptionally dysregulated genes in Nr5a2+/-mouse pancreas, a model of PDAC predisposition which displays a pre-inflammatory pancreatic state^63^. Using human orthologs for cross-species comparison (Supplementary Figure 8A), hypergeometric testing revealed significant enrichment of genes associated with increased chromatin accessibility in z3’-DpE-/- acinar cells among those upregulated in Nr5a2+/- pancreas (p ≤ 0.0001, Supplementary Figure 8B), suggesting convergence on inflammatory and epithelial plasticity pathways across species.

These findings demonstrate that z3’-DpE safeguards acinar cell identity by sustaining open chromatin at acinar and developmental regulatory elements while limiting accessibility at loci involved in proliferation, inflammation, and pancreatic cancer. Loss of this enhancer precipitates broad chromatin remodeling that potentially compromises lineage stability, and causes a shift toward a tumor-permissive transcriptional states, which aligns with the role of Ptf1a both as a regulator of pancreatic acinar cell identity^8,12,13^ and as a tumor suppressor^23,24^.

### z3’-DpE suppresses acinar-to-ductal metaplasia and early neoplastic transformation

Our chromatin accessibility analysis of z3’-DpE-/- acinar cells revealed increased accessibility in regions associated with genes involved in pancreatic cancer, aligning with the established role of Ptf1a as a tumor suppressor^23,24^. To determine whether the loss of z3’-DpE predisposes zebrafish to pancreatic neoplasia, we performed histological analysis of pancreata from adult (≥1 year) z3’-DpE-/- and z3’-DpE+/+ zebrafish. In wild-type animals, hematoxylin and eosin (H&E) staining revealed a characteristic exocrine pancreas, with densely packed acinar cells exhibiting basophilic basal cytoplasm and abundant eosinophilic apical zymogen granules (Figure 5A, panel a). Duct cells were sparse and distinguished by their lighter staining pattern with a higher nuclear-to-cytoplasmic ratio, forming structures with characteristic ductal morphology (Figure 5A, panel a). In contrast, pancreata from some z3’-DpE-/- mutants displayed widespread decrease of acinar tissue alongside the appearance of multiple irregularly arranged duct-like structures (larger, rounded tubular structures with a clearly visible central lumen), consistent with ADM. ADM is considered an early histological event in pancreatic neoplasia, often preceding pancreatic intraepithelial neoplasia (PanIN) and PDAC^64–66^. Moreover, these lesions exhibited a partially disorganized morphology that still resembled ductal structures and were frequently surrounded by lighter-staining stromal areas containing elongated, fibroblast-like nuclei (Figure 5A, panel c) indicative of early stromal activation or fibrotic remodeling. This fibroblast-rich stroma resembles desmoplasia, characterized by dense extracellular-matrix deposition and fibroblast proliferation, commonly seen in precancerous and neoplastic pancreatic lesions in mammals^67–69^. In zebrafish, a similar though milder fibroblastic response has also been reported around acinar lesions, particularly in injury or *KRAS*-driven models^27,70^.

**Figure 5.**
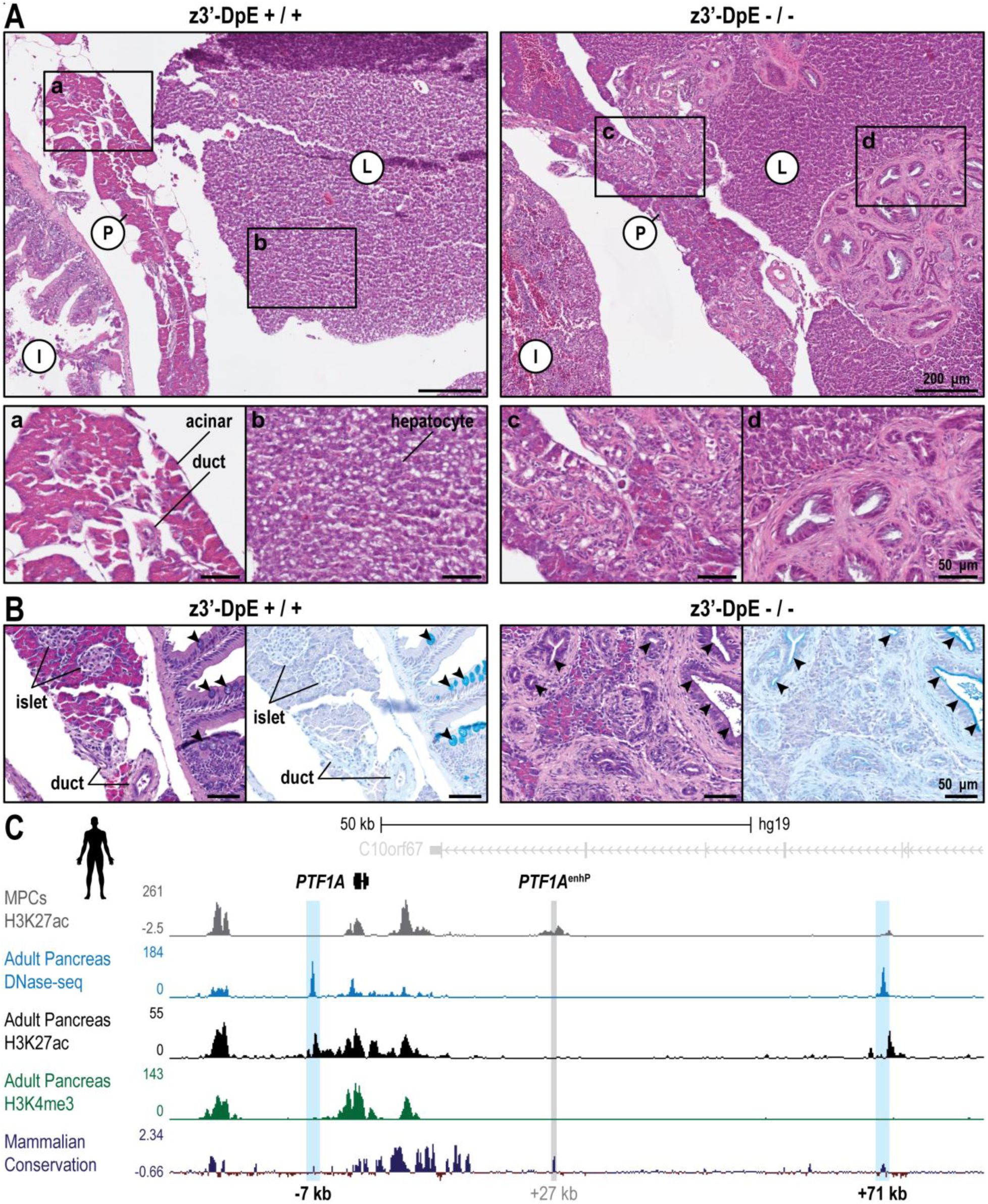
The zebrafish z3’-DpE reveals a potential PDAC susceptibility hotspot in the human *PTF1A* locus. **(A)** Representative hematoxylin and eosin (H&E) stained sections of adult pancreas and liver. In z3’-DpE+/+ (left) the exocrine pancreas consists of closely packed acinar cells, with abundant, granular, and strongly eosinophilic cytoplasm; the interspersed ducts display paler, lightly eosinophilic cytoplasm, higher nuclear-cytoplasmic ratio, and more uniform epithelial linings (a). Hepatocytes display polygonal morphology, uniform size, prominent central nuclei and homogeneous cytoplasmic eosin staining (b). In z3’-DpE-/- (right), the pancreas shows partial loss of acinar tissue with expansion of large, irregular, duct-like structures, distinguished by increased nuclear density, enlarged lumina, and surrounding eosinophilic fibrotic stroma (c); occasional invasion or ductal outgrowths extend into the adjacent hepatic parenchyma (d). **(B)** H&E and Alcian Blue staining showing acidic mucin accumulation (arrowheads) within atypical ductal structures in z3’-DpE-/- mutants. **(C)** The human *PTF1A* locus showing DNase-seq and ChIP-seq tracks for H3K27ac and H3K4me3 in adult pancreas, highlighting sites of regulatory activity. Candidate enhancers active in adult pancreas are shaded blue; the MPC-specific enhancer (PTF1A^enhP^)^6^ is indicated by the central grey shaded region; below, mammalian conservation is shown. L, liver; P, pancreas; I, anterior intestinal bulb.

In z3’-DpE-/- mutants, duct-like structures embedded in a fibrotic stroma were also detected within the liver parenchyma, adjacent to the pancreas (Figure 5A, panel d). These extra-pancreatic lesions are suggestive of invasive outgrowths from neoplastic pancreatic tissue. Alcian blue staining highlighted the emergence of mucin-producing ductal cells in z3’-DpE-/- pancreata, contrasting with the sparse, Alcian blue-negative ducts in wild types (Figure 5B). Alcian blue specifically stains acidic, gastric-type mucins, which are not typically produced by normal pancreatic tissue. Their presence is a recognized feature of metaplastic ductal lesions, including ADM, PanIN, and early pancreatic cancer^71–74^.

The combined H&E and Alcian blue findings robustly support an ADM phenotype, mucinous metaplasia, and early preneoplastic transformation, highlighting profound changes in both cell identity and tissue homeostasis resulting from the z3’-DpE enhancer loss. Notably, the histopathological features observed in z3’-DpE-/- mutants closely mirror the pancreatic lesions described in the *ptf1a:eGFP-KRAS^G12V^* zebrafish model of pancreatic cancer^27^. In that model, oncogenic KRAS disrupts acinar differentiation, leading to expansion of undifferentiated progenitors and aggressive, PDAC-like tumors, frequently resulting in invasion of the liver, gut, and ovary^27^. Together, these results demonstrate that z3’-DpE plays a critical role in safeguarding acinar cell fate and suppressing pathological plasticity. Its loss disrupts exocrine homeostasis, leading to widespread ADM and hallmarks of early neoplasia. The close phenotypic parallels with KRAS-driven oncogenesis emphasize Ptf1a’s central function in antagonizing tumorigenic reprogramming, likely through maintenance of a restrictive, acinar-specific chromatin environment.

## Discussion

In contrast to classical *ptf1a* null mutants, in which exocrine loss is uniform and fully penetrant with a consistent failure to form acinar tissue^10,28^, deletion of the z3’-DpE enhancer produces a graded spectrum of pancreatic phenotypes with variable reduction in acinar cell number, area or volume. This hypomorphic phenotype provides a powerful model to examine how suboptimal expression of the master regulator *ptf1a*, due to a cis-regulatory mutation, destabilizes cell identity and activates disease-associated programs, including repair, inflammatory, and EMT pathways. The graded nature of the observed phenotype closely mirrors the variable penetrance typically associated with non-coding mutations in human disease^6,75–77^ and supports the idea that hypomorphic regulatory alleles place gene-regulatory networks near critical thresholds. In this context, phenotypic outcomes are likely shaped by a combination of deterministic effects imposed by genetic background and network dosage, together with probabilistic fluctuations arising from stochastic transcriptional dynamics.

Our findings closely parallel *Ptf1a* enhancer functions described in mammals during early pancreas development. In humans, loss-of-function mutations or deletions of the main *PTF1A* MPC enhancer cause pancreatic agenesis, impaired pancreatic growth, and neonatal diabetes without affecting neural expression^6^. Similarly, mouse embryos lacking the orthologous enhancer display pancreatic hypoplasia and defective progenitor expansion^20^, mirroring the early developmental defects observed in zebrafish z3’-DpE-/-mutants. Furthermore, during early zebrafish development, z3’-DpE-/- MPCs prematurely commit to the acinar lineage at the expense of progenitor renewal. This is in agreement with the observed downregulation of Notch signaling, a canonical trigger for exit of progenitors from the undifferentiated state, accelerated differentiation and depletion of the pancreatic progenitor pool^78–83^. Consistently, Miguel-Escalada and colleagues^20^ provided direct experimental support for the downregulation of Notch pathway components in mouse *Ptf1a* enhancer mutant MPCs. Both systems provide strong evidence that enhancer-driven regulation of the Notch pathway is evolutionarily conserved as a mechanism for maintaining the MPC pool during early pancreas development. Loss of this regulation results in premature depletion of MPCs. Nevertheless, there is a notable difference between the zebrafish z3’-DpE enhancer and the corresponding enhancers in mouse and human in their activity profile during later development. In zebrafish, z3’-DpE maintains activity post-differentiation in mature acinar cells, consistent with endogenous *ptf1a* expression^10,26,27^. Conversely, the activity of the analogous mouse and human enhancers is predominantly restricted to MPCs^6,20^. Despite this divergence, functional analogy between the elements during early pancreas development is clear.

The deeply conserved role of *Ptf1a* in pancreatic development, lineage specification, and maintenance of acinar identity across vertebrates^18^ suggests that the regulatory elements governing its expression in MPCs and acinar cells are likewise functionally conserved, even if their nucleotide sequences and genomic positions have diverged. Consistent with this view, developmental cis-regulatory elements often display conserved activity across species despite extensive sequence divergence^84–89^. Several human and zebrafish enhancers with no detectable sequence similarity drive highly similar tissue-specific expression^22,90–94^. Moreover, comparative studies of other developmental loci, such as the *Drosophila* genes *eve*, *sna*, *krüppel*, and *svb*^95–98^, have proposed that shadow enhancers (partially redundant elements with overlapping activity) and enhancer turnover (the evolutionary gain and loss of cis-elements at a locus) can facilitate such regulatory flexibility, maintaining conserved transcriptional outputs despite rapid sequence evolution^99^. Through such mechanisms, the zebrafish and mammalian *Ptf1a* enhancers can exert comparable control over MPC programs, both driving expression during early pancreatic development and supporting progenitor expansion and acinar fate commitment, even though their underlying sequences are unrelated. Although the configuration of the ancestral *Ptf1a* regulatory landscape is currently unknown, its MPC- and acinar-specific functions are likely to have been preserved throughout vertebrate evolution. It is possible that an ancestral enhancer with combined MPC- and acinar-specific activity was retained in zebrafish but partitioned into distinct elements in mammals. Alternatively, multiple cell-type-specific enhancers may have existed early on and were differentially maintained in each lineage. Regardless of the ancestral organization, the conservation of both regulatory functions highlights the strong selective pressure to maintain *Ptf1a* expression in these two cellular contexts.

Given that the activity of the zebrafish *ptf1a* z3’-DpE enhancer spans both progenitors and mature acinar cells, its loss is expected to produce broader effects than those seen in mammalian enhancer deletions restricted to early stages. At post-embryonic stages, the z3’-DpE-/- mutants exhibited pronounced defects in the exocrine pancreas, most evident as reduction of acinar tissue and disruption of overall tissue architecture. Quantitative and imaging analyses revealed a significant depletion of acinar cells and a concurrent expansion of non-acinar cell populations, particularly pancreatic ducts, leading to irregular and fragmented organ morphology in adult mutants. The decline in acinar tissue in z3’-DpE-/- mutants recapitulates the exocrine loss seen in complete *Ptf1a*/*ptf1a* null mutants in both mouse and zebrafish. In these models, loss of Ptf1a activity leads to complete absence of acinar differentiation^10,28,100^, with Ptf1a-deficient MPCs diverted toward duodenal and bile-duct fates in mice^8,100,101^, or toward the gallbladder and other foregut derivatives in zebrafish^28^. Additionally, ablation of early *Ptf1a*/*ptf1a* expression in both species results in underdeveloped or absent endocrine islets and severe pancreatic duct reduction, reflecting MPC depletion before robust contribution to these lineages^8,10,100^. By contrast, deletion of z3’-DpE results in selective acinar loss with expansion of ductal and endocrine tissues. Likely, the expansion of non-acinar fates occurs as a consequence of the inability to activate the proper levels of *ptf1a* expression required for the acquisition of acinar identity. Moreover, the extent of non-acinar expansion exceeds the loss of acinar tissue, suggesting that the phenotype reflects not only biased fate allocation but also an expansion of late progenitors capable of generating ductal and endocrine lineages. This is consistent with the known progenitor potential of the ductal compartment, which harbors facultative progenitors capable of generating both endocrine and acinar lineages after injury or developmental perturbation^29,102,103^. Expansion of the ductal lineage at the expense of acinar cells may therefore increase the late progenitor pool, enabling generation of additional ductal and endocrine cells. Indeed, while speculative, such tissue-level responses are consistent with previous observations where selective ablation of acinar cells in zebrafish or mouse triggers adaptive expansion not only of new acinar cells but also of duct and beta-cells^103,104^, and residual progenitors or duct cells can proliferate and give rise to multiple pancreatic lineages, including beta-cells, during regeneration or after developmental perturbation^29,105,106^. This response may also explain why, while in mouse and humans the adult pancreas is overall reduced^6,20^, in zebrafish overall the pancreatic volume is increased.

Pancreatic differentiated cells retain subsets of regulatory elements in a poised, partially accessible chromatin state^107,108^. In acinar cells, this residual accessibility is preferentially associated with developmental and alternative lineage genes, albeit at lower levels than in progenitors or endocrine cells^108,109^, indicating that the acinar chromatin landscape remains permissive for transcriptional reprogramming. Consistent with a hypomorphic state, some z3’-DpE-deleted cells still differentiate as acinar. However, their regulatory programs show coordinated suppression of both acinar and endocrine regulatory signatures, disrupting the chromatin architecture that normally stabilizes acinar identity while retaining controlled plasticity^13,36,110–112^. Such lineage ambiguity increases susceptibility to metaplasia and tumorigenesis by rendering chromatin vulnerable to aberrant activation of oncogenic programs^113,114^, thereby predisposing cells to ADM and progression toward PanIN and PDAC^36,110,115^. Accordingly, z3’-DpE-/- acinar cells exhibit selective chromatin opening at loci linked to inflammation, proliferation, EMT, and neoplasia, consistent with maladaptive dedifferentiation and early neoplastic transformation. This enhancer-driven chromatin remodeling and lineage instability aligns with emerging models of metaplasia and preneoplastic progression^35,42–46^. Histologically, mutant z3’-DpE-/- pancreata show loss of compact acinar parenchyma, with irregular acinar clusters embedded in duct-like, mucinous, fibrotic structures resembling early PanIN lesions. These lesions recapitulate pathology seen in mouse models with conditional *Ptf1a* impairment in adult acinar cells^13,111^ and in zebrafish expressing oncogenic *KRAS^G12V^*, which display compromised acinar differentiation, mucinous ADM, and PanIN-like lesions that progress to carcinoma^27,70^. Although z3’-DpE loss is not oncogenic per se, it disrupts the same *ptf1a*-dependent transcriptional axis that counteracts KRAS-driven reprogramming, underscoring that defective lineage stabilization is a rate-limiting step in ADM and neoplastic susceptibility^13,42,46,116^.

Loss of a *ptf1a* enhancer with late acinar-restricted activity in zebrafish destabilizes acinar identity and promotes ADM and neoplastic progression, suggesting that a similar regulatory vulnerability may exist in humans. Motivated by these findings, we interrogated human chromatin datasets from MPCs and adult pancreas^20,117^ and identified three putative *PTF1A* regulatory regions: the known progenitor-specific enhancer^6^ (PTF1A^enhP^; +27 kb) and two adult-specific candidates located -7 kb upstream and +71 kb downstream of the gene (Figure 5C). The latter two display strong DNase hypersensitivity and H3K27ac in adult pancreas but not in pancreatic progenitors (Figure 5C), suggesting potential roles in maintaining *PTF1A* expression and acinar identity post-differentiation, analogous to the zebrafish z3’-DpE. Mutations in these elements may constitute regulatory vulnerability points that predispose to ADM, PanIN, and PDAC.

Together, our data identify z3’-DpE as a key regulatory safeguard of acinar lineage stability. Its loss destabilizes chromatin architecture, promotes duct-like remodeling and mucinous preneoplasia, and creates a permissive environment for neoplastic transformation. The evolutionary conservation of PTF1A-dependent programs suggests that analogous late-acting enhancers in humans may constitute previously unrecognized susceptibility loci for pancreatic disease and cancer.

## Materials and Methods

### Zebrafish husbandry and ethical approval

Zebrafish (*Danio rerio*) were handled in accordance with European and Portuguese regulations and standard husbandry protocols. The animal facility at i3S is licensed by the Portuguese Directorate-General for Food and Veterinary (DGAV) and accredited by AAALAC International (June 2018). It complies with Portuguese legislation (Portaria 1005/02 and Portaria 1131/97) and the European Directive 2010/63/EU and follows the FELASA (Federation of European Laboratory Animal Science Associations) guidelines and recommendations concerning laboratory animal welfare. All procedures were performed under DGAV project authorization (reference: 0421/000/000/2016).

Adult zebrafish were maintained in a recirculating aquatic system under a 14 h light/10 h dark cycle at 28°C. Water quality was continuously monitored, including pH (∼7) and salinity (∼900 μS). Embryos were obtained from natural matings set up as 3 females: 2 males or 1 female: 1 male, depending on experimental needs. Embryos were grown at 28°C in E3 medium or E3 supplemented with 0.003% 1-phenyl-2-thiourea (PTU; P7629, Sigma-Aldrich). E3 medium consisted of 5 mM NaCl (MB15901, NZYtech), 0.17 mM KCl (2676.298, VWR), 0.33 mM CaCl_2_·2H_2_O (C3881, Sigma-Aldrich), 0.33 mM MgSO_4_·7H_2_O (63140, Sigma-Aldrich) and 0.00015% methylene blue (66720, Sigma-Aldrich), pH ∼7. Embryos/larvae between 33 hpf and 13 dpf were used for immunohistochemistry and for the preparation of material for RNA-based assays and chromatin accessibility profiling, as indicated in the respective sections. For ATAC-seq from acinar cells, whole pancreata were dissected from adult zebrafish aged 8-16 months.

Euthanasia of adult fish was performed by overdose of tricaine methanesulfonate (MS-222; E10521, Sigma-Aldrich; 300 mg/L) in system water for 15 min, followed by decapitation. Embryos and larvae were euthanized by rapid chilling (ice-cold water immersion) or by MS-222 overdose (≥ 72 hpf), in accordance with institutional guidelines.

The following zebrafish lines were used in this study, previously established in our laboratory^22,118^: *Tg(ins:GFP)*, *Tg(sst:mCherry)*, *Tg(ela:mCherry)*, and *Tg(z3’-DpE:GFP)*. The z3’-DpE enhancer deletion allele was maintained as heterozygotes (z3’-DpE+/-) and experiments used z3’-DpE-/- animals and wild-type/heterozygous siblings obtained from incrosses of z3’-DpE+/- adults, unless otherwise specified. All lines were maintained on our laboratory Tübingen (TU) background. Embryos were staged by hours post-fertilization (hpf) and days post-fertilization (dpf) using standard morphological criteria. Sex was not determined for embryos/larvae; adult experiments used animals of both sexes unless otherwise specified.

### Genotyping

Genomic DNA was extracted from adult fin clips, larval tail clips, or whole embryos/larvae using Chelex-100 resin (C7901, Sigma-Aldrich). Tissue was incubated in 5% Chelex suspension supplemented with 1 mg/mL Proteinase K (GE010.0100, GRiSP) at 56°C for 3 h, followed by 95°C for 10 min. Samples were briefly centrifuged, and the supernatant was used directly as PCR template. PCR genotyping was performed using primers flanking the enhancer region: forward primer 5’-GCCTTTTCCAGTGCAGAGT-3’ and reverse primer 5’-GGAGTAGTGTTAGTTGTGGGC-3’. PCR amplification was performed using NZYTaq II DNA polymerase (MB354, NZYtech) with the following cycling conditions: 94°C for 3 min; 35 cycles of 94°C for 30 s, 59°C for 30 s, 72°C for 2 min 30 s; followed by a final extension at 72°C for 5 min. Wild-type alleles produce a 1,928 bp amplicon, whereas the z3’-DpE deletion alleles produce a 1,296 bp amplicon. Heterozygous animals were identified by the presence of both bands. Products were resolved on 1% agarose gels stained with GreenSafe Premium (MB13201, NZYtech) and visualized under UV transillumination.

### Embryo and larval immunostaining and confocal imaging

Whole-mount immunohistochemistry was performed as previously described^22^. Embryos were manually dechorionated at 24-48 hpf and fixed in 4% paraformaldehyde (PFA; Sigma-Aldrich) in 1× phosphate-buffered saline (PBS) at 4°C overnight. When necessary, the yolk was removed post-fixation. Larvae were fixed for 1 h at room temperature. PBS consisted of 137 mM NaCl (MB15901, NZYtech), 2.7 mM KCl (2676.298, VWR), 10 mM NaHPO_4_ (Merk), and 1.8 mM KH_2_PO_4_ (Merk). Following fixation, samples were washed in PBS containing 0.1% Triton X-100 (PBS-T, 0.1%), permeabilized in PBS containing 1% Triton X-100 for 2 h, washed again in PBS-T (0.1%), and blocked for 1 h in PBS-T (0.1%) supplemented with 5% bovine serum albumin (BSA; MB04602, NZYtech). Primary antibodies were diluted in blocking solution and incubated overnight at 4°C: mouse anti-rat Nkx6.1 (Developmental Studies Hybridoma Bank, F55A12; 1:50), mouse anti-zebrafish Zn8 (Developmental Studies Hybridoma Bank, ZN-8; 1:50), and rabbit anti-zebrafish insulin (Abcam, ab210560; 1:50). After extensive washing in PBS-T (0.1%), samples were incubated overnight at 4°C with DAPI (D1306, Invitrogen; 1:1000) and secondary antibodies diluted in blocking solution: goat anti-rabbit IgG Alexa Fluor 568 (A-11036, Invitrogen; 1:800), goat anti-mouse IgG Alexa Fluor 647 (A-21236, Invitrogen; 1:800). Samples were washed extensively in PBS-T (0.1%) and mounted in 50% glycerol in PBS. Confocal imaging was performed on a Leica TCS SP5 II confocal microscope using a 40× water-immersion objective. Z-stacks were acquired with a 1.5 µm (embryos) or 1.0 µm (larvae) step size, and imaging settings were kept constant within each experiment when comparing genotypes and/or developmental stages. Image processing and quantitative analyses were performed using Fiji/ImageJ.

### Histology and mucin staining

Adult zebrafish were euthanized as described above (six adults per genotype), and the abdominal cavity was opened to dissect the gastrointestinal tract region containing the pancreas. Dissected tissues were fixed in formalin for 1 h at room temperature and submitted to the Histology and Electron Microscopy Scientific Platform at i3S for processing, paraffin embedding, and microtome sectioning according to standard procedures. Serial paraffin sections were cut at 5 µm thickness. Hematoxylin and eosin (H&E) staining and Alcian Blue staining were performed by the Histology and Electron Microscopy Scientific Platform, with the assistance of Rossana Correia, using standard protocols. Brightfield whole-slide digital scanning was performed by the Platform, and representative images were exported for figure preparation and downstream analysis.

### Micro-CT imaging and pancreas morphometrics

Whole zebrafish bodies were contrast-stained with phosphomolybdic acid (PMA, hydrate; Sigma-Aldrich, 221856) by incubation for 6 days in 10 mL of 2.5% PMA in type I water at 4°C, in a 15 mL conical tube under agitation. Samples were briefly rinsed with deionized water and mounted in water-soaked foam to prevent dehydration and sample displacement within a 5 mL tube (Sarstedt, 62.558.201), sealed with Parafilm. To minimize staining fading and loss of staining homogeneity, scans were performed using short acquisition settings. Samples were scanned using a SkyScan 1276 micro-CT system (Bruker) with the following parameters: 55 kV tube voltage, 72 µA current, 0.25 mm aluminum filter, 6.5 µm voxel size, 0.2° rotation step over 360°, and 1900 ms exposure time. Images were acquired with frame averaging of 4 and an image size of 4032 × 2688 pixels. Three-dimensional reconstruction from X-ray projection images was performed using NRecon (v1.7.5.2; Bruker). Reconstruction settings were kept constant across samples, except for post-alignment parameters adjusted per sample, using smoothing = 0, ring artifact correction = 1, and beam hardening correction = 15%. DataViewer (v1.5.6.3; Bruker) was used to assess reconstruction quality and to align datasets for downstream analysis. Segmentation was performed in CTAn (CT-Analyser, v1.20.3.0; Bruker) using transverse sections. The pancreas was segmented from the abdominal region based on PMA-enhanced soft-tissue contrast and anatomical landmarks, and morphometric measurements including volume (voxel number) and surface area were extracted in CTAn. Graphs were generated using GraphPad Prism (v10.0.2).

### FACS purification of pancreatic cell populations

Pancreatic cell populations were purified by fluorescence-activated cell sorting (FACS) from adult and embryonic zebrafish reporter lines. For adult acinar preparations used for ATAC-seq, whole pancreata were dissected from 8-16-month-old *Tg(ela:mCherry)* zebrafish and fixed in 4% formaldehyde (Sigma-Aldrich) in PBS. For embryonic pancreatic progenitors, 48 hpf *Tg(z3’-DpE:GFP)* embryos were euthanized by rapid chilling and progenitor cells were released by gentle, repeated pipetting in 300 µL of Ginzburg fish Ringer’s solution; 55 mM NaCl (MB15901, NZYtech), 1.8 mM KCl (2676.298, VWR), 1.25 mM NaHCO_3_, Sigma-Aldrich). Embryos were allowed to settle by gravity and the supernatant containing detached pancreatic progenitor cells and yolk was collected and washed with PBS. For ATAC-seq, embryonic pancreatic progenitors were fixed for 10 min in 4% PFA in PBS prior to dissociation and sorting. Adult pancreatic tissue and embryonic cell preparations were dissociated on ice using a 15 mL Dounce homogenizer in 1 mL of ice-cold sort buffer and passed through a 40 µm cell strainer. Sort buffer consisted of 1× PBS supplemented with 0.5% BSA (MB04602, NZYtech), 25 mM HEPES (H0887, Sigma-Aldrich), and 2 mM EDTA. For RNA-seq of live pancreatic progenitors, sort buffer was additionally supplemented with 0.02 U/µL NZY Ribonuclease Inhibitor (NZYtech, MB08401). Cell sorting was performed by the Translational Cytometry Unit at i3S on a BD Biosciences FACSAria II cell sorter using an 85 µm nozzle. Cells were gated to exclude debris and doublets; for live pancreatic progenitor RNA-seq samples, dead cells were excluded using Fixable Viability Dye eFluor 780 (65-0865, Invitrogen; 1:1000). mCherry+ (adult acinar) or GFP+ (embryonic progenitor) populations were collected in sort buffer (for ATAC-seq) or in TRK lysis buffer (for RNA extraction and RNA-seq).

### ATAC-seq library preparation

Assay for transposase-accessible chromatin with high-throughput sequencing (ATAC-seq) was performed as previously described^119^, with minor modifications, using FACS-purified formaldehyde-fixed pancreatic cell populations. For adult acinar ATAC-seq, fixed mCherry+ acinar cells were collected by FACS (10,000-50,000 cells per replicate), lysed ice-cold lysis buffer (10mM Tris-HCl, pH7.5; 10mM NaCl; 3mM MgCl_2_; 0.1% IGEPAL), and incubated with homemade Tn5 transposase in TAPS-DMF buffer^120^ at 37°C for 30 min. For pancreatic progenitors ATAC-seq, fewer than 5,000 fixed GFP+ pancreatic progenitor cells were collected per replicate. Due to the low input, the tagmentation reaction was scaled down to a total volume of 10 µL^119^. Following transposition, crosslinks were reversed by incubation at 65°C overnight in reverse-crosslinking solution (50 mM Tris-Cl pH 8, 1 mM EDTA pH 8, 1% SDS, 0.2 M NaCl, and 5 ng/µL Proteinase K). DNA was purified using the MinElute PCR Purification Kit (QIAGEN). To minimize PCR bias, the optimal number of amplification cycles was determined by qPCR as the cycle corresponding to approximately half-maximal fluorescence. Libraries were amplified using KAPA HiFi HotStart ReadyMix (Roche) and Illumina/Nextera adapter primers (Ad1 and indexed Ad2 primers), and purified using the QIAquick PCR Purification Kit (QIAGEN).

### RNA-seq library preparation

After collection by flow cytometry, total RNA was extracted from live zebrafish pancreatic progenitor cells using the E.Z.N.A. MicroElute Total RNA Kit (R6831; Omega Bio-tek) according to the manufacturer’s instructions. The optional on-membrane DNase digestion step was performed during extraction using DNase I, RNase-free (EN0521; Thermo Scientific) in its supplied buffer; 30 min at 37°C; 1 µL DNase I and 0.5 µL NZY Ribonuclease Inhibitor (NZYtech, MB08401). RNA was eluted according to the kit protocol and stored at -80°C.

### scRNA-seq of whole adult pancreas

Single-cell RNA sequencing (scRNA-seq) was performed on whole pancreas tissue from adult wild-type and mutant zebrafish siblings (two adults per genotype) using the 10x Genomics Chromium platform and the Chromium Next GEM Single Cell 3’ Reagent Kits v3.1 workflow. Whole pancreata were mechanically dissociated on ice using a 15 mL Dounce homogenizer in 1 mL of ice-cold dissociation buffer and passed through a 40 µm cell strainer. Dissociation buffer consisted of 1× cOmplete, EDTA-free Protease Inhibitor Cocktail (11873580001, Roche) and 0.2 U/µL NZY Ribonuclease Inhibitor (NZYtech, MB08401) in 1× PBS. Cells were stained with Fixable Viability Dye eFluor 780 (65-0865, Invitrogen; 1:1000) prior to sorting. Sort buffer consisted of 1× PBS supplemented with 0.04% BSA (MB04602, NZYtech) and 0.2 U/µL NZY Ribonuclease Inhibitor (NZYtech, MB08401). Live single cells were purified by FACS on a BD Biosciences FACSAria II cell sorter using a 100 µm nozzle, gating to exclude debris, dead cells, and doublets. Approximately 20,000 live cells were collected per sample and immediately processed. Cell sorting was performed by the Translational Cytometry Unit at i3S, and libraries were generated by the i3S Genomics Scientific Platform, with the assistance of Ana Mafalda Rocha, following standard 10x Genomics recommendations.

### Quantification and statistical analysis

Quantifications were performed from independent biological samples; no statistical methods were used to predetermine sample size. Image processing and quantifications were performed in Fiji (ImageJ) using consistent acquisition and analysis settings within each experiment, and measurements were normalized to wild-type siblings within the same batch where indicated. Flow-cytometry quantifications were derived from gated single-cell populations and reported as fractions or normalized values, and micro-CT morphometrics (pancreas surface area and volume) were extracted from 3D segmentation outputs; graphs and statistical analyses for these experiments were performed in GraphPad Prism. Tests were two-sided and significance was defined as p ≤ 0.05; data are shown as mean ± SD unless otherwise indicated. For figures using asterisk notation: *p ≤ 0.05, **p ≤ 0.01, ***p ≤ 0.001, ****p ≤ 0.0001.

### RNA-seq analysis

Total RNA extracted from zebrafish pancreatic progenitors was sequenced on an Illumina HiSeq 2500 platform. Read quality was assessed using FastQC (v.0.11.8)^121^. Reads were aligned to the zebrafish reference genome (GRCz10/danRer10) using Bowtie2 (v.2.4.2)^122^ with the “--very-sensitive” parameter. Gene-level read counting was performed using featureCounts^123^, with default parameters. Differential expression analysis was carried out using DESeq2 (v.1.32)^124^, and genes were considered differentially expressed when adjusted p-value ≤ 0.05 and |log_2_ fold-change| ≥ 1. Gene ontology enrichment analysis was performed using PANTHER GO Enrichment Analysis^125^. For visualization, BAM files from each replicate were normalized using bamCoverage (deepTools v3.5.1)^126^ with the kilobase per million mapped reads (RPKM) method. Signal tracks from replicates were averaged using WiggleTools (v.1.2)^127^ (mean), and the resulting bedGraph files were converted to BigWig format using bedGraphToBigWig (v.377)^128^ for visualization in the UCSC Genome Browser.

### ATAC-seq analysis

ATAC-seq raw read quality was assessed using FastQC (v.0.11.8)^121^. Adapter sequences were trimmed using Skewer (v.0.2.1)^129^, and reads were aligned to the zebrafish reference genome (GRCz10/danRer10) using Bowtie2 (v.2.4.2)^122^. Duplicate mapped reads were removed using SAMtools^130^. Aligned reads were filtered by fragment size (≤120 bp) and mapping quality (MAPQ ≥10). Peak calling was performed using MACS2 (v.2.2.7)^131^ with the following parameters: “--nomodel, --keep-dup 1, --llocal 10000, --extsize 74, --shift – 37 and -p 0.07”. Reproducible peaks across replicates were defined using Irreproducible Discovery Rate (IDR; v2.0.4)^132^. Peaks with an IDR score reflecting a p ≤ 0.1 were retained for downstream analyses. Differential chromatin accessibility was assessed using edgeR^133^, and peaks were considered differentially accessible when adjusted p-value ≤ 0.05 and |log_2_ fold-change| ≥ 1. BAM files from each replicate were normalized to RPKM using bamCoverage (deepTools v3.5.1)^126^. Signal tracks were averaged across replicates using WiggleTools (v.1.2)^127^ (mean), converted from bedGraph to BigWig using bedGraphToBigWig (v.377)^128^, and visualized in the UCSC Genome Browser. Heatmaps and aggregate profiles were generated with deepTools (v.3.5.1)^126^ using computeMatrix (“--referencePoint center, -b 500, -a 500, --binSize 10, --sortRegions descend”) followed by plotHeatmap and plotProfile with default parameters. Genomic intersections were performed using BEDTools (v.2.29.2)^134^. Peak-to-gene association was performed using Genomic Regions Enrichment of Annotations Tool (GREAT)^135^, and GREAT-associated gene lists were subjected to gene ontology enrichment analysis using PANTHER^125^.

### Motif enrichment and TF-grouping strategy

Motif enrichment analysis was performed on differentially accessible peak sets from ATAC-seq of acinar cells using HOMER (v.4.11)^136^, with findMotifsGenome.pl. To compare motif enrichments between conditions, the predicted percentage of sequences containing each enriched motif was extracted from HOMER outputs and contrasted between datasets by subtracting the percentage in one condition from the other using a custom Python script. TFs corresponding to differentially enriched motifs were then manually grouped into functional categories based on curated evidence from the literature (Supplementary Table 6). To annotate TFs linked to pancreatic disease, TF-encoding genes implicated by motif enrichment (Up and Down TF lists) were cross-referenced against human gene-disease associations from DisGeNET (v6.0; all_gene_disease_associations.tsv)^137^ filtered for score > 0.1 and manually curated for pancreas-related terms “Pancreatic Neoplasm” and “Malignant neoplasm of pancreas”.

### Motif co-occurrence analysis

Downregulated peaks were scanned for known Ptf1a binding motifs (HOMER: Panc1-Ptf1a-ChIP-Seq, GSE47459)^138^ using FIMO^139^, with a p-value threshold of p < 10^- 4^. Based on the presence of at least one predicted Ptf1a binding site, downregulated peaks were classified as either Ptf1a-positive or Ptf1a-negative. Motif enrichment analyses were then performed using HOMER (v5.1)^136^ findMotifs.pl in two complementary comparisons. First, motifs enriched across all downregulated peaks were identified using a genome-matched background. Second, motifs specifically enriched in Ptf1a-positive peaks were identified using Ptf1a-negative peaks as a custom background. Candidate co-occurring motifs were identified as those reaching statistical significance in both analyses. To exclude motifs incorrectly identified as co-occurring due to sequence similarity with the Ptf1a motif, we assessed the positional overlap of candidate motif occurrences with predicted Ptf1a binding sites. Motif occurrences were considered overlapping if their intersection with any Ptf1a site exceeded 50% of the shorter motif’s length. Motifs were excluded from further analysis if more than 20% of their occurrences in Ptf1a-positive peaks overlapped with Ptf1a sites. For the remaining candidate motifs, enrichment was quantified as the ratio of motif frequency in Ptf1a-positive versus Ptf1a-negative peaks. Statistical significance was assessed using Fisher’s Exact Test, with p-values adjusted for multiple testing using the Benjamini-Hochberg procedure (FDR < 0.05)^140^.

### scRNA-seq analysis

Alignment of scRNA-seq reads was performed with Cell Ranger (v8.0.1; 10x Genomics) using the *Danio rerio* danRer10 reference transcriptome. For each sample (z3’-DpE+/+ and z3’-DpE-/-), a gene-cell count matrix was generated and imported into R (v4.2.2) for downstream analysis using Seurat (v5.0.1). Cells with fewer than 400 detected features, more than 60,000 UMIs, or > 15% mitochondrial gene expression were excluded. Doublets were identified using scDblFinder (v1.12) applied to SingleCellExperiment objects containing raw counts, and all predicted doublets were removed. After quality control, samples were annotated by condition and merged into a single Seurat object. Data were normalized using LogNormalize (scale factor = 10,000), highly variable genes were identified using the vst method (3,000 features), and the dataset was scaled using ScaleData. Principal component analysis (PCA) was computed using variable genes (100 PCs). Intrinsic dimensionality was estimated using maxLikGlobalDimEst, indicating ∼14 informative components. Based on this estimate and inspection of the elbow plot, the first 25 PCs were retained for downstream analyses. A shared nearest-neighbor graph was constructed using FindNeighbors (k = 20), and clustering was performed using the Leiden algorithm (algorithm = 4). Multiple resolutions (0.1-1.0) were evaluated using UMAP and t-SNE embeddings and silhouette scores, and a final resolution of 0.3 was selected. Dimensionality reduction for visualization was performed using UMAP (uwot implementation) and t-SNE (n.neighbors = 30, seed = 100). Differential expression analysis of each cluster for mutant versus wild-type cells was performed using the MAST test implemented in FindAllMarkers, considering genes expressed in at least 25% of cells in either group, with |log_2_FC| ≥ 1 and adjusted p < 0.05. Differentially expressed genes were filtered and exported using custom R functions, and top markers were visualized with heatmaps and FeaturePlot. Clusters were manually annotated based on canonical marker genes and expression patterns in UMAP and t-SNE embeddings, assigning identities such as stromal, T and NK, blood, and pancreatic IL-2-producing cells. Cell-type composition across z3’-DpE+/+ and z3’-DpE-/- samples was quantified by calculating the relative frequency of each annotated population, and log_2_ fold-changes were computed to compare abundances between conditions. To refine pancreatic populations, a second round of clustering was performed on the pancreatic cell cluster using the same parameters, yielding ∼10 informative components. Manual annotation is based on canonical marker genes and embedding patterns identified subpopulations corresponding to T and NK cells, acinar cells, endocrine cells, macrophages and monocytes, ribosomal cells, ductal cells, and endothelial cells. Cell-type composition across conditions was quantified as relative frequencies, and log_2_ fold-changes were computed to compare abundances between conditions.

## Acknowledgments

We thank all current and past members of the Vertebrate Development and Regeneration group at i3S for their contributions and critical discussions, particularly Fábio Ferreira. We are grateful to Juan Tena and Ana Burgos (CABD - Centro Andaluz de Biología del Desarrollo, Seville, Spain) for support with single-cell RNA-seq analysis. We acknowledge Isabel Guedes for assistance with zebrafish maintenance.

The authors acknowledge the support of the i3S Scientific Platform Advanced Light Microscopy, member of the national infrastructure PPBI - Portuguese Platform of BioImaging (POCI-01-0145-FEDER-022122) and a site of the Euro-BioImaging Node. Cell sorting was performed with the support of the i3S Translational Cytometry (TraCy) Scientific Platform. Histology processing, paraffin embedding and sectioning, H&E and Alcian Blue staining, and slide scanning were performed with the support of the Histology and Electron Microscopy i3S Scientific Platform (Portuguese Roadmap of Research Infrastructures, FCT). Single-cell RNA-seq library preparation was performed with the support of the Genomics i3S Scientific Platform. This work is a result of the GenomePT project (POCI-01-0145-FEDER-022184), supported by COMPETE 2020 - Operational Programme for Competitiveness and Internationalization (POCI), Lisboa Portugal Regional Operational Programme (Lisboa2020), Algarve Portugal Regional Operational Programme (CRESC Algarve2020), under the PORTUGAL 2020 Partnership Agreement, through the European Regional Development Fund (ERDF), and by Fundação para a Ciência e a Tecnologia (FCT).

The laboratory of JB was funded by national funds through FCT through projects COMPETE2030-FEDER-00731800-16163 and UID/4293/2025, and by “la Caixa” Foundation (HR21-01212). JB was supported by the FCT Investigator contract CEECIND/03482/2018. The laboratory of APP was funded by FCT through the project JumpIN (PTDC/BTM-MAT/4156/2021) and acknowledges the MOBILIsE Project, funded by the European Union’s Horizon 2020 research and innovation programme (grant agreement no. 951723). MD (SFRH/BD/135957/2018), JPA (SFRH/BD/145110/2019), and BC (SFRH/BD/145652/2019) were supported by FCT PhD fellowships. MD was also supported by Liga Portuguesa Contra o Cancro - Núcleo Regional do Norte through “Bolsas de Investigação em Oncologia LPCC-NRN 2025” (Bolsa de Iniciação à Investigação).

## Notes

### Competing Interest Statement

The authors have declared no competing interest.

