## Supplementary Figures and Tables for "A dual-phase enhancer couples progenitor maintenance and pancreatic lineage stability"

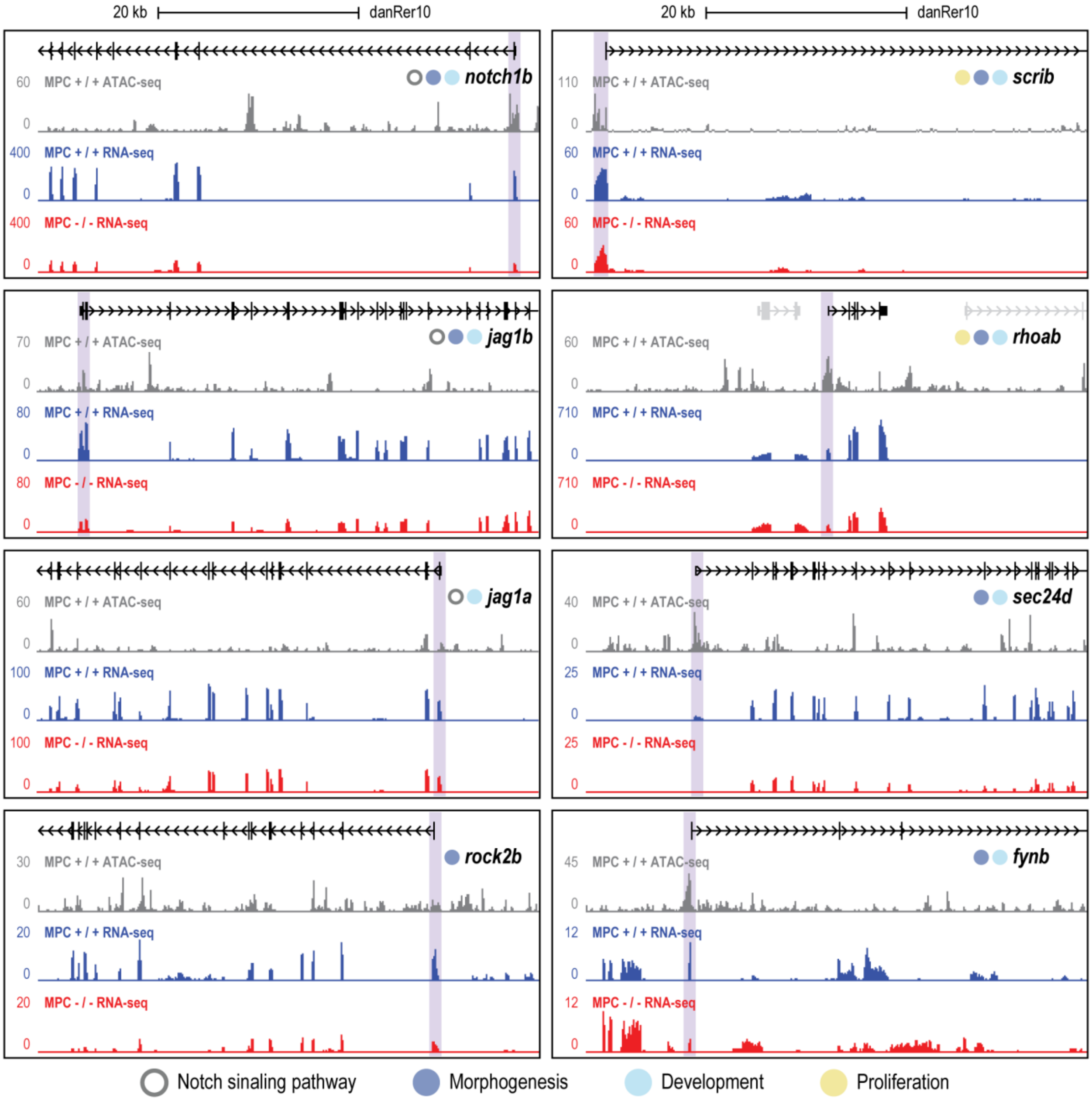

**Supplementary Figure 1. Examples of downregulated genes in z3′-DpE-/- MPCs linked to impaired Notch signaling, morphogenesis, development, and proliferation.**

Genome browser views of representative loci corresponding to genes associated with the GO categories Notch signaling (*notch1b, jag1a, jag1b*), morphogenesis (*scrib*), development (*sec24d, fynb*), and proliferation (*rohab, rock2b*). ATAC-seq read density from 48 hpf z3′-DpE+/+ MPCs (MPC+/+, gray) is shown above together with RNA-seq read density for z3′-DpE+/+ (blue) and z3′-DpE-/- (MPC-/-, red) MPCs below. Blue shaded boxes highlight TSS-proximal regions of downregulated genes in z3′-DpE-/- MPCs.

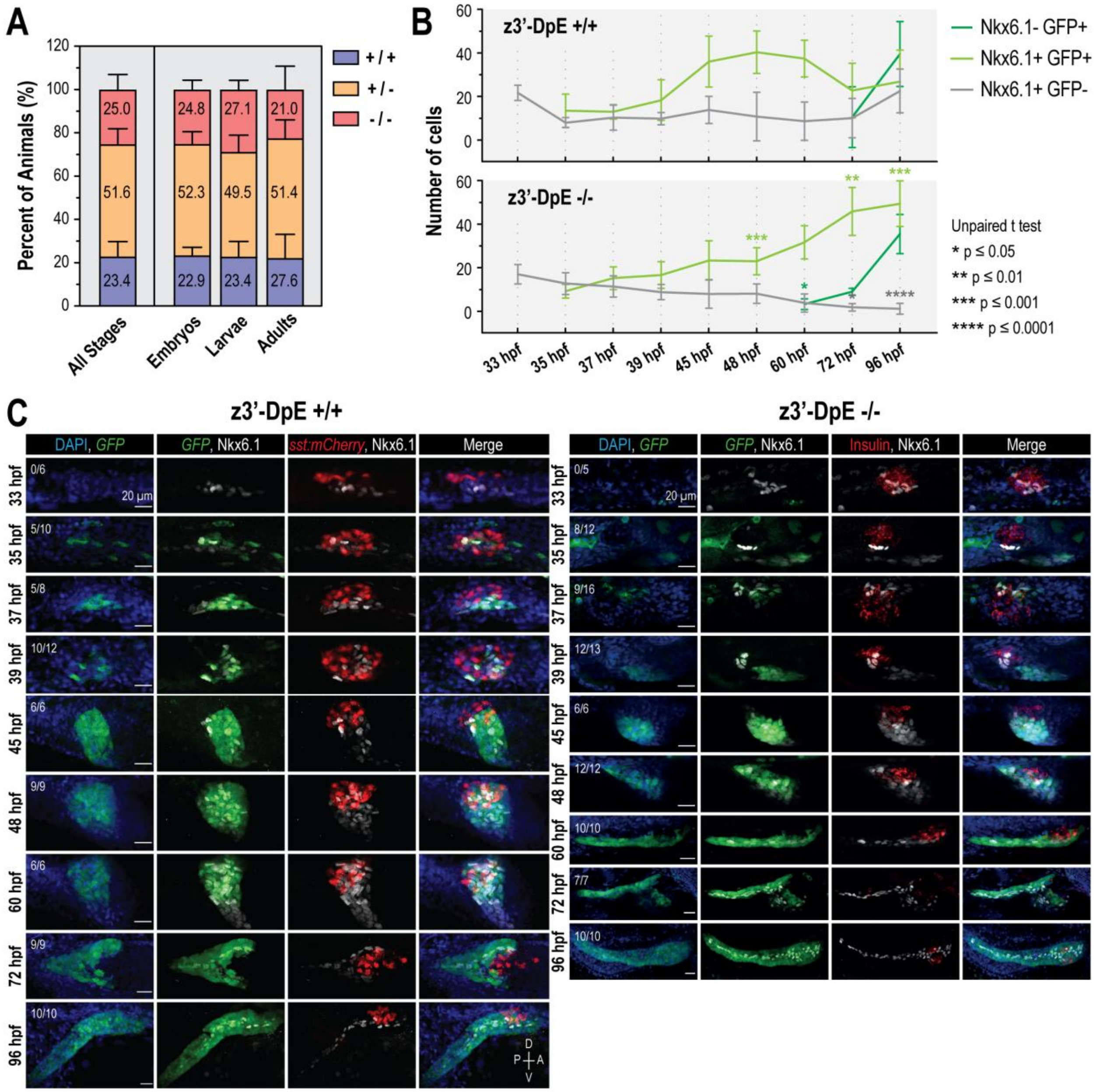

**Supplementary Figure 2. Genotype distribution and MPC composition in z3′-DpE mutants.**

**(A)** Percent distribution of z3′-DpE genotypes obtained from all animals genotyped in this study from heterozygous incrosses (n = 1,303), grouped by developmental stage: Embryos (48-96 hpf; n = 899), Larvae (9-12 dpf; n = 299), and Adults (3-6 months; n = 105). The values displayed inside each bar represent the mean genotype proportions averaged across multiple independent batches (Embryos, 7 batches; Larvae, 5 batches; Adults, 6 batches; minimum of 10 animals per batch). Genotypes were recovered at approximately Mendelian ratios at all stages, indicating that loss of z3′-DpE does not impair viability from embryogenesis to adulthood.

**(B)** Quantification of Nkx6.1+ GFP-, Nkx6.1+ GFP+, and Nkx6.1- GFP+ cell populations in the ventral pancreatic bud of z3′-DpE+/+ (top) and z3′-DpE-/- (bottom) embryos from 33 to 96 hpf (mean ± SD). Embryos were obtained from separate z3′-DpE+/+ and z3′-DpE-/- incrosses.

**(C)** Representative confocal projections showing Nkx6.1+ MPCs (white), z3′-DpE:GFP expression (green), and endocrine cells (sst:mCherry or insulin, red) across developmental stages (33 to 96 hpf) in embryos derived from z3′-DpE+/+ (left) and z3′-DpE-/- (right) incrosses. Numbers indicate the fraction of embryos displaying GFP expression over the total analyzed at each stage. Nuclei are labeled with DAPI (blue). D, dorsal; V, ventral; P, posterior; A, anterior.

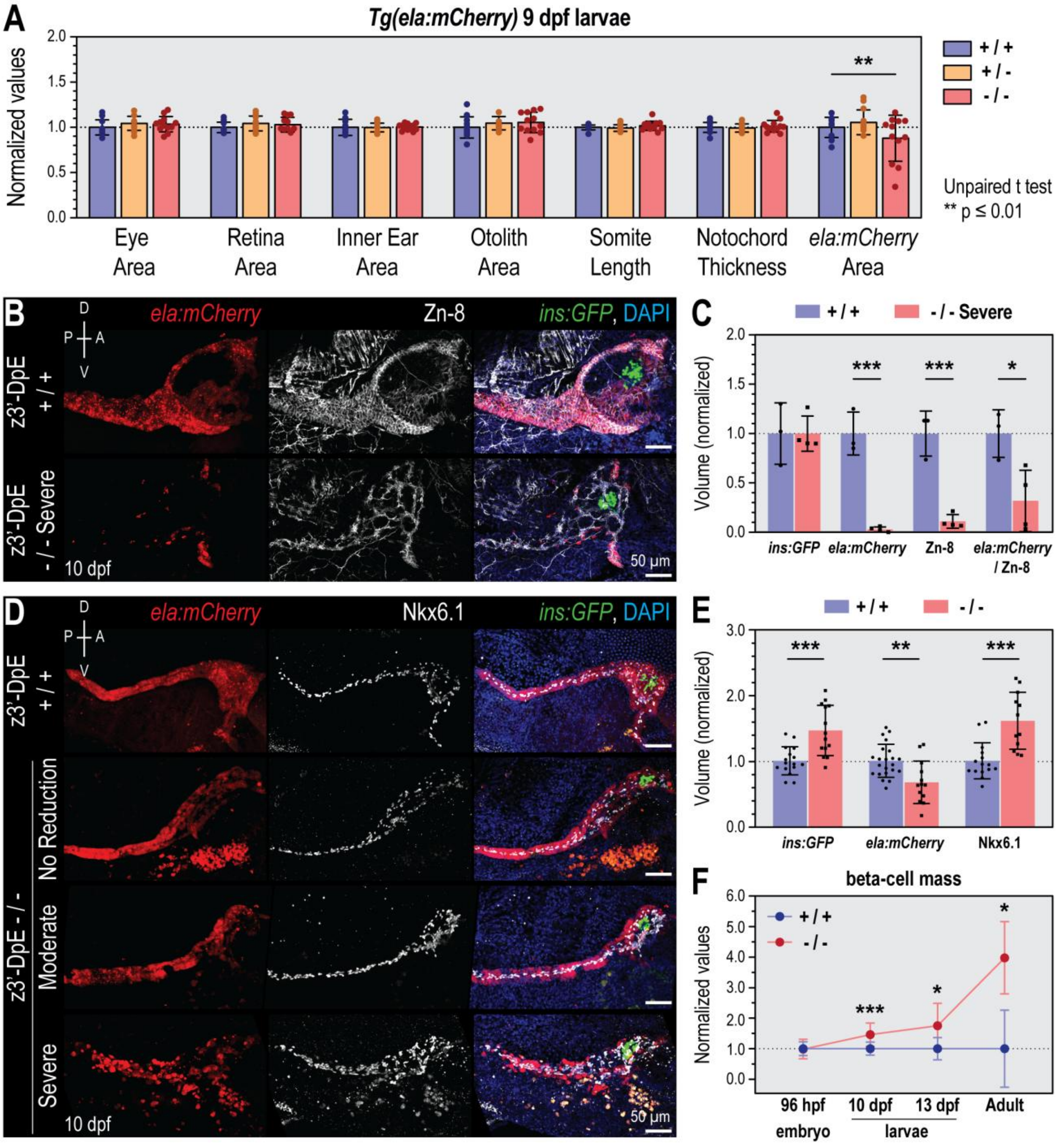

**Supplementary Figure 3. Larval pancreas defects in z3′-DpE mutants.**

**(A)** Morphometric analysis of non-pancreatic anatomical structures (eye area, retina area, inner ear area, otolith area, somite length, and notochord thickness) and *ela:mCherry*-positive pancreatic area in 9 dpf Tg(ela:mCherry) larvae. Measurements are normalized to wild-type siblings (mean ± SD). Sample sizes: z3′-DpE+/+ (n = 12), +/- (n = 13), -/- (n = 12).

**(B)** Representative confocal projections of 10 dpf *Tg(ins:GFP);Tg(ela:mCherry)* z3′-DpE+/+ and severely affected z3′-DpE-/- larvae. Zn-8 (white) marks pancreatic epithelial cells, including both acinar and duct cell populations, and is used here in combination with *ela:mCherry* (red) to distinguish acinar (*mCherry*+ Zn-8+) from duct (*mCherry*- Zn-8+) epithelium. Beta-cells are labeled by *ins:GFP* (green), and nuclei by DAPI (blue).

**(C)** Quantification of *ins:GFP*, *ela:mCherry*, Zn-8-positive epithelial volumes, and the *ela:mCherry*/Zn-8 ratio in z3′-DpE+/+ versus severely affected z3′-DpE-/- larvae at 10 dpf (normalized to wild type; mean ± SD). Sample sizes: z3′-DpE+/+ (n = 3), -/- (n = 5).

**(D)** Representative confocal projections of z3′-DpE+/+ and z3′-DpE-/- larvae at 10 dpf illustrating the range of phenotypic severity (no reduction, moderate, severe). Acinar cells: *ela:mCherry* (red); duct cells: Nkx6.1 (white); beta-cells: *ins:GFP* (green); nuclei: DAPI (blue).

**(E)** Quantification of beta-cell (*ins:GFP*), acinar (*ela:mCherry*), and ductal (Nkx6.1) volumes in z3′-DpE+/+ versus z3′-DpE-/- larvae at 10 dpf (normalized to wild type; mean ± SD). Sample sizes: z3′-DpE+/+ (n = 22), -/- (n = 14).

**(F)** Beta-cell mass across developmental stages (96 hpf, 10 dpf, 13 dpf, adult). For embryos and larvae, beta-cell mass corresponds to total insulin-positive volume; in adults, beta-cell mass corresponds to percentage of *ins:GFP*-positive cells within the pancreatic single cell population (flow cytometry). All values are normalized to wild-type siblings at each stage (mean ± SD). Sample sizes: z3′-DpE+/+ 96 hpf (n = 6), 10 dpf (n = 22), 13 dpf (n = 8), adult (n = 4); z3′-DpE-/- 96 hpf (n = 7), 10 dpf (n = 14), 13 dpf (n = 8), adult (n = 4).

D, dorsal; V, ventral; P, posterior; A, anterior.

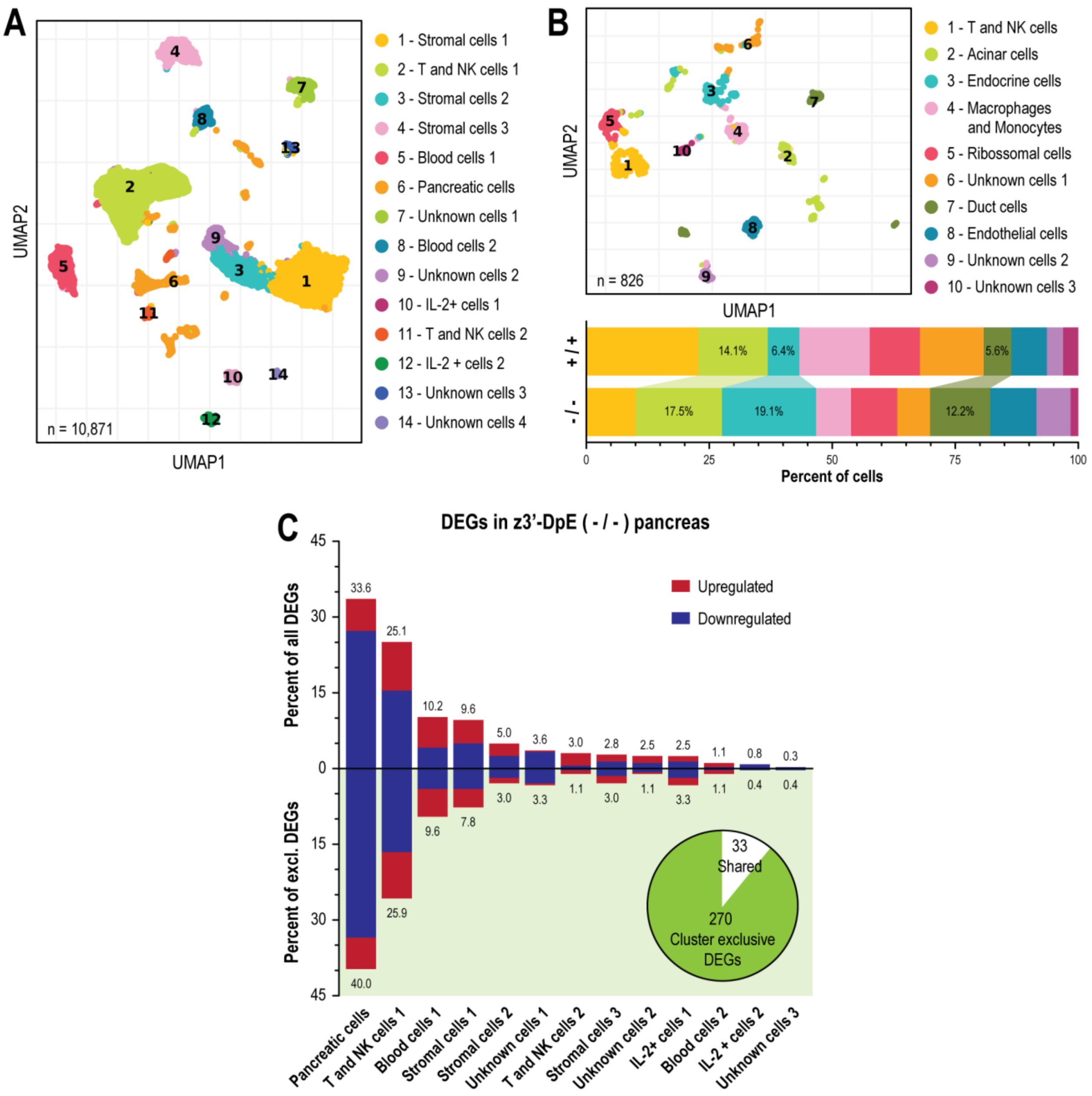

**Supplementary Figure 4. Single-cell transcriptomic profiling of adult z3′-DpE+/+ and z3′-DpE-/- pancreata.**

**(A)** UMAP visualization of the integrated scRNA-seq dataset from whole adult pancreata (n = 10,871 cells), showing 14 transcriptionally distinct clusters annotated as stromal (Clusters 1, 3, and 4), immune (Clusters 2, 5, 8, and 10-12), pancreatic (Cluster 6), and unknown cell populations (Clusters 7, 9, 13, and 14).

**(B)** Sub-clustering of Cluster 6 (pancreatic cells; n = 826 cells) resolves acinar, endocrine, ductal, endothelial, immune, and additional minor cell subpopulations. The stacked bar plots show the relative proportion of each subcluster in z3′-DpE+/+ and z3′-DpE-/- pancreata.

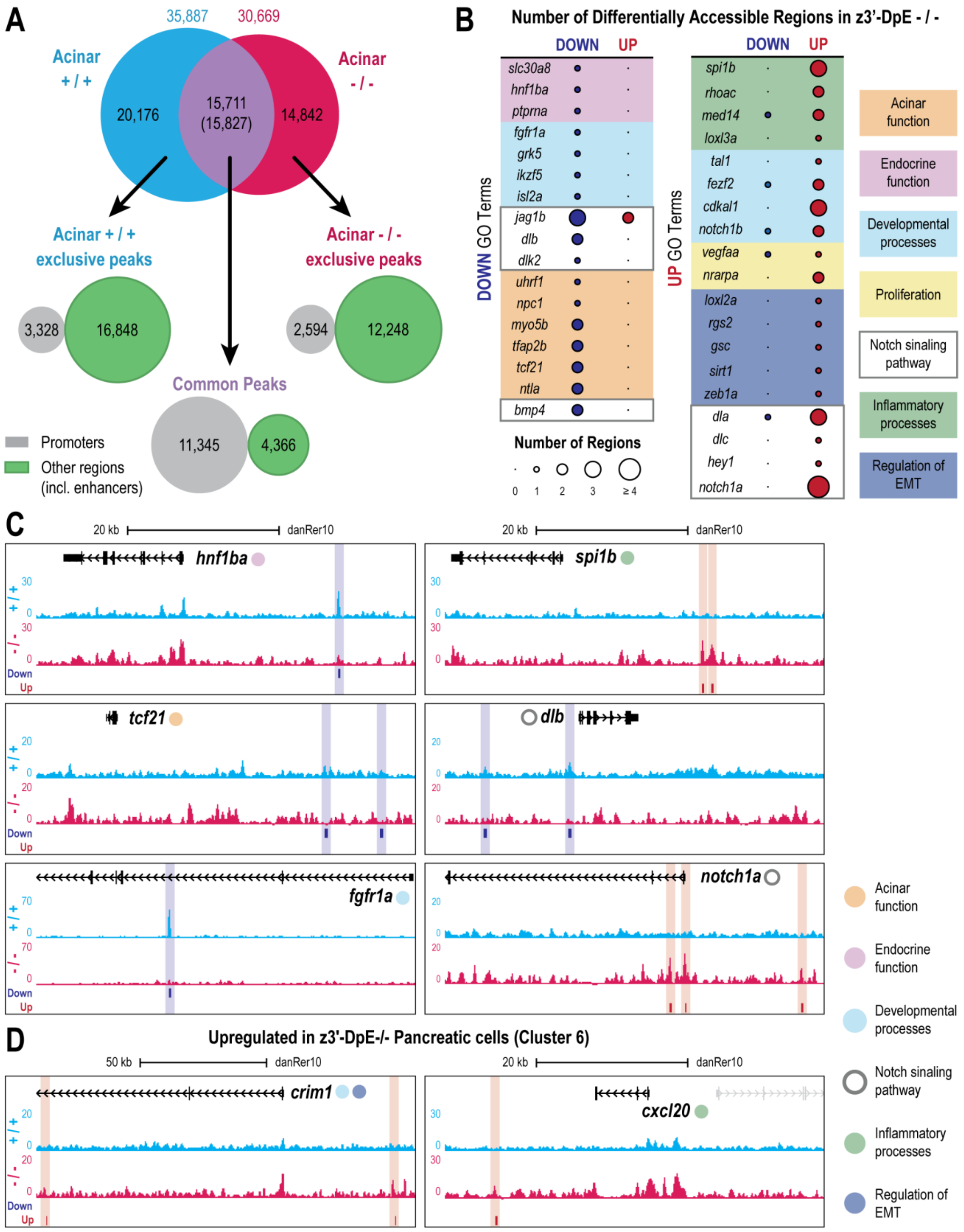
A
35,887
30,669
Acinar + / +
Acinar - / -
20,176
15,711 (15,827)
14,842
Acinar + / + exclusive peaks
Acinar - / - exclusive peaks
3,328
16,848
2,594
12,248
Common Peaks
11,345
4,366
Promoters
Other regions (incl. enhancers)
B
Number of Differentially Accessible Regions in z3'-DpE - / -
DOWN
UP
DOWN GO Terms
UP GO Terms
slc30a8
hnf1ba
ptprna
fgfr1a
grk5
ikzf5
isl2a
jag1b
dlb
dlk2
uhrf1
npc1
myo5b
tfap2b
tcf21
ntla
bmp4
spi1b
rhoac
med14
loxl3a
tal1
fezf2
cdkal1
notch1b
vegfaa
nrarpa
loxl2a
rgs2
gsc
sirt1
zeb1a
dla
dlc
hey1
notch1a
Number of Regions
0
1
2
3
≥ 4
Acinar function
Endocrine function
Developmental processes
Proliferation
Notch sinaling pathway
Inflammatory processes
Regulation of EMT
C
20 kb
danRer10
hnf1ba
spi1b
tcf21
dlb
fgfr1a
notch1a
+ / +
- / -
Down
Up
30
20
70
0
D
Upregulated in z3'-DpE-/- Pancreatic cells (Cluster 6)
50 kb
crim1
cxcl20
Acinar function
Endocrine function
Developmental processes
Notch sinaling pathway
Inflammatory processes
Regulation of EMT

**Supplementary Figure 5. Chromatin accessibility changes in adult z3′-DpE-/- acinar cells.**

**(A)** Overlap analysis of ATAC-seq peaks from FACS-purified acinar cells of adult z3′-DpE+/+ and z3′-DpE-/- pancreata. The Venn diagram shows peaks shared between genotypes and peaks exclusive to z3′-DpE+/+ or z3′-DpE-/-. Each peak set was intersected with annotated transcription start sites (TSSs) to determine the proportion of promoter-proximal (gray circles) and non-promoter regulatory regions (green circles), including putative enhancers.

**(B)** Representative genes associated with differentially accessible regions (DARs) within the selected GO terms (acinar function, endocrine function, developmental processes, Notch signaling, inflammatory processes, and regulation of EMT). Blue and red circles denote regions with decreased (DOWN) or increased (UP) accessibility in z3′-DpE-/- acinar cells, respectively; circle size reflects the number of DARs linked to each gene.

**(C)** Genome browser views of representative genes with altered accessibility linked to the GO terms highlighted in (B). ATAC-seq tracks from z3′-DpE+/+ (blue) and z3′-DpE-/- (red) samples are shown, with DARs highlighted in blue (Down) and red (Up).

**(D)** Genome browser views of representative genes upregulated in z3′-DpE-/- pancreatic cells (Cluster 6). ATAC-seq tracks from z3′-DpE+/+ (blue) and z3′-DpE-/- (red) acinar cells are shown, with regions of increased accessibility in mutants (Up DARs) highlighted in red. Shaded regions mark DARs associated with *crim1* (developmental/EMT-related) and *cxcl20* (inflammatory), illustrating concordant chromatin opening at stress- and inflammation-associated loci.

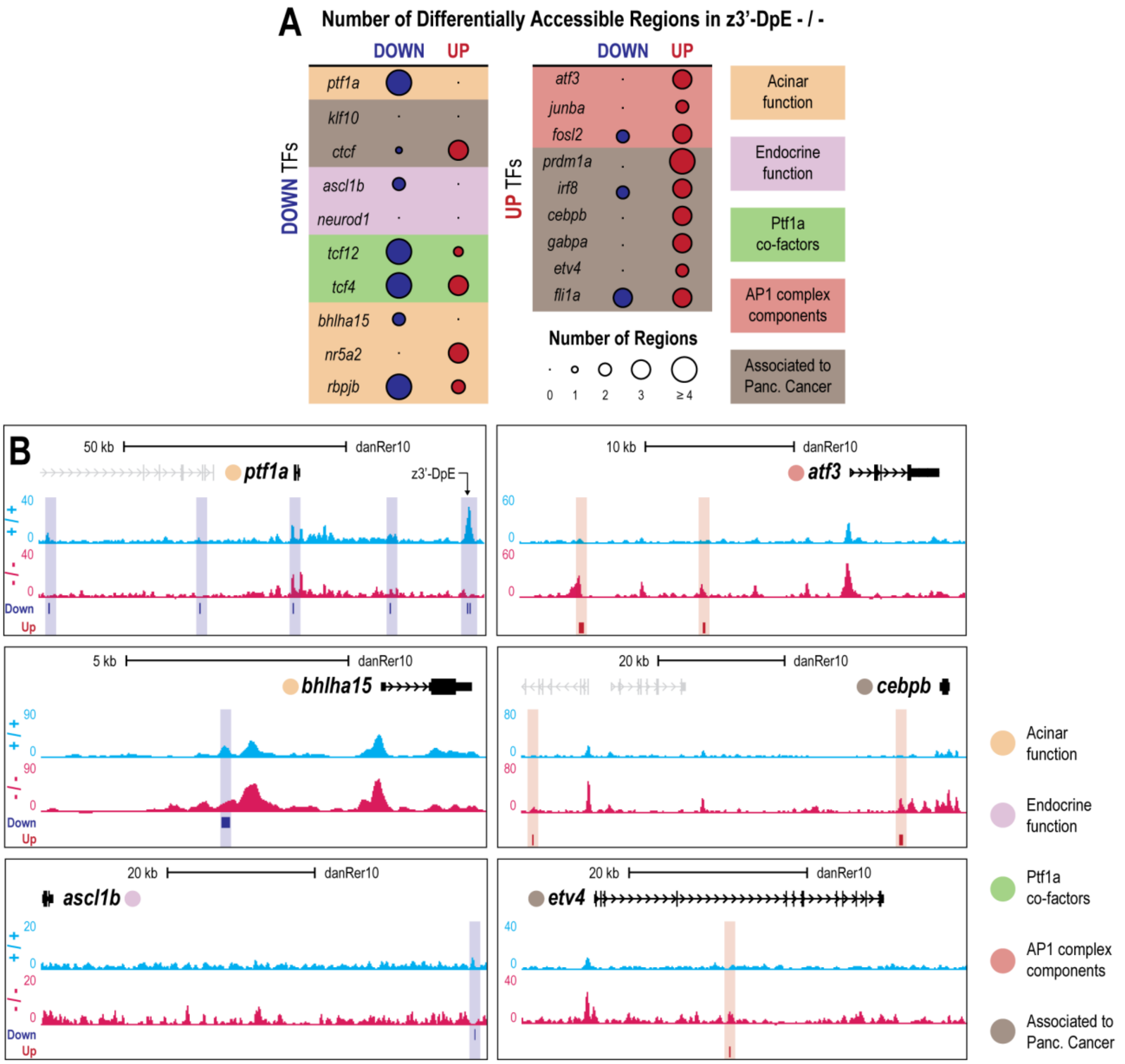

**Supplementary Figure 6. TF-associated chromatin changes in z3′-DpE-/- acinar cells.**

**(A)** Transcription factor (TF)-encoding genes associated with DARs in z3′-DpE-/- acinar cells. TFs are grouped into categories with related biological roles, including acinar-associated TFs, known Ptf1a co-factors, endocrine/progenitor-related TFs, AP-1 complex members, and TFs linked to pancreatic neoplasia (DisGeNET). Left: TF motifs enriched in regions that lose accessibility (DOWN TFs). Right: TF motifs enriched in regions that gain accessibility (UP TFs). Circle size indicates the number of DARs associated with each gene; blue and red circles denote decreased (DOWN) and increased (UP) accessibility, respectively.

**(B)** Genome browser views of representative TF-encoding genes whose motifs are enriched in DOWN (*ptf1a*, *bhlha15*, *ascl1b*) or UP DARs (*atf3*, *cebpb*, *etv4*). ATAC-seq tracks from z3′-DpE+/+ (blue) and z3′-DpE-/- (red) acinar cells are shown, with DARs highlighted in blue (Down) and red (Up).

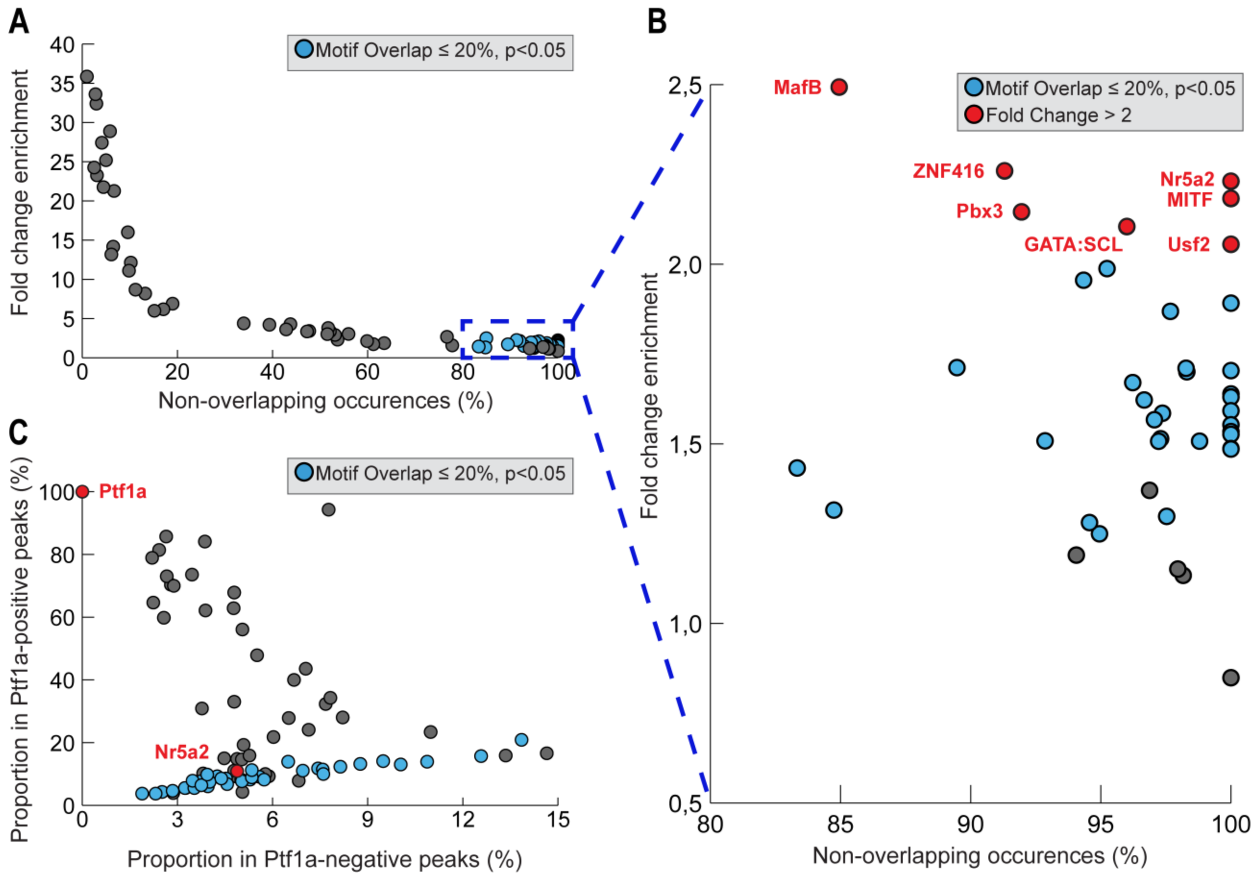

**Supplementary Figure 7. Identification of TF motifs co-enriched with Ptf1a in downregulated chromatin accessibility regions.**

**(A)** TF motifs enriched in Ptf1a-positive peaks relative to Ptf1a-negative peaks in regions that lose accessibility in z3′-DpE-/- acinar cells (Down DARs). Each dot represents one motif; the y-axis indicates enrichment (fold-change in motif frequency in Ptf1a-positive versus Ptf1a-negative peaks), and the x-axis indicates the percentage of motif occurrences not overlapping predicted Ptf1a sites. Motifs retained after filtering for sequence similarity to Ptf1a (≤ 20% of motif occurrences overlapping Ptf1a sites) and significant enrichment in Ptf1a-positive peaks (Fisher's exact test, Benjamini-Hochberg FDR < 0.05; see Methods) are highlighted in blue.

**(B)** Zoomed view of motifs with high non-overlapping occurrence (right-side region indicated in A). Motifs meeting selection criteria (blue) and exhibiting ≥ 2-fold enrichment in Ptf1a-positive versus Ptf1a-negative peaks are highlighted in red and labeled.

**(C)** Motif occurrence frequencies shown as the proportion of Down DARs containing each motif in Ptf1a-negative (x-axis) versus Ptf1a-positive (y-axis) peak sets. Ptf1a and Nr5a2 motifs are highlighted in red. The Ptf1a motif is shown for reference: 100% prevalence in Ptf1a-positive peaks and 0% in Ptf1a-negative peaks. Nr5a2, a known acinar identity regulator, shows enrichment in Ptf1a-positive peaks compared to Ptf1a-negative peaks, supporting independent co-occurrence with Ptf1a.

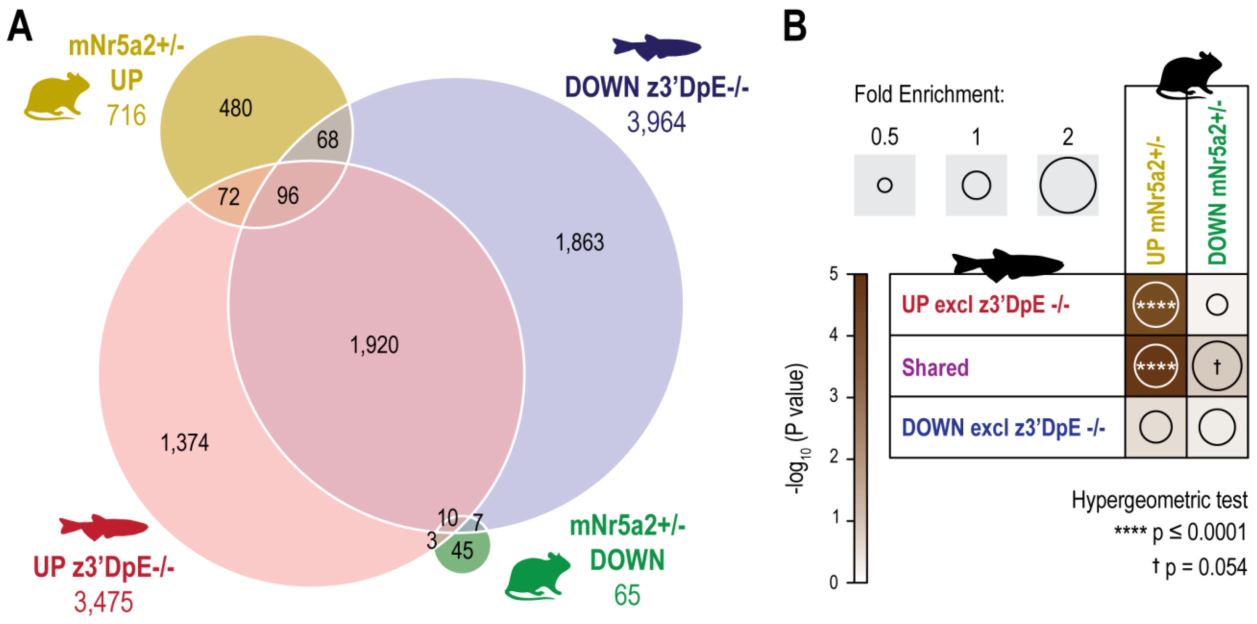

**Supplementary Figure 8. Cross-species comparison of z3'-DpE-/- zebrafish and Nr5a2+/- mouse model.**

**(A)** Venn diagram showing overlap between genes associated with differentially accessible chromatin regions in z3'-DpE-/- zebrafish acinar cells and differentially expressed genes in Nr5a2+/- mouse pancreas (whole tissue). Gene lists were converted to human orthologs for cross-species comparison. UP (yellow/red): increased accessibility or expression in mutants; DOWN (blue/green): decreased accessibility or expression.

**(B)** Hypergeometric enrichment analysis of gene overlap between models. Circle size represents fold enrichment; significance levels shown as -log10(p-value). Significant enrichment (****p ≤ 0.0001) was observed for genes with increased accessibility in z3'-DpE-/- acinar cells overlapping with upregulated genes in Nr5a2+/- pancreas, suggesting convergence on inflammatory pathways. †p = 0.054 indicates trend-level enrichment for shared genes showing bidirectional changes across models.

### Supplementary Tables

#### Supplementary Table 1. GO enrichment of downregulated genes in z3′-DpE-/- MPCs (48 hpf RNA-seq).

Source: RNA-seq of FACS-purified pancreatic multipotent progenitor cells (MPCs) at 48 hpf from z3′-DpE+/+ and z3′-DpE-/- animals (*Danio rerio*). Criteria: Differential expression (z3′-DpE-/- vs z3′-DpE+/+) defined as FDR ≤ 0.05 and |log2FC| ≥ 1 (DESeq2). GO Biological Process enrichment of downregulated genes performed using the PANTHER Overrepresentation Test (*Danio rerio* reference list; Fisher's exact test with FDR correction).

| GO term (biological process) | GO ID | Ref list count (N=26248) | Query list count (n=84) | Expected | Fold enrichment | Raw P-value | FDR |
|---|---|---|---|---|---|---|---|
| positive regulation of circadian rhythm | GO:0042753 | 4 | 2 | .01 | > 100 | 6.05E-05 | 1.59E-02 |
| CRD-mediated mRNA stabilization | GO:0070934 | 6 | 2 | .02 | > 100 | 1.51E-04 | 2.69E-02 |
| thyroid gland development | GO:0030878 | 18 | 3 | .06 | 52.08 | 2.49E-05 | 8.90E-03 |
| midbrain development | GO:0030901 | 39 | 3 | .12 | 24.04 | 2.66E-04 | 3.83E-02 |
| spindle localization | GO:0051653 | 40 | 3 | .13 | 23.44 | 2.87E-04 | 4.00E-02 |
| establishment of spindle localization | GO:0051293 | 40 | 3 | .13 | 23.44 | 2.87E-04 | 3.94E-02 |
| Notch signaling pathway | GO:0007219 | 81 | 4 | .26 | 15.43 | 1.35E-04 | 2.56E-02 |
| regulation of mRNA stability | GO:0043488 | 88 | 4 | .28 | 14.20 | 1.85E-04 | 2.96E-02 |
| regulation of RNA stability | GO:0043487 | 93 | 4 | .30 | 13.44 | 2.29E-04 | 3.53E-02 |
| regulation of mRNA catabolic process | GO:0061013 | 98 | 4 | .31 | 12.75 | 2.80E-04 | 3.98E-02 |
| protein polyubiquitination | GO:0000209 | 126 | 5 | .40 | 12.40 | 5.36E-05 | 1.50E-02 |
| cell population proliferation | GO:0008283 | 159 | 5 | .51 | 9.83 | 1.61E-04 | 2.76E-02 |
| stem cell differentiation | GO:0048863 | 237 | 7 | .76 | 9.23 | 1.12E-05 | 5.28E-03 |
| epithelial tube morphogenesis | GO:0060562 | 245 | 6 | .78 | 7.65 | 1.38E-04 | 2.51E-02 |
| cranial skeletal system development | GO:1904888 | 252 | 6 | .81 | 7.44 | 1.60E-04 | 2.81E-02 |
| protein ubiquitination | GO:0016567 | 409 | 8 | 1.31 | 6.11 | 5.04E-05 | 1.50E-02 |
| chordate embryonic development | GO:0043009 | 538 | 10 | 1.72 | 5.81 | 8.54E-06 | 4.77E-03 |
| embryo development ending in birth or egg hatching | GO:0009792 | 541 | 10 | 1.73 | 5.78 | 8.97E-06 | 4.45E-03 |
| protein modification by small protein conjugation | GO:0032446 | 451 | 8 | 1.44 | 5.54 | 9.94E-05 | 2.17E-02 |
| embryonic morphogenesis | GO:0048598 | 736 | 13 | 2.36 | 5.52 | 5.98E-07 | 5.93E-04 |
| actin cytoskeleton organization | GO:0030036 | 514 | 9 | 1.64 | 5.47 | 3.95E-05 | 1.31E-02 |

| | | | | | | | |
|---|---|---|---|---|---|---|---|
| tissue morphogenesis | GO:0048729 | 583 | 10 | 1.87 | 5.36 | 1.71E-05 | 6.64E-03 |
| actin filament-based process | GO:0030029 | 541 | 9 | 1.73 | 5.20 | 5.87E-05 | 1.59E-02 |
| tube morphogenesis | GO:0035239 | 603 | 10 | 1.93 | 5.18 | 2.28E-05 | 8.49E-03 |
| morphogenesis of an epithelium | GO:0002009 | 485 | 8 | 1.55 | 5.15 | 1.64E-04 | 2.76E-02 |
| embryo development | GO:0009790 | 1179 | 19 | 3.77 | 5.04 | 4.43E-09 | 9.89E-06 |
| embryonic organ development | GO:0048568 | 647 | 10 | 2.07 | 4.83 | 4.14E-05 | 1.27E-02 |
| head development | GO:0060322 | 587 | 9 | 1.88 | 4.79 | 1.09E-04 | 2.22E-02 |
| tube development | GO:0035295 | 798 | 12 | 2.55 | 4.70 | 8.72E-06 | 4.58E-03 |
| protein modification by small protein conjugation or removal | GO:0070647 | 538 | 8 | 1.72 | 4.65 | 3.29E-04 | 4.39E-02 |
| regulation of developmental process | GO:0050793 | 921 | 13 | 2.95 | 4.41 | 7.02E-06 | 4.18E-03 |
| epithelium development | GO:0060429 | 1006 | 14 | 3.22 | 4.35 | 3.51E-06 | 2.42E-03 |
| tissue development | GO:0009888 | 1619 | 22 | 5.18 | 4.25 | 4.85E-09 | 8.66E-06 |
| negative regulation of metabolic process | GO:0009892 | 931 | 12 | 2.98 | 4.03 | 3.99E-05 | 1.27E-02 |
| negative regulation of macromolecule metabolic process | GO:0010605 | 898 | 11 | 2.87 | 3.83 | 1.35E-04 | 2.61E-02 |
| positive regulation of metabolic process | GO:0009893 | 936 | 11 | 3.00 | 3.67 | 1.93E-04 | 3.02E-02 |
| circulatory system development | GO:0072359 | 1031 | 12 | 3.30 | 3.64 | 1.06E-04 | 2.24E-02 |
| animal organ development | GO:0048513 | 2668 | 30 | 8.54 | 3.51 | 3.06E-10 | 2.73E-06 |
| cell surface receptor signaling pathway | GO:0007166 | 1198 | 13 | 3.83 | 3.39 | 1.07E-04 | 2.23E-02 |
| regulation of signal transduction | GO:0009966 | 1391 | 14 | 4.45 | 3.14 | 1.26E-04 | 2.49E-02 |
| cell development | GO:0048468 | 1833 | 18 | 5.87 | 3.07 | 1.61E-05 | 6.84E-03 |
| anatomical structure morphogenesis | GO:0009653 | 2407 | 23 | 7.70 | 2.99 | 1.24E-06 | 1.11E-03 |
| multicellular organism development | GO:0007275 | 4053 | 36 | 12.97 | 2.78 | 1.72E-09 | 7.67E-06 |
| protein modification process | GO:0036211 | 1716 | 15 | 5.49 | 2.73 | 3.27E-04 | 4.43E-02 |
| system development | GO:0048731 | 3562 | 31 | 11.40 | 2.72 | 6.76E-08 | 7.55E-05 |
| positive regulation of biological process | GO:0048518 | 2342 | 20 | 7.49 | 2.67 | 3.78E-05 | 1.30E-02 |
| nervous system development | GO:0007399 | 2002 | 17 | 6.41 | 2.65 | 1.74E-04 | 2.83E-02 |
| cell differentiation | GO:0030154 | 2598 | 22 | 8.31 | 2.65 | 1.58E-05 | 7.06E-03 |
| cellular developmental process | GO:0048869 | 2603 | 22 | 8.33 | 2.64 | 1.63E-05 | 6.62E-03 |
| positive regulation of cellular process | GO:0048522 | 2129 | 17 | 6.81 | 2.50 | 3.60E-04 | 4.66E-02 |
| negative regulation of cellular process | GO:0048523 | 2260 | 18 | 7.23 | 2.49 | 2.41E-04 | 3.59E-02 |

| | | | | | | | |
|---|---|---|---|---|---|---|---|
| multicellular organismal process | GO:0032501 | 4946 | 39 | 15.83 | 2.46 | 7.66E-09 | 1.14E-05 |
| developmental process | GO:0032502 | 5270 | 41 | 16.87 | 2.43 | 3.45E-09 | 1.03E-05 |
| anatomical structure development | GO:0048856 | 5032 | 39 | 16.10 | 2.42 | 1.26E-08 | 1.61E-05 |
| negative regulation of biological process | GO:0048519 | 2347 | 18 | 7.51 | 2.40 | 3.83E-04 | 4.88E-02 |
| regulation of RNA metabolic process | GO:0051252 | 2978 | 22 | 9.53 | 2.31 | 1.66E-04 | 2.74E-02 |
| regulation of gene expression | GO:0010468 | 3465 | 25 | 11.09 | 2.25 | 6.16E-05 | 1.57E-02 |
| regulation of macromolecule biosynthetic process | GO:0010556 | 3503 | 25 | 11.21 | 2.23 | 7.17E-05 | 1.78E-02 |
| regulation of nucleobase-containing compound metabolic process | GO:0019219 | 3093 | 22 | 9.90 | 2.22 | 2.45E-04 | 3.58E-02 |
| regulation of biosynthetic process | GO:0009889 | 3565 | 25 | 11.41 | 2.19 | 9.25E-05 | 2.12E-02 |
| regulation of metabolic process | GO:0019222 | 4081 | 28 | 13.06 | 2.14 | 5.20E-05 | 1.50E-02 |
| regulation of macromolecule metabolic process | GO:0060255 | 3847 | 26 | 12.31 | 2.11 | 1.37E-04 | 2.55E-02 |
| cellular component organization | GO:0016043 | 3995 | 27 | 12.78 | 2.11 | 9.29E-05 | 2.07E-02 |
| cellular component organization or biogenesis | GO:0071840 | 4177 | 27 | 13.37 | 2.02 | 2.34E-04 | 3.54E-02 |
| regulation of cellular process | GO:0050794 | 9413 | 51 | 30.12 | 1.69 | 5.57E-06 | 3.55E-03 |
| biological regulation | GO:0065007 | 10229 | 55 | 32.74 | 1.68 | 1.43E-06 | 1.16E-03 |
| regulation of biological process | GO:0050789 | 9887 | 53 | 31.64 | 1.68 | 2.52E-06 | 1.88E-03 |
| cellular process | GO:0009987 | 14971 | 64 | 47.91 | 1.34 | 3.54E-04 | 4.64E-02 |
| biological_process | GO:0008150 | 19453 | 77 | 62.25 | 1.24 | 8.23E-05 | 1.99E-02 |

**Supplementary Table 2. scRNA-seq cell recovery and cluster composition of adult whole pancreata.**

Source: Single-cell RNA sequencing of dissociated whole adult pancreata from z3′-DpE+/+ (WT) and z3′-DpE-/- (Mut) animals; counts reflect cells retained after quality control and used for clustering and downstream analyses. Criteria: Cells were clustered and annotated based on marker gene expression as described in Methods. Cluster 6 corresponds to pancreatic cells; the subcluster table reports re-clustering of cluster 6. Percentages for cluster 6 subclusters are calculated within cluster 6 for each genotype.

| Cluster | n cells (WT) | n cells (Mut) |
|---|---|---|
| Stromal cells 1 | 1613 | 1175 |
| T and NK cells 1 | 1396 | 1026 |
| Stromal cells 2 | 408 | 692 |
| Stromal cells 3 | 690 | 329 |
| Blood cells 1 | 457 | 401 |
| Pancreatic cells | 391 | 435 |
| Unknown cells 1 | 239 | 191 |
| Blood cells 2 | 265 | 132 |
| Unknown cells 2 | 108 | 195 |
| ILC2 cells 1 | 81 | 129 |
| T and NK cells 2 | 72 | 98 |
| ILC2 cells 2 | 45 | 121 |
| Unknown cells 3 | 25 | 89 |
| Unknown cells 4 | 18 | 50 |
| **Subcluster of Pancreatic cells** | **n cells (WT)** | **n cells (Mut)** |
| T and NK cells | 89 | 44 |
| Acinar cells | 55 | 76 |
| Endocrine cells | 25 | 83 |
| Macrophages and monocytes cells | 56 | 31 |
| Ribossomal cells | 40 | 41 |
| Unknown cells 1 | 51 | 29 |
| Duct cells | 22 | 53 |
| Endothelial cells | 28 | 41 |
| Unknown cells 2 | 13 | 30 |
| Unknown cells 3 | 12 | 7 |

**Supplementary Table 3. Cluster-wise differentially expressed genes (scRNA-seq; z3′-DpE-/- vs z3′-DpE+/+).**

Source: Single-cell RNA sequencing (scRNA-seq) of whole adult pancreata from z3′-DpE+/+ and z3′-DpE-/- animals. Criteria: Differential expression was performed within each cluster using Seurat FindMarkers (Wilcoxon rank-sum test). Genes listed meet adjusted p-value (p_adj) ≤ 0.05.

| cluster | gene | p_val | avg_logFC | pct.1 | pct.2 | p_val_adj |
|---|---|---|---|---|---|---|
| 1 | nptna | 5.30293215925396E-160 | 3.37 | 0.52 | 0.08 | 1.36513382575675E-155 |
| 1 | ins | 8.70413486348684E-156 | 3.99 | 0.46 | 0.06 | 2.24070543790742E-151 |
| 1 | si:ch211-209n20.3 | 8.35979652402066E-140 | 3.88 | 0.42 | 0.05 | 2.15206241917864E-135 |
| 1 | junbb | 4.68918381292811E-60 | 1.71 | 0.51 | 0.23 | 1.20713658896208E-55 |
| 1 | si:dkey-78o7.3 | 5.22946791219384E-49 | 1.99 | 0.31 | 0.09 | 1.34622192463606E-44 |
| 1 | si:busm1-266f07.2 | 4.50693461089748E-44 | 1.03 | 0.63 | 0.45 | 1.16022017688334E-39 |
| 1 | si:dkey-112a7.4 | 8.68593453957113E-40 | 1.09 | 0.57 | 0.37 | 2.2360201285218E-35 |
| 1 | higd1a | 1.0591839958828E-36 | 1.26 | 0.46 | 0.24 | 2.72665736060109E-32 |
| 1 | cebpb | 1.15425066095122E-34 | 1.1 | 0.64 | 0.47 | 2.97138747648673E-30 |
| 1 | si:dkey-3h2.4 | 4.74873581865796E-33 | 1.09 | 0.52 | 0.33 | 1.22246706179712E-28 |
| 1 | ets2 | 5.0970171353203E-33 | 1.69 | 0.32 | 0.14 | 1.31212512114551E-28 |
| 1 | ccnl1b | 4.15392679049572E-29 | 1.18 | 0.4 | 0.22 | 1.06934537367731E-24 |
| 1 | tnfrsf9a | 7.02159788575869E-26 | 1.6 | 0.35 | 0.22 | 1.80756994373086E-21 |
| 1 | stat2 | 7.9782754687367E-25 | 1.44 | 0.32 | 0.17 | 2.05384745391689E-20 |
| 1 | npl | 1.46496999793318E-24 | 1.2 | 0.27 | 0.11 | 3.77127226567938E-20 |
| 1 | rc3h2 | 1.18186368807845E-23 | 1.03 | 0.39 | 0.24 | 3.04247169222034E-19 |
| 1 | mxra8b | 1.95348559498386E-22 | 1.52 | 0.26 | 0.14 | 5.02885796716695E-18 |
| 1 | tox | 1.96579576063608E-22 | 1.24 | 0.32 | 0.16 | 5.06054802660547E-18 |
| 1 | ccr9a | 5.10801265205973E-22 | 1.54 | 0.34 | 0.2 | 1.31495569701974E-17 |
| 1 | si:ch211-256a21.4 | 7.4419064870132E-22 | 1.05 | 0.34 | 0.18 | 1.91576998695181E-17 |
| 1 | tcf7 | 4.14302613277411E-20 | 1.08 | 0.42 | 0.27 | 1.06653921736004E-15 |
| 1 | hsp70.2 | 9.35609651115257E-18 | 1.82 | 0.26 | 0.15 | 2.40853992486601E-13 |
| 1 | iscub | 1.95052556525884E-15 | 1,00 | 0.31 | 0.19 | 5.02123796264584E-11 |
| 1 | tob1b | 1.96521526419931E-12 | 1.06 | 0.27 | 0.16 | 5.05905365462829E-08 |
| 1 | CABZ01045617.1 | 7.72992774462919E-12 | 1.08 | 0.5 | 0.45 | 1.98991529929989E-07 |
| 1 | gzmk | 4.07645758771375E-15 | -1.11 | 0.15 | 0.25 | 1.04940247680515E-10 |
| 1 | egr2b | 2.51803134876455E-17 | -1.25 | 0.19 | 0.32 | 6.48216810112459E-13 |
| 1 | kctd12.2 | 1.46335014119183E-22 | -1.5 | 0.13 | 0.26 | 3.76710226847013E-18 |
| 1 | nitr3a | 2.03478153769671E-23 | -1.39 | 0.15 | 0.3 | 5.23813811249264E-19 |
| 1 | si:ch211-165b19.5 | 1.62168281213885E-23 | -1.29 | 0.16 | 0.31 | 4.17469806328904E-19 |
| 1 | sh2d1ab | 1.35752587116815E-24 | -1.23 | 0.24 | 0.34 | 3.49467885014818E-20 |
| 1 | ccl38a.5 | 3.53143766556786E-25 | -1.37 | 0.16 | 0.33 | 9.09097998247135E-21 |
| 1 | ifng1-2 | 8.43585205511273E-27 | -1.28 | 0.23 | 0.4 | 2.17164139454767E-22 |
| 1 | p2rx1 | 2.73305164108773E-27 | -1.53 | 0.16 | 0.31 | 7.03569483965214E-23 |

| | | | | | | |
|---|---|---|---|---|---|---|
| 1 | zgc:103700 | 6.49243080665394E-28 | -1.6 | 0.1 | 0.26 | 1.67134646255692E-23 |
| 1 | tapbp.1 | 5.85370532276516E-28 | -1.17 | 0.16 | 0.34 | 1.50691936123943E-23 |
| 1 | si:ch211-165b19.8 | 4.42067588878234E-28 | -1.76 | 0.11 | 0.27 | 1.13801459404924E-23 |
| 1 | BX005223.1 | 7.46557890456924E-29 | -1.33 | 0.17 | 0.35 | 1.92186397740326E-24 |
| 1 | si:dkey-11f4.16 | 5.08656980952172E-29 | -1.27 | 0.22 | 0.4 | 1.30943566606518E-24 |
| 1 | lipia | 4.40295979226129E-29 | -1.95 | 0.11 | 0.26 | 1.13345393932182E-24 |
| 1 | igsf11 | 8.95729716644857E-34 | -1.56 | 0.14 | 0.33 | 2.30587700955885E-29 |
| 1 | ENSDARG00000079078 | 8.93774401702438E-35 | -1.04 | 0.29 | 0.52 | 2.30084344230259E-30 |
| 1 | cadm4 | 4.24432203738578E-37 | -1.36 | 0.09 | 0.29 | 1.09261582208422E-32 |
| 1 | gzm3 | 1.85295153671495E-38 | -1.73 | 0.17 | 0.35 | 4.77005314096529E-34 |
| 1 | ccl34b.4 | 6.79310342230848E-40 | -1.58 | 0.49 | 0.67 | 1.74874861400487E-35 |
| 1 | gig2i | 6.76408597106343E-40 | -1.47 | 0.19 | 0.4 | 1.74127865153086E-35 |
| 1 | hbba2 | 1.00722383514329E-48 | -1.99 | 0.08 | 0.29 | 2.59289631880938E-44 |
| 1 | zgc:153317 | 4.99475799926606E-73 | -1.37 | 0.21 | 0.55 | 1.28580055175106E-68 |
| 1 | hbba1 | 4.50010173305732E-81 | -1.84 | 0.18 | 0.52 | 1.15846118914095E-76 |
| 1 | hbaa1 | 3.17912424062086E-85 | -1.48 | 0.35 | 0.7 | 8.18401953263027E-81 |
| 1 | si:busm1-48c11.3 | 3.39997866671942E-130 | -10.66 | 0,00 | 0.3 | 8.7525650817358E-126 |
| 1 | si:busm1-194e12.12 | 2.09410009232158E-136 | -10.73 | 0,00 | 0.31 | 5.39084186766344E-132 |
| 1 | CABZ01074309.1 | 3.91019959277333E-151 | -2.19 | 0.26 | 0.71 | 1.00660268116764E-146 |
| 1 | hbba1.1 | 2.15203124408014E-156 | -1.64 | 0.49 | 0.89 | 5.5399740316355E-152 |
| 2 | CABZ01040076.1 | 4.83934443227059E-204 | 7.21 | 0.46 | 0,00 | 1.24579243719942E-199 |
| 2 | ms4a17a.9 | 1.64755710096238E-192 | 5.85 | 0.49 | 0.02 | 4.24130624500746E-188 |
| 2 | ins | 1.74347815761731E-131 | 4.32 | 0.53 | 0.09 | 4.48823582115423E-127 |
| 2 | junbb | 9.42299558558844E-94 | 1.81 | 0.71 | 0.36 | 2.42576175359803E-89 |
| 2 | si:busm1-266f07.2 | 1.14335239811761E-64 | 1.1 | 0.96 | 0.93 | 2.94333207847417E-60 |
| 2 | si:ch73-236c18.6 | 9.4549082223406E-64 | 2.24 | 0.33 | 0.07 | 2.43397702367714E-59 |
| 2 | si:dkey-19a16.2 | 2.20231704188002E-59 | 2.06 | 0.26 | 0.04 | 5.66942476091174E-55 |
| 2 | ms4a17a.11 | 1.02862776369321E-53 | 2.55 | 0.29 | 0.06 | 2.64799645207542E-49 |
| 2 | fkbp5 | 4.42354388031387E-49 | 1.89 | 0.3 | 0.07 | 1.1387529011092E-44 |
| 2 | ch25h | 1.75579650359384E-47 | 1.65 | 0.48 | 0.2 | 4.51994693920162E-43 |
| 2 | hspa5 | 5.20282207946606E-42 | 1.48 | 0.51 | 0.25 | 1.33936248791695E-37 |
| 2 | sytl3 | 2.50701682667399E-39 | 2.33 | 0.26 | 0.06 | 6.45381341690686E-35 |
| 2 | hsp70.1 | 2.91275246602414E-39 | 2.09 | 0.38 | 0.15 | 7.49829867328593E-35 |
| 2 | ctsba | 4.12162175364698E-39 | 1.44 | 0.61 | 0.4 | 1.06102908804134E-34 |
| 2 | ndufa4 | 1.89065542838096E-36 | 1.8 | 0.26 | 0.07 | 4.86711426928112E-32 |
| 2 | rc3h2 | 3.45176134240499E-33 | 1.13 | 0.51 | 0.27 | 8.88586922375317E-29 |
| 2 | yrk | 3.82475202170071E-33 | 1.33 | 0.72 | 0.58 | 9.84605912946414E-29 |
| 2 | si:dkey-263j23.1 | 8.35851097622272E-33 | 1.78 | 0.31 | 0.12 | 2.15173148060901E-28 |
| 2 | pdk2b | 1.7296035299954E-32 | 1.04 | 0.53 | 0.29 | 4.45251836726717E-28 |
| 2 | NPC2 (1 of many) | 1.83791749746208E-32 | 1.2 | 0.57 | 0.35 | 4.73135101371663E-28 |
| 2 | jdp2b | 1.52893414237099E-31 | 1.33 | 0.49 | 0.26 | 3.93593516270565E-27 |
| 2 | si:dkey-112a7.4 | 3.18061344598784E-31 | 1.21 | 0.48 | 0.26 | 8.18785319400651E-27 |
| 2 | hsp70.3 | 8.38219452933202E-30 | 1.76 | 0.37 | 0.17 | 2.15782833768594E-25 |

| | | | | | | |
|---|---|---|---|---|---|---|
| 2 | enpp1 | 1.59854306595852E-29 | 1.19 | 0.59 | 0.39 | 4.11512941469702E-25 |
| 2 | ccl35.1 | 2.82706985418781E-29 | 1.51 | 0.61 | 0.4 | 7.27772592563567E-25 |
| 2 | atp1a3a | 2.93640480879852E-29 | 1.28 | 0.38 | 0.17 | 7.55918689929003E-25 |
| 2 | hsp70.2 | 3.38017519670723E-28 | 1.36 | 0.47 | 0.26 | 8.70158500888342E-24 |
| 2 | si:dkey-188i13.7 | 3.49683636533982E-26 | 1.62 | 0.31 | 0.15 | 9.00190585529431E-22 |
| 2 | hsp70l | 5.05139803301881E-26 | 1.14 | 0.52 | 0.31 | 1.30038139564003E-21 |
| 2 | CR848788.1 | 8.20841032472102E-26 | 1.28 | 0.32 | 0.14 | 2.11309106989293E-21 |
| 2 | chordc1a | 1.04468120819333E-24 | 1.17 | 0.25 | 0.09 | 2.68932283425209E-20 |
| 2 | phf20a | 4.1560521371069E-24 | 1.13 | 0.29 | 0.12 | 1.06989250165543E-19 |
| 2 | hspa9 | 7.07389445174881E-23 | 1.11 | 0.37 | 0.18 | 1.8210326487137E-18 |
| 2 | sec24d | 7.66353137946737E-22 | 1.03 | 0.38 | 0.2 | 1.97282288301629E-17 |
| 2 | si:dkey-51e6.1 | 4.08716218172507E-21 | 1.33 | 0.28 | 0.13 | 1.05215816044148E-16 |
| 2 | nfil3-6 | 3.96198576782198E-19 | 1.26 | 0.25 | 0.11 | 1.01993399621041E-14 |
| 2 | bcam | 2.60574352625396E-17 | 1.4 | 0.38 | 0.27 | 6.70796555963558E-13 |
| 2 | zfand2a | 3.13265158907934E-17 | 1.08 | 0.36 | 0.2 | 8.06438498576695E-13 |
| 2 | timp4.2 | 8.9217923880046E-17 | 1.39 | 0.29 | 0.2 | 2.29673701444402E-12 |
| 2 | CABZ01076275.1 | 2.56308517065671E-16 | 1.33 | 0.27 | 0.14 | 6.59815015482156E-12 |
| 2 | hspbp1 | 4.36248329311035E-16 | 1.59 | 0.25 | 0.12 | 1.1230340741454E-11 |
| 2 | ccl34b.1 | 9.08553780844574E-15 | 1.19 | 0.35 | 0.21 | 2.33888999802819E-10 |
| 2 | dagla | 9.3499084232199E-15 | 1.32 | 0.26 | 0.14 | 2.4069469253895E-10 |
| 2 | ier5l | 1.6724572362533E-14 | 1.04 | 0.37 | 0.24 | 4.30540666328686E-10 |
| 2 | CABZ01045062.1 | 4.50773418861743E-14 | 1.14 | 0.32 | 0.19 | 1.16042601217579E-09 |
| 2 | oser1 | 7.87758457347486E-14 | 1.05 | 0.35 | 0.22 | 2.02792659674963E-09 |
| 2 | dennd4a | 6.89185137849543E-13 | 1.04 | 0.3 | 0.17 | 1.77416930036608E-08 |
| 2 | sst2 | 0.0000017533151633624 | -2.93 | 0.31 | 0.22 | 0.0451355922504383 |
| 2 | fam65b | 3.51985816951194E-10 | -1.13 | 0.25 | 0.32 | 9.06117088577459E-06 |
| 2 | map3k8 | 2.22369277898723E-10 | -1.02 | 0.15 | 0.26 | 5.72445232094682E-06 |
| 2 | tex2 | 1.0678320058953E-10 | -1.04 | 0.17 | 0.25 | 2.74891993277626E-06 |
| 2 | fam212ab | 8.98304912483519E-11 | -1.05 | 0.21 | 0.27 | 2.31250633620632E-06 |
| 2 | kdm2ba | 8.69820078848596E-11 | -1.22 | 0.17 | 0.26 | 2.23917782897994E-06 |
| 2 | fes | 2.24782253823682E-12 | -1.15 | 0.21 | 0.29 | 5.78656956018305E-08 |
| 2 | gba | 2.39817354388893E-13 | -1.1 | 0.22 | 0.32 | 6.17361815403326E-09 |
| 2 | sardh | 3.10900041937428E-14 | -1.23 | 0.14 | 0.26 | 8.00349977959521E-10 |
| 2 | sptlc2a | 7.29930485893539E-15 | -1.01 | 0.31 | 0.41 | 1.87906004983574E-10 |
| 2 | tp53inp1 | 5.77769689338525E-15 | -1.14 | 0.24 | 0.36 | 1.48735251126416E-10 |
| 2 | ptbp1b | 4.23351101194839E-15 | -1.47 | 0.15 | 0.25 | 1.08983273980587E-10 |
| 2 | gbp1 | 4.07785931556358E-15 | -1.05 | 0.25 | 0.37 | 1.04976332360553E-10 |
| 2 | hsd17b14 | 2.12097573266121E-15 | -1.39 | 0.15 | 0.25 | 5.46002782858974E-11 |
| 2 | ENSDARG00000101094 | 1.58039410283599E-15 | -1.19 | 0.17 | 0.3 | 4.0684085389307E-11 |
| 2 | irg1 | 2.29026664857608E-16 | -1.12 | 0.22 | 0.38 | 5.8958334334294E-12 |
| 2 | moxd1 | 2.71228516857931E-17 | -1.05 | 0.28 | 0.4 | 6.98223570947372E-13 |
| 2 | sppl3 | 1.17760167568303E-17 | -1.19 | 0.24 | 0.38 | 3.03149999371083E-13 |
| 2 | myofl | 1.38445122823622E-18 | -1.03 | 0.26 | 0.42 | 3.5639927968485E-14 |

| | | | | | | |
|---|---|---|---|---|---|---|
| 2 | si:ch211-171h4.5 | 3.41082971975612E-19 | -1.1 | 0.26 | 0.41 | 8.78049894756817E-15 |
| 2 | si:dkey-195m11.11 | 3.17122561663675E-19 | -1.66 | 0.13 | 0.26 | 8.16368610490799E-15 |
| 2 | dennd2da | 2.008337767066E-19 | -1.01 | 0.3 | 0.44 | 5.17006391375801E-15 |
| 2 | epdl1 | 5.52857796401728E-20 | -1.49 | 0.32 | 0.43 | 1.42322182527697E-15 |
| 2 | cox6b1 | 4.49104479248054E-20 | -1.05 | 0.27 | 0.39 | 1.15612966092827E-15 |
| 2 | tns3.2 | 2.40785338449553E-20 | -1.41 | 0.23 | 0.37 | 6.19853696770685E-16 |
| 2 | mtmr11 | 1.26545023831529E-20 | -1.05 | 0.29 | 0.46 | 3.25764854849505E-16 |
| 2 | ppm1j | 6.67660188851194E-21 | -1.01 | 0.32 | 0.47 | 1.71875762415963E-16 |
| 2 | ENSDARG00000077648 | 2.45803664792199E-22 | -1.45 | 0.2 | 0.36 | 6.32772374274557E-18 |
| 2 | hspa12b | 2.05359806309478E-22 | -1.1 | 0.28 | 0.43 | 5.2865774938249E-18 |
| 2 | si:ch1073-443f11.2 | 3.27633567901129E-23 | -1.03 | 0.29 | 0.45 | 8.43427093847876E-19 |
| 2 | trim35-31 | 3.01872661098551E-23 | -1.75 | 0.12 | 0.26 | 7.77110791466E-19 |
| 2 | si:dkey-33i11.9 | 2.28347522506954E-23 | -1.36 | 0.33 | 0.45 | 5.87835027189652E-19 |
| 2 | zmp:0000000608 | 1.95509977233586E-23 | -1.22 | 0.11 | 0.28 | 5.03301334392421E-19 |
| 2 | mtss1lb | 6.94875904327829E-24 | -1.16 | 0.18 | 0.37 | 1.78881904051113E-19 |
| 2 | gig2d | 3.30185940034768E-24 | -1.29 | 0.21 | 0.38 | 8.49997665431504E-20 |
| 2 | slc36a1 | 3.12581260777291E-24 | -1.12 | 0.41 | 0.57 | 8.0467793961898E-20 |
| 2 | ano9a | 2.75347572895636E-24 | -1.51 | 0.19 | 0.36 | 7.08827256905237E-20 |
| 2 | gig2i | 8.09230739282766E-25 | -1.79 | 0.12 | 0.26 | 2.08320269213562E-20 |
| 2 | si:ch73-22o12.1 | 3.307542479865E-25 | -1.11 | 0.38 | 0.55 | 8.51460660591647E-21 |
| 2 | si:dkey-32n7.4 | 2.69181457861485E-26 | -1.37 | 0.25 | 0.41 | 6.92953826972821E-22 |
| 2 | ENSDARG00000074117 | 8.62369399162141E-28 | -1.23 | 0.08 | 0.25 | 2.2199975442631E-23 |
| 2 | lect2l | 1.00430756808714E-28 | -1.8 | 0.2 | 0.39 | 2.58538897252673E-24 |
| 2 | pstpip2 | 2.67890958359213E-30 | -1.06 | 0.47 | 0.65 | 6.89631694104121E-26 |
| 2 | MFAP4 (1 of many).3 | 1.34320485443794E-30 | -1.28 | 0.45 | 0.56 | 3.4578122567796E-26 |
| 2 | si:ch211-181d7.3 | 5.30993777874761E-31 | -1.94 | 0.11 | 0.28 | 1.366937282383E-26 |
| 2 | zgc:154125 | 2.62873474533753E-31 | -1.48 | 0.2 | 0.4 | 6.76715185492241E-27 |
| 2 | ptenb | 3.6272856043828E-37 | -1.17 | 0.47 | 0.66 | 9.33772133136263E-33 |
| 2 | stard8 | 1.22462737415013E-37 | -2.67 | 0.08 | 0.26 | 3.15255824927469E-33 |
| 2 | gtf2f2a | 1.00180237521684E-38 | -1.2 | 0.35 | 0.52 | 2.5789398545207E-34 |
| 2 | plxna1a | 7.99421908791069E-39 | -2.03 | 0.11 | 0.32 | 2.05795181980085E-34 |
| 2 | CABZ01074397.1 | 9.17626058483341E-40 | -1.07 | 0.47 | 0.67 | 2.36224476235366E-35 |
| 2 | card9 | 5.865497254938E-41 | -1.11 | 0.46 | 0.66 | 1.50995495833869E-36 |
| 2 | me1 | 3.59293427078802E-41 | -1.17 | 0.55 | 0.76 | 9.24929069328959E-37 |
| 2 | CABZ01088134.1 | 5.72896990822621E-42 | -1.37 | 0.35 | 0.55 | 1.47480872347467E-37 |
| 2 | mxc | 1.69534392549772E-50 | -2.61 | 0.06 | 0.28 | 4.36432386740878E-46 |
| 2 | hbba2 | 7.2426418063522E-51 | -1.07 | 0.08 | 0.33 | 1.86447328020925E-46 |
| 2 | tcirg1b | 1.02747037645617E-52 | -1,00 | 0.71 | 0.82 | 2.64501699011113E-48 |
| 2 | ptprfb | 4.81391628887301E-54 | -2.81 | 0.08 | 0.32 | 1.23924647024458E-49 |
| 2 | si:dkey-1h24.6 | 1.26785950543118E-55 | -1.71 | 0.24 | 0.53 | 3.26385072483149E-51 |
| 2 | tapbp.1 | 8.45688428155844E-56 | -1.69 | 0.28 | 0.54 | 2.17705572060159E-51 |
| 2 | ccl39.3 | 9.12796203350515E-57 | -1.93 | 0.13 | 0.41 | 2.34981126628523E-52 |
| 2 | zgc:158446 | 4.9829553284128E-57 | -1.36 | 0.42 | 0.64 | 1.28276219019331E-52 |

| | | | | | | |
|---|---|---|---|---|---|---|
| 2 | aif1l | 7.26862744978628E-65 | -1.5 | 0.56 | 0.77 | 1.87116276439848E-60 |
| 2 | hbba1 | 2.0338161088278E-68 | -1.01 | 0.22 | 0.56 | 5.23565280895541E-64 |
| 2 | zgc:103700 | 9.52320490036521E-72 | -1.01 | 0.81 | 0.92 | 2.45155863750102E-67 |
| 2 | hbaa1 | 4.58494466231845E-74 | -1.44 | 0.37 | 0.72 | 1.18030230442064E-69 |
| 2 | mhc1uba | 1.35808499203344E-77 | -1.9 | 0.28 | 0.63 | 3.4961181949917E-73 |
| 2 | def8 | 1.9066583260467E-78 | -1.29 | 0.65 | 0.85 | 4.90831052874203E-74 |
| 2 | BX640512.3 | 1.22202489249424E-84 | -2.41 | 0.15 | 0.51 | 3.14585868074793E-80 |
| 2 | hbba1.1 | 5.53419967187477E-122 | -1.14 | 0.55 | 0.91 | 1.42466902153072E-117 |
| 2 | mhc2dab | 6.65198450384916E-139 | -1.96 | 0.47 | 0.83 | 1.71242037082589E-134 |
| 2 | grn2 | 1.37779027529798E-142 | -2.39 | 0.17 | 0.67 | 3.54684550569958E-138 |
| 2 | CABZ01074309.1 | 2.08377533108191E-146 | -2.43 | 0.28 | 0.75 | 5.36426283480417E-142 |
| 2 | si:busm1-48c11.3 | 0 | -11.83 | 0,00 | 0.75 | 0 |
| 2 | si:busm1-194e12.12 | 0 | -12.3 | 0,00 | 0.8 | 0 |
| 3 | ins | 9.62945969367752E-49 | 3.59 | 0.44 | 0.06 | 2.47891180894341E-44 |
| 3 | si:dkey-3h2.4 | 6.58096646479975E-36 | 1.6 | 0.72 | 0.42 | 1.6941381970334E-31 |
| 3 | si:dkey-3h2.2 | 8.22962333570617E-28 | 3.31 | 0.27 | 0.03 | 2.11855193531084E-23 |
| 3 | si:dkey-53k12.33 | 1.00627227613794E-25 | 1.09 | 0.79 | 0.6 | 2.59044672046191E-21 |
| 3 | nptna | 1.48999786389635E-25 | 1.33 | 0.61 | 0.29 | 3.83570150102837E-21 |
| 3 | si:ch211-209n20.3 | 1.47496741989648E-20 | 1.97 | 0.41 | 0.16 | 3.79700862903952E-16 |
| 3 | arl11 | 2.68583042292619E-20 | 1.6 | 0.45 | 0.2 | 6.91413325773888E-16 |
| 3 | CABZ01045617.1 | 1.61078486973659E-17 | 1.39 | 0.78 | 0.61 | 4.1466434901629E-13 |
| 3 | hsp70.1 | 1.20295314791343E-11 | 1.77 | 0.32 | 0.15 | 3.09676228867355E-07 |
| 3 | satb1b | 2.19497109992923E-11 | 1.31 | 0.32 | 0.14 | 5.65051410254782E-07 |
| 3 | hsp70l | 5.86790230736845E-11 | 1.15 | 0.69 | 0.57 | 1.51057409098586E-06 |
| 3 | fkbp5 | 1.47222265298878E-09 | 1.39 | 0.32 | 0.17 | 0.0000378994277558902 |
| 3 | nkl.2 | 4.283435172249E-08 | 1.89 | 0.81 | 0.72 | 0.00110268471639206 |
| 3 | si:busm1-266f07.2 | 1.87536927592851E-07 | 1.02 | 0.43 | 0.31 | 0.00482776312702275 |
| 3 | gip | 5.23400557965966E-07 | 1.25 | 0.36 | 0.23 | 0.0134739005637179 |
| 3 | hsp70.2 | 0.0000014000986922757 | 1.38 | 0.35 | 0.24 | 0.0360427406352533 |
| 3 | hpdb | 1.10336862818284E-08 | -1.63 | 0.17 | 0.26 | 0.000284040185953107 |
| 3 | si:dkey-10h3.2 | 1.32273424840719E-11 | -1.11 | 0.2 | 0.39 | 3.40511477567464E-07 |
| 3 | gig2i | 4.30962579676238E-12 | -1.33 | 0.12 | 0.3 | 1.10942696886054E-07 |
| 3 | cox6a2 | 1.14519123440728E-12 | -1.58 | 0.1 | 0.28 | 2.94806579473466E-08 |
| 3 | cmah | 3.50066232227668E-14 | -1.03 | 0.24 | 0.47 | 9.01175501623685E-10 |
| 3 | mt2 | 6.82313284602953E-16 | -1.51 | 0.13 | 0.34 | 1.75647908855338E-11 |
| 3 | sh3gl1a | 1.18005217791124E-18 | -1.2 | 0.2 | 0.47 | 3.03780832159692E-14 |
| 3 | hbba2 | 3.00591041654176E-19 | -2.27 | 0.06 | 0.25 | 7.73811518530344E-15 |
| 3 | hbaa1 | 2.74175819794663E-20 | -1.14 | 0.35 | 0.64 | 7.058108128974E-16 |
| 3 | si:dkey-32n7.4 | 2.64608919383414E-20 | -2.06 | 0.1 | 0.32 | 6.81182741168722E-16 |
| 3 | mibp2 | 3.66352218063325E-21 | -1.04 | 0.48 | 0.69 | 9.43100514960417E-17 |
| 3 | hbba1 | 2.73699642479998E-22 | -1.52 | 0.2 | 0.49 | 7.0458498963626E-18 |
| 3 | si:ch211-181d7.3 | 1.82870488801267E-27 | -2.55 | 0.07 | 0.32 | 4.70763499321101E-23 |
| 3 | ccl38a.5 | 9.85226010699926E-35 | -1.64 | 0.4 | 0.72 | 2.53626731934482E-30 |

| | | | | | | |
|---|---|---|---|---|---|---|
| 3 | hbba1.1 | 8.49714295302129E-42 | -1.41 | 0.48 | 0.83 | 2.18741951039627E-37 |
| 3 | cadm4 | 3.75422783610942E-46 | -2.97 | 0.1 | 0.48 | 9.66450871849649E-42 |
| 3 | si:busm1-194e12.12 | 8.72702277173693E-57 | -9.53 | 0,00 | 0.28 | 2.24659747212824E-52 |
| 3 | CABZ01074309.1 | 2.54178254692617E-67 | -1.95 | 0.35 | 0.79 | 6.54331081055204E-63 |
| 4 | sst3 | 1.47153723081921E-47 | 12.97 | 0.26 | 0,00 | 3.78817829329789E-43 |
| 4 | ins | 3.07869187871809E-43 | 3.57 | 0.42 | 0.07 | 7.92547650338397E-39 |
| 4 | zmat4b | 1.71209410928697E-21 | 2.35 | 0.32 | 0.09 | 4.40744386553746E-17 |
| 4 | tmem106ba | 2.74148224598166E-09 | 1.31 | 0.28 | 0.15 | 0.0000705739774583058 |
| 4 | ssuh2.1 | 0.000000136601656602 | 1.04 | 0.32 | 0.21 | 0.00351653644590529 |
| 4 | junbb | 1.44626649842227E-06 | 1.18 | 0.25 | 0.13 | 0.0372312384688846 |
| 4 | CABZ01074309.1 | 1.7222035922364E-08 | -1.32 | 0.13 | 0.29 | 0.000443346870749416 |
| 4 | tuba8l3 | 3.90792126747245E-11 | -1.63 | 0.12 | 0.31 | 1.00601617188543E-06 |
| 4 | rdh8a | 1.88473846725671E-12 | -1.54 | 0.14 | 0.35 | 4.85188223625894E-08 |
| 4 | slit3 | 4.0673713896981E-13 | -2.17 | 0.07 | 0.25 | 1.04706341684998E-08 |
| 4 | sc:d0202 | 1.7900102597627E-13 | -1.53 | 0.08 | 0.27 | 4.60802341170712E-09 |
| 4 | sgms1 | 4.65988223930456E-14 | -1.61 | 0.09 | 0.29 | 1.19959348486417E-09 |
| 4 | hbba2 | 3.72489587087248E-18 | -2.06 | 0.07 | 0.3 | 9.58899944038702E-14 |
| 4 | icn | 4.78938953230535E-23 | -1.96 | 0.2 | 0.5 | 1.23293254730137E-18 |
| 4 | hbba1 | 3.00803207510652E-23 | -1.64 | 0.22 | 0.55 | 7.74357697094671E-19 |
| 4 | hbaa1 | 3.92529647037406E-37 | -1.51 | 0.31 | 0.73 | 1.01048907036839E-32 |
| 4 | hbba1.1 | 3.09603890926562E-50 | -1.47 | 0.51 | 0.9 | 7.97013296412248E-46 |
| 5 | BX908782.1 | 5.38149972507E-87 | 3.94 | 0.76 | 0.15 | 1.38535947422477E-82 |
| 5 | gig2j | 1.04499853658154E-45 | 8.79 | 0.3 | 0,00 | 2.69013973272186E-41 |
| 5 | ndufa4 | 4.85027244094939E-38 | 2.89 | 0.49 | 0.13 | 1.2486056344736E-33 |
| 5 | si:busm1-266f07.2 | 3.13825697760123E-31 | 2.08 | 0.48 | 0.12 | 8.07881493743884E-27 |
| 5 | si:ch211-188c18.1 | 3.54339738000865E-27 | 2.61 | 0.43 | 0.14 | 9.12176787535626E-23 |
| 5 | nfkbiaa | 4.21952594852603E-27 | 1.38 | 0.85 | 0.68 | 1.08623256492906E-22 |
| 5 | ins | 7.89571316642602E-26 | 2.5 | 0.38 | 0.08 | 2.03259344043305E-21 |
| 5 | jun | 1.15121377666103E-24 | 1.6 | 0.64 | 0.31 | 2.9635696252585E-20 |
| 5 | junba | 4.10840659896478E-23 | 1.15 | 0.85 | 0.63 | 1.0576271107715E-18 |
| 5 | pim2 | 1.37088228928324E-19 | 1.03 | 0.77 | 0.51 | 3.52906227730184E-15 |
| 5 | tnip2 | 1.99610060435695E-19 | 1.38 | 0.52 | 0.23 | 5.13856178579609E-15 |
| 5 | dnajb1b | 6.15249406010188E-19 | 1.15 | 0.69 | 0.4 | 1.58383654589203E-14 |
| 5 | kcnh5a | 6.19966580461094E-18 | 2.41 | 0.26 | 0.05 | 1.595979968081E-13 |
| 5 | pim1 | 7.78645961306098E-17 | 1.4 | 0.53 | 0.27 | 2.00446829819029E-12 |
| 5 | mmp9 | 4.22837964174352E-16 | 1.23 | 0.64 | 0.5 | 1.08851177117404E-11 |
| 5 | hspa4a | 5.78070449144882E-16 | 1.1 | 0.55 | 0.27 | 1.48812675723367E-11 |
| 5 | hsp70l | 3.39438168627688E-15 | 1.44 | 0.44 | 0.18 | 8.73815677498257E-11 |
| 5 | map3k15 | 4.80212580801585E-14 | 1.13 | 0.53 | 0.28 | 1.23621124675752E-09 |
| 5 | kctd12.2 | 6.90991831626114E-14 | 1.28 | 0.49 | 0.29 | 1.77882027215511E-09 |
| 5 | ccr9a | 2.46704119261057E-11 | 1.18 | 0.42 | 0.21 | 6.35090414213738E-07 |
| 5 | si:ch211-163c2.2 | 4.47837670721549E-11 | 1.13 | 0.36 | 0.16 | 1.15286851573848E-06 |
| 5 | junbb | 7.93850370918584E-11 | 1.41 | 0.34 | 0.14 | 2.04360900985571E-06 |

| | | | | | | |
|---|---|---|---|---|---|---|
| 5 | zfand6 | 1.05672325906226E-10 | 1.41 | 0.35 | 0.16 | 2.72032268580397E-06 |
| 5 | si:ch211-117m20.5 | 1.99874033040932E-10 | 1.17 | 0.6 | 0.48 | 5.14535723257271E-06 |
| 5 | ahsa1b | 6.23948850219464E-10 | 1.09 | 0.4 | 0.2 | 0.0000160623152511997 |
| 5 | ets2 | 1.26419063694953E-09 | 1.02 | 0.47 | 0.28 | 0.0000325440595669917 |
| 5 | zgc:158343 | 7.83144440699558E-09 | 1.01 | 0.74 | 0.57 | 0.000201604873369287 |
| 5 | s100a10b | 4.19210441710868E-08 | 1.09 | 0.35 | 0.18 | 0.00107917344009629 |
| 5 | deptor | 1.89871525540484E-06 | -1.24 | 0.17 | 0.3 | 0.0488786268198867 |
| 5 | dhrs13b | 1.58792134075513E-06 | -1.13 | 0.13 | 0.27 | 0.0408778590750594 |
| 5 | klhl6 | 1.12282784982101E-07 | -1.24 | 0.17 | 0.33 | 0.00289049573379423 |
| 5 | CABZ01049925.1 | 3.37593737322256E-08 | -1.22 | 0.21 | 0.37 | 0.000869067557988685 |
| 5 | adam19a | 2.75996426363475E-08 | -1.07 | 0.37 | 0.54 | 0.000710497600387493 |
| 5 | ENSDARG00000101094 | 5.31863029441221E-09 | -1.03 | 0.33 | 0.51 | 0.000136917499669054 |
| 5 | hbba2 | 7.68553677065564E-10 | -1.72 | 0.12 | 0.29 | 0.0000197848773086988 |
| 5 | ppp1r12b | 1.93190914970882E-10 | -1.34 | 0.18 | 0.38 | 4.97331372409543E-06 |
| 5 | papss2b | 1.46830838919702E-10 | -1.22 | 0.26 | 0.41 | 3.77986628630989E-06 |
| 5 | si:dkey-11f4.16 | 7.72208535729313E-11 | -1.43 | 0.15 | 0.34 | 1.98789643352797E-06 |
| 5 | mhc1uba | 2.94062853307711E-11 | -1.92 | 0.1 | 0.27 | 7.57006003270039E-07 |
| 5 | ENSDARG00000079078 | 1.73842722749095E-12 | -1.49 | 0.28 | 0.51 | 4.47523321172996E-08 |
| 5 | f11r.1 | 7.85587620849468E-13 | -1.12 | 0.46 | 0.67 | 2.02233821235279E-08 |
| 5 | mmp25b | 1.07546411655357E-13 | -1.08 | 0.46 | 0.66 | 2.76856727524386E-09 |
| 5 | CABZ01001434.1 | 8.04364874577902E-14 | -1.54 | 0.09 | 0.3 | 2.07067649662589E-09 |
| 5 | hopx | 2.21534607313309E-14 | -1.28 | 0.29 | 0.54 | 5.70296539606651E-10 |
| 5 | def8 | 5.46257732495377E-16 | -2,00 | 0.16 | 0.39 | 1.40623128076285E-11 |
| 5 | hpgd | 4.99879507863277E-16 | -1.35 | 0.26 | 0.52 | 1.28683981709244E-11 |
| 5 | si:dkey-32n7.4 | 2.31943601502033E-18 | -1.6 | 0.26 | 0.52 | 5.97092413346683E-14 |
| 5 | zgc:123297 | 1.33636431152723E-20 | -2.52 | 0.08 | 0.33 | 3.44020264716455E-16 |
| 5 | hbba1 | 3.62856312836877E-29 | -2.24 | 0.22 | 0.58 | 9.34101006135973E-25 |
| 5 | CABZ01074309.1 | 5.17389549855678E-32 | -2.27 | 0.15 | 0.53 | 1.33191591819347E-27 |
| 5 | hbaa1 | 5.37778410776584E-34 | -2,00 | 0.34 | 0.71 | 1.38440296286216E-29 |
| 5 | hbba1.1 | 9.80881437442541E-51 | -2.02 | 0.52 | 0.9 | 2.52508308440833E-46 |
| 6 | ins | 4.8994496409157E-43 | 1.99 | 0.59 | 0.13 | 1.26126532106093E-38 |
| 6 | nkl.2 | 3.36346338374917E-17 | 2.05 | 0.26 | 0.06 | 8.65856378878549E-13 |
| 6 | clu | 5.85077987340642E-12 | 1.5 | 0.33 | 0.13 | 1.50616626281101E-07 |
| 6 | icn | 2.25466342005707E-11 | 1.71 | 0.58 | 0.51 | 5.80418004225291E-07 |
| 6 | junbb | 2.6541224688076E-10 | 1.76 | 0.37 | 0.21 | 6.83250747145141E-06 |
| 6 | dhrs1 | 1.31493991442728E-09 | 1.6 | 0.32 | 0.14 | 0.0000338504982171016 |
| 6 | foxp4 | 1.86705179295516E-09 | 1.07 | 0.61 | 0.4 | 0.0000480635143060446 |
| 6 | fkbp5 | 1.89545854198374E-09 | 2.21 | 0.29 | 0.14 | 0.0000487947892462874 |
| 6 | sdk1a | 3.94265614354932E-09 | 1.71 | 0.32 | 0.15 | 0.00010149579710339 |
| 6 | ENSDARG00000078145 | 2.88551046352797E-08 | 1.58 | 0.29 | 0.13 | 0.000742816958626006 |
| 6 | aldob | 3.78342024769494E-08 | 1.93 | 0.31 | 0.2 | 0.000973965874364108 |
| 6 | tox | 3.8189108209655E-08 | 1.03 | 0.32 | 0.15 | 0.00098310221264115 |
| 6 | crim1 | 6.76913573462211E-08 | 1.49 | 0.37 | 0.2 | 0.00174257861216377 |

| | | | | | | |
|---|---|---|---|---|---|---|
| 6 | map1lc3a | 8.02350576960033E-08 | 1.05 | 0.26 | 0.11 | 0.00206549109026821 |
| 6 | CABZ01045617.1 | 1.19984797607616E-07 | 1.28 | 0.71 | 0.69 | 0.00308876864481285 |
| 6 | dip2ca | 1.36589483538507E-07 | 1.48 | 0.36 | 0.19 | 0.00351622307473178 |
| 6 | scg5 | 1.67801596510089E-07 | 1.87 | 0.26 | 0.13 | 0.00431971649895921 |
| 6 | arnt2 | 2.81209670018097E-07 | 1.44 | 0.43 | 0.3 | 0.00723918053527586 |
| 6 | sst2 | 4.25131917455712E-07 | 1.31 | 0.41 | 0.25 | 0.0109441709510624 |
| 6 | gpx4a | 4.99489113374136E-07 | 1.21 | 0.48 | 0.33 | 0.0128583482455904 |
| 6 | cacna2d2a | 6.09102021858905E-07 | 1.39 | 0.25 | 0.18 | 0.0156801133487138 |
| 6 | gipc2 | 0.000000785971275905 | 1.27 | 0.28 | 0.13 | 0.0202332585556224 |
| 6 | cpt1ab | 1.24566292698267E-06 | 1.15 | 0.28 | 0.14 | 0.0320671007293149 |
| 6 | gstk1 | 1.41716783284786E-06 | 1.39 | 0.32 | 0.18 | 0.0364821515210024 |
| 6 | cxcl20 | 1.60323343150446E-06 | 1.45 | 0.26 | 0.13 | 0.0412720382272193 |
| 6 | cbx1b | 1.92702742378309E-06 | -1.2 | 0.18 | 0.3 | 0.0496074669704481 |
| 6 | ndc80 | 1.86165032317143E-06 | -1.16 | 0.16 | 0.31 | 0.0479244642694021 |
| 6 | aurka | 1.80776643075558E-06 | -1.16 | 0.12 | 0.26 | 0.0465373312269409 |
| 6 | armc1l | 1.76291095543829E-06 | -1.16 | 0.16 | 0.31 | 0.045382616725848 |
| 6 | fbxo5 | 1.55943621292778E-06 | -1.45 | 0.12 | 0.26 | 0.0401445664293998 |
| 6 | rasal3 | 0.000001308642745349 | -1.08 | 0.23 | 0.39 | 0.0336883901935193 |
| 6 | cks1b | 1.26484123642557E-06 | -1.09 | 0.19 | 0.34 | 0.0325608079493036 |
| 6 | arhgap11a | 1.02441054493321E-06 | -1.34 | 0.13 | 0.27 | 0.0263714006582156 |
| 6 | arhgef3l | 9.41569247198848E-07 | -1.06 | 0.15 | 0.31 | 0.02423881713064 |
| 6 | stra13 | 9.12740848274615E-07 | -1.11 | 0.25 | 0.41 | 0.0234966876571334 |
| 6 | ptprea | 8.16948545636141E-07 | -1.06 | 0.31 | 0.45 | 0.0210307064103112 |
| 6 | naga | 7.70114066435375E-07 | -1.39 | 0.25 | 0.36 | 0.0198250464122459 |
| 6 | g2e3 | 7.38416682258324E-07 | -1.36 | 0.12 | 0.26 | 0.019009060651376 |
| 6 | si:dkey-108k21.10 | 7.05531915554788E-07 | -1.08 | 0.13 | 0.27 | 0.0181625081021269 |
| 6 | snrpg | 6.57382313645984E-07 | -1.09 | 0.34 | 0.47 | 0.0169229929001886 |
| 6 | e2f8 | 5.60602153137577E-07 | -1.24 | 0.13 | 0.27 | 0.0144315812282207 |
| 6 | skap1 | 5.24977129933216E-07 | -1.33 | 0.15 | 0.28 | 0.0135144862558708 |
| 6 | FP085398.1 | 5.02751638275235E-07 | -1.26 | 0.13 | 0.28 | 0.0129423354241194 |
| 6 | tbc1d10c | 4.97893628575515E-07 | -1.63 | 0.12 | 0.26 | 0.0128172756804195 |
| 6 | gnpda1 | 4.37757825143163E-07 | -1.57 | 0.16 | 0.28 | 0.0112691996926605 |
| 6 | mapre1a | 4.16320153563471E-07 | -1.01 | 0.35 | 0.53 | 0.0107173297131844 |
| 6 | cdca5 | 2.8122598605568E-07 | -1.23 | 0.16 | 0.31 | 0.00723960055903137 |
| 6 | ypel5 | 2.70537859006354E-07 | -1.2 | 0.3 | 0.45 | 0.00696445610440057 |
| 6 | mhc1zea | 2.6290451738838E-07 | -1.04 | 0.22 | 0.39 | 0.00676795099112906 |
| 6 | plk1 | 2.49966063125933E-07 | -1.3 | 0.14 | 0.29 | 0.00643487636305089 |
| 6 | tuba1b | 2.46294261145701E-07 | -1.54 | 0.14 | 0.29 | 0.00634035316467378 |
| 6 | si:dkeyp-68b7.12 | 2.40976177257346E-07 | -1.05 | 0.28 | 0.45 | 0.00620344973113585 |
| 6 | si:dkey-109a10.2 | 2.25498727712624E-07 | -1.67 | 0.13 | 0.26 | 0.00580501374750608 |
| 6 | gmip | 2.14901584342971E-07 | -1.14 | 0.25 | 0.4 | 0.0055322114857411 |
| 6 | capgb | 2.1429631302831E-07 | -1.15 | 0.3 | 0.42 | 0.00551662998628779 |
| 6 | tacc3 | 2.06787969470306E-07 | -1.21 | 0.14 | 0.29 | 0.00532334269807408 |

| | | | | | | |
|---|---|---|---|---|---|---|
| 6 | nusap1 | 0.0000001998572287879 | -1.15 | 0.18 | 0.35 | 0.00514492464068692 |
| 6 | grn1 | 1.80894467347991E-07 | -1.48 | 0.21 | 0.39 | 0.00465676627293933 |
| 6 | ndufs2 | 1.49195160040852E-07 | -1.04 | 0.35 | 0.49 | 0.00384073100493167 |
| 6 | cdc20 | 1.37581240174772E-07 | -1.21 | 0.13 | 0.28 | 0.00354175386581916 |
| 6 | aurkb | 1.04402350385218E-07 | -1.25 | 0.18 | 0.35 | 0.00268762970596666 |
| 6 | si:ch211-165d12.4 | 1.04178712821215E-07 | -1.01 | 0.33 | 0.5 | 0.00268187260415655 |
| 6 | vrk1 | 7.54845593180134E-08 | -1.08 | 0.23 | 0.4 | 0.00194319901052362 |
| 6 | dock10 | 7.05105168787847E-08 | -1.1 | 0.21 | 0.38 | 0.00181515223601055 |
| 6 | tuba8l | 5.6590691262699E-08 | -1.19 | 0.41 | 0.57 | 0.00145681416517566 |
| 6 | si:ch73-40i7.2 | 5.38124935541478E-08 | -1.63 | 0.17 | 0.28 | 0.00138529502156443 |
| 6 | anln | 4.45940107483239E-08 | -1.69 | 0.12 | 0.28 | 0.0011479836186941 |
| 6 | ankrd44 | 3.84071353908259E-08 | -1.2 | 0.22 | 0.4 | 0.000988714886366031 |
| 6 | zgc:171506 | 3.27187387070906E-08 | -1.14 | 0.11 | 0.26 | 0.000842278490536634 |
| 6 | abi3a | 2.83552138496202E-08 | -1,00 | 0.16 | 0.33 | 0.000729948270130773 |
| 6 | kif23 | 2.73939132253744E-08 | -1.21 | 0.13 | 0.29 | 0.000705201508160812 |
| 6 | rpa2 | 2.66099627664697E-08 | -1.2 | 0.21 | 0.37 | 0.000685020271497229 |
| 6 | ENSDARG00000101094 | 2.46312584635228E-08 | -1.17 | 0.2 | 0.38 | 0.000634082486626467 |
| 6 | hmha1b | 2.4109664793216E-08 | -1.15 | 0.24 | 0.42 | 0.00062065510077176 |
| 6 | spdl1 | 2.22701639332171E-08 | -1.31 | 0.13 | 0.29 | 0.000573300830132808 |
| 6 | wu:fb44b02 | 1.93611885837276E-08 | -1.37 | 0.14 | 0.31 | 0.0004984150777109 |
| 6 | chaf1a | 1.76227374090576E-08 | -1.12 | 0.24 | 0.43 | 0.00045366212912137 |
| 6 | krtcap2 | 1.47457312872106E-08 | -1.19 | 0.12 | 0.28 | 0.000379599360526663 |
| 6 | birc5a | 1.37364568637266E-08 | -1.43 | 0.15 | 0.31 | 0.000353617609042915 |
| 6 | ncapg | 1.35510581180869E-08 | -1.09 | 0.17 | 0.34 | 0.000348844889133912 |
| 6 | kifc1 | 1.33766170506048E-08 | -1.24 | 0.15 | 0.32 | 0.000344354252733719 |
| 6 | arhgap15 | 1.32212973322911E-08 | -1.09 | 0.29 | 0.47 | 0.00034035585722517 |
| 6 | dlgap5 | 1.27376579538558E-08 | -1.26 | 0.19 | 0.36 | 0.00032790552870611 |
| 6 | ptges3b | 1.11437440071364E-08 | -1.2 | 0.24 | 0.4 | 0.000286873401975711 |
| 6 | trpv1 | 1.04953137450879E-08 | -1.52 | 0.13 | 0.29 | 0.000270180861739798 |
| 6 | kcnab2a | 9.44053423242867E-09 | -1.24 | 0.25 | 0.41 | 0.000243027672745411 |
| 6 | si:ch211-210g13.5 | 6.4518405468011E-09 | -1.54 | 0.17 | 0.34 | 0.000166089731196301 |
| 6 | psmd10 | 5.75713598841856E-09 | -1.57 | 0.13 | 0.27 | 0.000148205951749859 |
| 6 | il12rb2l | 5.26063752909107E-09 | -1.45 | 0.2 | 0.38 | 0.000135424591911391 |
| 6 | tpx2 | 4.8492844264902E-09 | -1.42 | 0.17 | 0.35 | 0.000124835128991137 |
| 6 | prr11 | 3.9424897479974E-09 | -1.61 | 0.1 | 0.26 | 0.000101491513582697 |
| 6 | ccna2 | 3.57728934737496E-09 | -1.08 | 0.18 | 0.37 | 0.0000920901596694736 |
| 6 | ttk | 3.09872031795354E-09 | -1.45 | 0.15 | 0.32 | 0.000079770357145078 |
| 6 | bub3 | 2.73194640456923E-09 | -1.1 | 0.24 | 0.44 | 0.0000703284962928257 |
| 6 | cenpe | 2.46618161260029E-09 | -1.73 | 0.13 | 0.29 | 0.0000634869132531693 |
| 6 | cdk1 | 2.04362792944869E-09 | -1.36 | 0.18 | 0.35 | 0.0000526091137877977 |
| 6 | CR847895.3 | 2.03839808635692E-09 | -1.15 | 0.32 | 0.5 | 0.0000524744819370862 |
| 6 | h2afx | 1.76865933514111E-09 | -1.53 | 0.17 | 0.35 | 0.0000455305972645376 |
| 6 | aspm | 1.23817208870529E-09 | -1.38 | 0.15 | 0.33 | 0.0000318742640795404 |

| | | | | | | |
|---|---|---|---|---|---|---|
| 6 | wasa | 1.23102855728203E-09 | -1.11 | 0.27 | 0.47 | 0.0000316903681501114 |
| 6 | si:dkey-171c9.3 | 9.88029422012763E-10 | -1.38 | 0.23 | 0.42 | 0.0000254348414108746 |
| 6 | cenpf | 7.69042628990438E-10 | -1.76 | 0.18 | 0.33 | 0.0000197974643981008 |
| 6 | mad2l1 | 7.17256314646317E-10 | -1.24 | 0.18 | 0.37 | 0.0000184643293079401 |
| 6 | smc2 | 6.24511215066891E-10 | -1.38 | 0.15 | 0.34 | 0.000016076792209467 |
| 6 | ncapd2 | 5.14905606234825E-10 | -1.74 | 0.13 | 0.3 | 0.0000132552150213031 |
| 6 | psme2 | 4.67885092745491E-10 | -1.04 | 0.39 | 0.55 | 0.0000120447659425472 |
| 6 | si:dkey-25o16.4 | 4.04814840378131E-10 | -2.03 | 0.1 | 0.25 | 0.0000104211484358542 |
| 6 | fam49ba | 2.63005172838082E-10 | -1.3 | 0.31 | 0.51 | 6.77054216437075E-06 |
| 6 | ms4a17a.6 | 2.47120166851751E-10 | -1.63 | 0.19 | 0.36 | 6.36161445526463E-06 |
| 6 | top2a | 2.24744650371963E-10 | -1.59 | 0.18 | 0.37 | 5.78560153452545E-06 |
| 6 | CABZ01058261.1 | 1.95666603876955E-10 | -1.35 | 0.22 | 0.4 | 5.03704538360446E-06 |
| 6 | NCKAP1L | 5.27025981164212E-11 | -1.3 | 0.22 | 0.4 | 1.35672298331103E-06 |
| 6 | lcp1 | 4.26584338216153E-11 | -1.14 | 0.38 | 0.58 | 1.09815606186984E-06 |
| 6 | lbr | 4.14847000246614E-11 | -1.1 | 0.29 | 0.51 | 1.06794063273486E-06 |
| 6 | ptpn6 | 3.601567575722E-11 | -1.19 | 0.35 | 0.56 | 9.27151541018116E-07 |
| 6 | arhgdig | 2.5494849843627E-11 | -1.04 | 0.46 | 0.65 | 6.56313919524489E-07 |
| 6 | msna | 1.93918935836296E-11 | -1.07 | 0.52 | 0.68 | 4.99205516523377E-07 |
| 6 | hmgb2a | 1.23883137795413E-11 | -1.21 | 0.66 | 0.78 | 3.18912361626733E-07 |
| 6 | txn | 1.15616669701074E-11 | -1.46 | 0.2 | 0.42 | 2.97631992811474E-07 |
| 6 | coro1a | 9.08039883271857E-12 | -1.28 | 0.39 | 0.6 | 2.33756707150674E-07 |
| 6 | tapbp.1 | 8.96351803395174E-12 | -1.2 | 0.27 | 0.48 | 2.3074784474802E-07 |
| 6 | mhc1uba | 6.64144396767409E-12 | -1.21 | 0.22 | 0.44 | 1.70970692059834E-07 |
| 6 | si:ch211-181d7.3 | 5.80730340595597E-12 | -1.44 | 0.09 | 0.27 | 1.49497411579525E-07 |
| 6 | ube2c | 5.32762701062419E-12 | -1.49 | 0.19 | 0.41 | 1.37149102134498E-07 |
| 6 | gig2i | 4.20505049980771E-12 | -1.87 | 0.12 | 0.31 | 1.0825061501655E-07 |
| 6 | BX005223.1 | 3.69804873419398E-12 | -2.04 | 0.1 | 0.27 | 9.51988685643556E-08 |
| 6 | mki67 | 2.73618676000402E-12 | -1.7 | 0.21 | 0.42 | 7.04376557627836E-08 |
| 6 | hbba2 | 1.80477413542368E-12 | -1.56 | 0.1 | 0.3 | 4.64603005682119E-08 |
| 6 | arhgap4a | 7.63748865538481E-13 | -1.31 | 0.13 | 0.35 | 1.96611870455571E-08 |
| 6 | hmgb2b | 2.37145508848393E-13 | -1.07 | 0.52 | 0.68 | 6.10483683428418E-09 |
| 6 | rac2 | 1.19695072802872E-13 | -1.2 | 0.41 | 0.59 | 3.08131025916433E-09 |
| 6 | h2afvb | 6.02300451070032E-14 | -1,00 | 0.55 | 0.71 | 1.55050205118958E-09 |
| 6 | si:ch211-119e14.1 | 1.22956690755949E-14 | -1.65 | 0.3 | 0.51 | 3.16527409013041E-10 |
| 6 | zgc:103700 | 8.76822889103901E-15 | -2.06 | 0.18 | 0.4 | 2.25720516342017E-10 |
| 6 | gtf2f2a | 7.05652658575097E-15 | -1.14 | 0.34 | 0.58 | 1.81656163896987E-10 |
| 6 | pycard | 4.61716836246225E-15 | -1.7 | 0.23 | 0.45 | 1.18859765154866E-10 |
| 6 | mhc2dab | 3.78505098577102E-15 | -2.84 | 0.09 | 0.28 | 9.74385675267032E-11 |
| 6 | grn2 | 2.11301171574339E-15 | -2.84 | 0.07 | 0.27 | 5.43952605983821E-11 |
| 6 | hmgn2 | 1.08090838920685E-16 | -1.37 | 0.71 | 0.85 | 2.7825824663352E-12 |
| 6 | hbba1 | 9.98127757450079E-19 | -1.81 | 0.24 | 0.55 | 2.56948028600374E-14 |
| 6 | hbaa1 | 6.62843482772327E-20 | -1.52 | 0.38 | 0.69 | 1.7063579777008E-15 |
| 6 | hbba1.1 | 1.42782832820175E-28 | -1.84 | 0.58 | 0.9 | 3.67565846528977E-24 |

| | | | | | | |
|---|---|---|---|---|---|---|
| 6 | CABZ01074309.1 | 1.81746533031735E-30 | -1.8 | 0.25 | 0.62 | 4.67870099983596E-26 |
| 6 | rrm2.1 | 2.22433506805022E-31 | -2.65 | 0.06 | 0.38 | 5.72610576568168E-27 |
| 6 | si:busm1-48c11.3 | 4.35053047327087E-58 | -8.64 | 0,00 | 0.38 | 1.11995705973412E-53 |
| 6 | si:busm1-194e12.12 | 6.88477316661383E-60 | -8.86 | 0,00 | 0.39 | 1.7723471562814E-55 |
| 7 | ins | 1.40013637261131E-31 | 5.04 | 0.52 | 0.05 | 3.60437106401329E-27 |
| 7 | CABZ01045617.1 | 2.20325274582542E-22 | 2.19 | 0.72 | 0.47 | 5.67183354357837E-18 |
| 7 | camk2d1 | 3.05450251033089E-09 | 2.29 | 0.34 | 0.11 | 0.000078632058123448 |
| 7 | si:dkey-53k12.33 | 1.80201474337422E-08 | 1.44 | 0.46 | 0.31 | 0.000463892655386825 |
| 7 | ENSDARG00000079078 | 8.2440023064304E-07 | -1.06 | 0.35 | 0.59 | 0.0212225351374438 |
| 7 | si:dkey-11f4.16 | 3.59188793457843E-07 | -1.55 | 0.09 | 0.29 | 0.00924659710998525 |
| 7 | hbba2 | 1.3476021725638E-07 | -1.68 | 0.11 | 0.33 | 0.003469132272831 |
| 7 | irf4b | 2.32065001747855E-08 | -1.18 | 0.21 | 0.48 | 0.000597404933999502 |
| 7 | mhc2dab | 1.08000362432929E-08 | -1.21 | 0.1 | 0.33 | 0.000278025333011089 |
| 7 | ier2a | 2.95433448225219E-09 | -1.18 | 0.23 | 0.52 | 0.0000760534325766182 |
| 7 | mych | 2.22876921465855E-09 | -1.36 | 0.32 | 0.62 | 0.000057375205892955 |
| 7 | egr3 | 2.15684560948756E-09 | -1.29 | 0.15 | 0.42 | 0.0000555236765250382 |
| 7 | ENSDARG00000103577 | 2.48653078940227E-10 | -2.09 | 0.08 | 0.34 | 6.40107621115827E-06 |
| 7 | igl3v5 | 1.36896033471777E-10 | -2.1 | 0.78 | 0.92 | 3.52411458966396E-06 |
| 7 | cd83 | 6.07935022188708E-11 | -1.84 | 0.05 | 0.29 | 1.56500712762039E-06 |
| 7 | CABZ01074309.1 | 5.69171631520897E-12 | -1.39 | 0.16 | 0.48 | 1.46521853102424E-07 |
| 7 | gtf2f2a | 4.14916995086655E-12 | -1.9 | 0.07 | 0.34 | 1.06812082045158E-07 |
| 7 | fosab | 2.95079658940709E-12 | -1.21 | 0.35 | 0.69 | 7.59623566011068E-08 |
| 7 | zgc:103700 | 3.65233929319521E-13 | -1.74 | 0.25 | 0.59 | 9.40221704247244E-09 |
| 7 | hbba1 | 1.88969552161213E-13 | -2.07 | 0.2 | 0.56 | 4.86464318128612E-09 |
| 7 | hbaa1 | 2.987411835147E-19 | -1.39 | 0.33 | 0.76 | 7.69049428721893E-15 |
| 7 | hbba1.1 | 2.31658581605945E-19 | -1.12 | 0.53 | 0.92 | 5.96358686628185E-15 |
| 7 | si:busm1-194e12.12 | 2.6418713605896E-42 | -8.88 | 0,00 | 0.52 | 6.8009694435658E-38 |
| 7 | si:busm1-48c11.3 | 6.66332990244629E-57 | -10.26 | 0,00 | 0.65 | 1.71534101678675E-52 |
| 8 | ins | 3.01145097215863E-24 | 3.31 | 0.54 | 0.08 | 7.75237823762796E-20 |
| 8 | ddx6 | 9.37074134410098E-09 | 2.11 | 0.31 | 0.14 | 0.000241230994421192 |
| 8 | eef2l2 | 1.0130386714257E-07 | 1.72 | 0.49 | 0.28 | 0.00260786545185118 |
| 8 | fkbp5 | 1.03824285751617E-07 | 1.28 | 0.33 | 0.1 | 0.00267274858810386 |
| 8 | lrrc7 | 3.70218277590524E-08 | -2.22 | 0.08 | 0.34 | 0.000953052912001286 |
| 9 | nptna | 3.84464051246836E-16 | 1.38 | 0.73 | 0.24 | 9.8972580712473E-12 |
| 9 | ins | 1.61201386459456E-12 | 3.34 | 0.42 | 0.06 | 4.14980729162578E-08 |
| 9 | ctsba | 5.78612498123006E-09 | 4.21 | 0.27 | 0.02 | 0.000148952215391805 |
| 9 | pla2g15 | 8.33377292013773E-09 | 1.8 | 0.35 | 0.07 | 0.000214536316283106 |
| 9 | nkl.2 | 5.56422066819745E-08 | 7.22 | 0.35 | 0.09 | 0.00143239732661407 |
| 9 | hsp70.1 | 3.44542757726826E-07 | 3.31 | 0.39 | 0.12 | 0.00886956421216167 |
| 9 | si:dkey-28k24.2.1 | 3.80287367154211E-07 | 1.26 | 0.72 | 0.42 | 0.00978973769265086 |
| 9 | rnf130 | 1.81374208935752E-06 | -1.18 | 0.55 | 0.77 | 0.0466911626063306 |
| 9 | hbba1 | 1.20476684321883E-08 | -2.22 | 0.16 | 0.47 | 0.000310143128449824 |
| 9 | mhc2dab | 2.12560725845658E-10 | -2.27 | 0.13 | 0.47 | 5.47195076544476E-06 |

| | | | | | | |
|---|---|---|---|---|---|---|
| 9 | hbaa1 | 5.73729703389831E-11 | -1.64 | 0.32 | 0.69 | 1.47695237543644E-06 |
| 9 | CABZ01074309.1 | 8.64833537968792E-12 | -2.05 | 0.28 | 0.6 | 2.22634097679306E-07 |
| 9 | efna2a | 7.91595012766417E-14 | -4.09 | 0.05 | 0.39 | 2.03780304136459E-09 |
| 9 | hbba1.1 | 5.89138753280008E-16 | -1.62 | 0.48 | 0.86 | 1.51661989256872E-11 |
| 9 | zgc:103700 | 1.94775039558224E-18 | -2.06 | 0.38 | 0.77 | 5.01409384334737E-14 |
| 9 | si:busm1-48c11.3 | 6.70291576044494E-25 | -8.09 | 0,00 | 0.42 | 1.72553160421134E-20 |
| 9 | si:busm1-194e12.12 | 1.15803691575964E-30 | -9.01 | 0,00 | 0.5 | 2.98113443224003E-26 |
| 10 | cd81a | 3.01713727489131E-17 | 3.68 | 0.67 | 0.11 | 7.76701648675271E-13 |
| 10 | MFAP4 (1 of many).7 | 1.82194776090068E-12 | 2.33 | 0.72 | 0.24 | 4.69024012088663E-08 |
| 10 | ca2 | 7.21037502952861E-11 | 1.8 | 0.68 | 0.21 | 1.85616684385155E-06 |
| 10 | ins | 1.87221933534388E-09 | 3.31 | 0.44 | 0.06 | 0.0000481965423497575 |
| 10 | MFAP4 (1 of many).6 | 2.91812137036496E-08 | 1.31 | 0.91 | 0.75 | 0.000751211984373051 |
| 10 | CR388373.3 | 1.59479819042634E-06 | 3.16 | 0.4 | 0.11 | 0.0410548898161454 |
| 10 | ppdpfa | 4.75508140839802E-07 | -2.49 | 0.12 | 0.42 | 0.012241006069639 |
| 10 | hbba2 | 4.05192800182955E-07 | -3.11 | 0.05 | 0.33 | 0.0104308782551098 |
| 10 | MFAP4 (1 of many) | 6.36411493701977E-08 | -1.04 | 0.85 | 0.96 | 0.001638314108237 |
| 10 | hbba1 | 2.70163915343732E-08 | -2.64 | 0.17 | 0.44 | 0.000695482967269369 |
| 10 | CABZ01074309.1 | 1.57244972843484E-08 | -2.25 | 0.18 | 0.52 | 0.000404795733590981 |
| 10 | alox5ap | 1.18492059100202E-09 | -1.68 | 0.3 | 0.7 | 0.000030503410774165 |
| 10 | hbaa1 | 5.70822708783753E-10 | -2.03 | 0.31 | 0.72 | 0.0000146946889922202 |
| 10 | si:dkey-11f4.16 | 1.7324062161825E-10 | -2.58 | 0.26 | 0.62 | 4.45973332231861E-06 |
| 10 | bzw1b | 1.77435365745164E-12 | -1.67 | 0.54 | 0.74 | 4.56771862037775E-08 |
| 10 | hbba1.1 | 1.3796892205621E-14 | -1.9 | 0.56 | 0.88 | 3.55173396049302E-10 |
| 10 | igsf21a | 4.58839980527926E-15 | -4.89 | 0.03 | 0.47 | 1.18119176187304E-10 |
| 11 | hsp70.2 | 1.4995895780919E-13 | 3.98 | 0.62 | 0.08 | 3.86039345088198E-09 |
| 11 | CABZ01040076.1 | 3.47495788915621E-12 | 6.62 | 0.39 | 0,00 | 8.94558409405482E-08 |
| 11 | cnksr2a | 1.68629262814983E-11 | 7.56 | 0.37 | 0,00 | 4.34102311264611E-07 |
| 11 | hsp70.1 | 8.98420397057335E-11 | 4.2 | 0.52 | 0.1 | 0.0000023128036281447 |
| 11 | hsp70l | 9.59962972617986E-11 | 3.18 | 0.64 | 0.17 | 2.47123268041048E-06 |
| 11 | hspa4a | 1.29012646036629E-10 | 2.41 | 0.75 | 0.26 | 3.32117254692094E-06 |
| 11 | si:busm1-266f07.2 | 3.21442202092153E-10 | 1.71 | 0.98 | 0.92 | 0.0000082748866084583 |
| 11 | ins | 6.72616225286134E-10 | 3.03 | 0.51 | 0.07 | 0.000017315159487541 |
| 11 | si:dkey-78o7.3 | 4.79415914157551E-09 | 2.28 | 0.74 | 0.32 | 0.000123416038781578 |
| 11 | hsp90aa1.2 | 1.87758995287892E-08 | 3.78 | 0.53 | 0.14 | 0.000483347981569619 |
| 11 | ccr9a | 1.7552187430365E-07 | 1.56 | 0.84 | 0.44 | 0.00451845961019885 |
| 11 | hsp70.3 | 1.81562037601373E-07 | 4.04 | 0.43 | 0.08 | 0.00467395153397214 |
| 11 | hspbp1 | 1.95971049357146E-07 | 4.37 | 0.37 | 0.04 | 0.00504488272360101 |
| 11 | si:ch211-209n20.3 | 1.07195172395635E-06 | 3.87 | 0.32 | 0.03 | 0.0275952532298084 |
| 11 | mhc2dab | 7.92811494439059E-07 | -2.07 | 0.33 | 0.63 | 0.0204093463013447 |
| 11 | hbba1 | 2.72892354410667E-07 | -2.3 | 0.25 | 0.63 | 0.00702506787959379 |
| 11 | CR385050.1 | 2.36355054097814E-08 | -2.9 | 0.11 | 0.5 | 0.000608448815764003 |
| 11 | hbaa1 | 1.50658920108603E-08 | -1.96 | 0.4 | 0.81 | 0.000387841258035578 |
| 11 | CABZ01074309.1 | 1.2527592227976E-09 | -2.58 | 0.25 | 0.67 | 0.0000322497806724787 |

| | | | | | | |
|---|---|---|---|---|---|---|
| 11 | hbba1.1 | 1.34669440152829E-13 | -2.31 | 0.51 | 0.92 | 3.46679539785427E-09 |
| 11 | zgc:113229 | 4.94983815365434E-14 | -9.16 | 0,00 | 0.4 | 1.27423683589524E-09 |
| 11 | si:busm1-48c11.3 | 1.97498959924785E-23 | -8.74 | 0,00 | 0.63 | 5.08421572534374E-19 |
| 11 | si:busm1-194e12.12 | 3.10636577089248E-29 | -9.5 | 0,00 | 0.74 | 7.99671740400852E-25 |
| 12 | CR847895.3 | 1.17387310600681E-07 | 2.06 | 0.6 | 0.13 | 0.00302190153679332 |
| 12 | sh3gl1a | 4.28788733505134E-07 | -1.77 | 0.6 | 0.73 | 0.0110383083666227 |
| 12 | mt-co3 | 7.59939751253382E-08 | -1.27 | 0.96 | 1,00 | 0.00195631290165158 |
| 12 | si:ch73-170d6.3 | 5.23339506806244E-08 | -4.41 | 0.01 | 0.31 | 0.00134723289237131 |
| 12 | CABZ01074309.1 | 1.25128631134789E-08 | -2.19 | 0.3 | 0.73 | 0.000322118635130288 |
| 12 | hbaa1 | 6.19964308478979E-09 | -3,00 | 0.24 | 0.69 | 0.000159597411931744 |
| 12 | hbba1.1 | 1.53699935503664E-12 | -2.74 | 0.36 | 0.89 | 3.95669743967083E-08 |
| 12 | ptprfb | 6.13583923580903E-13 | -9.17 | 0,00 | 0.4 | 1.57954909447432E-08 |
| 13 | hbaa1 | 1.62282717267693E-07 | -4.59 | 0.36 | 0.88 | 0.00417764399062222 |
| 13 | gstp1 | 5.41961021259991E-08 | -2,00 | 0.64 | 0.96 | 0.00139517025702959 |
| 13 | hbba1.1 | 4.2526552203053E-08 | -4.09 | 0.54 | 0.96 | 0.00109476103336319 |
| 14 | hbba1.1 | 2.66757708811558E-07 | -2.63 | 0.44 | 1,00 | 0.00686714369793594 |

p_val, p-value; avg_logFC, log2 fold-change of the average expression between group 1 and group 2; pct.1, fraction of cells that express the gene in group 1 (z3'-DpE-/-); pct.2 fraction of cells that express the gene in group 2 (z3'-DpE+/+); p_val_adj, q-value

**Supplementary Table 4. GO enrichment of GREAT-associated genes from chromatin regions losing accessibility in z3′-DpE-/- acinar cells (adult ATAC-seq, Down regions).**

Source: ATAC-seq of FACS-purified fixed mCherry+ adult acinar cells from *Tg(ela:mCherry)* zebrafish (*Danio rerio*), comparing z3′-DpE+/+ and z3′-DpE-/-. Criteria: Differential accessibility (z3′-DpE-/- vs z3′-DpE+/+) defined as FDR ≤ 0.05 and |log2FC| ≥ 1 (edgeR); Down regions defined as peaks with decreased accessibility in z3′-DpE-/- vs z3′-DpE+/+ (log2FC < -1). Peaks were assigned to putative target genes using GREAT (peak-to-gene association). GO Biological Process enrichment performed using the PANTHER Overrepresentation Test (*Danio rerio* reference list; Fisher's exact test with FDR correction).

| GO term (biological process) | GO ID | Ref list count (N=26353) | Query list count (n=3317) | Expected | Fold enrichment | Raw P-value | FDR |
|---|---|---|---|---|---|---|---|
| insulin secretion involved in cellular response to glucose stimulus | GO:0035773 | 3 | 3 | 0.38 | 7.94 | 1.99E-03 | 2.32E-02 |
| vagus nerve development | GO:0021564 | 3 | 3 | 0.38 | 7.94 | 1.99E-03 | 2.32E-02 |
| late distal convoluted tubule development | GO:0072068 | 3 | 3 | 0.38 | 7.94 | 1.99E-03 | 2.32E-02 |
| metaphase/anaphase transition of mitotic cell cycle | GO:0007091 | 3 | 3 | 0.38 | 7.94 | 1.99E-03 | 2.31E-02 |
| distal convoluted tubule development | GO:0072025 | 3 | 3 | 0.38 | 7.94 | 1.99E-03 | 2.31E-02 |
| negative regulation of synapse assembly | GO:0051964 | 3 | 3 | 0.38 | 7.94 | 1.99E-03 | 2.31E-02 |
| selective angioblast sprouting | GO:0035474 | 3 | 3 | 0.38 | 7.94 | 1.99E-03 | 2.31E-02 |
| metaphase/anaphase transition of cell cycle | GO:0044784 | 3 | 3 | 0.38 | 7.94 | 1.99E-03 | 2.30E-02 |
| regulation of Wnt signaling pathway, calcium modulating pathway | GO:0008591 | 3 | 3 | 0.38 | 7.94 | 1.99E-03 | 2.30E-02 |
| regulation of nuclear-transcribed mRNA catabolic process, nonsense-mediated decay | GO:2000622 | 3 | 3 | 0.38 | 7.94 | 1.99E-03 | 2.30E-02 |
| smooth muscle cell proliferation | GO:0048659 | 3 | 3 | 0.38 | 7.94 | 1.99E-03 | 2.29E-02 |
| DNA ligation involved in DNA repair | GO:0051103 | 3 | 3 | 0.38 | 7.94 | 1.99E-03 | 2.29E-02 |
| negative regulation of synapse organization | GO:1905809 | 3 | 3 | 0.38 | 7.94 | 1.99E-03 | 2.29E-02 |
| rhombomere 6 development | GO:0021572 | 4 | 4 | 0.50 | 7.94 | 2.51E-04 | 3.92E-03 |
| rhombomere 5 development | GO:0021571 | 4 | 4 | 0.50 | 7.94 | 2.51E-04 | 3.92E-03 |
| motor behavior | GO:0061744 | 4 | 4 | 0.50 | 7.94 | 2.51E-04 | 3.91E-03 |
| neural plate morphogenesis | GO:0001839 | 7 | 6 | 0.88 | 6.81 | 2.47E-05 | 5.37E-04 |
| locus ceruleus development | GO:0021703 | 6 | 5 | 0.76 | 6.62 | 1.69E-04 | 2.83E-03 |
| intracellular sphingolipid homeostasis | GO:0090156 | 5 | 4 | 0.63 | 6.36 | 1.13E-03 | 1.43E-02 |
| regulation of Wnt signaling pathway involved in dorsal/ventral axis specification | GO:2000053 | 5 | 4 | 0.63 | 6.36 | 1.13E-03 | 1.43E-02 |
| glial cell fate commitment | GO:0021781 | 5 | 4 | 0.63 | 6.36 | 1.13E-03 | 1.43E-02 |
| oligodendrocyte cell fate commitment | GO:0021779 | 5 | 4 | 0.63 | 6.36 | 1.13E-03 | 1.43E-02 |
| renal system pattern specification | GO:0072048 | 8 | 6 | 1.01 | 5.96 | 8.83E-05 | 1.65E-03 |
| distal tubule development | GO:0072017 | 8 | 6 | 1.01 | 5.96 | 8.83E-05 | 1.65E-03 |

| | | | | | | | |
|---|---|---|---|---|---|---|---|
| negative regulation of endodermal cell fate specification | GO:0042664 | 8 | 6 | 1.01 | 5.96 | 8.83E-05 | 1.64E-03 |
| motor neuron migration | GO:0097475 | 8 | 6 | 1.01 | 5.96 | 8.83E-05 | 1.64E-03 |
| proximal/distal pattern formation | GO:0009954 | 8 | 6 | 1.01 | 5.96 | 8.83E-05 | 1.64E-03 |
| insulin secretion | GO:0030073 | 8 | 6 | 1.01 | 5.96 | 8.83E-05 | 1.63E-03 |
| pattern specification involved in kidney development | GO:0061004 | 8 | 6 | 1.01 | 5.96 | 8.83E-05 | 1.63E-03 |
| pattern specification involved in pronephros development | GO:0039017 | 8 | 6 | 1.01 | 5.96 | 8.83E-05 | 1.63E-03 |
| synaptic transmission, GABAergic | GO:0051932 | 7 | 5 | 0.88 | 5.67 | 5.31E-04 | 7.50E-03 |
| coronary vasculature development | GO:0060976 | 7 | 5 | 0.88 | 5.67 | 5.31E-04 | 7.48E-03 |
| pronephric distal tubule development | GO:0035777 | 6 | 4 | 0.76 | 5.30 | 3.04E-03 | 3.30E-02 |
| anterior/posterior pattern specification involved in kidney development | GO:0072098 | 6 | 4 | 0.76 | 5.30 | 3.04E-03 | 3.30E-02 |
| peptidyl-histidine modification | GO:0018202 | 6 | 4 | 0.76 | 5.30 | 3.04E-03 | 3.30E-02 |
| regulation of gene silencing by regulatory ncRNA | GO:0060966 | 6 | 4 | 0.76 | 5.30 | 3.04E-03 | 3.29E-02 |
| anterior/posterior pattern specification involved in pronephros development | GO:0034672 | 6 | 4 | 0.76 | 5.30 | 3.04E-03 | 3.29E-02 |
| positive regulation of peptide hormone secretion | GO:0090277 | 6 | 4 | 0.76 | 5.30 | 3.04E-03 | 3.28E-02 |
| regulation of retinal ganglion cell axon guidance | GO:0090259 | 6 | 4 | 0.76 | 5.30 | 3.04E-03 | 3.28E-02 |
| regulation of muscle organ development | GO:0048634 | 6 | 4 | 0.76 | 5.30 | 3.04E-03 | 3.28E-02 |
| pineal gland development | GO:0021982 | 6 | 4 | 0.76 | 5.30 | 3.04E-03 | 3.27E-02 |
| positive regulation of peptide secretion | GO:0002793 | 6 | 4 | 0.76 | 5.30 | 3.04E-03 | 3.27E-02 |
| protein histidyl modification to diphthamide | GO:0017183 | 6 | 4 | 0.76 | 5.30 | 3.04E-03 | 3.26E-02 |
| neuron cell-cell adhesion | GO:0007158 | 9 | 6 | 1.13 | 5.30 | 2.37E-04 | 3.72E-03 |
| negative regulation of endodermal cell differentiation | GO:1903225 | 9 | 6 | 1.13 | 5.30 | 2.37E-04 | 3.72E-03 |
| regulation of non-canonical Wnt signaling pathway | GO:2000050 | 17 | 11 | 2.14 | 5.14 | 7.37E-07 | 2.29E-05 |
| urogenital system development | GO:0001655 | 14 | 9 | 1.76 | 5.11 | 8.65E-06 | 2.06E-04 |
| hormone secretion | GO:0046879 | 11 | 7 | 1.38 | 5.06 | 1.03E-04 | 1.87E-03 |
| peptide hormone secretion | GO:0030072 | 11 | 7 | 1.38 | 5.06 | 1.03E-04 | 1.86E-03 |
| peptide secretion | GO:0002790 | 11 | 7 | 1.38 | 5.06 | 1.03E-04 | 1.86E-03 |
| gamma-aminobutyric acid signaling pathway | GO:0007214 | 16 | 10 | 2.01 | 4.97 | 3.83E-06 | 1.03E-04 |
| angioblast cell migration | GO:0035476 | 16 | 10 | 2.01 | 4.97 | 3.83E-06 | 1.02E-04 |
| DNA topological change | GO:0006265 | 8 | 5 | 1.01 | 4.97 | 1.27E-03 | 1.58E-02 |
| venous blood vessel morphogenesis | GO:0048845 | 8 | 5 | 1.01 | 4.97 | 1.27E-03 | 1.58E-02 |
| hypothalamus cell differentiation | GO:0021979 | 8 | 5 | 1.01 | 4.97 | 1.27E-03 | 1.58E-02 |
| regulation of nodal signaling pathway | GO:1900107 | 13 | 8 | 1.64 | 4.89 | 4.46E-05 | 9.06E-04 |
| regulation of cell-matrix adhesion | GO:0001952 | 10 | 6 | 1.26 | 4.77 | 5.28E-04 | 7.50E-03 |
| response to auditory stimulus | GO:0010996 | 10 | 6 | 1.26 | 4.77 | 5.28E-04 | 7.49E-03 |
| regulation of axon guidance | GO:1902667 | 10 | 6 | 1.26 | 4.77 | 5.28E-04 | 7.48E-03 |
| hormone transport | GO:0009914 | 12 | 7 | 1.51 | 4.63 | 2.21E-04 | 3.55E-03 |
| pituitary gland development | GO:0021983 | 19 | 11 | 2.39 | 4.60 | 3.53E-06 | 9.58E-05 |
| rhombomere development | GO:0021546 | 21 | 12 | 2.64 | 4.54 | 1.51E-06 | 4.42E-05 |
| epithalamus development | GO:0021538 | 21 | 12 | 2.64 | 4.54 | 1.51E-06 | 4.41E-05 |

| | | | | | | | |
|---|---|---|---|---|---|---|---|
| S-adenosylmethionine metabolic process | GO:0046500 | 14 | 8 | 1.76 | 4.54 | 9.26E-05 | 1.70E-03 |
| cardioblast differentiation | GO:0010002 | 9 | 5 | 1.13 | 4.41 | 2.56E-03 | 2.86E-02 |
| angioblast cell migration from lateral mesoderm to midline | GO:0035479 | 9 | 5 | 1.13 | 4.41 | 2.56E-03 | 2.85E-02 |
| heme transport | GO:0015886 | 9 | 5 | 1.13 | 4.41 | 2.56E-03 | 2.85E-02 |
| regulation of cell-substrate junction organization | GO:0150116 | 9 | 5 | 1.13 | 4.41 | 2.56E-03 | 2.85E-02 |
| regulation of focal adhesion assembly | GO:0051893 | 9 | 5 | 1.13 | 4.41 | 2.56E-03 | 2.84E-02 |
| anterior lateral line development | GO:0048899 | 9 | 5 | 1.13 | 4.41 | 2.56E-03 | 2.84E-02 |
| regulation of cell-substrate junction assembly | GO:0090109 | 9 | 5 | 1.13 | 4.41 | 2.56E-03 | 2.84E-02 |
| habenula development | GO:0021986 | 18 | 10 | 2.27 | 4.41 | 1.64E-05 | 3.68E-04 |
| body morphogenesis | GO:0010171 | 18 | 10 | 2.27 | 4.41 | 1.64E-05 | 3.67E-04 |
| face morphogenesis | GO:0060325 | 9 | 5 | 1.13 | 4.41 | 2.56E-03 | 2.83E-02 |
| cell migration in hindbrain | GO:0021535 | 11 | 6 | 1.38 | 4.33 | 1.04E-03 | 1.35E-02 |
| neural plate development | GO:0001840 | 11 | 6 | 1.38 | 4.33 | 1.04E-03 | 1.35E-02 |
| vascular endothelial growth factor signaling pathway | GO:0038084 | 11 | 6 | 1.38 | 4.33 | 1.04E-03 | 1.34E-02 |
| benzene-containing compound metabolic process | GO:0042537 | 11 | 6 | 1.38 | 4.33 | 1.04E-03 | 1.34E-02 |
| positive regulation of phosphatidylinositol 3-kinase/protein kinase B signal transduction | GO:0051897 | 11 | 6 | 1.38 | 4.33 | 1.04E-03 | 1.34E-02 |
| inhibitory synapse assembly | GO:1904862 | 11 | 6 | 1.38 | 4.33 | 1.04E-03 | 1.34E-02 |
| negative regulation of cell fate specification | GO:0009996 | 11 | 6 | 1.38 | 4.33 | 1.04E-03 | 1.34E-02 |
| cloaca development | GO:0035844 | 11 | 6 | 1.38 | 4.33 | 1.04E-03 | 1.33E-02 |
| face development | GO:0060324 | 11 | 6 | 1.38 | 4.33 | 1.04E-03 | 1.33E-02 |
| head morphogenesis | GO:0060323 | 11 | 6 | 1.38 | 4.33 | 1.04E-03 | 1.33E-02 |
| vascular endothelial growth factor receptor signaling pathway | GO:0048010 | 15 | 8 | 1.89 | 4.24 | 1.76E-04 | 2.92E-03 |
| cardiac muscle tissue regeneration | GO:0061026 | 15 | 8 | 1.89 | 4.24 | 1.76E-04 | 2.91E-03 |
| regulation of activin receptor signaling pathway | GO:0032925 | 17 | 9 | 2.14 | 4.21 | 7.35E-05 | 1.41E-03 |
| muscle cell proliferation | GO:0033002 | 17 | 9 | 2.14 | 4.21 | 7.35E-05 | 1.40E-03 |
| adenohypophysis development | GO:0021984 | 17 | 9 | 2.14 | 4.21 | 7.35E-05 | 1.40E-03 |
| rhombomere morphogenesis | GO:0021593 | 19 | 10 | 2.39 | 4.18 | 3.08E-05 | 6.47E-04 |
| retinoic acid receptor signaling pathway | GO:0048384 | 23 | 12 | 2.89 | 4.15 | 5.46E-06 | 1.38E-04 |
| anterior/posterior axon guidance | GO:0033564 | 12 | 6 | 1.51 | 3.97 | 1.86E-03 | 2.20E-02 |
| proximal tubule development | GO:0072014 | 12 | 6 | 1.51 | 3.97 | 1.86E-03 | 2.19E-02 |
| cellular response to vascular endothelial growth factor stimulus | GO:0035924 | 14 | 7 | 1.76 | 3.97 | 7.60E-04 | 1.02E-02 |
| response to zinc ion | GO:0010043 | 16 | 8 | 2.01 | 3.97 | 3.14E-04 | 4.79E-03 |
| pons development | GO:0021548 | 18 | 9 | 2.27 | 3.97 | 1.31E-04 | 2.27E-03 |
| establishment of tissue polarity | GO:0007164 | 10 | 5 | 1.26 | 3.97 | 4.58E-03 | 4.56E-02 |
| cardiac muscle tissue growth | GO:0055017 | 16 | 8 | 2.01 | 3.97 | 3.14E-04 | 4.78E-03 |
| negative regulation of extrinsic apoptotic signaling pathway | GO:2001237 | 16 | 8 | 2.01 | 3.97 | 3.14E-04 | 4.77E-03 |
| positive regulation of BMP signaling pathway | GO:0030513 | 18 | 9 | 2.27 | 3.97 | 1.31E-04 | 2.26E-03 |
| Schwann cell development | GO:0014044 | 20 | 10 | 2.52 | 3.97 | 5.46E-05 | 1.08E-03 |
| venous blood vessel development | GO:0060841 | 18 | 9 | 2.27 | 3.97 | 1.31E-04 | 2.26E-03 |

| | | | | | | | |
|---|---|---|---|---|---|---|---|
| iron coordination entity transport | GO:1901678 | 10 | 5 | 1.26 | 3.97 | 4.58E-03 | 4.56E-02 |
| embryonic retina morphogenesis in camera-type eye | GO:0060059 | 29 | 14 | 3.65 | 3.84 | 2.96E-06 | 8.19E-05 |
| hypothalamus development | GO:0021854 | 27 | 13 | 3.40 | 3.83 | 6.96E-06 | 1.74E-04 |
| cAMP-mediated signaling | GO:0019933 | 23 | 11 | 2.89 | 3.80 | 3.89E-05 | 7.98E-04 |
| response to glucose | GO:0009749 | 17 | 8 | 2.14 | 3.74 | 5.28E-04 | 7.55E-03 |
| response to hexose | GO:0009746 | 17 | 8 | 2.14 | 3.74 | 5.28E-04 | 7.54E-03 |
| response to carbohydrate | GO:0009743 | 17 | 8 | 2.14 | 3.74 | 5.28E-04 | 7.53E-03 |
| response to monosaccharide | GO:0034284 | 17 | 8 | 2.14 | 3.74 | 5.28E-04 | 7.51E-03 |
| heart growth | GO:0060419 | 17 | 8 | 2.14 | 3.74 | 5.28E-04 | 7.50E-03 |
| semaphorin-plexin signaling pathway | GO:0071526 | 47 | 22 | 5.92 | 3.72 | 9.18E-09 | 3.55E-07 |
| cell volume homeostasis | GO:0006884 | 15 | 7 | 1.89 | 3.71 | 1.27E-03 | 1.58E-02 |
| steroid hormone mediated signaling pathway | GO:0043401 | 15 | 7 | 1.89 | 3.71 | 1.27E-03 | 1.58E-02 |
| diencephalon development | GO:0021536 | 86 | 40 | 10.82 | 3.70 | 1.13E-14 | 7.79E-13 |
| limbic system development | GO:0021761 | 28 | 13 | 3.52 | 3.69 | 1.15E-05 | 2.66E-04 |
| olfactory placode development | GO:0071698 | 13 | 6 | 1.64 | 3.67 | 3.08E-03 | 3.28E-02 |
| posterior lateral line neuromast hair cell differentiation | GO:0048923 | 13 | 6 | 1.64 | 3.67 | 3.08E-03 | 3.27E-02 |
| anterior lateral line system development | GO:0048898 | 13 | 6 | 1.64 | 3.67 | 3.08E-03 | 3.27E-02 |
| neural nucleus development | GO:0048857 | 26 | 12 | 3.27 | 3.67 | 2.70E-05 | 5.78E-04 |
| embryonic neurocranium morphogenesis | GO:0048702 | 26 | 12 | 3.27 | 3.67 | 2.70E-05 | 5.77E-04 |
| negative regulation of signal transduction by p53 class mediator | GO:1901797 | 13 | 6 | 1.64 | 3.67 | 3.08E-03 | 3.27E-02 |
| mesendoderm development | GO:0048382 | 13 | 6 | 1.64 | 3.67 | 3.08E-03 | 3.26E-02 |
| hematopoietic or lymphoid organ development | GO:0048534 | 24 | 11 | 3.02 | 3.64 | 6.37E-05 | 1.25E-03 |
| cranial nerve morphogenesis | GO:0021602 | 24 | 11 | 3.02 | 3.64 | 6.37E-05 | 1.24E-03 |
| Schwann cell differentiation | GO:0014037 | 22 | 10 | 2.77 | 3.61 | 1.50E-04 | 2.55E-03 |
| zinc ion transmembrane transport | GO:0071577 | 20 | 9 | 2.52 | 3.58 | 3.56E-04 | 5.31E-03 |
| negative regulation of intrinsic apoptotic signaling pathway | GO:2001243 | 20 | 9 | 2.52 | 3.58 | 3.56E-04 | 5.30E-03 |
| midbrain development | GO:0030901 | 38 | 17 | 4.78 | 3.55 | 9.96E-07 | 3.03E-05 |
| regulation of vascular permeability | GO:0043114 | 18 | 8 | 2.27 | 3.53 | 8.45E-04 | 1.12E-02 |
| nuclear migration | GO:0007097 | 18 | 8 | 2.27 | 3.53 | 8.45E-04 | 1.12E-02 |
| nucleus localization | GO:0051647 | 18 | 8 | 2.27 | 3.53 | 8.45E-04 | 1.12E-02 |
| inner ear receptor cell development | GO:0060119 | 41 | 18 | 5.16 | 3.49 | 6.76E-07 | 2.12E-05 |
| inner ear receptor cell differentiation | GO:0060113 | 48 | 21 | 6.04 | 3.48 | 8.64E-08 | 3.07E-06 |
| gland morphogenesis | GO:0022612 | 16 | 7 | 2.01 | 3.48 | 2.01E-03 | 2.31E-02 |
| peripheral nervous system axon ensheathment | GO:0032292 | 16 | 7 | 2.01 | 3.48 | 2.01E-03 | 2.31E-02 |
| zinc ion transport | GO:0006829 | 23 | 10 | 2.89 | 3.45 | 2.36E-04 | 3.73E-03 |
| closure of optic fissure | GO:0061386 | 23 | 10 | 2.89 | 3.45 | 2.36E-04 | 3.72E-03 |
| cartilage morphogenesis | GO:0060536 | 30 | 13 | 3.78 | 3.44 | 2.88E-05 | 6.10E-04 |
| regulation of vascular endothelial growth factor receptor signaling pathway | GO:0030947 | 14 | 6 | 1.76 | 3.40 | 4.82E-03 | 4.76E-02 |
| myelination in peripheral nervous system | GO:0022011 | 14 | 6 | 1.76 | 3.40 | 4.82E-03 | 4.76E-02 |
| rhombomere formation | GO:0021594 | 14 | 6 | 1.76 | 3.40 | 4.82E-03 | 4.75E-02 |

| | | | | | | | |
|---|---|---|---|---|---|---|---|
| cardiac muscle cell proliferation | GO:0060038 | 14 | 6 | 1.76 | 3.40 | 4.82E-03 | 4.75E-02 |
| embryonic camera-type eye development | GO:0031076 | 56 | 24 | 7.05 | 3.40 | 1.71E-08 | 6.42E-07 |
| positive regulation of macroautophagy | GO:0016239 | 14 | 6 | 1.76 | 3.40 | 4.82E-03 | 4.74E-02 |
| regulation of peptide hormone secretion | GO:0090276 | 14 | 6 | 1.76 | 3.40 | 4.82E-03 | 4.74E-02 |
| striated muscle cell proliferation | GO:0014855 | 14 | 6 | 1.76 | 3.40 | 4.82E-03 | 4.73E-02 |
| negative regulation of cell fate commitment | GO:0010454 | 14 | 6 | 1.76 | 3.40 | 4.82E-03 | 4.73E-02 |
| regulation of peptide transport | GO:0090087 | 14 | 6 | 1.76 | 3.40 | 4.82E-03 | 4.72E-02 |
| regulation of peptide secretion | GO:0002791 | 14 | 6 | 1.76 | 3.40 | 4.82E-03 | 4.72E-02 |
| neuromast hair cell development | GO:0035675 | 21 | 9 | 2.64 | 3.40 | 5.54E-04 | 7.76E-03 |
| regulation of extrinsic apoptotic signaling pathway | GO:2001236 | 21 | 9 | 2.64 | 3.40 | 5.54E-04 | 7.75E-03 |
| intracellular receptor signaling pathway | GO:0030522 | 42 | 18 | 5.29 | 3.40 | 1.04E-06 | 3.17E-05 |
| immune system development | GO:0002520 | 35 | 15 | 4.41 | 3.40 | 8.28E-06 | 1.99E-04 |
| determination of ventral identity | GO:0048264 | 21 | 9 | 2.64 | 3.40 | 5.54E-04 | 7.74E-03 |
| negative regulation of axon extension | GO:0030517 | 40 | 17 | 5.03 | 3.38 | 2.39E-06 | 6.76E-05 |
| regulation of endodermal cell differentiation | GO:1903224 | 19 | 8 | 2.39 | 3.35 | 1.30E-03 | 1.61E-02 |
| central nervous system projection neuron axonogenesis | GO:0021952 | 19 | 8 | 2.39 | 3.35 | 1.30E-03 | 1.61E-02 |
| ectodermal placode development | GO:0071696 | 43 | 18 | 5.41 | 3.33 | 1.59E-06 | 4.59E-05 |
| positive regulation of transmembrane receptor protein serine/threonine kinase signaling pathway | GO:0090100 | 24 | 10 | 3.02 | 3.31 | 3.59E-04 | 5.34E-03 |
| notochord morphogenesis | GO:0048570 | 29 | 12 | 3.65 | 3.29 | 1.01E-04 | 1.83E-03 |
| thyroid gland development | GO:0030878 | 17 | 7 | 2.14 | 3.27 | 3.06E-03 | 3.27E-02 |
| regulation of endodermal cell fate specification | GO:0042663 | 17 | 7 | 2.14 | 3.27 | 3.06E-03 | 3.27E-02 |
| determination of pancreatic left/right asymmetry | GO:0035469 | 17 | 7 | 2.14 | 3.27 | 3.06E-03 | 3.27E-02 |
| medial fin morphogenesis | GO:0035141 | 17 | 7 | 2.14 | 3.27 | 3.06E-03 | 3.26E-02 |
| feeding behavior | GO:0007631 | 17 | 7 | 2.14 | 3.27 | 3.06E-03 | 3.26E-02 |
| positive regulation of axonogenesis | GO:0050772 | 34 | 14 | 4.28 | 3.27 | 2.87E-05 | 6.09E-04 |
| dorsal/ventral axis specification | GO:0009950 | 22 | 9 | 2.77 | 3.25 | 8.33E-04 | 1.11E-02 |
| blood vessel endothelial cell migration | GO:0043534 | 22 | 9 | 2.77 | 3.25 | 8.33E-04 | 1.11E-02 |
| ephrin receptor signaling pathway | GO:0048013 | 27 | 11 | 3.40 | 3.24 | 2.32E-04 | 3.67E-03 |
| cardiac chamber morphogenesis | GO:0003206 | 27 | 11 | 3.40 | 3.24 | 2.32E-04 | 3.67E-03 |
| neuron recognition | GO:0008038 | 32 | 13 | 4.03 | 3.23 | 6.54E-05 | 1.27E-03 |
| midbrain-hindbrain boundary development | GO:0030917 | 42 | 17 | 5.29 | 3.22 | 5.36E-06 | 1.37E-04 |
| negative regulation of axonogenesis | GO:0050771 | 42 | 17 | 5.29 | 3.22 | 5.36E-06 | 1.36E-04 |
| embryonic camera-type eye morphogenesis | GO:0048596 | 47 | 19 | 5.92 | 3.21 | 1.55E-06 | 4.49E-05 |
| embryonic eye morphogenesis | GO:0048048 | 57 | 23 | 7.17 | 3.21 | 1.31E-07 | 4.45E-06 |
| regulation of actomyosin structure organization | GO:0110020 | 25 | 10 | 3.15 | 3.18 | 5.32E-04 | 7.49E-03 |
| auditory receptor cell development | GO:0060117 | 20 | 8 | 2.52 | 3.18 | 1.93E-03 | 2.27E-02 |
| response to alcohol | GO:0097305 | 20 | 8 | 2.52 | 3.18 | 1.93E-03 | 2.27E-02 |
| digestive system process | GO:0022600 | 25 | 10 | 3.15 | 3.18 | 5.32E-04 | 7.48E-03 |
| thymus development | GO:0048538 | 20 | 8 | 2.52 | 3.18 | 1.93E-03 | 2.26E-02 |

| | | | | | | | |
|---|---|---|---|---|---|---|---|
| ectodermal placode morphogenesis | GO:0071697 | 35 | 14 | 4.41 | 3.18 | 4.23E-05 | 8.65E-04 |
| mechanoreceptor differentiation | GO:0042490 | 70 | 28 | 8.81 | 3.18 | 7.31E-09 | 2.86E-07 |
| negative regulation of axon extension involved in axon guidance | GO:0048843 | 33 | 13 | 4.15 | 3.13 | 9.56E-05 | 1.74E-03 |
| optic cup morphogenesis involved in camera-type eye development | GO:0002072 | 28 | 11 | 3.52 | 3.12 | 3.39E-04 | 5.09E-03 |
| otic placode formation | GO:0043049 | 23 | 9 | 2.89 | 3.11 | 1.22E-03 | 1.53E-02 |
| negative regulation of developmental growth | GO:0048640 | 46 | 18 | 5.79 | 3.11 | 5.05E-06 | 1.30E-04 |
| cardiac chamber development | GO:0003205 | 46 | 18 | 5.79 | 3.11 | 5.05E-06 | 1.29E-04 |
| establishment of spindle orientation | GO:0051294 | 23 | 9 | 2.89 | 3.11 | 1.22E-03 | 1.52E-02 |
| optic nerve development | GO:0021554 | 18 | 7 | 2.27 | 3.09 | 4.46E-03 | 4.47E-02 |
| activin receptor signaling pathway | GO:0032924 | 18 | 7 | 2.27 | 3.09 | 4.46E-03 | 4.46E-02 |
| myeloid leukocyte activation | GO:0002274 | 18 | 7 | 2.27 | 3.09 | 4.46E-03 | 4.46E-02 |
| organ growth | GO:0035265 | 31 | 12 | 3.90 | 3.08 | 2.15E-04 | 3.48E-03 |
| cyclic-nucleotide-mediated signaling | GO:0019935 | 31 | 12 | 3.90 | 3.08 | 2.15E-04 | 3.47E-03 |
| ear morphogenesis | GO:0042471 | 96 | 37 | 12.08 | 3.06 | 1.07E-10 | 5.19E-09 |
| embryonic skeletal system morphogenesis | GO:0048704 | 187 | 72 | 23.54 | 3.06 | 2.06E-19 | 1.80E-17 |
| thigmotaxis | GO:0001966 | 26 | 10 | 3.27 | 3.06 | 7.67E-04 | 1.03E-02 |
| nuclear-transcribed mRNA catabolic process, nonsense-mediated decay | GO:0000184 | 26 | 10 | 3.27 | 3.06 | 7.67E-04 | 1.03E-02 |
| morphogenesis of a branching structure | GO:0001763 | 26 | 10 | 3.27 | 3.06 | 7.67E-04 | 1.02E-02 |
| establishment of mitotic spindle localization | GO:0040001 | 26 | 10 | 3.27 | 3.06 | 7.67E-04 | 1.02E-02 |
| establishment of mitotic spindle orientation | GO:0000132 | 21 | 8 | 2.64 | 3.03 | 2.78E-03 | 3.05E-02 |
| retinal cone cell differentiation | GO:0042670 | 21 | 8 | 2.64 | 3.03 | 2.78E-03 | 3.04E-02 |
| peptide transport | GO:0015833 | 21 | 8 | 2.64 | 3.03 | 2.78E-03 | 3.04E-02 |
| semicircular canal morphogenesis | GO:0048752 | 21 | 8 | 2.64 | 3.03 | 2.78E-03 | 3.04E-02 |
| embryonic viscerocranium morphogenesis | GO:0048703 | 108 | 41 | 13.59 | 3.02 | 1.88E-11 | 1.01E-09 |
| regulation of phosphatidylinositol 3-kinase/protein kinase B signal transduction | GO:0051896 | 29 | 11 | 3.65 | 3.01 | 4.84E-04 | 7.01E-03 |
| positive regulation of neuron projection development | GO:0010976 | 29 | 11 | 3.65 | 3.01 | 4.84E-04 | 7.00E-03 |
| chloride transmembrane transport | GO:1902476 | 58 | 22 | 7.30 | 3.01 | 8.67E-07 | 2.64E-05 |
| neuromast hair cell differentiation | GO:0048886 | 29 | 11 | 3.65 | 3.01 | 4.84E-04 | 6.99E-03 |
| semicircular canal development | GO:0060872 | 29 | 11 | 3.65 | 3.01 | 4.84E-04 | 6.98E-03 |
| inner ear morphogenesis | GO:0042472 | 95 | 36 | 11.96 | 3.01 | 3.32E-10 | 1.50E-08 |
| posterior lateral line neuromast development | GO:0048919 | 37 | 14 | 4.66 | 3.01 | 8.71E-05 | 1.63E-03 |
| determination of dorsal/ventral asymmetry | GO:0048262 | 37 | 14 | 4.66 | 3.01 | 8.71E-05 | 1.63E-03 |
| skeletal system morphogenesis | GO:0048705 | 217 | 82 | 27.31 | 3.00 | 2.67E-21 | 2.86E-19 |
| morphogenesis of embryonic epithelium | GO:0016331 | 69 | 26 | 8.68 | 2.99 | 1.05E-07 | 3.68E-06 |
| neural tube patterning | GO:0021532 | 56 | 21 | 7.05 | 2.98 | 1.90E-06 | 5.43E-05 |
| central nervous system neuron axonogenesis | GO:0021955 | 24 | 9 | 3.02 | 2.98 | 1.73E-03 | 2.09E-02 |
| rostrocaudal neural tube patterning | GO:0021903 | 48 | 18 | 6.04 | 2.98 | 1.02E-05 | 2.39E-04 |
| hair cell differentiation | GO:0035315 | 64 | 24 | 8.06 | 2.98 | 3.57E-07 | 1.15E-05 |
| regulation of fibroblast growth factor receptor signaling pathway | GO:0040036 | 32 | 12 | 4.03 | 2.98 | 3.05E-04 | 4.69E-03 |

| | | | | | | | |
|---|---|---|---|---|---|---|---|
| endothelial cell migration | GO:0043542 | 32 | 12 | 4.03 | 2.98 | 3.05E-04 | 4.68E-03 |
| embryonic cranial skeleton morphogenesis | GO:0048701 | 171 | 64 | 21.52 | 2.97 | 1.06E-16 | 8.18E-15 |
| negative regulation of cell projection organization | GO:0031345 | 54 | 20 | 6.80 | 2.94 | 4.16E-06 | 1.09E-04 |
| inner ear auditory receptor cell differentiation | GO:0042491 | 27 | 10 | 3.40 | 2.94 | 1.08E-03 | 1.38E-02 |
| negative regulation of chemotaxis | GO:0050922 | 38 | 14 | 4.78 | 2.93 | 1.22E-04 | 2.13E-03 |
| embryonic skeletal system development | GO:0048706 | 204 | 75 | 25.68 | 2.92 | 8.45E-19 | 7.31E-17 |
| ear development | GO:0043583 | 188 | 69 | 23.66 | 2.92 | 2.29E-17 | 1.86E-15 |
| neural precursor cell proliferation | GO:0061351 | 30 | 11 | 3.78 | 2.91 | 6.78E-04 | 9.27E-03 |
| ectodermal placode formation | GO:0060788 | 30 | 11 | 3.78 | 2.91 | 6.78E-04 | 9.26E-03 |
| cardiac muscle tissue development | GO:0048738 | 101 | 37 | 12.71 | 2.91 | 5.83E-10 | 2.57E-08 |
| posterior lateral line development | GO:0048916 | 71 | 26 | 8.94 | 2.91 | 2.05E-07 | 6.82E-06 |
| positive regulation of neurogenesis | GO:0050769 | 71 | 26 | 8.94 | 2.91 | 2.05E-07 | 6.79E-06 |
| negative chemotaxis | GO:0050919 | 41 | 15 | 5.16 | 2.91 | 7.72E-05 | 1.46E-03 |
| retinal ganglion cell axon guidance | GO:0031290 | 44 | 16 | 5.54 | 2.89 | 4.89E-05 | 9.84E-04 |
| medial fin development | GO:0033338 | 22 | 8 | 2.77 | 2.89 | 3.89E-03 | 3.97E-02 |
| negative regulation of apoptotic signaling pathway | GO:2001234 | 47 | 17 | 5.92 | 2.87 | 3.10E-05 | 6.48E-04 |
| neural tube development | GO:0021915 | 108 | 39 | 13.59 | 2.87 | 3.31E-10 | 1.50E-08 |
| cranial nerve development | GO:0021545 | 61 | 22 | 7.68 | 2.87 | 2.34E-06 | 6.62E-05 |
| inner ear development | GO:0048839 | 186 | 67 | 23.41 | 2.86 | 2.00E-16 | 1.49E-14 |
| nephron tubule development | GO:0072080 | 25 | 9 | 3.15 | 2.86 | 2.41E-03 | 2.71E-02 |
| vesicle cytoskeletal trafficking | GO:0099518 | 25 | 9 | 3.15 | 2.86 | 2.41E-03 | 2.70E-02 |
| motor neuron axon guidance | GO:0008045 | 50 | 18 | 6.29 | 2.86 | 1.96E-05 | 4.32E-04 |
| striated muscle tissue development | GO:0014706 | 103 | 37 | 12.96 | 2.85 | 1.10E-09 | 4.73E-08 |
| tissue migration | GO:0090130 | 53 | 19 | 6.67 | 2.85 | 1.25E-05 | 2.85E-04 |
| negative regulation of growth | GO:0045926 | 56 | 20 | 7.05 | 2.84 | 7.91E-06 | 1.92E-04 |
| posterior lateral line system development | GO:0048915 | 84 | 30 | 10.57 | 2.84 | 4.69E-08 | 1.70E-06 |
| regulation of axon extension involved in axon guidance | GO:0048841 | 42 | 15 | 5.29 | 2.84 | 1.06E-04 | 1.91E-03 |
| cartilage development | GO:0051216 | 157 | 56 | 19.76 | 2.83 | 9.26E-14 | 6.17E-12 |
| mechanosensory lateral line system development | GO:0048881 | 87 | 31 | 10.95 | 2.83 | 3.00E-08 | 1.10E-06 |
| neuron projection guidance | GO:0097485 | 262 | 93 | 32.98 | 2.82 | 1.00E-21 | 1.13E-19 |
| axon guidance | GO:0007411 | 262 | 93 | 32.98 | 2.82 | 1.00E-21 | 1.11E-19 |
| brain morphogenesis | GO:0048854 | 34 | 12 | 4.28 | 2.80 | 5.83E-04 | 8.11E-03 |
| negative regulation of neuron projection development | GO:0010977 | 51 | 18 | 6.42 | 2.80 | 2.68E-05 | 5.77E-04 |
| negative regulation of cell growth | GO:0030308 | 51 | 18 | 6.42 | 2.80 | 2.68E-05 | 5.76E-04 |
| protein secretion | GO:0009306 | 37 | 13 | 4.66 | 2.79 | 3.65E-04 | 5.41E-03 |
| establishment of protein localization to extracellular region | GO:0035592 | 37 | 13 | 4.66 | 2.79 | 3.65E-04 | 5.40E-03 |
| animal organ formation | GO:0048645 | 37 | 13 | 4.66 | 2.79 | 3.65E-04 | 5.40E-03 |
| regulation of chemotaxis | GO:0050920 | 74 | 26 | 9.31 | 2.79 | 5.24E-07 | 1.66E-05 |
| morphogenesis of an epithelial sheet | GO:0002011 | 97 | 34 | 12.21 | 2.78 | 1.06E-08 | 4.04E-07 |
| carbohydrate homeostasis | GO:0033500 | 40 | 14 | 5.03 | 2.78 | 2.29E-04 | 3.66E-03 |
| glucose homeostasis | GO:0042593 | 40 | 14 | 5.03 | 2.78 | 2.29E-04 | 3.66E-03 |

| | | | | | | | |
|---|---|---|---|---|---|---|---|
| neuromast development | GO:0048884 | 86 | 30 | 10.82 | 2.77 | 8.63E-08 | 3.08E-06 |
| epithelial tube formation | GO:0072175 | 66 | 23 | 8.31 | 2.77 | 2.75E-06 | 7.68E-05 |
| regulation of axonogenesis | GO:0050770 | 95 | 33 | 11.96 | 2.76 | 2.24E-08 | 8.26E-07 |
| connective tissue development | GO:0061448 | 170 | 59 | 21.40 | 2.76 | 8.13E-14 | 5.50E-12 |
| enteric nervous system development | GO:0048484 | 49 | 17 | 6.17 | 2.76 | 5.73E-05 | 1.13E-03 |
| lateral line development | GO:0048882 | 124 | 43 | 15.61 | 2.76 | 1.88E-10 | 8.81E-09 |
| cellular response to steroid hormone stimulus | GO:0071383 | 26 | 9 | 3.27 | 2.75 | 3.27E-03 | 3.44E-02 |
| renal tubule development | GO:0061326 | 26 | 9 | 3.27 | 2.75 | 3.27E-03 | 3.44E-02 |
| swim bladder development | GO:0048794 | 26 | 9 | 3.27 | 2.75 | 3.27E-03 | 3.43E-02 |
| forebrain development | GO:0030900 | 148 | 51 | 18.63 | 2.74 | 5.23E-12 | 3.00E-10 |
| embryonic organ morphogenesis | GO:0048562 | 442 | 152 | 55.63 | 2.73 | 6.53E-33 | 1.51E-30 |
| embryonic epithelial tube formation | GO:0001838 | 35 | 12 | 4.41 | 2.72 | 7.86E-04 | 1.05E-02 |
| protein localization to extracellular region | GO:0071692 | 38 | 13 | 4.78 | 2.72 | 4.91E-04 | 7.07E-03 |
| bone morphogenesis | GO:0060349 | 38 | 13 | 4.78 | 2.72 | 4.91E-04 | 7.06E-03 |
| cellular response to BMP stimulus | GO:0071773 | 44 | 15 | 5.54 | 2.71 | 1.93E-04 | 3.16E-03 |
| response to BMP | GO:0071772 | 44 | 15 | 5.54 | 2.71 | 1.93E-04 | 3.15E-03 |
| BMP signaling pathway | GO:0030509 | 44 | 15 | 5.54 | 2.71 | 1.93E-04 | 3.15E-03 |
| epithelium migration | GO:0090132 | 47 | 16 | 5.92 | 2.70 | 1.22E-04 | 2.13E-03 |
| autonomic nervous system development | GO:0048483 | 62 | 21 | 7.80 | 2.69 | 1.22E-05 | 2.82E-04 |
| hindbrain development | GO:0030902 | 136 | 46 | 17.12 | 2.69 | 1.16E-10 | 5.59E-09 |
| embryonic heart tube development | GO:0035050 | 172 | 58 | 21.65 | 2.68 | 5.36E-13 | 3.37E-11 |
| chloride transport | GO:0006821 | 92 | 31 | 11.58 | 2.68 | 1.31E-07 | 4.45E-06 |
| mesenchymal cell migration | GO:0090497 | 110 | 37 | 13.85 | 2.67 | 8.76E-09 | 3.41E-07 |
| skeletal system development | GO:0001501 | 372 | 125 | 46.82 | 2.67 | 3.24E-26 | 4.95E-24 |
| heart looping | GO:0001947 | 126 | 42 | 15.86 | 2.65 | 1.22E-09 | 5.21E-08 |
| neural tube formation | GO:0001841 | 33 | 11 | 4.15 | 2.65 | 1.68E-03 | 2.04E-02 |
| regulation of neuron projection development | GO:0010975 | 165 | 55 | 20.77 | 2.65 | 3.58E-12 | 2.08E-10 |
| spindle localization | GO:0051653 | 33 | 11 | 4.15 | 2.65 | 1.68E-03 | 2.03E-02 |
| regulation of cell junction assembly | GO:1901888 | 33 | 11 | 4.15 | 2.65 | 1.68E-03 | 2.03E-02 |
| establishment of spindle localization | GO:0051293 | 33 | 11 | 4.15 | 2.65 | 1.68E-03 | 2.03E-02 |
| sex differentiation | GO:0007548 | 33 | 11 | 4.15 | 2.65 | 1.68E-03 | 2.03E-02 |
| otic vesicle formation | GO:0030916 | 27 | 9 | 3.40 | 2.65 | 4.37E-03 | 4.42E-02 |
| hindbrain morphogenesis | GO:0021575 | 39 | 13 | 4.91 | 2.65 | 6.53E-04 | 8.99E-03 |
| endothelial cell proliferation | GO:0001935 | 27 | 9 | 3.40 | 2.65 | 4.37E-03 | 4.41E-02 |
| very long-chain fatty acid metabolic process | GO:0000038 | 27 | 9 | 3.40 | 2.65 | 4.37E-03 | 4.41E-02 |
| neural crest cell migration | GO:0001755 | 108 | 36 | 13.59 | 2.65 | 1.81E-08 | 6.77E-07 |
| GPI anchor biosynthetic process | GO:0006506 | 27 | 9 | 3.40 | 2.65 | 4.37E-03 | 4.40E-02 |
| positive regulation of proteolysis involved in protein catabolic process | GO:1903052 | 27 | 9 | 3.40 | 2.65 | 4.37E-03 | 4.40E-02 |
| regulation of Notch signaling pathway | GO:0008593 | 60 | 20 | 7.55 | 2.65 | 2.56E-05 | 5.54E-04 |
| protein localization to lysosome | GO:0061462 | 27 | 9 | 3.40 | 2.65 | 4.37E-03 | 4.39E-02 |
| otolith development | GO:0048840 | 60 | 20 | 7.55 | 2.65 | 2.56E-05 | 5.52E-04 |

| | | | | | | | |
|---|---|---|---|---|---|---|---|
| heart formation | GO:0060914 | 30 | 10 | 3.78 | 2.65 | 2.70E-03 | 2.98E-02 |
| otolith morphogenesis | GO:0032474 | 27 | 9 | 3.40 | 2.65 | 4.37E-03 | 4.39E-02 |
| receptor localization to synapse | GO:0097120 | 36 | 12 | 4.53 | 2.65 | 1.05E-03 | 1.34E-02 |
| receptor internalization | GO:0031623 | 30 | 10 | 3.78 | 2.65 | 2.70E-03 | 2.98E-02 |
| digestion | GO:0007586 | 30 | 10 | 3.78 | 2.65 | 2.70E-03 | 2.97E-02 |
| lateral mesoderm development | GO:0048368 | 27 | 9 | 3.40 | 2.65 | 4.37E-03 | 4.38E-02 |
| synapse assembly | GO:0007416 | 42 | 14 | 5.29 | 2.65 | 4.09E-04 | 6.03E-03 |
| positive regulation of cell development | GO:0010720 | 94 | 31 | 11.83 | 2.62 | 2.27E-07 | 7.48E-06 |
| tube formation | GO:0035148 | 73 | 24 | 9.19 | 2.61 | 5.41E-06 | 1.37E-04 |
| lateral line system development | GO:0048925 | 140 | 46 | 17.62 | 2.61 | 3.48E-10 | 1.57E-08 |
| negative regulation of nervous system development | GO:0051961 | 61 | 20 | 7.68 | 2.60 | 3.36E-05 | 7.02E-04 |
| endocrine system development | GO:0035270 | 116 | 38 | 14.60 | 2.60 | 1.27E-08 | 4.79E-07 |
| epiboly involved in gastrulation with mouth forming second | GO:0055113 | 46 | 15 | 5.79 | 2.59 | 3.37E-04 | 5.08E-03 |
| sensory perception of sound | GO:0007605 | 46 | 15 | 5.79 | 2.59 | 3.37E-04 | 5.07E-03 |
| positive regulation of nervous system development | GO:0051962 | 86 | 28 | 10.82 | 2.59 | 1.15E-06 | 3.44E-05 |
| epithelial cell migration | GO:0010631 | 43 | 14 | 5.41 | 2.59 | 5.37E-04 | 7.54E-03 |
| epithelial tube morphogenesis | GO:0060562 | 249 | 81 | 31.34 | 2.58 | 1.55E-16 | 1.18E-14 |
| neural crest cell development | GO:0014032 | 148 | 48 | 18.63 | 2.58 | 2.41E-10 | 1.12E-08 |
| presynaptic endocytosis | GO:0140238 | 37 | 12 | 4.66 | 2.58 | 1.37E-03 | 1.69E-02 |
| stem cell development | GO:0048864 | 148 | 48 | 18.63 | 2.58 | 2.41E-10 | 1.11E-08 |
| synaptic vesicle endocytosis | GO:0048488 | 37 | 12 | 4.66 | 2.58 | 1.37E-03 | 1.68E-02 |
| inorganic anion transmembrane transport | GO:0098661 | 74 | 24 | 9.31 | 2.58 | 7.06E-06 | 1.76E-04 |
| embryonic heart tube morphogenesis | GO:0003143 | 142 | 46 | 17.87 | 2.57 | 5.91E-10 | 2.59E-08 |
| regulation of cell size | GO:0008361 | 105 | 34 | 13.22 | 2.57 | 9.97E-08 | 3.52E-06 |
| transmission of nerve impulse | GO:0019226 | 34 | 11 | 4.28 | 2.57 | 2.20E-03 | 2.51E-02 |
| regulation of axon extension | GO:0030516 | 68 | 22 | 8.56 | 2.57 | 1.75E-05 | 3.89E-04 |
| retina morphogenesis in camera-type eye | GO:0060042 | 130 | 42 | 16.36 | 2.57 | 3.55E-09 | 1.45E-07 |
| regulation of postsynaptic membrane neurotransmitter receptor levels | GO:0099072 | 31 | 10 | 3.90 | 2.56 | 3.54E-03 | 3.69E-02 |
| positive regulation of vasculature development | GO:1904018 | 31 | 10 | 3.90 | 2.56 | 3.54E-03 | 3.69E-02 |
| gastrulation with mouth forming second | GO:0001702 | 56 | 18 | 7.05 | 2.55 | 1.10E-04 | 1.95E-03 |
| endoderm development | GO:0007492 | 56 | 18 | 7.05 | 2.55 | 1.10E-04 | 1.95E-03 |
| neural crest cell differentiation | GO:0014033 | 165 | 53 | 20.77 | 2.55 | 4.31E-11 | 2.21E-09 |
| ossification | GO:0001503 | 81 | 26 | 10.20 | 2.55 | 3.69E-06 | 9.98E-05 |
| monoatomic anion transmembrane transport | GO:0098656 | 72 | 23 | 9.06 | 2.54 | 1.44E-05 | 3.26E-04 |
| cranial skeletal system development | GO:1904888 | 245 | 78 | 30.84 | 2.53 | 2.15E-15 | 1.56E-13 |
| retina homeostasis | GO:0001895 | 44 | 14 | 5.54 | 2.53 | 6.97E-04 | 9.49E-03 |
| synaptic vesicle recycling | GO:0036465 | 44 | 14 | 5.54 | 2.53 | 6.97E-04 | 9.48E-03 |
| cell population proliferation | GO:0008283 | 151 | 48 | 19.01 | 2.53 | 5.20E-10 | 2.32E-08 |
| gland development | GO:0048732 | 252 | 80 | 31.72 | 2.52 | 1.13E-15 | 8.33E-14 |
| camera-type eye photoreceptor cell differentiation | GO:0060219 | 41 | 13 | 5.16 | 2.52 | 1.11E-03 | 1.42E-02 |

| | | | | | | | |
|---|---|---|---|---|---|---|---|
| cellular ketone metabolic process | GO:0042180 | 41 | 13 | 5.16 | 2.52 | 1.11E-03 | 1.42E-02 |
| limb development | GO:0060173 | 79 | 25 | 9.94 | 2.51 | 7.49E-06 | 1.85E-04 |
| regulation of cell morphogenesis | GO:0022604 | 79 | 25 | 9.94 | 2.51 | 7.49E-06 | 1.84E-04 |
| purine nucleoside bisphosphate biosynthetic process | GO:0034033 | 38 | 12 | 4.78 | 2.51 | 1.78E-03 | 2.13E-02 |
| ribonucleoside bisphosphate biosynthetic process | GO:0034030 | 38 | 12 | 4.78 | 2.51 | 1.78E-03 | 2.13E-02 |
| otic vesicle development | GO:0071599 | 57 | 18 | 7.17 | 2.51 | 1.41E-04 | 2.42E-03 |
| embryonic camera-type eye formation | GO:0060900 | 38 | 12 | 4.78 | 2.51 | 1.78E-03 | 2.13E-02 |
| actin filament-based movement | GO:0030048 | 38 | 12 | 4.78 | 2.51 | 1.78E-03 | 2.13E-02 |
| nucleoside bisphosphate biosynthetic process | GO:0033866 | 38 | 12 | 4.78 | 2.51 | 1.78E-03 | 2.12E-02 |
| hormone-mediated signaling pathway | GO:0009755 | 54 | 17 | 6.80 | 2.50 | 2.24E-04 | 3.59E-03 |
| pronephros development | GO:0048793 | 143 | 45 | 18.00 | 2.50 | 2.55E-09 | 1.06E-07 |
| oocyte differentiation | GO:0009994 | 35 | 11 | 4.41 | 2.50 | 2.85E-03 | 3.11E-02 |
| positive regulation of cell projection organization | GO:0031346 | 86 | 27 | 10.82 | 2.49 | 3.89E-06 | 1.04E-04 |
| response to mechanical stimulus | GO:0009612 | 51 | 16 | 6.42 | 2.49 | 3.55E-04 | 5.30E-03 |
| bone mineralization | GO:0030282 | 32 | 10 | 4.03 | 2.48 | 4.58E-03 | 4.57E-02 |
| metencephalon development | GO:0022037 | 61 | 19 | 7.68 | 2.47 | 1.15E-04 | 2.02E-03 |
| epidermis development | GO:0008544 | 106 | 33 | 13.34 | 2.47 | 4.28E-07 | 1.37E-05 |
| tissue regeneration | GO:0042246 | 135 | 42 | 16.99 | 2.47 | 1.24E-08 | 4.73E-07 |
| response to fibroblast growth factor | GO:0071774 | 45 | 14 | 5.66 | 2.47 | 8.96E-04 | 1.18E-02 |
| regulation of BMP signaling pathway | GO:0030510 | 90 | 28 | 11.33 | 2.47 | 3.17E-06 | 8.64E-05 |
| cellular response to fibroblast growth factor stimulus | GO:0044344 | 45 | 14 | 5.66 | 2.47 | 8.96E-04 | 1.17E-02 |
| central nervous system neuron development | GO:0021954 | 45 | 14 | 5.66 | 2.47 | 8.96E-04 | 1.17E-02 |
| ameboidal-type cell migration | GO:0001667 | 235 | 73 | 29.58 | 2.47 | 8.47E-14 | 5.69E-12 |
| fin regeneration | GO:0031101 | 87 | 27 | 10.95 | 2.47 | 4.97E-06 | 1.28E-04 |
| negative regulation of neurogenesis | GO:0050768 | 58 | 18 | 7.30 | 2.47 | 1.81E-04 | 2.99E-03 |
| epiboly | GO:0090504 | 71 | 22 | 8.94 | 2.46 | 3.73E-05 | 7.73E-04 |
| cardiocyte differentiation | GO:0035051 | 71 | 22 | 8.94 | 2.46 | 3.73E-05 | 7.71E-04 |
| mesenchymal cell differentiation | GO:0048762 | 184 | 57 | 23.16 | 2.46 | 5.73E-11 | 2.87E-09 |
| epidermal cell differentiation | GO:0009913 | 84 | 26 | 10.57 | 2.46 | 7.80E-06 | 1.90E-04 |
| liver development | GO:0001889 | 152 | 47 | 19.13 | 2.46 | 3.41E-09 | 1.40E-07 |
| sensory perception of mechanical stimulus | GO:0050954 | 55 | 17 | 6.92 | 2.46 | 2.86E-04 | 4.44E-03 |
| sensory organ morphogenesis | GO:0090596 | 324 | 100 | 40.78 | 2.45 | 2.89E-18 | 2.43E-16 |
| regulation of cellular response to growth factor stimulus | GO:0090287 | 162 | 50 | 20.39 | 2.45 | 1.05E-09 | 4.50E-08 |
| mitotic cytokinesis | GO:0000281 | 39 | 12 | 4.91 | 2.44 | 2.28E-03 | 2.60E-02 |
| smoothened signaling pathway | GO:0007224 | 39 | 12 | 4.91 | 2.44 | 2.28E-03 | 2.59E-02 |
| response to growth factor | GO:0070848 | 156 | 48 | 19.64 | 2.44 | 2.49E-09 | 1.04E-07 |
| oligodendrocyte differentiation | GO:0048709 | 52 | 16 | 6.55 | 2.44 | 4.53E-04 | 6.62E-03 |
| hepaticobiliary system development | GO:0061008 | 156 | 48 | 19.64 | 2.44 | 2.49E-09 | 1.04E-07 |
| nerve development | GO:0021675 | 78 | 24 | 9.82 | 2.44 | 1.92E-05 | 4.24E-04 |
| regulation of cell shape | GO:0008360 | 65 | 20 | 8.18 | 2.44 | 9.26E-05 | 1.70E-03 |

| | | | | | | | |
|---|---|---|---|---|---|---|---|
| embryo development ending in birth or egg hatching | GO:0009792 | 530 | 163 | 66.71 | 2.44 | 8.77E-29 | 1.64E-26 |
| embryonic organ development | GO:0048568 | 628 | 193 | 79.05 | 2.44 | 9.88E-34 | 2.61E-31 |
| heart morphogenesis | GO:0003007 | 241 | 74 | 30.33 | 2.44 | 9.72E-14 | 6.43E-12 |
| sprouting angiogenesis | GO:0002040 | 114 | 35 | 14.35 | 2.44 | 2.83E-07 | 9.22E-06 |
| calcium ion transmembrane import into cytosol | GO:0097553 | 49 | 15 | 6.17 | 2.43 | 7.19E-04 | 9.74E-03 |
| axon extension | GO:0048675 | 49 | 15 | 6.17 | 2.43 | 7.19E-04 | 9.73E-03 |
| monoatomic anion transport | GO:0006820 | 121 | 37 | 15.23 | 2.43 | 2.29E-07 | 7.51E-06 |
| regulation of synaptic transmission, glutamatergic | GO:0051966 | 36 | 11 | 4.53 | 2.43 | 3.64E-03 | 3.77E-02 |
| regulation of ubiquitin-dependent protein catabolic process | GO:2000058 | 36 | 11 | 4.53 | 2.43 | 3.64E-03 | 3.76E-02 |
| regulation of epithelial cell proliferation | GO:0050678 | 36 | 11 | 4.53 | 2.43 | 3.64E-03 | 3.76E-02 |
| epithelial cell proliferation | GO:0050673 | 36 | 11 | 4.53 | 2.43 | 3.64E-03 | 3.75E-02 |
| axonogenesis | GO:0007409 | 390 | 119 | 49.09 | 2.42 | 5.50E-21 | 5.69E-19 |
| notochord development | GO:0030903 | 82 | 25 | 10.32 | 2.42 | 1.55E-05 | 3.50E-04 |
| response to steroid hormone | GO:0048545 | 46 | 14 | 5.79 | 2.42 | 1.14E-03 | 1.44E-02 |
| regulation of transmembrane receptor protein serine/threonine kinase signaling pathway | GO:0090092 | 125 | 38 | 15.73 | 2.42 | 1.67E-07 | 5.61E-06 |
| chordate embryonic development | GO:0043009 | 527 | 160 | 66.33 | 2.41 | 1.56E-27 | 2.70E-25 |
| cellular response to growth factor stimulus | GO:0071363 | 155 | 47 | 19.51 | 2.41 | 5.22E-09 | 2.11E-07 |
| peripheral nervous system development | GO:0007422 | 66 | 20 | 8.31 | 2.41 | 1.17E-04 | 2.06E-03 |
| fibroblast growth factor receptor signaling pathway | GO:0008543 | 43 | 13 | 5.41 | 2.40 | 1.81E-03 | 2.16E-02 |
| regulation of neurogenesis | GO:0050767 | 169 | 51 | 21.27 | 2.40 | 1.31E-09 | 5.54E-08 |
| regeneration | GO:0031099 | 189 | 57 | 23.79 | 2.40 | 1.41E-10 | 6.73E-09 |
| transmembrane receptor protein serine/threonine kinase signaling pathway | GO:0007178 | 93 | 28 | 11.71 | 2.39 | 1.00E-05 | 2.36E-04 |
| embryonic hemopoiesis | GO:0035162 | 93 | 28 | 11.71 | 2.39 | 1.00E-05 | 2.36E-04 |
| embryonic morphogenesis | GO:0048598 | 722 | 217 | 90.88 | 2.39 | 2.96E-36 | 8.88E-34 |
| axon development | GO:0061564 | 433 | 130 | 54.50 | 2.39 | 4.24E-22 | 4.89E-20 |
| kidney development | GO:0001822 | 210 | 63 | 26.43 | 2.38 | 1.90E-11 | 1.01E-09 |
| pectoral fin development | GO:0033339 | 70 | 21 | 8.81 | 2.38 | 9.40E-05 | 1.72E-03 |
| cell morphogenesis involved in neuron differentiation | GO:0048667 | 415 | 124 | 52.24 | 2.37 | 5.88E-21 | 6.01E-19 |
| mesenchyme development | GO:0060485 | 241 | 72 | 30.33 | 2.37 | 1.12E-12 | 6.83E-11 |
| renal system development | GO:0072001 | 211 | 63 | 26.56 | 2.37 | 3.64E-11 | 1.87E-09 |
| morphogenesis of an epithelium | GO:0002009 | 479 | 143 | 60.29 | 2.37 | 8.68E-24 | 1.07E-21 |
| regulation of proteolysis involved in protein catabolic process | GO:1903050 | 47 | 14 | 5.92 | 2.37 | 1.44E-03 | 1.76E-02 |
| pronephric nephron development | GO:0039019 | 47 | 14 | 5.92 | 2.37 | 1.44E-03 | 1.76E-02 |
| endosome to lysosome transport | GO:0008333 | 37 | 11 | 4.66 | 2.36 | 4.61E-03 | 4.58E-02 |
| digestive tract morphogenesis | GO:0048546 | 37 | 11 | 4.66 | 2.36 | 4.61E-03 | 4.57E-02 |
| regulation of developmental growth | GO:0048638 | 111 | 33 | 13.97 | 2.36 | 1.69E-06 | 4.86E-05 |
| negative regulation of canonical Wnt signaling pathway | GO:0090090 | 54 | 16 | 6.80 | 2.35 | 7.22E-04 | 9.76E-03 |
| tissue morphogenesis | GO:0048729 | 571 | 169 | 71.87 | 2.35 | 1.49E-27 | 2.64E-25 |
| positive regulation of kinase activity | GO:0033674 | 98 | 29 | 12.34 | 2.35 | 8.30E-06 | 1.99E-04 |

| | | | | | | | |
|---|---|---|---|---|---|---|---|
| embryonic pattern specification | GO:0009880 | 71 | 21 | 8.94 | 2.35 | 1.88E-04 | 3.08E-03 |
| neuron migration | GO:0001764 | 44 | 13 | 5.54 | 2.35 | 2.28E-03 | 2.59E-02 |
| neural retina development | GO:0003407 | 105 | 31 | 13.22 | 2.35 | 4.04E-06 | 1.07E-04 |
| determination of heart left/right asymmetry | GO:0061371 | 204 | 60 | 25.68 | 2.34 | 1.68E-10 | 7.95E-09 |
| positive regulation of Wnt signaling pathway | GO:0030177 | 68 | 20 | 8.56 | 2.34 | 2.90E-04 | 4.48E-03 |
| positive regulation of transferase activity | GO:0051347 | 102 | 30 | 12.84 | 2.34 | 6.24E-06 | 1.57E-04 |
| tube development | GO:0035295 | 780 | 229 | 98.18 | 2.33 | 1.41E-36 | 4.54E-34 |
| positive regulation of phosphorylation | GO:0042327 | 174 | 51 | 21.90 | 2.33 | 5.26E-09 | 2.11E-07 |
| one-carbon metabolic process | GO:0006730 | 41 | 12 | 5.16 | 2.33 | 3.62E-03 | 3.75E-02 |
| regulation of extent of cell growth | GO:0061387 | 82 | 24 | 10.32 | 2.33 | 6.38E-05 | 1.24E-03 |
| camera-type eye morphogenesis | GO:0048593 | 195 | 57 | 24.54 | 2.32 | 6.08E-10 | 2.65E-08 |
| dorsal/ventral pattern formation | GO:0009953 | 154 | 45 | 19.38 | 2.32 | 5.44E-08 | 1.96E-06 |
| transmembrane receptor protein tyrosine kinase signaling pathway | GO:0007169 | 243 | 71 | 30.59 | 2.32 | 4.23E-12 | 2.44E-10 |
| animal organ morphogenesis | GO:0009887 | 839 | 245 | 105.60 | 2.32 | 1.10E-38 | 4.50E-36 |
| non-proteinogenic amino acid metabolic process | GO:0170041 | 48 | 14 | 6.04 | 2.32 | 1.80E-03 | 2.14E-02 |
| inorganic anion transport | GO:0015698 | 120 | 35 | 15.10 | 2.32 | 1.19E-06 | 3.54E-05 |
| enzyme-linked receptor protein signaling pathway | GO:0007167 | 350 | 102 | 44.05 | 2.32 | 1.66E-16 | 1.25E-14 |
| plasma membrane bounded cell projection morphogenesis | GO:0120039 | 447 | 130 | 56.26 | 2.31 | 1.15E-20 | 1.12E-18 |
| negative regulation of developmental process | GO:0051093 | 172 | 50 | 21.65 | 2.31 | 9.06E-09 | 3.52E-07 |
| cell projection morphogenesis | GO:0048858 | 451 | 131 | 56.77 | 2.31 | 8.42E-21 | 8.33E-19 |
| mesoderm development | GO:0007498 | 100 | 29 | 12.59 | 2.30 | 1.14E-05 | 2.64E-04 |
| axis specification | GO:0009798 | 76 | 22 | 9.57 | 2.30 | 1.52E-04 | 2.58E-03 |
| central nervous system neuron differentiation | GO:0021953 | 121 | 35 | 15.23 | 2.30 | 1.44E-06 | 4.22E-05 |
| neuron projection morphogenesis | GO:0048812 | 446 | 129 | 56.14 | 2.30 | 2.28E-20 | 2.16E-18 |
| convergent extension involved in gastrulation | GO:0060027 | 83 | 24 | 10.45 | 2.30 | 7.33E-05 | 1.41E-03 |
| transforming growth factor beta receptor superfamily signaling pathway | GO:0141091 | 90 | 26 | 11.33 | 2.30 | 3.57E-05 | 7.44E-04 |
| cardiac muscle cell differentiation | GO:0055007 | 52 | 15 | 6.55 | 2.29 | 2.33E-03 | 2.62E-02 |
| lymph vessel development | GO:0001945 | 66 | 19 | 8.31 | 2.29 | 4.97E-04 | 7.12E-03 |
| anterior/posterior pattern specification | GO:0009952 | 247 | 71 | 31.09 | 2.28 | 1.15E-11 | 6.33E-10 |
| anatomical structure homeostasis | GO:0060249 | 87 | 25 | 10.95 | 2.28 | 5.53E-05 | 1.10E-03 |
| tube morphogenesis | GO:0035239 | 589 | 169 | 74.14 | 2.28 | 6.25E-26 | 8.93E-24 |
| angiogenesis | GO:0001525 | 279 | 80 | 35.12 | 2.28 | 5.71E-13 | 3.57E-11 |
| regulation of cell projection organization | GO:0031344 | 231 | 66 | 29.08 | 2.27 | 8.30E-11 | 4.08E-09 |
| tissue homeostasis | GO:0001894 | 84 | 24 | 10.57 | 2.27 | 8.55E-05 | 1.61E-03 |
| retina development in camera-type eye | GO:0060041 | 273 | 78 | 34.36 | 2.27 | 1.35E-12 | 8.05E-11 |
| columnar/cuboidal epithelial cell differentiation | GO:0002065 | 42 | 12 | 5.29 | 2.27 | 4.50E-03 | 4.49E-02 |
| neuron fate commitment | GO:0048663 | 49 | 14 | 6.17 | 2.27 | 3.59E-03 | 3.73E-02 |
| positive regulation of phosphorus metabolic process | GO:0010562 | 186 | 53 | 23.41 | 2.26 | 5.70E-09 | 2.28E-07 |
| positive regulation of phosphate metabolic process | GO:0045937 | 186 | 53 | 23.41 | 2.26 | 5.70E-09 | 2.27E-07 |

| | | | | | | | |
|---|---|---|---|---|---|---|---|
| Wnt signaling pathway | GO:0016055 | 158 | 45 | 19.89 | 2.26 | 8.81E-08 | 3.12E-06 |
| negative regulation of cell differentiation | GO:0045596 | 123 | 35 | 15.48 | 2.26 | 3.23E-06 | 8.80E-05 |
| cell migration involved in gastrulation | GO:0042074 | 81 | 23 | 10.20 | 2.26 | 1.32E-04 | 2.28E-03 |
| regulation of plasma membrane bounded cell projection organization | GO:0120035 | 229 | 65 | 28.82 | 2.26 | 1.47E-10 | 7.00E-09 |
| negative regulation of cell development | GO:0010721 | 74 | 21 | 9.31 | 2.25 | 2.71E-04 | 4.21E-03 |
| growth | GO:0040007 | 296 | 84 | 37.26 | 2.25 | 3.22E-13 | 2.06E-11 |
| developmental growth | GO:0048589 | 296 | 84 | 37.26 | 2.25 | 3.22E-13 | 2.04E-11 |
| positive regulation of transcription by RNA polymerase II | GO:0045944 | 335 | 95 | 42.17 | 2.25 | 1.28E-14 | 8.73E-13 |
| negative regulation of cellular response to growth factor stimulus | GO:0090288 | 60 | 17 | 7.55 | 2.25 | 1.19E-03 | 1.50E-02 |
| appendage development | GO:0048736 | 120 | 34 | 15.10 | 2.25 | 4.90E-06 | 1.27E-04 |
| eye morphogenesis | GO:0048592 | 240 | 68 | 30.21 | 2.25 | 5.70E-11 | 2.87E-09 |
| lysosomal transport | GO:0007041 | 53 | 15 | 6.67 | 2.25 | 2.54E-03 | 2.85E-02 |
| brain development | GO:0007420 | 527 | 149 | 66.33 | 2.25 | 2.52E-22 | 2.98E-20 |
| negative regulation of response to external stimulus | GO:0032102 | 92 | 26 | 11.58 | 2.25 | 5.01E-05 | 1.01E-03 |
| swimming behavior | GO:0036269 | 71 | 20 | 8.94 | 2.24 | 4.19E-04 | 6.16E-03 |
| regulation of mRNA processing | GO:0050684 | 71 | 20 | 8.94 | 2.24 | 4.19E-04 | 6.15E-03 |
| regulation of hormone levels | GO:0010817 | 96 | 27 | 12.08 | 2.23 | 3.88E-05 | 7.97E-04 |
| cell-cell signaling by wnt | GO:0198738 | 160 | 45 | 20.14 | 2.23 | 1.23E-07 | 4.21E-06 |
| regulation of nervous system development | GO:0051960 | 196 | 55 | 24.67 | 2.23 | 6.45E-09 | 2.53E-07 |
| stem cell differentiation | GO:0048863 | 232 | 65 | 29.20 | 2.23 | 2.44E-10 | 1.12E-08 |
| pattern specification process | GO:0007389 | 644 | 180 | 81.06 | 2.22 | 5.14E-26 | 7.58E-24 |
| transition metal ion transport | GO:0000041 | 68 | 19 | 8.56 | 2.22 | 6.48E-04 | 8.93E-03 |
| regionalization | GO:0003002 | 629 | 175 | 79.17 | 2.21 | 4.20E-25 | 5.56E-23 |
| head development | GO:0060322 | 547 | 152 | 68.85 | 2.21 | 7.78E-22 | 8.86E-20 |
| central nervous system development | GO:0007417 | 738 | 205 | 92.89 | 2.21 | 4.04E-29 | 7.91E-27 |
| regulation of apoptotic signaling pathway | GO:2001233 | 83 | 23 | 10.45 | 2.20 | 1.84E-04 | 3.03E-03 |
| epithelium development | GO:0060429 | 942 | 261 | 118.57 | 2.20 | 7.48E-37 | 2.49E-34 |
| positive regulation of cell migration | GO:0030335 | 112 | 31 | 14.10 | 2.20 | 1.90E-05 | 4.20E-04 |
| cell fate commitment | GO:0045165 | 174 | 48 | 21.90 | 2.19 | 1.05E-07 | 3.68E-06 |
| synapse organization | GO:0050808 | 127 | 35 | 15.99 | 2.19 | 5.27E-06 | 1.35E-04 |
| regulation of cell development | GO:0060284 | 276 | 76 | 34.74 | 2.19 | 2.52E-11 | 1.31E-09 |
| determination of bilateral symmetry | GO:0009855 | 320 | 88 | 40.28 | 2.18 | 8.14E-13 | 5.02E-11 |
| specification of symmetry | GO:0009799 | 320 | 88 | 40.28 | 2.18 | 8.14E-13 | 4.98E-11 |
| RNA catabolic process | GO:0006401 | 91 | 25 | 11.45 | 2.18 | 1.72E-04 | 2.88E-03 |
| establishment of cell polarity | GO:0030010 | 51 | 14 | 6.42 | 2.18 | 4.39E-03 | 4.40E-02 |
| fin development | GO:0033333 | 113 | 31 | 14.22 | 2.18 | 2.14E-05 | 4.67E-04 |
| segmentation | GO:0035282 | 124 | 34 | 15.61 | 2.18 | 8.04E-06 | 1.94E-04 |
| microtubule cytoskeleton organization involved in mitosis | GO:1902850 | 73 | 20 | 9.19 | 2.18 | 5.72E-04 | 7.99E-03 |
| cellular response to organic cyclic compound | GO:0071407 | 73 | 20 | 9.19 | 2.18 | 5.72E-04 | 7.98E-03 |
| determination of left/right symmetry | GO:0007368 | 292 | 80 | 36.75 | 2.18 | 8.08E-12 | 4.49E-10 |

| | | | | | | | |
|---|---|---|---|---|---|---|---|
| embryo development | GO:0009790 | 1212 | 332 | 152.55 | 2.18 | 8.98E-46 | 4.49E-43 |
| cell morphogenesis | GO:0000902 | 548 | 150 | 68.98 | 2.17 | 6.83E-21 | 6.90E-19 |
| left/right pattern formation | GO:0060972 | 296 | 81 | 37.26 | 2.17 | 6.21E-12 | 3.52E-10 |
| neuron projection development | GO:0031175 | 549 | 150 | 69.10 | 2.17 | 7.68E-21 | 7.68E-19 |
| somitogenesis | GO:0001756 | 103 | 28 | 12.96 | 2.16 | 6.64E-05 | 1.29E-03 |
| glial cell differentiation | GO:0010001 | 140 | 38 | 17.62 | 2.16 | 4.14E-06 | 1.09E-04 |
| cellular response to external stimulus | GO:0071496 | 59 | 16 | 7.43 | 2.15 | 2.39E-03 | 2.69E-02 |
| heart development | GO:0007507 | 583 | 158 | 73.38 | 2.15 | 1.53E-21 | 1.66E-19 |
| convergent extension | GO:0060026 | 155 | 42 | 19.51 | 2.15 | 1.10E-06 | 3.32E-05 |
| blood vessel morphogenesis | GO:0048514 | 358 | 97 | 45.06 | 2.15 | 1.16E-13 | 7.62E-12 |
| nephron development | GO:0072006 | 96 | 26 | 12.08 | 2.15 | 1.36E-04 | 2.35E-03 |
| vascular process in circulatory system | GO:0003018 | 74 | 20 | 9.31 | 2.15 | 6.75E-04 | 9.24E-03 |
| neuron differentiation | GO:0030182 | 903 | 244 | 113.66 | 2.15 | 1.94E-32 | 4.26E-30 |
| cell differentiation in spinal cord | GO:0021515 | 63 | 17 | 7.93 | 2.14 | 1.79E-03 | 2.14E-02 |
| regulation of growth | GO:0040008 | 141 | 38 | 17.75 | 2.14 | 4.53E-06 | 1.17E-04 |
| regulation of cellular component size | GO:0032535 | 208 | 56 | 26.18 | 2.14 | 2.17E-08 | 8.02E-07 |
| neuron development | GO:0048666 | 725 | 195 | 91.25 | 2.14 | 8.82E-26 | 1.22E-23 |
| tissue development | GO:0009888 | 1511 | 406 | 190.19 | 2.13 | 8.11E-54 | 4.30E-51 |
| positive regulation of RNA biosynthetic process | GO:1902680 | 488 | 131 | 61.42 | 2.13 | 1.16E-17 | 9.60E-16 |
| positive regulation of DNA-templated transcription | GO:0045893 | 488 | 131 | 61.42 | 2.13 | 1.16E-17 | 9.51E-16 |
| muscle tissue development | GO:0060537 | 220 | 59 | 27.69 | 2.13 | 1.51E-08 | 5.67E-07 |
| glial cell development | GO:0021782 | 97 | 26 | 12.21 | 2.13 | 1.53E-04 | 2.58E-03 |
| cellular response to extracellular stimulus | GO:0031668 | 56 | 15 | 7.05 | 2.13 | 3.69E-03 | 3.79E-02 |
| generation of neurons | GO:0048699 | 934 | 250 | 117.56 | 2.13 | 1.08E-32 | 2.42E-30 |
| cell fate specification | GO:0001708 | 101 | 27 | 12.71 | 2.12 | 1.14E-04 | 2.02E-03 |
| cellular anatomical entity morphogenesis | GO:0032989 | 562 | 150 | 70.74 | 2.12 | 8.77E-20 | 7.90E-18 |
| synaptic vesicle cycle | GO:0099504 | 90 | 24 | 11.33 | 2.12 | 3.12E-04 | 4.78E-03 |
| vesicle-mediated transport in synapse | GO:0099003 | 90 | 24 | 11.33 | 2.12 | 3.12E-04 | 4.77E-03 |
| regulation of lipid metabolic process | GO:0019216 | 60 | 16 | 7.55 | 2.12 | 2.77E-03 | 3.04E-02 |
| signal release | GO:0023061 | 75 | 20 | 9.44 | 2.12 | 1.24E-03 | 1.55E-02 |
| gliogenesis | GO:0042063 | 150 | 40 | 18.88 | 2.12 | 2.86E-06 | 7.95E-05 |
| blood vessel development | GO:0001568 | 413 | 110 | 51.98 | 2.12 | 9.32E-15 | 6.45E-13 |
| mRNA catabolic process | GO:0006402 | 79 | 21 | 9.94 | 2.11 | 8.87E-04 | 1.17E-02 |
| circulatory system development | GO:0072359 | 981 | 260 | 123.48 | 2.11 | 2.99E-33 | 7.69E-31 |
| nervous system development | GO:0007399 | 1684 | 445 | 211.96 | 2.10 | 5.68E-57 | 3.93E-54 |
| neuron projection extension | GO:1990138 | 57 | 15 | 7.17 | 2.09 | 4.26E-03 | 4.32E-02 |
| neurogenesis | GO:0022008 | 1070 | 281 | 134.68 | 2.09 | 4.92E-35 | 1.43E-32 |
| anatomical structure formation involved in morphogenesis | GO:0048646 | 808 | 212 | 101.70 | 2.08 | 1.68E-26 | 2.61E-24 |
| regulation of mRNA splicing, via spliceosome | GO:0048024 | 61 | 16 | 7.68 | 2.08 | 3.22E-03 | 3.39E-02 |
| vasculature development | GO:0001944 | 492 | 129 | 61.93 | 2.08 | 1.96E-16 | 1.47E-14 |
| vacuolar transport | GO:0007034 | 103 | 27 | 12.96 | 2.08 | 1.51E-04 | 2.56E-03 |

| | | | | | | | |
|---|---|---|---|---|---|---|---|
| regulation of anatomical structure size | GO:0090066 | 264 | 69 | 33.23 | 2.08 | 1.97E-09 | 8.27E-08 |
| kidney epithelium development | GO:0072073 | 69 | 18 | 8.68 | 2.07 | 2.79E-03 | 3.05E-02 |
| nuclear-transcribed mRNA catabolic process | GO:0000956 | 69 | 18 | 8.68 | 2.07 | 2.79E-03 | 3.04E-02 |
| ceramide metabolic process | GO:0006672 | 77 | 20 | 9.69 | 2.06 | 1.46E-03 | 1.78E-02 |
| protein lipidation | GO:0006497 | 81 | 21 | 10.20 | 2.06 | 1.08E-03 | 1.37E-02 |
| positive regulation of cell differentiation | GO:0045597 | 178 | 46 | 22.40 | 2.05 | 1.94E-06 | 5.53E-05 |
| positive regulation of cell motility | GO:2000147 | 120 | 31 | 15.10 | 2.05 | 7.70E-05 | 1.46E-03 |
| regulation of locomotion | GO:0040012 | 271 | 70 | 34.11 | 2.05 | 2.97E-09 | 1.23E-07 |
| regulation of cell migration | GO:0030334 | 233 | 60 | 29.33 | 2.05 | 4.34E-08 | 1.57E-06 |
| positive regulation of developmental process | GO:0051094 | 268 | 69 | 33.73 | 2.05 | 4.39E-09 | 1.78E-07 |
| glomerulus development | GO:0032835 | 70 | 18 | 8.81 | 2.04 | 3.01E-03 | 3.28E-02 |
| digestive tract development | GO:0048565 | 105 | 27 | 13.22 | 2.04 | 2.85E-04 | 4.43E-03 |
| receptor-mediated endocytosis | GO:0006898 | 74 | 19 | 9.31 | 2.04 | 2.20E-03 | 2.51E-02 |
| negative regulation of locomotion | GO:0040013 | 78 | 20 | 9.82 | 2.04 | 1.62E-03 | 1.97E-02 |
| gastrulation | GO:0007369 | 242 | 62 | 30.46 | 2.04 | 2.85E-08 | 1.05E-06 |
| regulation of Wnt signaling pathway | GO:0030111 | 172 | 44 | 21.65 | 2.03 | 4.17E-06 | 1.09E-04 |
| regulation of cell differentiation | GO:0045595 | 493 | 126 | 62.05 | 2.03 | 2.74E-15 | 1.97E-13 |
| anatomical structure morphogenesis | GO:0009653 | 2211 | 565 | 278.29 | 2.03 | 5.30E-68 | 3.98E-65 |
| cell migration | GO:0016477 | 619 | 158 | 77.91 | 2.03 | 9.91E-19 | 8.50E-17 |
| negative regulation of intracellular signal transduction | GO:1902532 | 149 | 38 | 18.75 | 2.03 | 1.68E-05 | 3.74E-04 |
| positive regulation of RNA metabolic process | GO:0051254 | 557 | 142 | 70.11 | 2.03 | 5.83E-17 | 4.60E-15 |
| positive regulation of protein phosphorylation | GO:0001934 | 110 | 28 | 13.85 | 2.02 | 2.30E-04 | 3.66E-03 |
| cellular response to endogenous stimulus | GO:0071495 | 385 | 98 | 48.46 | 2.02 | 5.91E-12 | 3.37E-10 |
| regulation of canonical Wnt signaling pathway | GO:0060828 | 118 | 30 | 14.85 | 2.02 | 1.29E-04 | 2.25E-03 |
| positive regulation of locomotion | GO:0040017 | 126 | 32 | 15.86 | 2.02 | 7.48E-05 | 1.42E-03 |
| regulation of anatomical structure morphogenesis | GO:0022603 | 327 | 83 | 41.16 | 2.02 | 2.39E-10 | 1.11E-08 |
| response to hypoxia | GO:0001666 | 71 | 18 | 8.94 | 2.01 | 3.30E-03 | 3.46E-02 |
| lipoprotein biosynthetic process | GO:0042158 | 83 | 21 | 10.45 | 2.01 | 1.37E-03 | 1.68E-02 |
| negative regulation of Wnt signaling pathway | GO:0030178 | 83 | 21 | 10.45 | 2.01 | 1.37E-03 | 1.68E-02 |
| regulation of cell growth | GO:0001558 | 103 | 26 | 12.96 | 2.01 | 4.61E-04 | 6.70E-03 |
| response to organic cyclic compound | GO:0014070 | 123 | 31 | 15.48 | 2.00 | 1.69E-04 | 2.84E-03 |
| positive regulation of nucleobase-containing compound metabolic process | GO:0045935 | 596 | 150 | 75.02 | 2.00 | 2.54E-17 | 2.04E-15 |
| sensory organ development | GO:0007423 | 748 | 188 | 94.15 | 2.00 | 2.68E-21 | 2.84E-19 |
| positive regulation of cellular metabolic process | GO:0031325 | 926 | 232 | 116.55 | 1.99 | 8.57E-26 | 1.20E-23 |
| cell growth | GO:0016049 | 68 | 17 | 8.56 | 1.99 | 4.96E-03 | 4.84E-02 |
| spinal cord development | GO:0021510 | 88 | 22 | 11.08 | 1.99 | 1.75E-03 | 2.11E-02 |
| response to decreased oxygen levels | GO:0036293 | 72 | 18 | 9.06 | 1.99 | 3.66E-03 | 3.77E-02 |
| cell motility | GO:0048870 | 670 | 167 | 84.33 | 1.98 | 1.03E-18 | 8.78E-17 |
| negative regulation of response to stimulus | GO:0048585 | 542 | 135 | 68.22 | 1.98 | 3.39E-15 | 2.42E-13 |

| | | | | | | | |
|---|---|---|---|---|---|---|---|
| positive regulation of cell population proliferation | GO:0008284 | 161 | 40 | 20.26 | 1.97 | 2.08E-05 | 4.55E-04 |
| pancreas development | GO:0031016 | 157 | 39 | 19.76 | 1.97 | 2.80E-05 | 5.95E-04 |
| photoreceptor cell differentiation | GO:0046530 | 109 | 27 | 13.72 | 1.97 | 4.28E-04 | 6.27E-03 |
| germ cell development | GO:0007281 | 101 | 25 | 12.71 | 1.97 | 7.54E-04 | 1.01E-02 |
| secretion | GO:0046903 | 190 | 47 | 23.91 | 1.97 | 4.46E-06 | 1.16E-04 |
| response to endogenous stimulus | GO:0009719 | 437 | 108 | 55.00 | 1.96 | 3.23E-12 | 1.89E-10 |
| negative regulation of transcription by RNA polymerase II | GO:0000122 | 255 | 63 | 32.10 | 1.96 | 1.07E-07 | 3.72E-06 |
| response to oxygen levels | GO:0070482 | 77 | 19 | 9.69 | 1.96 | 3.08E-03 | 3.26E-02 |
| positive regulation of cellular component organization | GO:0051130 | 223 | 55 | 28.07 | 1.96 | 7.22E-07 | 2.25E-05 |
| regulation of actin cytoskeleton organization | GO:0032956 | 187 | 46 | 23.54 | 1.95 | 6.51E-06 | 1.63E-04 |
| regulation of multicellular organismal development | GO:2000026 | 456 | 112 | 57.40 | 1.95 | 1.95E-12 | 1.15E-10 |
| chromosome segregation | GO:0007059 | 143 | 35 | 18.00 | 1.94 | 1.09E-04 | 1.95E-03 |
| calcium ion transmembrane transport | GO:0070588 | 143 | 35 | 18.00 | 1.94 | 1.09E-04 | 1.95E-03 |
| negative regulation of cellular component organization | GO:0051129 | 184 | 45 | 23.16 | 1.94 | 9.51E-06 | 2.25E-04 |
| positive regulation of macromolecule biosynthetic process | GO:0010557 | 655 | 160 | 82.44 | 1.94 | 5.14E-17 | 4.10E-15 |
| regulation of actin filament organization | GO:0110053 | 131 | 32 | 16.49 | 1.94 | 1.76E-04 | 2.92E-03 |
| neurotransmitter transport | GO:0006836 | 82 | 20 | 10.32 | 1.94 | 3.77E-03 | 3.86E-02 |
| regulation of actin filament-based process | GO:0032970 | 189 | 46 | 23.79 | 1.93 | 7.97E-06 | 1.93E-04 |
| canonical Wnt signaling pathway | GO:0060070 | 74 | 18 | 9.31 | 1.93 | 4.62E-03 | 4.58E-02 |
| monosaccharide metabolic process | GO:0005996 | 74 | 18 | 9.31 | 1.93 | 4.62E-03 | 4.57E-02 |
| regulation of transferase activity | GO:0051338 | 185 | 45 | 23.29 | 1.93 | 1.05E-05 | 2.45E-04 |
| membrane lipid metabolic process | GO:0006643 | 144 | 35 | 18.12 | 1.93 | 1.17E-04 | 2.05E-03 |
| negative regulation of signal transduction | GO:0009968 | 445 | 108 | 56.01 | 1.93 | 9.53E-12 | 5.26E-10 |
| positive regulation of cellular biosynthetic process | GO:0031328 | 668 | 162 | 84.08 | 1.93 | 6.39E-17 | 5.00E-15 |
| negative regulation of cell communication | GO:0010648 | 458 | 111 | 57.65 | 1.93 | 6.34E-12 | 3.57E-10 |
| negative regulation of signaling | GO:0023057 | 458 | 111 | 57.65 | 1.93 | 6.34E-12 | 3.54E-10 |
| sodium ion transport | GO:0006814 | 128 | 31 | 16.11 | 1.92 | 2.59E-04 | 4.03E-03 |
| epithelial cell differentiation | GO:0030855 | 256 | 62 | 32.22 | 1.92 | 3.23E-07 | 1.05E-05 |
| positive regulation of metabolic process | GO:0009893 | 1037 | 251 | 130.53 | 1.92 | 1.40E-25 | 1.91E-23 |
| positive regulation of biosynthetic process | GO:0009891 | 670 | 162 | 84.33 | 1.92 | 7.71E-17 | 5.98E-15 |
| somite development | GO:0061053 | 149 | 36 | 18.75 | 1.92 | 9.52E-05 | 1.74E-03 |
| regulation of developmental process | GO:0050793 | 895 | 215 | 112.65 | 1.91 | 1.37E-21 | 1.50E-19 |
| positive regulation of nitrogen compound metabolic process | GO:0051173 | 854 | 205 | 107.49 | 1.91 | 1.65E-20 | 1.58E-18 |
| secretion by cell | GO:0032940 | 175 | 42 | 22.03 | 1.91 | 2.94E-05 | 6.21E-04 |
| system development | GO:0048731 | 3290 | 787 | 414.11 | 1.90 | 4.80E-83 | 4.80E-80 |
| regulation of secretion | GO:0051046 | 134 | 32 | 16.87 | 1.90 | 3.31E-04 | 4.99E-03 |
| regulation of membrane potential | GO:0042391 | 172 | 41 | 21.65 | 1.89 | 4.27E-05 | 8.71E-04 |
| regulation of cell motility | GO:2000145 | 252 | 60 | 31.72 | 1.89 | 7.98E-07 | 2.45E-05 |
| phosphatidylinositol biosynthetic process | GO:0006661 | 84 | 20 | 10.57 | 1.89 | 4.36E-03 | 4.42E-02 |

| | | | | | | | |
|---|---|---|---|---|---|---|---|
| cell-cell signaling | GO:0007267 | 534 | 127 | 67.21 | 1.89 | 7.60E-13 | 4.71E-11 |
| establishment of organelle localization | GO:0051656 | 177 | 42 | 22.28 | 1.89 | 5.14E-05 | 1.03E-03 |
| developmental growth involved in morphogenesis | GO:0060560 | 118 | 28 | 14.85 | 1.89 | 7.33E-04 | 9.89E-03 |
| establishment or maintenance of cell polarity | GO:0007163 | 114 | 27 | 14.35 | 1.88 | 9.70E-04 | 1.26E-02 |
| regulation of phosphorylation | GO:0042325 | 279 | 66 | 35.12 | 1.88 | 3.91E-07 | 1.25E-05 |
| sensory system development | GO:0048880 | 664 | 157 | 83.58 | 1.88 | 2.01E-15 | 1.47E-13 |
| plasma membrane bounded cell projection organization | GO:0120036 | 885 | 209 | 111.39 | 1.88 | 4.87E-20 | 4.57E-18 |
| regulation of cell population proliferation | GO:0042127 | 305 | 72 | 38.39 | 1.88 | 1.19E-07 | 4.10E-06 |
| wound healing | GO:0042060 | 89 | 21 | 11.20 | 1.87 | 3.58E-03 | 3.72E-02 |
| multicellular organism development | GO:0007275 | 3850 | 908 | 484.59 | 1.87 | 2.07E-94 | 3.11E-91 |
| cellular response to organonitrogen compound | GO:0071417 | 102 | 24 | 12.84 | 1.87 | 2.33E-03 | 2.63E-02 |
| muscle structure development | GO:0061061 | 374 | 88 | 47.07 | 1.87 | 5.81E-09 | 2.30E-07 |
| regulation of RNA splicing | GO:0043484 | 102 | 24 | 12.84 | 1.87 | 2.33E-03 | 2.63E-02 |
| positive regulation of proteolysis | GO:0045862 | 85 | 20 | 10.70 | 1.87 | 4.77E-03 | 4.72E-02 |
| regulation of actin filament polymerization | GO:0030833 | 98 | 23 | 12.34 | 1.86 | 3.15E-03 | 3.33E-02 |
| positive regulation of macromolecule metabolic process | GO:0010604 | 930 | 218 | 117.06 | 1.86 | 1.49E-20 | 1.44E-18 |
| actin filament organization | GO:0007015 | 252 | 59 | 31.72 | 1.86 | 2.04E-06 | 5.80E-05 |
| cell development | GO:0048468 | 1824 | 427 | 229.58 | 1.86 | 5.31E-40 | 2.27E-37 |
| endocytosis | GO:0006897 | 278 | 65 | 34.99 | 1.86 | 6.26E-07 | 1.97E-05 |
| cell projection organization | GO:0030030 | 907 | 212 | 114.16 | 1.86 | 8.74E-20 | 7.95E-18 |
| muscle cell differentiation | GO:0042692 | 261 | 61 | 32.85 | 1.86 | 1.27E-06 | 3.75E-05 |
| sphingolipid metabolic process | GO:0006665 | 107 | 25 | 13.47 | 1.86 | 1.88E-03 | 2.22E-02 |
| cellular developmental process | GO:0048869 | 2509 | 586 | 315.80 | 1.86 | 9.02E-56 | 5.80E-53 |
| regulation of supramolecular fiber organization | GO:1902903 | 167 | 39 | 21.02 | 1.86 | 1.41E-04 | 2.42E-03 |
| cell differentiation | GO:0030154 | 2505 | 585 | 315.30 | 1.86 | 1.24E-55 | 7.47E-53 |
| blood circulation | GO:0008015 | 223 | 52 | 28.07 | 1.85 | 9.52E-06 | 2.25E-04 |
| regulation of kinase activity | GO:0043549 | 176 | 41 | 22.15 | 1.85 | 8.30E-05 | 1.57E-03 |
| animal organ development | GO:0048513 | 2532 | 589 | 318.70 | 1.85 | 1.71E-55 | 9.61E-53 |
| developmental process involved in reproduction | GO:0003006 | 172 | 40 | 21.65 | 1.85 | 1.11E-04 | 1.96E-03 |
| positive regulation of protein modification process | GO:0031401 | 138 | 32 | 17.37 | 1.84 | 6.73E-04 | 9.23E-03 |
| cell surface receptor signaling pathway | GO:0007166 | 1075 | 248 | 135.31 | 1.83 | 4.04E-22 | 4.72E-20 |
| regulation of actin polymerization or depolymerization | GO:0008064 | 100 | 23 | 12.59 | 1.83 | 3.62E-03 | 3.75E-02 |
| mitotic cell cycle process | GO:1903047 | 244 | 56 | 30.71 | 1.82 | 6.36E-06 | 1.59E-04 |
| camera-type eye development | GO:0043010 | 475 | 109 | 59.79 | 1.82 | 3.04E-10 | 1.39E-08 |
| membrane lipid biosynthetic process | GO:0046467 | 109 | 25 | 13.72 | 1.82 | 3.27E-03 | 3.44E-02 |
| actin cytoskeleton organization | GO:0030036 | 463 | 106 | 58.28 | 1.82 | 7.05E-10 | 3.06E-08 |
| multicellular organismal-level homeostasis | GO:0048871 | 249 | 57 | 31.34 | 1.82 | 7.45E-06 | 1.84E-04 |
| glycerophospholipid biosynthetic process | GO:0046474 | 118 | 27 | 14.85 | 1.82 | 1.89E-03 | 2.24E-02 |
| actin filament-based process | GO:0030029 | 481 | 110 | 60.54 | 1.82 | 3.82E-10 | 1.71E-08 |
| regulation of secretion by cell | GO:1903530 | 127 | 29 | 15.99 | 1.81 | 1.17E-03 | 1.47E-02 |

| | | | | | | | |
|---|---|---|---|---|---|---|---|
| cytoskeleton-dependent intracellular transport | GO:0030705 | 101 | 23 | 12.71 | 1.81 | 3.93E-03 | 4.01E-02 |
| anatomical structure development | GO:0048856 | 4903 | 1116 | 617.13 | 1.81 | 5.50E-110 | 9.90E-107 |
| small GTPase-mediated signal transduction | GO:0007264 | 176 | 40 | 22.15 | 1.81 | 2.13E-04 | 3.45E-03 |
| export from cell | GO:0140352 | 198 | 45 | 24.92 | 1.81 | 8.56E-05 | 1.61E-03 |
| circulatory system process | GO:0003013 | 229 | 52 | 28.82 | 1.80 | 2.05E-05 | 4.50E-04 |
| developmental process | GO:0032502 | 5052 | 1144 | 635.89 | 1.80 | 7.06E-112 | 1.59E-108 |
| digestive system development | GO:0055123 | 146 | 33 | 18.38 | 1.80 | 6.60E-04 | 9.07E-03 |
| eye development | GO:0001654 | 549 | 124 | 69.10 | 1.79 | 6.92E-11 | 3.44E-09 |
| visual system development | GO:0150063 | 549 | 124 | 69.10 | 1.79 | 6.92E-11 | 3.42E-09 |
| regulation of actin filament length | GO:0030832 | 102 | 23 | 12.84 | 1.79 | 4.30E-03 | 4.36E-02 |
| phospholipid biosynthetic process | GO:0008654 | 142 | 32 | 17.87 | 1.79 | 8.66E-04 | 1.14E-02 |
| regulation of protein polymerization | GO:0032271 | 111 | 25 | 13.97 | 1.79 | 3.61E-03 | 3.75E-02 |
| cellular response to nitrogen compound | GO:1901699 | 129 | 29 | 16.24 | 1.79 | 1.90E-03 | 2.23E-02 |
| cell division | GO:0051301 | 263 | 59 | 33.10 | 1.78 | 8.68E-06 | 2.07E-04 |
| regulation of response to external stimulus | GO:0032101 | 290 | 65 | 36.50 | 1.78 | 2.85E-06 | 7.94E-05 |
| muscle organ development | GO:0007517 | 193 | 43 | 24.29 | 1.77 | 1.76E-04 | 2.92E-03 |
| calcium ion transport | GO:0006816 | 162 | 36 | 20.39 | 1.77 | 7.47E-04 | 1.01E-02 |
| regulation of cell communication | GO:0010646 | 1363 | 302 | 171.56 | 1.76 | 5.52E-24 | 6.90E-22 |
| response to wounding | GO:0009611 | 136 | 30 | 17.12 | 1.75 | 1.75E-03 | 2.10E-02 |
| regulation of signal transduction | GO:0009966 | 1215 | 268 | 152.93 | 1.75 | 5.01E-21 | 5.25E-19 |
| regulation of signaling | GO:0023051 | 1368 | 301 | 172.19 | 1.75 | 2.39E-23 | 2.90E-21 |
| organelle localization | GO:0051640 | 250 | 55 | 31.47 | 1.75 | 3.10E-05 | 6.49E-04 |
| carbohydrate derivative catabolic process | GO:1901136 | 141 | 31 | 17.75 | 1.75 | 1.96E-03 | 2.29E-02 |
| regulation of phosphate metabolic process | GO:0019220 | 314 | 69 | 39.52 | 1.75 | 3.02E-06 | 8.28E-05 |
| regulation of phosphorus metabolic process | GO:0051174 | 314 | 69 | 39.52 | 1.75 | 3.02E-06 | 8.26E-05 |
| positive regulation of multicellular organismal process | GO:0051240 | 296 | 65 | 37.26 | 1.74 | 5.75E-06 | 1.45E-04 |
| behavior | GO:0007610 | 237 | 52 | 29.83 | 1.74 | 6.81E-05 | 1.31E-03 |
| nucleobase-containing compound catabolic process | GO:0034655 | 196 | 43 | 24.67 | 1.74 | 3.05E-04 | 4.69E-03 |
| multicellular organismal process | GO:0032501 | 4814 | 1056 | 605.93 | 1.74 | 8.21E-92 | 1.06E-88 |
| mRNA metabolic process | GO:0016071 | 383 | 84 | 48.21 | 1.74 | 3.33E-07 | 1.08E-05 |
| regulation of biological quality | GO:0065008 | 1082 | 237 | 136.19 | 1.74 | 3.01E-18 | 2.51E-16 |
| cellular homeostasis | GO:0019725 | 274 | 60 | 34.49 | 1.74 | 1.43E-05 | 3.23E-04 |
| positive regulation of catabolic process | GO:0009896 | 128 | 28 | 16.11 | 1.74 | 3.01E-03 | 3.28E-02 |
| regulation of response to stimulus | GO:0048583 | 1568 | 343 | 197.36 | 1.74 | 4.27E-26 | 6.40E-24 |
| aromatic compound catabolic process | GO:0019439 | 243 | 53 | 30.59 | 1.73 | 5.63E-05 | 1.11E-03 |
| cell cycle process | GO:0022402 | 424 | 92 | 53.37 | 1.72 | 1.23E-07 | 4.21E-06 |
| homeostatic process | GO:0042592 | 655 | 142 | 82.44 | 1.72 | 5.02E-11 | 2.55E-09 |
| cellular response to hormone stimulus | GO:0032870 | 180 | 39 | 22.66 | 1.72 | 6.28E-04 | 8.68E-03 |
| heterocycle catabolic process | GO:0046700 | 222 | 48 | 27.94 | 1.72 | 1.48E-04 | 2.53E-03 |
| cellular nitrogen compound catabolic process | GO:0044270 | 222 | 48 | 27.94 | 1.72 | 1.48E-04 | 2.53E-03 |

| | | | | | | | |
|---|---|---|---|---|---|---|---|
| modulation of chemical synaptic transmission | GO:0050804 | 153 | 33 | 19.26 | 1.71 | 1.98E-03 | 2.31E-02 |
| regulation of trans-synaptic signaling | GO:0099177 | 153 | 33 | 19.26 | 1.71 | 1.98E-03 | 2.31E-02 |
| cell junction organization | GO:0034330 | 232 | 50 | 29.20 | 1.71 | 1.33E-04 | 2.30E-03 |
| synaptic signaling | GO:0099536 | 297 | 64 | 37.38 | 1.71 | 1.36E-05 | 3.08E-04 |
| negative regulation of RNA biosynthetic process | GO:1902679 | 372 | 80 | 46.82 | 1.71 | 1.25E-06 | 3.72E-05 |
| negative regulation of DNA-templated transcription | GO:0045892 | 372 | 80 | 46.82 | 1.71 | 1.25E-06 | 3.70E-05 |
| cell-cell adhesion | GO:0098609 | 335 | 72 | 42.17 | 1.71 | 4.25E-06 | 1.11E-04 |
| phosphatidylinositol metabolic process | GO:0046488 | 135 | 29 | 16.99 | 1.71 | 3.79E-03 | 3.87E-02 |
| regulation of GTPase activity | GO:0043087 | 163 | 35 | 20.52 | 1.71 | 1.29E-03 | 1.59E-02 |
| RNA splicing | GO:0008380 | 247 | 53 | 31.09 | 1.70 | 9.70E-05 | 1.76E-03 |
| regulation of cellular component biogenesis | GO:0044087 | 280 | 60 | 35.24 | 1.70 | 3.78E-05 | 7.78E-04 |
| multicellular organism reproduction | GO:0032504 | 178 | 38 | 22.40 | 1.70 | 9.25E-04 | 1.21E-02 |
| chemical homeostasis | GO:0048878 | 389 | 83 | 48.96 | 1.70 | 1.40E-06 | 4.13E-05 |
| import into cell | GO:0098657 | 408 | 87 | 51.35 | 1.69 | 7.80E-07 | 2.40E-05 |
| apoptotic process | GO:0006915 | 249 | 53 | 31.34 | 1.69 | 1.07E-04 | 1.92E-03 |
| regulation of multicellular organismal process | GO:0051239 | 838 | 178 | 105.48 | 1.69 | 1.16E-12 | 7.03E-11 |
| regulation of transcription by RNA polymerase II | GO:0006357 | 1810 | 384 | 227.82 | 1.69 | 1.28E-26 | 2.03E-24 |
| cell cycle | GO:0007049 | 632 | 134 | 79.55 | 1.68 | 8.85E-10 | 3.83E-08 |
| organic cyclic compound catabolic process | GO:1901361 | 250 | 53 | 31.47 | 1.68 | 1.14E-04 | 2.02E-03 |
| negative regulation of RNA metabolic process | GO:0051253 | 406 | 86 | 51.10 | 1.68 | 1.10E-06 | 3.31E-05 |
| organic acid catabolic process | GO:0016054 | 156 | 33 | 19.64 | 1.68 | 2.32E-03 | 2.63E-02 |
| carboxylic acid catabolic process | GO:0046395 | 156 | 33 | 19.64 | 1.68 | 2.32E-03 | 2.63E-02 |
| regulation of cellular component organization | GO:0051128 | 785 | 166 | 98.81 | 1.68 | 1.16E-11 | 6.34E-10 |
| metal ion transport | GO:0030001 | 484 | 102 | 60.92 | 1.67 | 1.18E-07 | 4.08E-06 |
| positive regulation of cellular process | GO:0048522 | 1918 | 404 | 241.41 | 1.67 | 2.47E-27 | 4.19E-25 |
| cell adhesion | GO:0007155 | 570 | 120 | 71.74 | 1.67 | 9.99E-09 | 3.82E-07 |
| inorganic ion transmembrane transport | GO:0098660 | 513 | 108 | 64.57 | 1.67 | 5.76E-08 | 2.07E-06 |
| hemopoiesis | GO:0030097 | 457 | 96 | 57.52 | 1.67 | 3.49E-07 | 1.13E-05 |
| microtubule-based movement | GO:0007018 | 219 | 46 | 27.57 | 1.67 | 4.46E-04 | 6.52E-03 |
| chemical synaptic transmission | GO:0007268 | 281 | 59 | 35.37 | 1.67 | 6.25E-05 | 1.23E-03 |
| anterograde trans-synaptic signaling | GO:0098916 | 281 | 59 | 35.37 | 1.67 | 6.25E-05 | 1.23E-03 |
| regulation of cytoskeleton organization | GO:0051493 | 267 | 56 | 33.61 | 1.67 | 1.23E-04 | 2.14E-03 |
| mRNA processing | GO:0006397 | 310 | 65 | 39.02 | 1.67 | 3.04E-05 | 6.42E-04 |
| cell-cell adhesion via plasma-membrane adhesion molecules | GO:0098742 | 186 | 39 | 23.41 | 1.67 | 1.19E-03 | 1.49E-02 |
| regulation of system process | GO:0044057 | 148 | 31 | 18.63 | 1.66 | 3.94E-03 | 4.01E-02 |
| multicellular organismal reproductive process | GO:0048609 | 172 | 36 | 21.65 | 1.66 | 2.45E-03 | 2.75E-02 |
| negative regulation of multicellular organismal process | GO:0051241 | 206 | 43 | 25.93 | 1.66 | 6.96E-04 | 9.49E-03 |
| response to oxygen-containing compound | GO:1901700 | 345 | 72 | 43.42 | 1.66 | 1.28E-05 | 2.93E-04 |
| striated muscle cell differentiation | GO:0051146 | 211 | 44 | 26.56 | 1.66 | 7.57E-04 | 1.02E-02 |
| monoatomic ion transport | GO:0006811 | 764 | 159 | 96.16 | 1.65 | 9.43E-11 | 4.61E-09 |

| | | | | | | | |
|---|---|---|---|---|---|---|---|
| mitotic cell cycle | GO:0000278 | 309 | 64 | 38.89 | 1.65 | 6.19E-05 | 1.22E-03 |
| trans-synaptic signaling | GO:0099537 | 285 | 59 | 35.87 | 1.64 | 1.01E-04 | 1.83E-03 |
| response to hormone | GO:0009725 | 232 | 48 | 29.20 | 1.64 | 4.57E-04 | 6.65E-03 |
| regulation of proteolysis | GO:0030162 | 170 | 35 | 21.40 | 1.64 | 3.40E-03 | 3.56E-02 |
| monoatomic ion transmembrane transport | GO:0034220 | 545 | 112 | 68.60 | 1.63 | 1.40E-07 | 4.73E-06 |
| regulation of cell cycle process | GO:0010564 | 219 | 45 | 27.57 | 1.63 | 9.51E-04 | 1.24E-02 |
| DNA-templated transcription | GO:0006351 | 190 | 39 | 23.91 | 1.63 | 1.94E-03 | 2.27E-02 |
| regulation of mRNA metabolic process | GO:1903311 | 156 | 32 | 19.64 | 1.63 | 4.99E-03 | 4.86E-02 |
| cell death | GO:0008219 | 278 | 57 | 34.99 | 1.63 | 1.75E-04 | 2.91E-03 |
| programmed cell death | GO:0012501 | 278 | 57 | 34.99 | 1.63 | 1.75E-04 | 2.91E-03 |
| regulation of catabolic process | GO:0009894 | 264 | 54 | 33.23 | 1.63 | 3.50E-04 | 5.25E-03 |
| regulation of cellular metabolic process | GO:0031323 | 3302 | 675 | 415.62 | 1.62 | 1.39E-42 | 6.24E-40 |
| lipid biosynthetic process | GO:0008610 | 362 | 74 | 45.56 | 1.62 | 2.08E-05 | 4.55E-04 |
| regulation of catalytic activity | GO:0050790 | 511 | 104 | 64.32 | 1.62 | 7.17E-07 | 2.24E-05 |
| regulation of metabolic process | GO:0019222 | 3554 | 723 | 447.33 | 1.62 | 3.47E-45 | 1.64E-42 |
| regulation of RNA metabolic process | GO:0051252 | 2445 | 497 | 307.75 | 1.61 | 6.31E-30 | 1.35E-27 |
| positive regulation of biological process | GO:0048518 | 2136 | 434 | 268.85 | 1.61 | 6.06E-26 | 8.79E-24 |
| positive regulation of protein metabolic process | GO:0051247 | 271 | 55 | 34.11 | 1.61 | 3.02E-04 | 4.65E-03 |
| regulation of protein phosphorylation | GO:0001932 | 207 | 42 | 26.05 | 1.61 | 1.52E-03 | 1.85E-02 |
| regulation of DNA-templated transcription | GO:0006355 | 2249 | 455 | 283.08 | 1.61 | 7.90E-27 | 1.32E-24 |
| organophosphate biosynthetic process | GO:0090407 | 351 | 71 | 44.18 | 1.61 | 4.55E-05 | 9.22E-04 |
| regulation of RNA biosynthetic process | GO:2001141 | 2251 | 455 | 283.33 | 1.61 | 1.16E-26 | 1.87E-24 |
| dephosphorylation | GO:0016311 | 218 | 44 | 27.44 | 1.60 | 1.38E-03 | 1.69E-02 |
| regulation of gene expression | GO:0010468 | 2792 | 562 | 351.42 | 1.60 | 6.07E-33 | 1.44E-30 |
| positive regulation of catalytic activity | GO:0043085 | 313 | 63 | 39.40 | 1.60 | 1.46E-04 | 2.49E-03 |
| regulation of biosynthetic process | GO:0009889 | 2882 | 580 | 362.75 | 1.60 | 5.49E-34 | 1.50E-31 |
| regulation of nucleobase-containing compound metabolic process | GO:0019219 | 2545 | 512 | 320.33 | 1.60 | 1.08E-29 | 2.26E-27 |
| response to lipid | GO:0033993 | 184 | 37 | 23.16 | 1.60 | 3.53E-03 | 3.68E-02 |
| glycerophospholipid metabolic process | GO:0006650 | 199 | 40 | 25.05 | 1.60 | 2.50E-03 | 2.80E-02 |
| regulation of macromolecule biosynthetic process | GO:0010556 | 2832 | 569 | 356.46 | 1.60 | 3.57E-33 | 8.68E-31 |
| negative regulation of nucleobase-containing compound metabolic process | GO:0045934 | 448 | 90 | 56.39 | 1.60 | 7.16E-06 | 1.77E-04 |
| carbohydrate metabolic process | GO:0005975 | 289 | 58 | 36.38 | 1.59 | 3.24E-04 | 4.90E-03 |
| regulation of cellular biosynthetic process | GO:0031326 | 2861 | 574 | 360.11 | 1.59 | 3.14E-33 | 7.85E-31 |
| intracellular signaling cassette | GO:0141124 | 369 | 74 | 46.45 | 1.59 | 4.86E-05 | 9.80E-04 |
| monoatomic cation transport | GO:0006812 | 559 | 112 | 70.36 | 1.59 | 5.71E-07 | 1.80E-05 |
| sulfur compound metabolic process | GO:0006790 | 200 | 40 | 25.17 | 1.59 | 2.62E-03 | 2.89E-02 |
| regulation of molecular function | GO:0065009 | 665 | 133 | 83.70 | 1.59 | 4.22E-08 | 1.54E-06 |
| RNA biosynthetic process | GO:0032774 | 195 | 39 | 24.54 | 1.59 | 3.21E-03 | 3.39E-02 |
| negative regulation of macromolecule biosynthetic process | GO:0010558 | 605 | 121 | 76.15 | 1.59 | 2.15E-07 | 7.13E-06 |

| | | | | | | | |
|---|---|---|---|---|---|---|---|
| regulation of primary metabolic process | GO:0080090 | 3151 | 630 | 396.61 | 1.59 | 2.60E-36 | 8.07E-34 |
| regulation of macromolecule metabolic process | GO:0060255 | 3292 | 658 | 414.36 | 1.59 | 3.81E-38 | 1.49E-35 |
| negative regulation of biosynthetic process | GO:0009890 | 612 | 122 | 77.03 | 1.58 | 1.88E-07 | 6.30E-06 |
| regulation of response to stress | GO:0080134 | 276 | 55 | 34.74 | 1.58 | 4.89E-04 | 7.05E-03 |
| monoatomic cation transmembrane transport | GO:0098655 | 457 | 91 | 57.52 | 1.58 | 8.84E-06 | 2.10E-04 |
| intracellular chemical homeostasis | GO:0055082 | 221 | 44 | 27.82 | 1.58 | 2.07E-03 | 2.37E-02 |
| muscle cell development | GO:0055001 | 216 | 43 | 27.19 | 1.58 | 2.60E-03 | 2.88E-02 |
| negative regulation of cellular biosynthetic process | GO:0031327 | 608 | 121 | 76.53 | 1.58 | 2.38E-07 | 7.81E-06 |
| positive regulation of molecular function | GO:0044093 | 367 | 73 | 46.19 | 1.58 | 6.63E-05 | 1.29E-03 |
| plasma membrane bounded cell projection assembly | GO:0120031 | 327 | 65 | 41.16 | 1.58 | 2.00E-04 | 3.24E-03 |
| regulation of nitrogen compound metabolic process | GO:0051171 | 3102 | 616 | 390.44 | 1.58 | 1.53E-34 | 4.29E-32 |
| inorganic cation transmembrane transport | GO:0098662 | 454 | 90 | 57.14 | 1.57 | 1.16E-05 | 2.67E-04 |
| transmembrane transport | GO:0055085 | 1014 | 201 | 127.63 | 1.57 | 2.81E-11 | 1.46E-09 |
| cytoskeleton organization | GO:0007010 | 853 | 169 | 107.37 | 1.57 | 1.25E-09 | 5.32E-08 |
| small molecule catabolic process | GO:0044282 | 202 | 40 | 25.43 | 1.57 | 3.75E-03 | 3.85E-02 |
| negative regulation of cellular process | GO:0048523 | 1824 | 361 | 229.58 | 1.57 | 1.73E-19 | 1.53E-17 |
| inorganic ion homeostasis | GO:0098771 | 213 | 42 | 26.81 | 1.57 | 3.42E-03 | 3.58E-02 |
| phospholipid metabolic process | GO:0006644 | 264 | 52 | 33.23 | 1.56 | 1.01E-03 | 1.32E-02 |
| cellular component organization | GO:0016043 | 3514 | 690 | 442.30 | 1.56 | 1.62E-37 | 6.09E-35 |
| cell projection assembly | GO:0030031 | 347 | 68 | 43.68 | 1.56 | 2.22E-04 | 3.57E-03 |
| endomembrane system organization | GO:0010256 | 245 | 48 | 30.84 | 1.56 | 1.82E-03 | 2.16E-02 |
| negative regulation of biological process | GO:0048519 | 1957 | 383 | 246.32 | 1.55 | 7.14E-20 | 6.56E-18 |
| cellular response to oxygen-containing compound | GO:1901701 | 241 | 47 | 30.33 | 1.55 | 2.32E-03 | 2.63E-02 |
| regulation of protein modification process | GO:0031399 | 267 | 52 | 33.61 | 1.55 | 1.50E-03 | 1.82E-02 |
| cilium assembly | GO:0060271 | 303 | 59 | 38.14 | 1.55 | 6.28E-04 | 8.67E-03 |
| cellular component organization or biogenesis | GO:0071840 | 3697 | 718 | 465.33 | 1.54 | 1.73E-37 | 6.23E-35 |
| response to nitrogen compound | GO:1901698 | 228 | 44 | 28.70 | 1.53 | 3.50E-03 | 3.65E-02 |
| reproductive process | GO:0022414 | 301 | 58 | 37.89 | 1.53 | 8.50E-04 | 1.12E-02 |
| positive regulation of cell communication | GO:0010647 | 479 | 92 | 60.29 | 1.53 | 3.67E-05 | 7.64E-04 |
| positive regulation of signaling | GO:0023056 | 479 | 92 | 60.29 | 1.53 | 3.67E-05 | 7.62E-04 |
| transport | GO:0006810 | 2656 | 510 | 334.31 | 1.53 | 1.24E-24 | 1.57E-22 |
| response to abiotic stimulus | GO:0009628 | 360 | 69 | 45.31 | 1.52 | 3.96E-04 | 5.85E-03 |
| RNA metabolic process | GO:0016070 | 935 | 179 | 117.69 | 1.52 | 6.43E-09 | 2.54E-07 |
| reproduction | GO:0000003 | 304 | 58 | 38.26 | 1.52 | 1.21E-03 | 1.52E-02 |
| establishment of localization | GO:0051234 | 2783 | 530 | 350.29 | 1.51 | 8.96E-25 | 1.15E-22 |
| establishment of localization in cell | GO:0051649 | 940 | 179 | 118.32 | 1.51 | 9.55E-09 | 3.68E-07 |
| regulation of transport | GO:0051049 | 484 | 92 | 60.92 | 1.51 | 5.46E-05 | 1.09E-03 |
| localization | GO:0051179 | 3034 | 576 | 381.88 | 1.51 | 8.74E-27 | 1.43E-24 |
| regulation of intracellular signal transduction | GO:1902531 | 569 | 108 | 71.62 | 1.51 | 1.24E-05 | 2.86E-04 |
| regulation of localization | GO:0032879 | 570 | 108 | 71.74 | 1.51 | 1.27E-05 | 2.90E-04 |

| | | | | | | | |
|---|---|---|---|---|---|---|---|
| vesicle-mediated transport | GO:0016192 | 840 | 159 | 105.73 | 1.50 | 1.07E-07 | 3.72E-06 |
| monoatomic ion homeostasis | GO:0050801 | 254 | 48 | 31.97 | 1.50 | 4.12E-03 | 4.19E-02 |
| supramolecular fiber organization | GO:0097435 | 457 | 86 | 57.52 | 1.50 | 1.16E-04 | 2.04E-03 |
| membrane organization | GO:0061024 | 357 | 67 | 44.93 | 1.49 | 7.14E-04 | 9.70E-03 |
| phosphorus metabolic process | GO:0006793 | 1654 | 310 | 208.19 | 1.49 | 1.49E-13 | 9.71E-12 |
| nucleobase-containing compound biosynthetic process | GO:0034654 | 427 | 80 | 53.75 | 1.49 | 2.90E-04 | 4.48E-03 |
| phosphate-containing compound metabolic process | GO:0006796 | 1639 | 307 | 206.30 | 1.49 | 2.11E-13 | 1.36E-11 |
| organophosphate metabolic process | GO:0019637 | 642 | 120 | 80.81 | 1.49 | 7.56E-06 | 1.85E-04 |
| cell communication | GO:0007154 | 3403 | 636 | 428.33 | 1.48 | 7.00E-28 | 1.26E-25 |
| regulation of organelle organization | GO:0033043 | 445 | 83 | 56.01 | 1.48 | 2.22E-04 | 3.57E-03 |
| cellular localization | GO:0051641 | 1433 | 267 | 180.37 | 1.48 | 1.95E-11 | 1.03E-09 |
| cilium organization | GO:0044782 | 323 | 60 | 40.66 | 1.48 | 1.75E-03 | 2.11E-02 |
| response to organic substance | GO:0010033 | 727 | 135 | 91.51 | 1.48 | 2.99E-06 | 8.24E-05 |
| positive regulation of signal transduction | GO:0009967 | 438 | 81 | 55.13 | 1.47 | 3.51E-04 | 5.26E-03 |
| carbohydrate derivative metabolic process | GO:1901135 | 725 | 134 | 91.25 | 1.47 | 3.91E-06 | 1.04E-04 |
| signaling | GO:0023052 | 3333 | 614 | 419.52 | 1.46 | 3.68E-25 | 4.94E-23 |
| regulation of biological process | GO:0050789 | 8353 | 1538 | 1051.38 | 1.46 | 8.69E-80 | 7.82E-77 |
| heterocycle biosynthetic process | GO:0018130 | 489 | 90 | 61.55 | 1.46 | 1.92E-04 | 3.14E-03 |
| organic cyclic compound biosynthetic process | GO:1901362 | 560 | 103 | 70.49 | 1.46 | 7.91E-05 | 1.50E-03 |
| aromatic compound biosynthetic process | GO:0019438 | 484 | 89 | 60.92 | 1.46 | 2.31E-04 | 3.67E-03 |
| regulation of protein metabolic process | GO:0051246 | 593 | 109 | 74.64 | 1.46 | 4.36E-05 | 8.89E-04 |
| carbohydrate derivative biosynthetic process | GO:1901137 | 468 | 86 | 58.91 | 1.46 | 3.14E-04 | 4.77E-03 |
| biological regulation | GO:0065007 | 8661 | 1591 | 1090.14 | 1.46 | 3.76E-83 | 4.24E-80 |
| cellular response to organic substance | GO:0071310 | 490 | 90 | 61.68 | 1.46 | 1.97E-04 | 3.21E-03 |
| regulation of cellular process | GO:0050794 | 7831 | 1436 | 985.67 | 1.46 | 1.18E-70 | 9.64E-68 |
| cellular catabolic process | GO:0044248 | 671 | 123 | 84.46 | 1.46 | 1.55E-05 | 3.50E-04 |
| nucleic acid metabolic process | GO:0090304 | 1353 | 248 | 170.30 | 1.46 | 5.22E-10 | 2.32E-08 |
| system process | GO:0003008 | 836 | 153 | 105.23 | 1.45 | 1.66E-06 | 4.79E-05 |
| negative regulation of macromolecule metabolic process | GO:0010605 | 800 | 146 | 100.69 | 1.45 | 2.99E-06 | 8.25E-05 |
| nucleotide metabolic process | GO:0009117 | 318 | 58 | 40.03 | 1.45 | 3.67E-03 | 3.77E-02 |
| microtubule-based process | GO:0007017 | 495 | 90 | 62.30 | 1.44 | 3.52E-04 | 5.27E-03 |
| phosphorylation | GO:0016310 | 960 | 174 | 120.83 | 1.44 | 5.07E-07 | 1.61E-05 |
| cellular lipid metabolic process | GO:0044255 | 624 | 113 | 78.54 | 1.44 | 6.68E-05 | 1.29E-03 |
| negative regulation of nitrogen compound metabolic process | GO:0051172 | 641 | 116 | 80.68 | 1.44 | 5.03E-05 | 1.01E-03 |
| negative regulation of cellular metabolic process | GO:0031324 | 763 | 138 | 96.04 | 1.44 | 1.11E-05 | 2.59E-04 |
| organic substance transport | GO:0071702 | 1225 | 221 | 154.19 | 1.43 | 1.90E-08 | 7.05E-07 |
| protein modification process | GO:0036211 | 1569 | 283 | 197.49 | 1.43 | 1.72E-10 | 8.13E-09 |
| nucleobase-containing compound metabolic process | GO:0006139 | 1742 | 314 | 219.26 | 1.43 | 1.45E-11 | 7.80E-10 |
| lipid metabolic process | GO:0006629 | 794 | 143 | 99.94 | 1.43 | 7.88E-06 | 1.92E-04 |
| organelle organization | GO:0006996 | 1935 | 348 | 243.55 | 1.43 | 1.45E-12 | 8.61E-11 |

| | | | | | | | |
|---|---|---|---|---|---|---|---|
| macromolecule modification | GO:0043412 | 1706 | 306 | 214.73 | 1.43 | 5.54E-11 | 2.80E-09 |
| organic cyclic compound metabolic process | GO:1901360 | 1986 | 356 | 249.97 | 1.42 | 1.25E-12 | 7.48E-11 |
| signal transduction | GO:0007165 | 3086 | 553 | 388.43 | 1.42 | 1.09E-19 | 9.76E-18 |
| organonitrogen compound catabolic process | GO:1901565 | 776 | 139 | 97.67 | 1.42 | 1.69E-05 | 3.76E-04 |
| negative regulation of metabolic process | GO:0009892 | 832 | 149 | 104.72 | 1.42 | 7.53E-06 | 1.85E-04 |
| intracellular transport | GO:0046907 | 771 | 138 | 97.04 | 1.42 | 1.62E-05 | 3.63E-04 |
| cellular aromatic compound metabolic process | GO:0006725 | 1860 | 332 | 234.11 | 1.42 | 1.29E-11 | 6.99E-10 |
| macromolecule catabolic process | GO:0009057 | 623 | 111 | 78.42 | 1.42 | 1.43E-04 | 2.45E-03 |
| protein modification by small protein conjugation or removal | GO:0070647 | 584 | 104 | 73.51 | 1.41 | 2.41E-04 | 3.78E-03 |
| heterocycle metabolic process | GO:0046483 | 1848 | 329 | 232.60 | 1.41 | 2.37E-11 | 1.24E-09 |
| post-translational protein modification | GO:0043687 | 602 | 107 | 75.77 | 1.41 | 2.32E-04 | 3.68E-03 |
| response to external stimulus | GO:0009605 | 772 | 137 | 97.17 | 1.41 | 2.70E-05 | 5.78E-04 |
| RNA processing | GO:0006396 | 637 | 113 | 80.18 | 1.41 | 1.68E-04 | 2.81E-03 |
| organic substance biosynthetic process | GO:1901576 | 2515 | 446 | 316.56 | 1.41 | 4.73E-15 | 3.33E-13 |
| cellular process | GO:0009987 | 12722 | 2251 | 1601.29 | 1.41 | 1.03E-130 | 3.10E-127 |
| organic substance catabolic process | GO:1901575 | 1080 | 191 | 135.94 | 1.41 | 8.08E-07 | 2.47E-05 |
| biosynthetic process | GO:0009058 | 2552 | 451 | 321.22 | 1.40 | 6.93E-15 | 4.83E-13 |
| cellular response to stimulus | GO:0051716 | 3832 | 677 | 482.33 | 1.40 | 7.56E-23 | 9.08E-21 |
| carboxylic acid metabolic process | GO:0019752 | 493 | 87 | 62.05 | 1.40 | 1.22E-03 | 1.53E-02 |
| protein localization | GO:0008104 | 981 | 173 | 123.48 | 1.40 | 3.70E-06 | 9.97E-05 |
| response to stimulus | GO:0050896 | 4693 | 827 | 590.70 | 1.40 | 2.39E-28 | 4.39E-26 |
| cellular macromolecule localization | GO:0070727 | 983 | 173 | 123.73 | 1.40 | 3.83E-06 | 1.02E-04 |
| intracellular signal transduction | GO:0035556 | 927 | 163 | 116.68 | 1.40 | 8.53E-06 | 2.04E-04 |
| macromolecule localization | GO:0033036 | 1271 | 223 | 159.98 | 1.39 | 1.80E-07 | 6.05E-06 |
| cellular response to stress | GO:0033554 | 667 | 117 | 83.95 | 1.39 | 1.86E-04 | 3.06E-03 |
| cellular biosynthetic process | GO:0044249 | 2429 | 426 | 305.73 | 1.39 | 1.60E-13 | 1.04E-11 |
| cellular metabolic process | GO:0044237 | 4901 | 859 | 616.88 | 1.39 | 7.74E-29 | 1.48E-26 |
| oxoacid metabolic process | GO:0043436 | 497 | 87 | 62.56 | 1.39 | 1.30E-03 | 1.61E-02 |
| protein phosphorylation | GO:0006468 | 423 | 74 | 53.24 | 1.39 | 3.07E-03 | 3.27E-02 |
| organonitrogen compound biosynthetic process | GO:1901566 | 1056 | 184 | 132.92 | 1.38 | 4.06E-06 | 1.07E-04 |
| positive regulation of response to stimulus | GO:0048584 | 598 | 104 | 75.27 | 1.38 | 5.84E-04 | 8.10E-03 |
| cellular nitrogen compound metabolic process | GO:0034641 | 2151 | 374 | 270.74 | 1.38 | 2.03E-11 | 1.07E-09 |
| primary metabolic process | GO:0044238 | 5319 | 922 | 669.49 | 1.38 | 1.49E-29 | 3.04E-27 |
| protein modification by small protein conjugation | GO:0032446 | 508 | 88 | 63.94 | 1.38 | 1.83E-03 | 2.17E-02 |
| metabolic process | GO:0008152 | 6420 | 1112 | 808.07 | 1.38 | 2.97E-37 | 1.03E-34 |
| catabolic process | GO:0009056 | 1275 | 220 | 160.48 | 1.37 | 7.57E-07 | 2.34E-05 |
| protein transport | GO:0015031 | 644 | 111 | 81.06 | 1.37 | 5.84E-04 | 8.11E-03 |
| nitrogen compound metabolic process | GO:0006807 | 4830 | 831 | 607.94 | 1.37 | 4.91E-25 | 6.40E-23 |
| macromolecule biosynthetic process | GO:0009059 | 1815 | 312 | 228.45 | 1.37 | 4.02E-09 | 1.64E-07 |
| nitrogen compound transport | GO:0071705 | 915 | 157 | 115.17 | 1.36 | 4.72E-05 | 9.54E-04 |

| | | | | | | | |
|---|---|---|---|---|---|---|---|
| organic substance metabolic process | GO:0071704 | 5633 | 965 | 709.01 | 1.36 | 2.85E-29 | 5.71E-27 |
| protein ubiquitination | GO:0016567 | 467 | 80 | 58.78 | 1.36 | 3.85E-03 | 3.93E-02 |
| macromolecule metabolic process | GO:0043170 | 4105 | 702 | 516.69 | 1.36 | 5.87E-20 | 5.44E-18 |
| response to stress | GO:0006950 | 1295 | 221 | 163.00 | 1.36 | 1.74E-06 | 4.99E-05 |
| proteolysis involved in protein catabolic process | GO:0051603 | 475 | 81 | 59.79 | 1.35 | 4.15E-03 | 4.22E-02 |
| small molecule metabolic process | GO:0044281 | 1052 | 179 | 132.41 | 1.35 | 2.30E-05 | 5.00E-04 |
| proteolysis | GO:0006508 | 937 | 159 | 117.94 | 1.35 | 7.20E-05 | 1.39E-03 |
| cellular response to chemical stimulus | GO:0070887 | 755 | 128 | 95.03 | 1.35 | 4.37E-04 | 6.39E-03 |
| biological_process | GO:0008150 | 17206 | 2915 | 2165.69 | 1.35 | 1.30E-217 | 1.17E-213 |
| protein catabolic process | GO:0030163 | 496 | 84 | 62.43 | 1.35 | 4.95E-03 | 4.84E-02 |
| response to chemical | GO:0042221 | 1184 | 200 | 149.03 | 1.34 | 1.07E-05 | 2.49E-04 |
| cellular component assembly | GO:0022607 | 1362 | 230 | 171.43 | 1.34 | 2.47E-06 | 6.93E-05 |
| organelle assembly | GO:0070925 | 594 | 100 | 74.77 | 1.34 | 2.60E-03 | 2.88E-02 |
| establishment of protein localization | GO:0045184 | 733 | 123 | 92.26 | 1.33 | 8.38E-04 | 1.11E-02 |
| organonitrogen compound metabolic process | GO:1901564 | 3602 | 604 | 453.38 | 1.33 | 3.81E-15 | 2.70E-13 |
| gene expression | GO:0010467 | 1439 | 241 | 181.12 | 1.33 | 2.46E-06 | 6.93E-05 |
| cellular component biogenesis | GO:0044085 | 1550 | 259 | 195.10 | 1.33 | 1.15E-06 | 3.44E-05 |
| protein metabolic process | GO:0019538 | 2806 | 464 | 353.19 | 1.31 | 1.08E-10 | 5.23E-09 |
| cellular nitrogen compound biosynthetic process | GO:0044271 | 879 | 145 | 110.64 | 1.31 | 6.24E-04 | 8.64E-03 |

**Supplementary Table 5. GO enrichment of GREAT-associated genes from chromatin regions gaining accessibility in z3′-DpE-/- acinar cells (adult ATAC-seq, Up regions).**

Source: ATAC-seq of FACS-purified fixed mCherry+ adult acinar cells from *Tg(ela:mCherry)* zebrafish (*Danio rerio*), comparing z3′-DpE+/+ and z3′-DpE-/-. Criteria: Differential accessibility (z3′-DpE-/- vs z3′-DpE+/+) defined as FDR ≤ 0.05 and |log2FC| ≥ 1 (edgeR); Up regions defined as peaks with increased accessibility in z3′-DpE-/- vs z3′-DpE+/+ (log2FC > 1). Peaks were assigned to putative target genes using GREAT (peak-to-gene association). GO Biological Process enrichment performed using the PANTHER Overrepresentation Test (*Danio rerio* reference list; Fisher's exact test with FDR correction).

| GO term (biological process) | GO ID | Ref list count (N=26353) | Query list count (n=2895) | Expected | Fold enrichment | Raw P-value | FDR |
|---|---|---|---|---|---|---|---|
| negative regulation of reactive oxygen species biosynthetic process | GO:1903427 | 3 | 3 | .33 | 9.10 | 1.32E-03 | 2.25E-02 |
| late distal convoluted tubule development | GO:0072068 | 3 | 3 | .33 | 9.10 | 1.32E-03 | 2.25E-02 |
| distal convoluted tubule development | GO:0072025 | 3 | 3 | .33 | 9.10 | 1.32E-03 | 2.25E-02 |
| octopamine biosynthetic process | GO:0006589 | 3 | 3 | .33 | 9.10 | 1.32E-03 | 2.24E-02 |
| motor behavior | GO:0061744 | 4 | 4 | .44 | 9.10 | 1.45E-04 | 3.48E-03 |
| pore complex assembly | GO:0046931 | 3 | 3 | .33 | 9.10 | 1.32E-03 | 2.24E-02 |
| clathrin-dependent synaptic vesicle endocytosis | GO:0150007 | 3 | 3 | .33 | 9.10 | 1.32E-03 | 2.23E-02 |
| norepinephrine biosynthetic process | GO:0042421 | 3 | 3 | .33 | 9.10 | 1.32E-03 | 2.23E-02 |
| dopamine catabolic process | GO:0042420 | 3 | 3 | .33 | 9.10 | 1.32E-03 | 2.22E-02 |
| positive regulation of mitochondrial calcium ion concentration | GO:0051561 | 3 | 3 | .33 | 9.10 | 1.32E-03 | 2.22E-02 |
| octopamine metabolic process | GO:0046333 | 3 | 3 | .33 | 9.10 | 1.32E-03 | 2.22E-02 |
| nuclear pore complex assembly | GO:0051292 | 3 | 3 | .33 | 9.10 | 1.32E-03 | 2.21E-02 |
| response to selenium ion | GO:0010269 | 3 | 3 | .33 | 9.10 | 1.32E-03 | 2.21E-02 |
| regulation of motor neuron axon guidance | GO:1905812 | 3 | 3 | .33 | 9.10 | 1.32E-03 | 2.20E-02 |
| negative regulation of macrophage differentiation | GO:0045650 | 3 | 3 | .33 | 9.10 | 1.32E-03 | 2.20E-02 |
| regulation of reactive oxygen species biosynthetic process | GO:1903426 | 5 | 4 | .55 | 7.28 | 6.63E-04 | 1.26E-02 |
| macrophage activation involved in immune response | GO:0002281 | 5 | 4 | .55 | 7.28 | 6.63E-04 | 1.26E-02 |
| skeletal muscle satellite cell migration | GO:1902766 | 5 | 4 | .55 | 7.28 | 6.63E-04 | 1.26E-02 |
| regulation of Wnt signaling pathway involved in dorsal/ventral axis specification | GO:2000053 | 5 | 4 | .55 | 7.28 | 6.63E-04 | 1.26E-02 |
| neuron cell-cell adhesion | GO:0007158 | 9 | 7 | .99 | 7.08 | 5.65E-06 | 1.96E-04 |
| negative regulation of myeloid leukocyte differentiation | GO:0002762 | 8 | 6 | .88 | 6.83 | 4.02E-05 | 1.15E-03 |
| epiboly involved in wound healing | GO:0090505 | 7 | 5 | .77 | 6.50 | 2.77E-04 | 6.10E-03 |
| wound healing, spreading of cells | GO:0044319 | 7 | 5 | .77 | 6.50 | 2.77E-04 | 6.09E-03 |
| presynaptic membrane assembly | GO:0097105 | 7 | 5 | .77 | 6.50 | 2.77E-04 | 6.07E-03 |

| | | | | | | | |
|---|---|---|---|---|---|---|---|
| postsynaptic membrane assembly | GO:0097104 | 7 | 5 | .77 | 6.50 | 2.77E-04 | 6.06E-03 |
| presynaptic membrane organization | GO:0097090 | 7 | 5 | .77 | 6.50 | 2.77E-04 | 6.04E-03 |
| postsynapse assembly | GO:0099068 | 7 | 5 | .77 | 6.50 | 2.77E-04 | 6.03E-03 |
| phenol-containing compound catabolic process | GO:0019336 | 7 | 5 | .77 | 6.50 | 2.77E-04 | 6.01E-03 |
| L-proline biosynthetic process | GO:0055129 | 6 | 4 | .66 | 6.07 | 1.82E-03 | 2.83E-02 |
| proline biosynthetic process | GO:0006561 | 6 | 4 | .66 | 6.07 | 1.82E-03 | 2.83E-02 |
| sphingolipid mediated signaling pathway | GO:0090520 | 6 | 4 | .66 | 6.07 | 1.82E-03 | 2.82E-02 |
| catecholamine catabolic process | GO:0042424 | 6 | 4 | .66 | 6.07 | 1.82E-03 | 2.82E-02 |
| positive regulation of cell morphogenesis | GO:0010770 | 6 | 4 | .66 | 6.07 | 1.82E-03 | 2.81E-02 |
| swim bladder inflation | GO:0048798 | 6 | 4 | .66 | 6.07 | 1.82E-03 | 2.81E-02 |
| swim bladder maturation | GO:0048796 | 6 | 4 | .66 | 6.07 | 1.82E-03 | 2.80E-02 |
| regulation of retinoic acid biosynthetic process | GO:1900052 | 6 | 4 | .66 | 6.07 | 1.82E-03 | 2.80E-02 |
| regulation of isoprenoid metabolic process | GO:0019747 | 6 | 4 | .66 | 6.07 | 1.82E-03 | 2.79E-02 |
| catechol-containing compound catabolic process | GO:0019614 | 6 | 4 | .66 | 6.07 | 1.82E-03 | 2.79E-02 |
| skin epidermis development | GO:0098773 | 6 | 4 | .66 | 6.07 | 1.82E-03 | 2.78E-02 |
| amine catabolic process | GO:0009310 | 11 | 7 | 1.21 | 5.79 | 4.23E-05 | 1.19E-03 |
| cellular biogenic amine catabolic process | GO:0042402 | 11 | 7 | 1.21 | 5.79 | 4.23E-05 | 1.19E-03 |
| heart rudiment development | GO:0003313 | 11 | 7 | 1.21 | 5.79 | 4.23E-05 | 1.19E-03 |
| negative regulation of leukocyte differentiation | GO:1902106 | 11 | 7 | 1.21 | 5.79 | 4.23E-05 | 1.18E-03 |
| negative regulation of hemopoiesis | GO:1903707 | 11 | 7 | 1.21 | 5.79 | 4.23E-05 | 1.18E-03 |
| head morphogenesis | GO:0060323 | 11 | 7 | 1.21 | 5.79 | 4.23E-05 | 1.18E-03 |
| distal tubule development | GO:0072017 | 8 | 5 | .88 | 5.69 | 6.71E-04 | 1.27E-02 |
| motor neuron migration | GO:0097475 | 8 | 5 | .88 | 5.69 | 6.71E-04 | 1.27E-02 |
| kynurenine metabolic process | GO:0070189 | 8 | 5 | .88 | 5.69 | 6.71E-04 | 1.26E-02 |
| retinoic acid receptor signaling pathway | GO:0048384 | 23 | 14 | 2.53 | 5.54 | 1.13E-08 | 5.93E-07 |
| establishment of tissue polarity | GO:0007164 | 10 | 6 | 1.10 | 5.46 | 2.48E-04 | 5.58E-03 |
| response to hydrogen peroxide | GO:0042542 | 10 | 6 | 1.10 | 5.46 | 2.48E-04 | 5.56E-03 |
| regulation of axon guidance | GO:1902667 | 10 | 6 | 1.10 | 5.46 | 2.48E-04 | 5.55E-03 |
| iron coordination entity transport | GO:1901678 | 10 | 6 | 1.10 | 5.46 | 2.48E-04 | 5.54E-03 |
| negative regulation of myeloid cell differentiation | GO:0045638 | 12 | 7 | 1.32 | 5.31 | 9.19E-05 | 2.31E-03 |
| establishment of planar polarity | GO:0001736 | 9 | 5 | .99 | 5.06 | 1.37E-03 | 2.26E-02 |
| heme transport | GO:0015886 | 9 | 5 | .99 | 5.06 | 1.37E-03 | 2.26E-02 |
| regulation of cell-substrate junction organization | GO:0150116 | 9 | 5 | .99 | 5.06 | 1.37E-03 | 2.25E-02 |
| regulation of focal adhesion assembly | GO:0051893 | 9 | 5 | .99 | 5.06 | 1.37E-03 | 2.25E-02 |
| regulation of cell-substrate junction assembly | GO:0090109 | 9 | 5 | .99 | 5.06 | 1.37E-03 | 2.25E-02 |
| body morphogenesis | GO:0010171 | 18 | 10 | 1.98 | 5.06 | 4.79E-06 | 1.72E-04 |
| face morphogenesis | GO:0060325 | 9 | 5 | .99 | 5.06 | 1.37E-03 | 2.24E-02 |
| benzene-containing compound metabolic process | GO:0042537 | 11 | 6 | 1.21 | 4.97 | 4.95E-04 | 1.00E-02 |
| face development | GO:0060324 | 11 | 6 | 1.21 | 4.97 | 4.95E-04 | 9.98E-03 |
| postsynaptic membrane organization | GO:0001941 | 15 | 8 | 1.65 | 4.85 | 6.62E-05 | 1.75E-03 |

| | | | | | | | |
|---|---|---|---|---|---|---|---|
| stem cell population maintenance | GO:0019827 | 15 | 8 | 1.65 | 4.85 | 6.62E-05 | 1.74E-03 |
| maintenance of cell number | GO:0098727 | 17 | 9 | 1.87 | 4.82 | 2.45E-05 | 7.48E-04 |
| anterior/posterior axon guidance | GO:0033564 | 12 | 6 | 1.32 | 4.55 | 8.97E-04 | 1.62E-02 |
| regulation of cell-matrix adhesion | GO:0001952 | 10 | 5 | 1.10 | 4.55 | 2.50E-03 | 3.60E-02 |
| proximal tubule development | GO:0072014 | 12 | 6 | 1.32 | 4.55 | 8.97E-04 | 1.62E-02 |
| urogenital system development | GO:0001655 | 14 | 7 | 1.54 | 4.55 | 3.26E-04 | 6.98E-03 |
| regulation of epithelial to mesenchymal transition | GO:0010717 | 14 | 7 | 1.54 | 4.55 | 3.26E-04 | 6.96E-03 |
| presynapse organization | GO:0099172 | 12 | 6 | 1.32 | 4.55 | 8.97E-04 | 1.61E-02 |
| presynapse assembly | GO:0099054 | 12 | 6 | 1.32 | 4.55 | 8.97E-04 | 1.61E-02 |
| pronephric nephron tubule development | GO:0039020 | 17 | 8 | 1.87 | 4.28 | 2.04E-04 | 4.74E-03 |
| dorsal aorta development | GO:0035907 | 15 | 7 | 1.65 | 4.25 | 5.54E-04 | 1.09E-02 |
| regulation of T cell differentiation | GO:0045580 | 13 | 6 | 1.43 | 4.20 | 1.51E-03 | 2.41E-02 |
| regulation of nodal signaling pathway | GO:1900107 | 13 | 6 | 1.43 | 4.20 | 1.51E-03 | 2.41E-02 |
| mesendoderm development | GO:0048382 | 13 | 6 | 1.43 | 4.20 | 1.51E-03 | 2.40E-02 |
| regulation of glial cell differentiation | GO:0045685 | 13 | 6 | 1.43 | 4.20 | 1.51E-03 | 2.40E-02 |
| dorsal/ventral axis specification | GO:0009950 | 22 | 10 | 2.42 | 4.14 | 4.67E-05 | 1.28E-03 |
| activin receptor signaling pathway | GO:0032924 | 18 | 8 | 1.98 | 4.05 | 3.32E-04 | 7.07E-03 |
| tRNA wobble base modification | GO:0002097 | 18 | 8 | 1.98 | 4.05 | 3.32E-04 | 7.06E-03 |
| animal organ regeneration | GO:0031100 | 25 | 11 | 2.75 | 4.01 | 2.83E-05 | 8.45E-04 |
| tRNA wobble uridine modification | GO:0002098 | 16 | 7 | 1.76 | 3.98 | 8.91E-04 | 1.61E-02 |
| formation of anatomical boundary | GO:0048859 | 16 | 7 | 1.76 | 3.98 | 8.91E-04 | 1.61E-02 |
| rhombomere formation | GO:0021594 | 14 | 6 | 1.54 | 3.90 | 2.40E-03 | 3.49E-02 |
| phosphate ion transport | GO:0006817 | 21 | 9 | 2.31 | 3.90 | 1.97E-04 | 4.58E-03 |
| rhombomere morphogenesis | GO:0021593 | 19 | 8 | 2.09 | 3.83 | 5.19E-04 | 1.05E-02 |
| regulation of lymphocyte differentiation | GO:0045619 | 19 | 8 | 2.09 | 3.83 | 5.19E-04 | 1.04E-02 |
| hematopoietic or lymphoid organ development | GO:0048534 | 24 | 10 | 2.64 | 3.79 | 1.15E-04 | 2.81E-03 |
| postsynapse organization | GO:0099173 | 24 | 10 | 2.64 | 3.79 | 1.15E-04 | 2.80E-03 |
| regulation of activin receptor signaling pathway | GO:0032925 | 17 | 7 | 1.87 | 3.75 | 1.37E-03 | 2.24E-02 |
| synaptic vesicle recycling | GO:0036465 | 44 | 18 | 4.83 | 3.72 | 3.15E-07 | 1.38E-05 |
| presynaptic endocytosis | GO:0140238 | 37 | 15 | 4.06 | 3.69 | 3.48E-06 | 1.29E-04 |
| synaptic vesicle endocytosis | GO:0048488 | 37 | 15 | 4.06 | 3.69 | 3.48E-06 | 1.28E-04 |
| intracellular receptor signaling pathway | GO:0030522 | 42 | 17 | 4.61 | 3.68 | 8.02E-07 | 3.33E-05 |
| nephron tubule development | GO:0072080 | 25 | 10 | 2.75 | 3.64 | 1.73E-04 | 4.08E-03 |
| immune system development | GO:0002520 | 35 | 14 | 3.84 | 3.64 | 8.87E-06 | 2.99E-04 |
| cell volume homeostasis | GO:0006884 | 15 | 6 | 1.65 | 3.64 | 3.63E-03 | 4.92E-02 |
| monoatomic anion homeostasis | GO:0055081 | 20 | 8 | 2.20 | 3.64 | 7.82E-04 | 1.44E-02 |
| artery morphogenesis | GO:0048844 | 20 | 8 | 2.20 | 3.64 | 7.82E-04 | 1.44E-02 |
| pancreas regeneration | GO:1990798 | 20 | 8 | 2.20 | 3.64 | 7.82E-04 | 1.44E-02 |
| thymus development | GO:0048538 | 20 | 8 | 2.20 | 3.64 | 7.82E-04 | 1.43E-02 |
| positive regulation of axon extension | GO:0045773 | 15 | 6 | 1.65 | 3.64 | 3.63E-03 | 4.92E-02 |
| midbrain development | GO:0030901 | 38 | 15 | 4.17 | 3.59 | 5.17E-06 | 1.84E-04 |

| | | | | | | | |
|---|---|---|---|---|---|---|---|
| cellular response to oxygen levels | GO:0071453 | 18 | 7 | 1.98 | 3.54 | 2.03E-03 | 3.06E-02 |
| mitochondrion localization | GO:0051646 | 18 | 7 | 1.98 | 3.54 | 2.03E-03 | 3.06E-02 |
| aorta development | GO:0035904 | 18 | 7 | 1.98 | 3.54 | 2.03E-03 | 3.05E-02 |
| regulation of leukocyte differentiation | GO:1902105 | 44 | 17 | 4.83 | 3.52 | 1.74E-06 | 6.79E-05 |
| renal tubule development | GO:0061326 | 26 | 10 | 2.86 | 3.50 | 2.54E-04 | 5.65E-03 |
| regulation of myeloid leukocyte differentiation | GO:0002761 | 26 | 10 | 2.86 | 3.50 | 2.54E-04 | 5.64E-03 |
| rhombomere development | GO:0021546 | 21 | 8 | 2.31 | 3.47 | 1.14E-03 | 1.99E-02 |
| epithalamus development | GO:0021538 | 21 | 8 | 2.31 | 3.47 | 1.14E-03 | 1.99E-02 |
| endothelial cell proliferation | GO:0001935 | 27 | 10 | 2.97 | 3.37 | 3.63E-04 | 7.57E-03 |
| peroxisomal membrane transport | GO:0015919 | 19 | 7 | 2.09 | 3.35 | 2.92E-03 | 4.12E-02 |
| morphogenesis of a polarized epithelium | GO:0001738 | 19 | 7 | 2.09 | 3.35 | 2.92E-03 | 4.12E-02 |
| regulation of endothelial cell migration | GO:0010594 | 19 | 7 | 2.09 | 3.35 | 2.92E-03 | 4.11E-02 |
| peroxisomal transport | GO:0043574 | 19 | 7 | 2.09 | 3.35 | 2.92E-03 | 4.10E-02 |
| ventral spinal cord interneuron differentiation | GO:0021514 | 25 | 9 | 2.75 | 3.28 | 9.08E-04 | 1.63E-02 |
| neuron fate specification | GO:0048665 | 31 | 11 | 3.41 | 3.23 | 2.87E-04 | 6.23E-03 |
| brain morphogenesis | GO:0048854 | 34 | 12 | 3.74 | 3.21 | 1.62E-04 | 3.85E-03 |
| establishment of spindle orientation | GO:0051294 | 23 | 8 | 2.53 | 3.17 | 2.25E-03 | 3.32E-02 |
| neuron fate commitment | GO:0048663 | 49 | 17 | 5.38 | 3.16 | 9.63E-06 | 3.20E-04 |
| establishment of mitotic spindle localization | GO:0040001 | 26 | 9 | 2.86 | 3.15 | 1.25E-03 | 2.14E-02 |
| positive regulation of neuron projection development | GO:0010976 | 29 | 10 | 3.19 | 3.14 | 7.02E-04 | 1.32E-02 |
| notochord morphogenesis | GO:0048570 | 29 | 10 | 3.19 | 3.14 | 7.02E-04 | 1.31E-02 |
| extracellular transport | GO:0006858 | 35 | 12 | 3.84 | 3.12 | 2.22E-04 | 5.11E-03 |
| epithelial cilium movement involved in extracellular fluid movement | GO:0003351 | 35 | 12 | 3.84 | 3.12 | 2.22E-04 | 5.09E-03 |
| artery development | GO:0060840 | 38 | 13 | 4.17 | 3.11 | 1.26E-04 | 3.03E-03 |
| spindle localization | GO:0051653 | 33 | 11 | 3.63 | 3.03 | 5.34E-04 | 1.06E-02 |
| establishment of spindle localization | GO:0051293 | 33 | 11 | 3.63 | 3.03 | 5.34E-04 | 1.06E-02 |
| response to yeast | GO:0001878 | 24 | 8 | 2.64 | 3.03 | 3.06E-03 | 4.27E-02 |
| inner ear receptor cell differentiation | GO:0060113 | 48 | 16 | 5.27 | 3.03 | 3.10E-05 | 9.09E-04 |
| inner ear auditory receptor cell differentiation | GO:0042491 | 27 | 9 | 2.97 | 3.03 | 1.70E-03 | 2.68E-02 |
| hypothalamus development | GO:0021854 | 27 | 9 | 2.97 | 3.03 | 1.70E-03 | 2.67E-02 |
| limbic system development | GO:0021761 | 28 | 9 | 3.08 | 2.93 | 2.26E-03 | 3.33E-02 |
| tissue migration | GO:0090130 | 53 | 17 | 5.82 | 2.92 | 3.12E-05 | 9.11E-04 |
| inner ear receptor cell development | GO:0060119 | 41 | 13 | 4.50 | 2.89 | 2.98E-04 | 6.44E-03 |
| diencephalon development | GO:0021536 | 86 | 27 | 9.45 | 2.86 | 2.68E-07 | 1.21E-05 |
| semicircular canal development | GO:0060872 | 29 | 9 | 3.19 | 2.83 | 2.97E-03 | 4.16E-02 |
| regulation of mRNA processing | GO:0050684 | 71 | 22 | 7.80 | 2.82 | 4.26E-06 | 1.55E-04 |
| regulation of neuron projection development | GO:0010975 | 165 | 51 | 18.13 | 2.81 | 3.19E-12 | 2.41E-10 |
| regulation of alternative mRNA splicing, via spliceosome | GO:0000381 | 39 | 12 | 4.28 | 2.80 | 6.87E-04 | 1.29E-02 |
| neural crest cell migration | GO:0001755 | 108 | 33 | 11.86 | 2.78 | 2.78E-08 | 1.39E-06 |
| epithelial cell proliferation | GO:0050673 | 36 | 11 | 3.95 | 2.78 | 1.21E-03 | 2.09E-02 |

| | | | | | | | |
|---|---|---|---|---|---|---|---|
| regulation of cell morphogenesis | GO:0022604 | 79 | 24 | 8.68 | 2.77 | 2.35E-06 | 8.91E-05 |
| establishment or maintenance of epithelial cell apical/basal polarity | GO:0045197 | 33 | 10 | 3.63 | 2.76 | 2.15E-03 | 3.20E-02 |
| negative regulation of axon extension | GO:0030517 | 40 | 12 | 4.39 | 2.73 | 8.84E-04 | 1.60E-02 |
| mesenchymal cell migration | GO:0090497 | 110 | 33 | 12.08 | 2.73 | 4.57E-08 | 2.25E-06 |
| regulation of hemopoiesis | GO:1903706 | 67 | 20 | 7.36 | 2.72 | 2.13E-05 | 6.51E-04 |
| ventral spinal cord development | GO:0021517 | 57 | 17 | 6.26 | 2.71 | 8.77E-05 | 2.23E-03 |
| positive regulation of cell development | GO:0010720 | 94 | 28 | 10.33 | 2.71 | 5.47E-07 | 2.32E-05 |
| pronephric nephron development | GO:0039019 | 47 | 14 | 5.16 | 2.71 | 3.66E-04 | 7.58E-03 |
| neural crest cell development | GO:0014032 | 148 | 44 | 16.26 | 2.71 | 3.89E-10 | 2.52E-08 |
| stem cell development | GO:0048864 | 148 | 44 | 16.26 | 2.71 | 3.89E-10 | 2.50E-08 |
| regulation of chemotaxis | GO:0050920 | 74 | 22 | 8.13 | 2.71 | 9.02E-06 | 3.02E-04 |
| regulation of peptidyl-tyrosine phosphorylation | GO:0050730 | 37 | 11 | 4.06 | 2.71 | 1.56E-03 | 2.47E-02 |
| inorganic anion transmembrane transport | GO:0098661 | 74 | 22 | 8.13 | 2.71 | 9.02E-06 | 3.01E-04 |
| neural crest cell differentiation | GO:0014033 | 165 | 49 | 18.13 | 2.70 | 4.19E-11 | 2.88E-09 |
| hormone-mediated signaling pathway | GO:0009755 | 54 | 16 | 5.93 | 2.70 | 1.53E-04 | 3.64E-03 |
| positive regulation of neurogenesis | GO:0050769 | 71 | 21 | 7.80 | 2.69 | 1.56E-05 | 4.93E-04 |
| carbohydrate transport | GO:0008643 | 44 | 13 | 4.83 | 2.69 | 6.42E-04 | 1.23E-02 |
| regulation of axonogenesis | GO:0050770 | 95 | 28 | 10.44 | 2.68 | 6.95E-07 | 2.90E-05 |
| regulation of axon extension | GO:0030516 | 68 | 20 | 7.47 | 2.68 | 2.71E-05 | 8.20E-04 |
| positive regulation of axonogenesis | GO:0050772 | 34 | 10 | 3.74 | 2.68 | 2.75E-03 | 3.93E-02 |
| chloride transmembrane transport | GO:1902476 | 58 | 17 | 6.37 | 2.67 | 1.11E-04 | 2.74E-03 |
| kidney epithelium development | GO:0072073 | 69 | 20 | 7.58 | 2.64 | 3.42E-05 | 9.93E-04 |
| bone morphogenesis | GO:0060349 | 38 | 11 | 4.17 | 2.64 | 1.98E-03 | 2.99E-02 |
| central nervous system neuron differentiation | GO:0021953 | 121 | 35 | 13.29 | 2.63 | 5.03E-08 | 2.46E-06 |
| mesenchymal cell differentiation | GO:0048762 | 184 | 53 | 20.21 | 2.62 | 2.48E-11 | 1.73E-09 |
| neural tube patterning | GO:0021532 | 56 | 16 | 6.15 | 2.60 | 2.44E-04 | 5.52E-03 |
| regulation of cell size | GO:0008361 | 105 | 30 | 11.53 | 2.60 | 5.92E-07 | 2.49E-05 |
| cell differentiation in spinal cord | GO:0021515 | 63 | 18 | 6.92 | 2.60 | 1.02E-04 | 2.55E-03 |
| establishment or maintenance of apical/basal cell polarity | GO:0035088 | 35 | 10 | 3.84 | 2.60 | 3.48E-03 | 4.74E-02 |
| skeletal system morphogenesis | GO:0048705 | 217 | 62 | 23.84 | 2.60 | 7.75E-13 | 6.06E-11 |
| establishment or maintenance of bipolar cell polarity | GO:0061245 | 35 | 10 | 3.84 | 2.60 | 3.48E-03 | 4.73E-02 |
| negative regulation of axonogenesis | GO:0050771 | 42 | 12 | 4.61 | 2.60 | 1.42E-03 | 2.30E-02 |
| synapse assembly | GO:0007416 | 42 | 12 | 4.61 | 2.60 | 1.42E-03 | 2.30E-02 |
| negative regulation of cell development | GO:0010721 | 74 | 21 | 8.13 | 2.58 | 3.12E-05 | 9.09E-04 |
| positive regulation of transcription by RNA polymerase II | GO:0045944 | 335 | 95 | 36.80 | 2.58 | 1.14E-18 | 1.56E-16 |
| synapse organization | GO:0050808 | 127 | 36 | 13.95 | 2.58 | 5.78E-08 | 2.78E-06 |
| mitotic cytokinesis | GO:0000281 | 39 | 11 | 4.28 | 2.57 | 2.49E-03 | 3.60E-02 |
| regulation of myeloid cell differentiation | GO:0045637 | 39 | 11 | 4.28 | 2.57 | 2.49E-03 | 3.59E-02 |
| ear development | GO:0043583 | 188 | 53 | 20.65 | 2.57 | 6.04E-11 | 4.12E-09 |
| nephron epithelium development | GO:0072009 | 57 | 16 | 6.26 | 2.56 | 3.05E-04 | 6.55E-03 |
| regulation of extent of cell growth | GO:0061387 | 82 | 23 | 9.01 | 2.55 | 1.65E-05 | 5.14E-04 |

| | | | | | | | |
|---|---|---|---|---|---|---|---|
| epithelial cell migration | GO:0010631 | 43 | 12 | 4.72 | 2.54 | 1.78E-03 | 2.78E-02 |
| regulation of mRNA splicing, via spliceosome | GO:0048024 | 61 | 17 | 6.70 | 2.54 | 2.20E-04 | 5.06E-03 |
| response to decreased oxygen levels | GO:0036293 | 72 | 20 | 7.91 | 2.53 | 6.67E-05 | 1.75E-03 |
| negative regulation of cell projection organization | GO:0031345 | 54 | 15 | 5.93 | 2.53 | 5.27E-04 | 1.05E-02 |
| negative regulation of canonical Wnt signaling pathway | GO:0090090 | 54 | 15 | 5.93 | 2.53 | 5.27E-04 | 1.05E-02 |
| neural tube development | GO:0021915 | 108 | 30 | 11.86 | 2.53 | 1.14E-06 | 4.67E-05 |
| epithelium migration | GO:0090132 | 47 | 13 | 5.16 | 2.52 | 1.27E-03 | 2.17E-02 |
| inorganic anion transport | GO:0015698 | 120 | 33 | 13.18 | 2.50 | 4.38E-07 | 1.89E-05 |
| embryonic cranial skeleton morphogenesis | GO:0048701 | 171 | 47 | 18.79 | 2.50 | 1.79E-09 | 1.06E-07 |
| negative regulation of neuron projection development | GO:0010977 | 51 | 14 | 5.60 | 2.50 | 9.09E-04 | 1.62E-02 |
| negative regulation of cell growth | GO:0030308 | 51 | 14 | 5.60 | 2.50 | 9.09E-04 | 1.62E-02 |
| establishment of cell polarity | GO:0030010 | 51 | 14 | 5.60 | 2.50 | 9.09E-04 | 1.62E-02 |
| inner ear development | GO:0048839 | 186 | 51 | 20.43 | 2.50 | 4.08E-10 | 2.60E-08 |
| retinal ganglion cell axon guidance | GO:0031290 | 44 | 12 | 4.83 | 2.48 | 2.20E-03 | 3.26E-02 |
| response to oxygen levels | GO:0070482 | 77 | 21 | 8.46 | 2.48 | 5.95E-05 | 1.60E-03 |
| embryonic skeletal system morphogenesis | GO:0048704 | 187 | 51 | 20.54 | 2.48 | 8.27E-10 | 5.07E-08 |
| ameboidal-type cell migration | GO:0001667 | 235 | 64 | 25.82 | 2.48 | 4.67E-12 | 3.51E-10 |
| regulation of cell development | GO:0060284 | 276 | 75 | 30.32 | 2.47 | 6.62E-14 | 5.62E-12 |
| mechanoreceptor differentiation | GO:0042490 | 70 | 19 | 7.69 | 2.47 | 1.41E-04 | 3.38E-03 |
| response to hypoxia | GO:0001666 | 71 | 19 | 7.80 | 2.44 | 1.73E-04 | 4.08E-03 |
| positive regulation of nervous system development | GO:0051962 | 86 | 23 | 9.45 | 2.43 | 3.80E-05 | 1.10E-03 |
| endocrine system development | GO:0035270 | 116 | 31 | 12.74 | 2.43 | 1.89E-06 | 7.33E-05 |
| stem cell differentiation | GO:0048863 | 232 | 62 | 25.49 | 2.43 | 1.99E-11 | 1.41E-09 |
| MAPK cascade | GO:0000165 | 45 | 12 | 4.94 | 2.43 | 2.71E-03 | 3.87E-02 |
| Notch signaling pathway | GO:0007219 | 75 | 20 | 8.24 | 2.43 | 1.24E-04 | 3.01E-03 |
| regulation of lipid metabolic process | GO:0019216 | 60 | 16 | 6.59 | 2.43 | 5.73E-04 | 1.12E-02 |
| regulation of Notch signaling pathway | GO:0008593 | 60 | 16 | 6.59 | 2.43 | 5.73E-04 | 1.12E-02 |
| otolith development | GO:0048840 | 60 | 16 | 6.59 | 2.43 | 5.73E-04 | 1.12E-02 |
| limb development | GO:0060173 | 79 | 21 | 8.68 | 2.42 | 8.94E-05 | 2.27E-03 |
| pronephros development | GO:0048793 | 143 | 38 | 15.71 | 2.42 | 2.18E-07 | 9.92E-06 |
| mesenchyme development | GO:0060485 | 241 | 64 | 26.47 | 2.42 | 1.23E-11 | 8.89E-10 |
| axon extension | GO:0048675 | 49 | 13 | 5.38 | 2.42 | 1.92E-03 | 2.94E-02 |
| enteric nervous system development | GO:0048484 | 49 | 13 | 5.38 | 2.42 | 1.92E-03 | 2.93E-02 |
| embryonic skeletal system development | GO:0048706 | 204 | 54 | 22.41 | 2.41 | 5.66E-10 | 3.54E-08 |
| positive regulation of cell differentiation | GO:0045597 | 178 | 47 | 19.55 | 2.40 | 8.26E-09 | 4.40E-07 |
| axis specification | GO:0009798 | 76 | 20 | 8.35 | 2.40 | 1.51E-04 | 3.61E-03 |
| embryonic eye morphogenesis | GO:0048048 | 57 | 15 | 6.26 | 2.40 | 9.78E-04 | 1.72E-02 |
| sprouting angiogenesis | GO:0002040 | 114 | 30 | 12.52 | 2.40 | 5.55E-06 | 1.95E-04 |
| somitogenesis | GO:0001756 | 103 | 27 | 11.32 | 2.39 | 1.86E-05 | 5.74E-04 |
| embryonic heart tube development | GO:0035050 | 172 | 45 | 18.90 | 2.38 | 2.24E-08 | 1.14E-06 |
| spinal cord development | GO:0021510 | 88 | 23 | 9.67 | 2.38 | 9.13E-05 | 2.30E-03 |

| | | | | | | | |
|---|---|---|---|---|---|---|---|
| regulation of developmental growth | GO:0048638 | 111 | 29 | 12.19 | 2.38 | 8.89E-06 | 2.99E-04 |
| chloride transport | GO:0006821 | 92 | 24 | 10.11 | 2.37 | 6.24E-05 | 1.66E-03 |
| negative regulation of developmental growth | GO:0048640 | 46 | 12 | 5.05 | 2.37 | 3.30E-03 | 4.52E-02 |
| skeletal system development | GO:0001501 | 372 | 97 | 40.87 | 2.37 | 3.57E-16 | 3.65E-14 |
| nephron development | GO:0072006 | 96 | 25 | 10.55 | 2.37 | 4.29E-05 | 1.18E-03 |
| negative regulation of mitotic cell cycle phase transition | GO:1901991 | 50 | 13 | 5.49 | 2.37 | 2.34E-03 | 3.42E-02 |
| motor neuron axon guidance | GO:0008045 | 50 | 13 | 5.49 | 2.37 | 2.34E-03 | 3.42E-02 |
| embryonic viscerocranium morphogenesis | GO:0048703 | 108 | 28 | 11.86 | 2.36 | 1.43E-05 | 4.58E-04 |
| carbohydrate biosynthetic process | GO:0016051 | 58 | 15 | 6.37 | 2.35 | 1.19E-03 | 2.06E-02 |
| angiogenesis | GO:0001525 | 279 | 72 | 30.65 | 2.35 | 3.01E-12 | 2.30E-10 |
| segmentation | GO:0035282 | 124 | 32 | 13.62 | 2.35 | 3.48E-06 | 1.29E-04 |
| embryonic hemopoiesis | GO:0035162 | 93 | 24 | 10.22 | 2.35 | 6.85E-05 | 1.79E-03 |
| Golgi organization | GO:0007030 | 66 | 17 | 7.25 | 2.34 | 6.05E-04 | 1.17E-02 |
| hindbrain development | GO:0030902 | 136 | 35 | 14.94 | 2.34 | 1.24E-06 | 5.03E-05 |
| alcohol biosynthetic process | GO:0046165 | 70 | 18 | 7.69 | 2.34 | 7.19E-04 | 1.34E-02 |
| forebrain development | GO:0030900 | 148 | 38 | 16.26 | 2.34 | 4.51E-07 | 1.93E-05 |
| synaptic vesicle cycle | GO:0099504 | 90 | 23 | 9.89 | 2.33 | 1.10E-04 | 2.71E-03 |
| vesicle-mediated transport in synapse | GO:0099003 | 90 | 23 | 9.89 | 2.33 | 1.10E-04 | 2.71E-03 |
| regulation of neurogenesis | GO:0050767 | 169 | 43 | 18.57 | 2.32 | 1.59E-07 | 7.39E-06 |
| definitive hemopoiesis | GO:0060216 | 59 | 15 | 6.48 | 2.31 | 1.43E-03 | 2.30E-02 |
| primitive hemopoiesis | GO:0060215 | 59 | 15 | 6.48 | 2.31 | 1.43E-03 | 2.29E-02 |
| regulation of anatomical structure morphogenesis | GO:0022603 | 327 | 83 | 35.92 | 2.31 | 1.84E-13 | 1.49E-11 |
| swimming behavior | GO:0036269 | 71 | 18 | 7.80 | 2.31 | 7.75E-04 | 1.43E-02 |
| embryonic heart tube morphogenesis | GO:0003143 | 142 | 36 | 15.60 | 2.31 | 1.22E-06 | 4.98E-05 |
| convergent extension involved in gastrulation | GO:0060027 | 83 | 21 | 9.12 | 2.30 | 2.54E-04 | 5.63E-03 |
| fin regeneration | GO:0031101 | 87 | 22 | 9.56 | 2.30 | 1.77E-04 | 4.15E-03 |
| regulation of cell growth | GO:0001558 | 103 | 26 | 11.32 | 2.30 | 4.26E-05 | 1.18E-03 |
| neuron projection guidance | GO:0097485 | 262 | 66 | 28.78 | 2.29 | 9.67E-11 | 6.55E-09 |
| axon guidance | GO:0007411 | 262 | 66 | 28.78 | 2.29 | 9.67E-11 | 6.50E-09 |
| positive regulation of RNA biosynthetic process | GO:1902680 | 488 | 122 | 53.61 | 2.28 | 1.27E-18 | 1.68E-16 |
| positive regulation of DNA-templated transcription | GO:0045893 | 488 | 122 | 53.61 | 2.28 | 1.27E-18 | 1.66E-16 |
| monoatomic anion transmembrane transport | GO:0098656 | 72 | 18 | 7.91 | 2.28 | 8.51E-04 | 1.55E-02 |
| embryonic organ morphogenesis | GO:0048562 | 442 | 110 | 48.56 | 2.27 | 9.79E-17 | 1.12E-14 |
| kidney development | GO:0001822 | 210 | 52 | 23.07 | 2.25 | 2.10E-08 | 1.08E-06 |
| regulation of cell differentiation | GO:0045595 | 493 | 122 | 54.16 | 2.25 | 3.40E-18 | 4.19E-16 |
| cell migration involved in gastrulation | GO:0042074 | 81 | 20 | 8.90 | 2.25 | 4.60E-04 | 9.39E-03 |
| regulation of cell projection organization | GO:0031344 | 231 | 57 | 25.38 | 2.25 | 3.87E-09 | 2.19E-07 |
| chordate embryonic development | GO:0043009 | 527 | 130 | 57.89 | 2.25 | 4.44E-19 | 6.45E-17 |
| microtubule cytoskeleton organization involved in mitosis | GO:1902850 | 73 | 18 | 8.02 | 2.24 | 9.46E-04 | 1.67E-02 |
| renal system development | GO:0072001 | 211 | 52 | 23.18 | 2.24 | 2.26E-08 | 1.14E-06 |

| | | | | | | | |
|---|---|---|---|---|---|---|---|
| tRNA modification | GO:0006400 | 69 | 17 | 7.58 | 2.24 | 1.36E-03 | 2.25E-02 |
| tube development | GO:0035295 | 780 | 192 | 85.69 | 2.24 | 1.04E-27 | 3.74E-25 |
| regulation of cell shape | GO:0008360 | 65 | 16 | 7.14 | 2.24 | 1.97E-03 | 2.99E-02 |
| heart looping | GO:0001947 | 126 | 31 | 13.84 | 2.24 | 1.89E-05 | 5.79E-04 |
| gland development | GO:0048732 | 252 | 62 | 27.68 | 2.24 | 7.99E-10 | 4.92E-08 |
| embryo development ending in birth or egg hatching | GO:0009792 | 530 | 130 | 58.22 | 2.23 | 5.95E-19 | 8.37E-17 |
| regulation of RNA splicing | GO:0043484 | 102 | 25 | 11.21 | 2.23 | 9.29E-05 | 2.32E-03 |
| heart morphogenesis | GO:0003007 | 241 | 59 | 26.47 | 2.23 | 2.52E-09 | 1.45E-07 |
| regulation of plasma membrane bounded cell projection organization | GO:0120035 | 229 | 56 | 25.16 | 2.23 | 6.94E-09 | 3.76E-07 |
| tissue regeneration | GO:0042246 | 135 | 33 | 14.83 | 2.23 | 9.95E-06 | 3.29E-04 |
| positive regulation of cell projection organization | GO:0031346 | 86 | 21 | 9.45 | 2.22 | 3.68E-04 | 7.60E-03 |
| notochord development | GO:0030903 | 82 | 20 | 9.01 | 2.22 | 5.23E-04 | 1.05E-02 |
| axonogenesis | GO:0007409 | 390 | 95 | 42.84 | 2.22 | 4.79E-14 | 4.14E-12 |
| regeneration | GO:0031099 | 189 | 46 | 20.76 | 2.22 | 2.28E-07 | 1.03E-05 |
| axon development | GO:0061564 | 433 | 105 | 47.57 | 2.21 | 3.19E-15 | 3.05E-13 |
| embryonic organ development | GO:0048568 | 628 | 152 | 68.99 | 2.20 | 2.15E-21 | 3.79E-19 |
| bone development | GO:0060348 | 62 | 15 | 6.81 | 2.20 | 3.16E-03 | 4.39E-02 |
| cilium movement | GO:0003341 | 91 | 22 | 10.00 | 2.20 | 3.00E-04 | 6.45E-03 |
| regulation of growth | GO:0040008 | 141 | 34 | 15.49 | 2.20 | 8.76E-06 | 2.96E-04 |
| negative regulation of Wnt signaling pathway | GO:0030178 | 83 | 20 | 9.12 | 2.19 | 5.98E-04 | 1.16E-02 |
| epithelial tube morphogenesis | GO:0060562 | 249 | 60 | 27.35 | 2.19 | 4.37E-09 | 2.44E-07 |
| neuron projection morphogenesis | GO:0048812 | 446 | 107 | 49.00 | 2.18 | 4.18E-15 | 3.92E-13 |
| regulation of nervous system development | GO:0051960 | 196 | 47 | 21.53 | 2.18 | 2.15E-07 | 9.87E-06 |
| ear morphogenesis | GO:0042471 | 96 | 23 | 10.55 | 2.18 | 2.47E-04 | 5.57E-03 |
| plasma membrane bounded cell projection morphogenesis | GO:0120039 | 447 | 107 | 49.11 | 2.18 | 4.68E-15 | 4.35E-13 |
| response to oxidative stress | GO:0006979 | 67 | 16 | 7.36 | 2.17 | 2.46E-03 | 3.56E-02 |
| positive regulation of RNA metabolic process | GO:0051254 | 557 | 133 | 61.19 | 2.17 | 2.69E-18 | 3.36E-16 |
| cell morphogenesis involved in neuron differentiation | GO:0048667 | 415 | 99 | 45.59 | 2.17 | 7.40E-14 | 6.23E-12 |
| retina morphogenesis in camera-type eye | GO:0060042 | 130 | 31 | 14.28 | 2.17 | 2.77E-05 | 8.35E-04 |
| blood vessel morphogenesis | GO:0048514 | 358 | 85 | 39.33 | 2.16 | 5.15E-12 | 3.83E-10 |
| cell projection morphogenesis | GO:0048858 | 451 | 107 | 49.54 | 2.16 | 1.15E-14 | 1.03E-12 |
| liver development | GO:0001889 | 152 | 36 | 16.70 | 2.16 | 1.02E-05 | 3.34E-04 |
| establishment or maintenance of cell polarity | GO:0007163 | 114 | 27 | 12.52 | 2.16 | 1.12E-04 | 2.74E-03 |
| epidermis development | GO:0008544 | 106 | 25 | 11.64 | 2.15 | 2.27E-04 | 5.17E-03 |
| negative regulation of cell differentiation | GO:0045596 | 123 | 29 | 13.51 | 2.15 | 6.21E-05 | 1.66E-03 |
| cartilage development | GO:0051216 | 157 | 37 | 17.25 | 2.15 | 7.63E-06 | 2.61E-04 |
| transition metal ion transport | GO:0000041 | 68 | 16 | 7.47 | 2.14 | 2.77E-03 | 3.95E-02 |
| positive regulation of developmental process | GO:0051094 | 268 | 63 | 29.44 | 2.14 | 5.03E-09 | 2.80E-07 |
| somite development | GO:0061053 | 149 | 35 | 16.37 | 2.14 | 1.55E-05 | 4.92E-04 |
| regulation of multicellular organismal development | GO:2000026 | 456 | 107 | 50.09 | 2.14 | 1.87E-14 | 1.63E-12 |

| | | | | | | | |
|---|---|---|---|---|---|---|---|
| cellular response to hormone stimulus | GO:0032870 | 180 | 42 | 19.77 | 2.12 | 1.89E-06 | 7.31E-05 |
| regulation of BMP signaling pathway | GO:0030510 | 90 | 21 | 9.89 | 2.12 | 9.60E-04 | 1.69E-02 |
| cranial skeletal system development | GO:1904888 | 245 | 57 | 26.91 | 2.12 | 4.38E-08 | 2.18E-06 |
| negative regulation of developmental process | GO:0051093 | 172 | 40 | 18.90 | 2.12 | 3.69E-06 | 1.35E-04 |
| blood vessel development | GO:0001568 | 413 | 96 | 45.37 | 2.12 | 8.57E-13 | 6.65E-11 |
| morphogenesis of embryonic epithelium | GO:0016331 | 69 | 16 | 7.58 | 2.11 | 3.15E-03 | 4.37E-02 |
| central nervous system development | GO:0007417 | 738 | 171 | 81.07 | 2.11 | 8.71E-22 | 1.63E-19 |
| inner ear morphogenesis | GO:0042472 | 95 | 22 | 10.44 | 2.11 | 7.20E-04 | 1.34E-02 |
| positive regulation of nucleobase-containing compound metabolic process | GO:0045935 | 596 | 138 | 65.47 | 2.11 | 1.20E-17 | 1.46E-15 |
| regulation of developmental process | GO:0050793 | 895 | 207 | 98.32 | 2.11 | 6.23E-26 | 1.93E-23 |
| tube morphogenesis | GO:0035239 | 589 | 136 | 64.70 | 2.10 | 2.59E-17 | 3.07E-15 |
| regulation of cellular component size | GO:0032535 | 208 | 48 | 22.85 | 2.10 | 5.40E-07 | 2.30E-05 |
| hepaticobiliary system development | GO:0061008 | 156 | 36 | 17.14 | 2.10 | 1.43E-05 | 4.58E-04 |
| positive regulation of macromolecule biosynthetic process | GO:0010557 | 655 | 151 | 71.95 | 2.10 | 4.07E-19 | 6.10E-17 |
| transmembrane receptor protein tyrosine kinase signaling pathway | GO:0007169 | 243 | 56 | 26.69 | 2.10 | 7.23E-08 | 3.45E-06 |
| organic hydroxy compound biosynthetic process | GO:1901617 | 100 | 23 | 10.99 | 2.09 | 5.54E-04 | 1.09E-02 |
| connective tissue development | GO:0061448 | 170 | 39 | 18.68 | 2.09 | 6.60E-06 | 2.27E-04 |
| positive regulation of cellular biosynthetic process | GO:0031328 | 668 | 153 | 73.38 | 2.08 | 4.90E-19 | 7.01E-17 |
| animal organ morphogenesis | GO:0009887 | 839 | 192 | 92.17 | 2.08 | 1.66E-23 | 3.93E-21 |
| pectoral fin development | GO:0033339 | 70 | 16 | 7.69 | 2.08 | 3.58E-03 | 4.86E-02 |
| head development | GO:0060322 | 547 | 125 | 60.09 | 2.08 | 1.36E-15 | 1.38E-13 |
| sensory organ morphogenesis | GO:0090596 | 324 | 74 | 35.59 | 2.08 | 8.83E-10 | 5.37E-08 |
| positive regulation of biosynthetic process | GO:0009891 | 670 | 153 | 73.60 | 2.08 | 5.98E-19 | 8.28E-17 |
| developmental pigmentation | GO:0048066 | 79 | 18 | 8.68 | 2.07 | 3.05E-03 | 4.26E-02 |
| cell fate specification | GO:0001708 | 101 | 23 | 11.10 | 2.07 | 6.13E-04 | 1.18E-02 |
| brain development | GO:0007420 | 527 | 120 | 57.89 | 2.07 | 4.88E-15 | 4.48E-13 |
| epithelium development | GO:0060429 | 942 | 214 | 103.48 | 2.07 | 7.25E-26 | 2.17E-23 |
| pigmentation | GO:0043473 | 141 | 32 | 15.49 | 2.07 | 6.16E-05 | 1.65E-03 |
| left/right pattern formation | GO:0060972 | 296 | 67 | 32.52 | 2.06 | 9.68E-09 | 5.13E-07 |
| determination of left/right symmetry | GO:0007368 | 292 | 66 | 32.08 | 2.06 | 1.37E-08 | 7.16E-07 |
| neuron projection development | GO:0031175 | 549 | 124 | 60.31 | 2.06 | 3.24E-15 | 3.07E-13 |
| convergent extension | GO:0060026 | 155 | 35 | 17.03 | 2.06 | 2.80E-05 | 8.41E-04 |
| camera-type eye morphogenesis | GO:0048593 | 195 | 44 | 21.42 | 2.05 | 2.86E-06 | 1.07E-04 |
| morphogenesis of an epithelium | GO:0002009 | 479 | 108 | 52.62 | 2.05 | 2.68E-13 | 2.12E-11 |
| cell population proliferation | GO:0008283 | 151 | 34 | 16.59 | 2.05 | 3.90E-05 | 1.12E-03 |
| eye morphogenesis | GO:0048592 | 240 | 54 | 26.37 | 2.05 | 3.62E-07 | 1.58E-05 |
| regulation of actin cytoskeleton organization | GO:0032956 | 187 | 42 | 20.54 | 2.04 | 5.54E-06 | 1.95E-04 |
| anatomical structure formation involved in morphogenesis | GO:0048646 | 808 | 181 | 88.76 | 2.04 | 3.22E-21 | 5.58E-19 |

| | | | | | | | |
|---|---|---|---|---|---|---|---|
| regulation of transmembrane receptor protein serine/threonine kinase signaling pathway | GO:0090092 | 125 | 28 | 13.73 | 2.04 | 2.43E-04 | 5.50E-03 |
| pattern specification process | GO:0007389 | 644 | 144 | 70.75 | 2.04 | 4.63E-17 | 5.41E-15 |
| retina development in camera-type eye | GO:0060041 | 273 | 61 | 29.99 | 2.03 | 8.04E-08 | 3.79E-06 |
| monoatomic anion transport | GO:0006820 | 121 | 27 | 13.29 | 2.03 | 3.43E-04 | 7.19E-03 |
| regulation of small GTPase mediated signal transduction | GO:0051056 | 121 | 27 | 13.29 | 2.03 | 3.43E-04 | 7.17E-03 |
| heart development | GO:0007507 | 583 | 130 | 64.05 | 2.03 | 2.20E-15 | 2.15E-13 |
| pancreas development | GO:0031016 | 157 | 35 | 17.25 | 2.03 | 4.91E-05 | 1.34E-03 |
| anterior/posterior pattern specification | GO:0009952 | 247 | 55 | 27.13 | 2.03 | 3.13E-07 | 1.38E-05 |
| regionalization | GO:0003002 | 629 | 140 | 69.10 | 2.03 | 1.93E-16 | 2.07E-14 |
| transforming growth factor beta receptor superfamily signaling pathway | GO:0141091 | 90 | 20 | 9.89 | 2.02 | 1.91E-03 | 2.92E-02 |
| regulation of actin filament-based process | GO:0032970 | 189 | 42 | 20.76 | 2.02 | 1.01E-05 | 3.34E-04 |
| determination of bilateral symmetry | GO:0009855 | 320 | 71 | 35.15 | 2.02 | 7.27E-09 | 3.92E-07 |
| specification of symmetry | GO:0009799 | 320 | 71 | 35.15 | 2.02 | 7.27E-09 | 3.89E-07 |
| tissue morphogenesis | GO:0048729 | 571 | 126 | 62.73 | 2.01 | 1.22E-14 | 1.09E-12 |
| determination of heart left/right asymmetry | GO:0061371 | 204 | 45 | 22.41 | 2.01 | 4.59E-06 | 1.66E-04 |
| microtubule-based transport | GO:0099111 | 109 | 24 | 11.97 | 2.00 | 9.70E-04 | 1.71E-02 |
| enzyme-linked receptor protein signaling pathway | GO:0007167 | 350 | 77 | 38.45 | 2.00 | 2.41E-09 | 1.40E-07 |
| response to hormone | GO:0009725 | 232 | 51 | 25.49 | 2.00 | 1.29E-06 | 5.20E-05 |
| RNA catabolic process | GO:0006401 | 91 | 20 | 10.00 | 2.00 | 2.12E-03 | 3.17E-02 |
| cell migration | GO:0016477 | 619 | 136 | 68.00 | 2.00 | 1.75E-15 | 1.73E-13 |
| positive regulation of cellular metabolic process | GO:0031325 | 926 | 203 | 101.73 | 2.00 | 1.61E-22 | 3.29E-20 |
| neural retina development | GO:0003407 | 105 | 23 | 11.53 | 1.99 | 1.37E-03 | 2.24E-02 |
| digestive tract development | GO:0048565 | 105 | 23 | 11.53 | 1.99 | 1.37E-03 | 2.23E-02 |
| cell morphogenesis | GO:0000902 | 548 | 120 | 60.20 | 1.99 | 9.55E-14 | 7.96E-12 |
| positive regulation of phosphorylation | GO:0042327 | 174 | 38 | 19.11 | 1.99 | 4.18E-05 | 1.18E-03 |
| regulation of angiogenesis | GO:0045765 | 87 | 19 | 9.56 | 1.99 | 2.98E-03 | 4.17E-02 |
| response to growth factor | GO:0070848 | 156 | 34 | 17.14 | 1.98 | 8.69E-05 | 2.22E-03 |
| embryo development | GO:0009790 | 1212 | 264 | 133.14 | 1.98 | 1.01E-28 | 3.96E-26 |
| cardiac muscle tissue development | GO:0048738 | 101 | 22 | 11.10 | 1.98 | 1.95E-03 | 2.96E-02 |
| circulatory system development | GO:0072359 | 981 | 213 | 107.77 | 1.98 | 5.97E-23 | 1.31E-20 |
| cellular anatomical entity morphogenesis | GO:0032989 | 562 | 122 | 61.74 | 1.98 | 1.15E-13 | 9.43E-12 |
| tRNA processing | GO:0008033 | 106 | 23 | 11.64 | 1.98 | 1.46E-03 | 2.34E-02 |
| appendage development | GO:0048736 | 120 | 26 | 13.18 | 1.97 | 6.23E-04 | 1.20E-02 |
| regulation of cellular response to growth factor stimulus | GO:0090287 | 162 | 35 | 17.80 | 1.97 | 7.59E-05 | 1.97E-03 |
| cellular response to endogenous stimulus | GO:0071495 | 385 | 83 | 42.29 | 1.96 | 1.73E-09 | 1.03E-07 |
| neuron development | GO:0048666 | 725 | 156 | 79.64 | 1.96 | 1.24E-16 | 1.38E-14 |
| positive regulation of phosphorus metabolic process | GO:0010562 | 186 | 40 | 20.43 | 1.96 | 2.85E-05 | 8.48E-04 |
| positive regulation of phosphate metabolic process | GO:0045937 | 186 | 40 | 20.43 | 1.96 | 2.85E-05 | 8.46E-04 |
| positive regulation of cell migration | GO:0030335 | 112 | 24 | 12.30 | 1.95 | 1.22E-03 | 2.09E-02 |

| | | | | | | | |
|---|---|---|---|---|---|---|---|
| neuron differentiation | GO:0030182 | 903 | 193 | 99.20 | 1.95 | 3.64E-20 | 5.85E-18 |
| striated muscle tissue development | GO:0014706 | 103 | 22 | 11.32 | 1.94 | 2.21E-03 | 3.26E-02 |
| vasculature development | GO:0001944 | 492 | 105 | 54.05 | 1.94 | 1.77E-11 | 1.27E-09 |
| embryonic morphogenesis | GO:0048598 | 722 | 154 | 79.32 | 1.94 | 3.33E-16 | 3.44E-14 |
| nervous system development | GO:0007399 | 1684 | 359 | 185.00 | 1.94 | 4.73E-37 | 2.66E-34 |
| cellular response to growth factor stimulus | GO:0071363 | 155 | 33 | 17.03 | 1.94 | 2.37E-04 | 5.39E-03 |
| growth | GO:0040007 | 296 | 63 | 32.52 | 1.94 | 2.77E-07 | 1.23E-05 |
| developmental growth | GO:0048589 | 296 | 63 | 32.52 | 1.94 | 2.77E-07 | 1.23E-05 |
| response to endogenous stimulus | GO:0009719 | 437 | 93 | 48.01 | 1.94 | 3.44E-10 | 2.25E-08 |
| negative regulation of cellular component organization | GO:0051129 | 184 | 39 | 20.21 | 1.93 | 7.04E-05 | 1.84E-03 |
| cell motility | GO:0048870 | 670 | 142 | 73.60 | 1.93 | 8.57E-15 | 7.79E-13 |
| negative regulation of transcription by RNA polymerase II | GO:0000122 | 255 | 54 | 28.01 | 1.93 | 2.45E-06 | 9.27E-05 |
| generation of neurons | GO:0048699 | 934 | 197 | 102.60 | 1.92 | 8.79E-20 | 1.34E-17 |
| RNA splicing | GO:0008380 | 247 | 52 | 27.13 | 1.92 | 4.71E-06 | 1.70E-04 |
| tissue development | GO:0009888 | 1511 | 318 | 165.99 | 1.92 | 1.02E-31 | 4.61E-29 |
| microtubule-based movement | GO:0007018 | 219 | 46 | 24.06 | 1.91 | 1.66E-05 | 5.15E-04 |
| myeloid cell homeostasis | GO:0002262 | 124 | 26 | 13.62 | 1.91 | 1.25E-03 | 2.14E-02 |
| developmental process involved in reproduction | GO:0003006 | 172 | 36 | 18.90 | 1.91 | 1.24E-04 | 3.00E-03 |
| modulation of chemical synaptic transmission | GO:0050804 | 153 | 32 | 16.81 | 1.90 | 3.64E-04 | 7.57E-03 |
| regulation of trans-synaptic signaling | GO:0099177 | 153 | 32 | 16.81 | 1.90 | 3.64E-04 | 7.56E-03 |
| positive regulation of protein phosphorylation | GO:0001934 | 110 | 23 | 12.08 | 1.90 | 2.03E-03 | 3.06E-02 |
| neurogenesis | GO:0022008 | 1070 | 223 | 117.54 | 1.90 | 9.91E-22 | 1.78E-19 |
| regulation of anatomical structure size | GO:0090066 | 264 | 55 | 29.00 | 1.90 | 3.75E-06 | 1.36E-04 |
| dorsal/ventral pattern formation | GO:0009953 | 154 | 32 | 16.92 | 1.89 | 3.87E-04 | 7.97E-03 |
| glial cell differentiation | GO:0010001 | 140 | 29 | 15.38 | 1.89 | 9.35E-04 | 1.66E-02 |
| cell fate commitment | GO:0045165 | 174 | 36 | 19.11 | 1.88 | 2.06E-04 | 4.76E-03 |
| behavior | GO:0007610 | 237 | 49 | 26.04 | 1.88 | 1.37E-05 | 4.43E-04 |
| gliogenesis | GO:0042063 | 150 | 31 | 16.48 | 1.88 | 5.36E-04 | 1.06E-02 |
| positive regulation of macromolecule metabolic process | GO:0010604 | 930 | 192 | 102.16 | 1.88 | 2.62E-18 | 3.32E-16 |
| positive regulation of metabolic process | GO:0009893 | 1037 | 214 | 113.92 | 1.88 | 3.25E-20 | 5.32E-18 |
| RNA modification | GO:0009451 | 126 | 26 | 13.84 | 1.88 | 1.43E-03 | 2.30E-02 |
| positive regulation of cell population proliferation | GO:0008284 | 161 | 33 | 17.69 | 1.87 | 3.53E-04 | 7.37E-03 |
| positive regulation of nitrogen compound metabolic process | GO:0051173 | 854 | 175 | 93.82 | 1.87 | 1.78E-16 | 1.93E-14 |
| anatomical structure morphogenesis | GO:0009653 | 2211 | 453 | 242.89 | 1.87 | 1.32E-42 | 9.91E-40 |
| negative regulation of multicellular organismal process | GO:0051241 | 206 | 42 | 22.63 | 1.86 | 7.23E-05 | 1.88E-03 |
| plasma membrane bounded cell projection organization | GO:0120036 | 885 | 180 | 97.22 | 1.85 | 1.44E-16 | 1.58E-14 |
| regulation of canonical Wnt signaling pathway | GO:0060828 | 118 | 24 | 12.96 | 1.85 | 2.74E-03 | 3.92E-02 |
| regulation of locomotion | GO:0040012 | 271 | 55 | 29.77 | 1.85 | 8.19E-06 | 2.79E-04 |
| regulation of multicellular organismal process | GO:0051239 | 838 | 170 | 92.06 | 1.85 | 1.55E-15 | 1.55E-13 |
| cell development | GO:0048468 | 1824 | 370 | 200.37 | 1.85 | 1.84E-33 | 8.72E-31 |

| | | | | | | | |
|---|---|---|---|---|---|---|---|
| Wnt signaling pathway | GO:0016055 | 158 | 32 | 17.36 | 1.84 | 7.49E-04 | 1.39E-02 |
| actin filament organization | GO:0007015 | 252 | 51 | 27.68 | 1.84 | 1.62E-05 | 5.10E-04 |
| inorganic ion homeostasis | GO:0098771 | 213 | 43 | 23.40 | 1.84 | 9.26E-05 | 2.32E-03 |
| system development | GO:0048731 | 3290 | 664 | 361.42 | 1.84 | 4.06E-62 | 4.57E-59 |
| cell projection organization | GO:0030030 | 907 | 183 | 99.64 | 1.84 | 2.05E-16 | 2.17E-14 |
| aromatic compound catabolic process | GO:0019439 | 243 | 49 | 26.69 | 1.84 | 2.87E-05 | 8.49E-04 |
| lateral line development | GO:0048882 | 124 | 25 | 13.62 | 1.84 | 2.31E-03 | 3.39E-02 |
| sensory organ development | GO:0007423 | 748 | 150 | 82.17 | 1.83 | 2.20E-13 | 1.75E-11 |
| regulation of cellular component organization | GO:0051128 | 785 | 157 | 86.24 | 1.82 | 5.95E-14 | 5.10E-12 |
| cell-cell signaling by wnt | GO:0198738 | 160 | 32 | 17.58 | 1.82 | 8.20E-04 | 1.50E-02 |
| positive regulation of cell motility | GO:2000147 | 120 | 24 | 13.18 | 1.82 | 3.18E-03 | 4.39E-02 |
| cell-cell adhesion | GO:0098609 | 335 | 67 | 36.80 | 1.82 | 1.48E-06 | 5.91E-05 |
| positive regulation of multicellular organismal process | GO:0051240 | 296 | 59 | 32.52 | 1.81 | 5.57E-06 | 1.95E-04 |
| epithelial cell differentiation | GO:0030855 | 256 | 51 | 28.12 | 1.81 | 2.97E-05 | 8.75E-04 |
| hemopoiesis | GO:0030097 | 457 | 91 | 50.20 | 1.81 | 1.62E-08 | 8.37E-07 |
| regulation of mRNA metabolic process | GO:1903311 | 156 | 31 | 17.14 | 1.81 | 1.12E-03 | 1.96E-02 |
| digestive system development | GO:0055123 | 146 | 29 | 16.04 | 1.81 | 1.95E-03 | 2.96E-02 |
| regulation of cytoskeleton organization | GO:0051493 | 267 | 53 | 29.33 | 1.81 | 1.87E-05 | 5.76E-04 |
| regulation of actin filament organization | GO:0110053 | 131 | 26 | 14.39 | 1.81 | 2.84E-03 | 4.02E-02 |
| multicellular organism development | GO:0007275 | 3850 | 762 | 422.94 | 1.80 | 4.69E-69 | 7.03E-66 |
| regulation of supramolecular fiber organization | GO:1902903 | 167 | 33 | 18.35 | 1.80 | 7.27E-04 | 1.35E-02 |
| regulation of cell migration | GO:0030334 | 233 | 46 | 25.60 | 1.80 | 8.05E-05 | 2.07E-03 |
| animal organ development | GO:0048513 | 2532 | 499 | 278.15 | 1.79 | 2.91E-42 | 2.02E-39 |
| negative regulation of RNA metabolic process | GO:0051253 | 406 | 80 | 44.60 | 1.79 | 2.18E-07 | 9.94E-06 |
| mRNA processing | GO:0006397 | 310 | 61 | 34.05 | 1.79 | 5.95E-06 | 2.06E-04 |
| regulation of GTPase activity | GO:0043087 | 163 | 32 | 17.91 | 1.79 | 9.89E-04 | 1.73E-02 |
| organic cyclic compound catabolic process | GO:1901361 | 250 | 49 | 27.46 | 1.78 | 5.97E-05 | 1.60E-03 |
| homeostasis of number of cells | GO:0048872 | 148 | 29 | 16.26 | 1.78 | 2.10E-03 | 3.15E-02 |
| positive regulation of protein modification process | GO:0031401 | 138 | 27 | 15.16 | 1.78 | 2.51E-03 | 3.62E-02 |
| cellular developmental process | GO:0048869 | 2509 | 489 | 275.63 | 1.77 | 5.94E-40 | 3.82E-37 |
| cell differentiation | GO:0030154 | 2505 | 488 | 275.19 | 1.77 | 8.28E-40 | 4.97E-37 |
| negative regulation of intracellular signal transduction | GO:1902532 | 149 | 29 | 16.37 | 1.77 | 2.21E-03 | 3.26E-02 |
| negative regulation of biosynthetic process | GO:0009890 | 612 | 119 | 67.23 | 1.77 | 5.00E-10 | 3.14E-08 |
| actin cytoskeleton organization | GO:0030036 | 463 | 90 | 50.86 | 1.77 | 7.19E-08 | 3.44E-06 |
| gamete generation | GO:0007276 | 170 | 33 | 18.68 | 1.77 | 1.19E-03 | 2.06E-02 |
| cell junction organization | GO:0034330 | 232 | 45 | 25.49 | 1.77 | 1.86E-04 | 4.36E-03 |
| negative regulation of RNA biosynthetic process | GO:1902679 | 372 | 72 | 40.87 | 1.76 | 1.56E-06 | 6.18E-05 |
| negative regulation of DNA-templated transcription | GO:0045892 | 372 | 72 | 40.87 | 1.76 | 1.56E-06 | 6.15E-05 |
| cell surface receptor signaling pathway | GO:0007166 | 1075 | 208 | 118.09 | 1.76 | 2.16E-16 | 2.26E-14 |
| cellular homeostasis | GO:0019725 | 274 | 53 | 30.10 | 1.76 | 3.84E-05 | 1.11E-03 |

| | | | | | | | |
|---|---|---|---|---|---|---|---|
| small GTPase-mediated signal transduction | GO:0007264 | 176 | 34 | 19.33 | 1.76 | 9.79E-04 | 1.72E-02 |
| positive regulation of cellular component organization | GO:0051130 | 223 | 43 | 24.50 | 1.76 | 2.24E-04 | 5.11E-03 |
| mitotic cell cycle process | GO:1903047 | 244 | 47 | 26.80 | 1.75 | 1.21E-04 | 2.94E-03 |
| methylation | GO:0032259 | 203 | 39 | 22.30 | 1.75 | 6.21E-04 | 1.19E-02 |
| regulation of Wnt signaling pathway | GO:0030111 | 172 | 33 | 18.90 | 1.75 | 1.32E-03 | 2.20E-02 |
| multicellular organismal reproductive process | GO:0048609 | 172 | 33 | 18.90 | 1.75 | 1.32E-03 | 2.19E-02 |
| multicellular organism reproduction | GO:0032504 | 178 | 34 | 19.55 | 1.74 | 1.54E-03 | 2.45E-02 |
| endocytosis | GO:0006897 | 278 | 53 | 30.54 | 1.74 | 6.43E-05 | 1.70E-03 |
| mRNA metabolic process | GO:0016071 | 383 | 73 | 42.07 | 1.74 | 3.14E-06 | 1.17E-04 |
| negative regulation of macromolecule biosynthetic process | GO:0010558 | 605 | 115 | 66.46 | 1.73 | 3.52E-09 | 2.00E-07 |
| actin filament-based process | GO:0030029 | 481 | 91 | 52.84 | 1.72 | 2.02E-07 | 9.35E-06 |
| heterocycle catabolic process | GO:0046700 | 222 | 42 | 24.39 | 1.72 | 4.84E-04 | 9.84E-03 |
| cellular nitrogen compound catabolic process | GO:0044270 | 222 | 42 | 24.39 | 1.72 | 4.84E-04 | 9.82E-03 |
| negative regulation of cellular biosynthetic process | GO:0031327 | 608 | 115 | 66.79 | 1.72 | 5.42E-09 | 2.98E-07 |
| monoatomic ion homeostasis | GO:0050801 | 254 | 48 | 27.90 | 1.72 | 1.67E-04 | 3.96E-03 |
| positive regulation of cellular process | GO:0048522 | 1918 | 362 | 210.70 | 1.72 | 2.43E-26 | 8.41E-24 |
| synaptic signaling | GO:0099536 | 297 | 56 | 32.63 | 1.72 | 5.15E-05 | 1.41E-03 |
| response to oxygen-containing compound | GO:1901700 | 345 | 65 | 37.90 | 1.72 | 1.27E-05 | 4.11E-04 |
| anatomical structure development | GO:0048856 | 4903 | 923 | 538.62 | 1.71 | 9.10E-75 | 1.64E-71 |
| developmental process | GO:0032502 | 5052 | 949 | 554.99 | 1.71 | 8.51E-77 | 1.92E-73 |
| muscle cell differentiation | GO:0042692 | 261 | 49 | 28.67 | 1.71 | 1.98E-04 | 4.61E-03 |
| chemical homeostasis | GO:0048878 | 389 | 73 | 42.73 | 1.71 | 5.62E-06 | 1.96E-04 |
| eye development | GO:0001654 | 549 | 103 | 60.31 | 1.71 | 5.51E-08 | 2.68E-06 |
| visual system development | GO:0150063 | 549 | 103 | 60.31 | 1.71 | 5.51E-08 | 2.67E-06 |
| negative regulation of nucleobase-containing compound metabolic process | GO:0045934 | 448 | 84 | 49.21 | 1.71 | 8.75E-07 | 3.60E-05 |
| regulation of cell population proliferation | GO:0042127 | 305 | 57 | 33.51 | 1.70 | 6.32E-05 | 1.68E-03 |
| plasma membrane bounded cell projection assembly | GO:0120031 | 327 | 61 | 35.92 | 1.70 | 3.58E-05 | 1.04E-03 |
| regulation of cell motility | GO:2000145 | 252 | 47 | 27.68 | 1.70 | 3.42E-04 | 7.20E-03 |
| homeostatic process | GO:0042592 | 655 | 122 | 71.95 | 1.70 | 4.32E-09 | 2.43E-07 |
| vesicle organization | GO:0016050 | 173 | 32 | 19.00 | 1.68 | 3.12E-03 | 4.34E-02 |
| mitotic cell cycle | GO:0000278 | 309 | 57 | 33.95 | 1.68 | 1.04E-04 | 2.57E-03 |
| cell projection assembly | GO:0030031 | 347 | 64 | 38.12 | 1.68 | 2.98E-05 | 8.76E-04 |
| multicellular organismal process | GO:0032501 | 4814 | 885 | 528.84 | 1.67 | 1.31E-65 | 1.69E-62 |
| regulation of transferase activity | GO:0051338 | 185 | 34 | 20.32 | 1.67 | 2.87E-03 | 4.06E-02 |
| positive regulation of biological process | GO:0048518 | 2136 | 391 | 234.65 | 1.67 | 9.08E-26 | 2.55E-23 |
| chromatin organization | GO:0006325 | 317 | 58 | 34.82 | 1.67 | 9.13E-05 | 2.31E-03 |
| regulation of phosphorylation | GO:0042325 | 279 | 51 | 30.65 | 1.66 | 3.24E-04 | 6.94E-03 |
| cell division | GO:0051301 | 263 | 48 | 28.89 | 1.66 | 4.61E-04 | 9.39E-03 |
| cell adhesion | GO:0007155 | 570 | 104 | 62.62 | 1.66 | 2.08E-07 | 9.58E-06 |

| | | | | | | | |
|---|---|---|---|---|---|---|---|
| chromatin remodeling | GO:0006338 | 203 | 37 | 22.30 | 1.66 | 2.12E-03 | 3.17E-02 |
| sensory system development | GO:0048880 | 664 | 121 | 72.94 | 1.66 | 2.40E-08 | 1.21E-06 |
| regulation of transcription by RNA polymerase II | GO:0006357 | 1810 | 329 | 198.84 | 1.65 | 4.00E-21 | 6.79E-19 |
| cell-cell signaling | GO:0007267 | 534 | 97 | 58.66 | 1.65 | 5.78E-07 | 2.44E-05 |
| chemical synaptic transmission | GO:0007268 | 281 | 51 | 30.87 | 1.65 | 3.42E-04 | 7.24E-03 |
| anterograde trans-synaptic signaling | GO:0098916 | 281 | 51 | 30.87 | 1.65 | 3.42E-04 | 7.22E-03 |
| small molecule biosynthetic process | GO:0044283 | 281 | 51 | 30.87 | 1.65 | 3.42E-04 | 7.20E-03 |
| camera-type eye development | GO:0043010 | 475 | 86 | 52.18 | 1.65 | 3.63E-06 | 1.33E-04 |
| mRNA splicing, via spliceosome | GO:0000398 | 188 | 34 | 20.65 | 1.65 | 3.24E-03 | 4.47E-02 |
| RNA splicing, via transesterification reactions with bulged adenosine as nucleophile | GO:0000377 | 188 | 34 | 20.65 | 1.65 | 3.24E-03 | 4.46E-02 |
| RNA splicing, via transesterification reactions | GO:0000375 | 188 | 34 | 20.65 | 1.65 | 3.24E-03 | 4.45E-02 |
| response to abiotic stimulus | GO:0009628 | 360 | 65 | 39.55 | 1.64 | 5.78E-05 | 1.57E-03 |
| regulation of organelle organization | GO:0033043 | 445 | 80 | 48.89 | 1.64 | 1.11E-05 | 3.61E-04 |
| negative regulation of cellular metabolic process | GO:0031324 | 763 | 137 | 83.82 | 1.63 | 6.45E-09 | 3.52E-07 |
| regulation of response to external stimulus | GO:0032101 | 290 | 52 | 31.86 | 1.63 | 4.23E-04 | 8.68E-03 |
| negative regulation of response to stimulus | GO:0048585 | 542 | 97 | 59.54 | 1.63 | 1.38E-06 | 5.53E-05 |
| trans-synaptic signaling | GO:0099537 | 285 | 51 | 31.31 | 1.63 | 5.42E-04 | 1.07E-02 |
| dephosphorylation | GO:0016311 | 218 | 39 | 23.95 | 1.63 | 2.15E-03 | 3.20E-02 |
| small molecule catabolic process | GO:0044282 | 202 | 36 | 22.19 | 1.62 | 3.17E-03 | 4.39E-02 |
| reproduction | GO:0000003 | 304 | 54 | 33.40 | 1.62 | 4.08E-04 | 8.38E-03 |
| negative regulation of signal transduction | GO:0009968 | 445 | 79 | 48.89 | 1.62 | 1.63E-05 | 5.13E-04 |
| organic hydroxy compound metabolic process | GO:1901615 | 220 | 39 | 24.17 | 1.61 | 2.35E-03 | 3.42E-02 |
| negative regulation of cell communication | GO:0010648 | 458 | 81 | 50.31 | 1.61 | 1.52E-05 | 4.84E-04 |
| negative regulation of signaling | GO:0023057 | 458 | 81 | 50.31 | 1.61 | 1.52E-05 | 4.82E-04 |
| negative regulation of metabolic process | GO:0009892 | 832 | 147 | 91.40 | 1.61 | 5.06E-09 | 2.79E-07 |
| muscle structure development | GO:0061061 | 374 | 66 | 41.09 | 1.61 | 1.13E-04 | 2.76E-03 |
| reproductive process | GO:0022414 | 301 | 53 | 33.07 | 1.60 | 5.48E-04 | 1.08E-02 |
| muscle cell development | GO:0055001 | 216 | 38 | 23.73 | 1.60 | 3.08E-03 | 4.29E-02 |
| monoatomic cation homeostasis | GO:0055080 | 239 | 42 | 26.26 | 1.60 | 2.37E-03 | 3.45E-02 |
| regulation of signaling | GO:0023051 | 1368 | 240 | 150.28 | 1.60 | 1.12E-13 | 9.26E-12 |
| cilium assembly | GO:0060271 | 303 | 53 | 33.29 | 1.59 | 5.90E-04 | 1.15E-02 |
| cellular component organization | GO:0016043 | 3514 | 614 | 386.03 | 1.59 | 1.45E-35 | 7.26E-33 |
| response to organic substance | GO:0010033 | 727 | 127 | 79.86 | 1.59 | 1.36E-07 | 6.38E-06 |
| regulation of cell communication | GO:0010646 | 1363 | 238 | 149.73 | 1.59 | 2.04E-13 | 1.64E-11 |
| positive regulation of molecular function | GO:0044093 | 367 | 64 | 40.32 | 1.59 | 1.95E-04 | 4.56E-03 |
| protein-DNA complex organization | GO:0071824 | 350 | 61 | 38.45 | 1.59 | 2.74E-04 | 6.06E-03 |
| import into cell | GO:0098657 | 408 | 71 | 44.82 | 1.58 | 8.47E-05 | 2.16E-03 |
| regulation of biological quality | GO:0065008 | 1082 | 188 | 118.86 | 1.58 | 1.56E-10 | 1.04E-08 |
| microtubule-based process | GO:0007017 | 495 | 86 | 54.38 | 1.58 | 1.65E-05 | 5.15E-04 |
| cellular component organization or biogenesis | GO:0071840 | 3697 | 642 | 406.13 | 1.58 | 1.81E-36 | 9.60E-34 |

| | | | | | | | |
|---|---|---|---|---|---|---|---|
| negative regulation of gene expression | GO:0010629 | 242 | 42 | 26.58 | 1.58 | 2.61E-03 | 3.74E-02 |
| gastrulation | GO:0007369 | 242 | 42 | 26.58 | 1.58 | 2.61E-03 | 3.74E-02 |
| cytoskeleton organization | GO:0007010 | 853 | 148 | 93.71 | 1.58 | 1.53E-08 | 7.94E-07 |
| intracellular signaling cassette | GO:0141124 | 369 | 64 | 40.54 | 1.58 | 2.09E-04 | 4.82E-03 |
| regulation of RNA metabolic process | GO:0051252 | 2445 | 424 | 268.59 | 1.58 | 3.49E-23 | 7.85E-21 |
| regulation of response to stimulus | GO:0048583 | 1568 | 271 | 172.25 | 1.57 | 1.23E-14 | 1.08E-12 |
| apoptotic process | GO:0006915 | 249 | 43 | 27.35 | 1.57 | 2.92E-03 | 4.11E-02 |
| transmembrane transport | GO:0055085 | 1014 | 175 | 111.39 | 1.57 | 1.17E-09 | 7.07E-08 |
| negative regulation of macromolecule metabolic process | GO:0010605 | 800 | 138 | 87.88 | 1.57 | 7.62E-08 | 3.61E-06 |
| cell cycle | GO:0007049 | 632 | 109 | 69.43 | 1.57 | 1.66E-06 | 6.54E-05 |
| negative regulation of cellular process | GO:0048523 | 1824 | 314 | 200.37 | 1.57 | 1.21E-16 | 1.36E-14 |
| regulation of biosynthetic process | GO:0009889 | 2882 | 495 | 316.60 | 1.56 | 2.51E-26 | 8.37E-24 |
| negative regulation of biological process | GO:0048519 | 1957 | 336 | 214.99 | 1.56 | 1.27E-17 | 1.52E-15 |
| regulation of DNA-templated transcription | GO:0006355 | 2249 | 386 | 247.06 | 1.56 | 3.74E-20 | 5.90E-18 |
| negative regulation of nitrogen compound metabolic process | GO:0051172 | 641 | 110 | 70.42 | 1.56 | 1.95E-06 | 7.47E-05 |
| regulation of RNA biosynthetic process | GO:2001141 | 2251 | 386 | 247.28 | 1.56 | 3.90E-20 | 6.05E-18 |
| RNA processing | GO:0006396 | 637 | 109 | 69.98 | 1.56 | 2.50E-06 | 9.42E-05 |
| regulation of nucleobase-containing compound metabolic process | GO:0019219 | 2545 | 435 | 279.58 | 1.56 | 1.53E-22 | 3.21E-20 |
| membrane organization | GO:0061024 | 357 | 61 | 39.22 | 1.56 | 4.48E-04 | 9.17E-03 |
| regulation of cellular metabolic process | GO:0031323 | 3302 | 564 | 362.74 | 1.55 | 1.32E-29 | 5.39E-27 |
| supramolecular fiber organization | GO:0097435 | 457 | 78 | 50.20 | 1.55 | 7.93E-05 | 2.05E-03 |
| regulation of cellular biosynthetic process | GO:0031326 | 2861 | 488 | 314.29 | 1.55 | 3.64E-25 | 9.65E-23 |
| regulation of macromolecule biosynthetic process | GO:0010556 | 2832 | 483 | 311.11 | 1.55 | 6.23E-25 | 1.60E-22 |
| regulation of gene expression | GO:0010468 | 2792 | 476 | 306.71 | 1.55 | 1.69E-24 | 4.21E-22 |
| phospholipid metabolic process | GO:0006644 | 264 | 45 | 29.00 | 1.55 | 2.83E-03 | 4.03E-02 |
| sexual reproduction | GO:0019953 | 264 | 45 | 29.00 | 1.55 | 2.83E-03 | 4.02E-02 |
| carbohydrate metabolic process | GO:0005975 | 289 | 49 | 31.75 | 1.54 | 2.30E-03 | 3.38E-02 |
| regulation of signal transduction | GO:0009966 | 1215 | 206 | 133.47 | 1.54 | 1.76E-10 | 1.17E-08 |
| regulation of metabolic process | GO:0019222 | 3554 | 602 | 390.42 | 1.54 | 6.95E-31 | 2.98E-28 |
| positive regulation of catalytic activity | GO:0043085 | 313 | 53 | 34.38 | 1.54 | 1.38E-03 | 2.23E-02 |
| regulation of phosphate metabolic process | GO:0019220 | 314 | 53 | 34.49 | 1.54 | 1.42E-03 | 2.29E-02 |
| regulation of phosphorus metabolic process | GO:0051174 | 314 | 53 | 34.49 | 1.54 | 1.42E-03 | 2.29E-02 |
| glycoprotein metabolic process | GO:0009100 | 273 | 46 | 29.99 | 1.53 | 3.32E-03 | 4.54E-02 |
| regulation of catalytic activity | GO:0050790 | 511 | 86 | 56.14 | 1.53 | 5.77E-05 | 1.57E-03 |
| regulation of primary metabolic process | GO:0080090 | 3151 | 529 | 346.15 | 1.53 | 7.82E-26 | 2.27E-23 |
| cell cycle process | GO:0022402 | 424 | 71 | 46.58 | 1.52 | 2.97E-04 | 6.42E-03 |
| cellular response to organic substance | GO:0071310 | 490 | 82 | 53.83 | 1.52 | 1.03E-04 | 2.56E-03 |
| cilium organization | GO:0044782 | 323 | 54 | 35.48 | 1.52 | 1.65E-03 | 2.60E-02 |
| regulation of macromolecule metabolic process | GO:0060255 | 3292 | 549 | 361.64 | 1.52 | 3.63E-26 | 1.17E-23 |

| | | | | | | | |
|---|---|---|---|---|---|---|---|
| regulation of nitrogen compound metabolic process | GO:0051171 | 3102 | 517 | 340.77 | 1.52 | 1.82E-24 | 4.43E-22 |
| cellular catabolic process | GO:0044248 | 671 | 111 | 73.71 | 1.51 | 1.08E-05 | 3.55E-04 |
| regulation of molecular function | GO:0065009 | 665 | 110 | 73.05 | 1.51 | 1.31E-05 | 4.25E-04 |
| regulation of intracellular signal transduction | GO:1902531 | 569 | 94 | 62.51 | 1.50 | 5.81E-05 | 1.57E-03 |
| nucleic acid metabolic process | GO:0090304 | 1353 | 223 | 148.63 | 1.50 | 3.43E-10 | 2.25E-08 |
| RNA metabolic process | GO:0016070 | 935 | 154 | 102.71 | 1.50 | 2.76E-07 | 1.24E-05 |
| regulation of immune system process | GO:0002682 | 378 | 62 | 41.53 | 1.49 | 1.20E-03 | 2.07E-02 |
| intracellular signal transduction | GO:0035556 | 927 | 152 | 101.84 | 1.49 | 4.37E-07 | 1.89E-05 |
| transport | GO:0006810 | 2656 | 434 | 291.77 | 1.49 | 1.18E-18 | 1.58E-16 |
| cellular response to chemical stimulus | GO:0070887 | 755 | 123 | 82.94 | 1.48 | 8.54E-06 | 2.90E-04 |
| organophosphate biosynthetic process | GO:0090407 | 351 | 57 | 38.56 | 1.48 | 2.53E-03 | 3.64E-02 |
| localization | GO:0051179 | 3034 | 491 | 333.30 | 1.47 | 2.35E-20 | 3.92E-18 |
| cell communication | GO:0007154 | 3403 | 550 | 373.84 | 1.47 | 8.05E-23 | 1.73E-20 |
| signaling | GO:0023052 | 3333 | 538 | 366.15 | 1.47 | 3.58E-22 | 7.01E-20 |
| establishment of localization | GO:0051234 | 2783 | 449 | 305.73 | 1.47 | 2.58E-18 | 3.32E-16 |
| DNA metabolic process | GO:0006259 | 441 | 71 | 48.45 | 1.47 | 9.30E-04 | 1.65E-02 |
| organic cyclic compound metabolic process | GO:1901360 | 1986 | 318 | 218.17 | 1.46 | 1.42E-12 | 1.09E-10 |
| signal transduction | GO:0007165 | 3086 | 492 | 339.01 | 1.45 | 4.41E-19 | 6.51E-17 |
| phosphorylation | GO:0016310 | 960 | 153 | 105.46 | 1.45 | 2.03E-06 | 7.76E-05 |
| cellular aromatic compound metabolic process | GO:0006725 | 1860 | 296 | 204.33 | 1.45 | 2.34E-11 | 1.65E-09 |
| heterocycle metabolic process | GO:0046483 | 1848 | 294 | 203.01 | 1.45 | 2.67E-11 | 1.85E-09 |
| organelle organization | GO:0006996 | 1935 | 307 | 212.57 | 1.44 | 1.15E-11 | 8.40E-10 |
| cellular response to stimulus | GO:0051716 | 3832 | 607 | 420.96 | 1.44 | 2.84E-23 | 6.56E-21 |
| nucleobase-containing compound metabolic process | GO:0006139 | 1742 | 274 | 191.37 | 1.43 | 4.22E-10 | 2.67E-08 |
| macromolecule modification | GO:0043412 | 1706 | 268 | 187.41 | 1.43 | 7.89E-10 | 4.90E-08 |
| phosphate-containing compound metabolic process | GO:0006796 | 1639 | 257 | 180.05 | 1.43 | 2.27E-09 | 1.33E-07 |
| phosphorus metabolic process | GO:0006793 | 1654 | 259 | 181.70 | 1.43 | 2.10E-09 | 1.24E-07 |
| biological regulation | GO:0065007 | 8661 | 1356 | 951.45 | 1.43 | 1.36E-61 | 1.36E-58 |
| regulation of biological process | GO:0050789 | 8353 | 1307 | 917.62 | 1.42 | 4.08E-58 | 3.67E-55 |
| regulation of cellular process | GO:0050794 | 7831 | 1223 | 860.27 | 1.42 | 3.31E-52 | 2.70E-49 |
| vesicle-mediated transport | GO:0016192 | 840 | 131 | 92.28 | 1.42 | 3.97E-05 | 1.13E-03 |
| protein modification by small protein conjugation or removal | GO:0070647 | 584 | 91 | 64.16 | 1.42 | 6.15E-04 | 1.19E-02 |
| monoatomic ion transport | GO:0006811 | 764 | 119 | 83.93 | 1.42 | 8.05E-05 | 2.07E-03 |
| organic substance transport | GO:0071702 | 1225 | 190 | 134.57 | 1.41 | 8.19E-07 | 3.38E-05 |
| response to stimulus | GO:0050896 | 4693 | 727 | 515.55 | 1.41 | 1.49E-25 | 4.05E-23 |
| response to chemical | GO:0042221 | 1184 | 183 | 130.07 | 1.41 | 1.47E-06 | 5.87E-05 |
| cellular lipid metabolic process | GO:0044255 | 624 | 96 | 68.55 | 1.40 | 7.22E-04 | 1.34E-02 |
| organic substance catabolic process | GO:1901575 | 1080 | 166 | 118.64 | 1.40 | 7.23E-06 | 2.48E-04 |
| protein modification process | GO:0036211 | 1569 | 241 | 172.36 | 1.40 | 4.54E-08 | 2.25E-06 |
| protein-containing complex organization | GO:0043933 | 918 | 141 | 100.85 | 1.40 | 4.18E-05 | 1.19E-03 |
| cellular process | GO:0009987 | 12722 | 1952 | 1397.57 | 1.40 | 2.42E-107 | 7.25E-104 |

| | | | | | | | |
|---|---|---|---|---|---|---|---|
| metal ion transport | GO:0030001 | 484 | 74 | 53.17 | 1.39 | 3.27E-03 | 4.49E-02 |
| organophosphate metabolic process | GO:0019637 | 642 | 98 | 70.53 | 1.39 | 8.55E-04 | 1.56E-02 |
| inorganic ion transmembrane transport | GO:0098660 | 513 | 78 | 56.36 | 1.38 | 3.35E-03 | 4.57E-02 |
| biosynthetic process | GO:0009058 | 2552 | 387 | 280.35 | 1.38 | 9.45E-12 | 6.97E-10 |
| organelle assembly | GO:0070925 | 594 | 90 | 65.25 | 1.38 | 1.75E-03 | 2.74E-02 |
| organic substance biosynthetic process | GO:1901576 | 2515 | 381 | 276.28 | 1.38 | 1.74E-11 | 1.25E-09 |
| cellular localization | GO:0051641 | 1433 | 217 | 157.42 | 1.38 | 6.90E-07 | 2.89E-05 |
| post-translational protein modification | GO:0043687 | 602 | 91 | 66.13 | 1.38 | 1.88E-03 | 2.87E-02 |
| establishment of localization in cell | GO:0051649 | 940 | 142 | 103.26 | 1.38 | 8.11E-05 | 2.08E-03 |
| catabolic process | GO:0009056 | 1275 | 192 | 140.06 | 1.37 | 5.16E-06 | 1.84E-04 |
| cellular metabolic process | GO:0044237 | 4901 | 738 | 538.40 | 1.37 | 1.75E-22 | 3.50E-20 |
| immune system process | GO:0002376 | 739 | 111 | 81.18 | 1.37 | 6.49E-04 | 1.24E-02 |
| organic cyclic compound biosynthetic process | GO:1901362 | 560 | 84 | 61.52 | 1.37 | 3.23E-03 | 4.45E-02 |
| cellular nitrogen compound metabolic process | GO:0034641 | 2151 | 322 | 236.30 | 1.36 | 3.29E-09 | 1.88E-07 |
| cellular component assembly | GO:0022607 | 1362 | 203 | 149.62 | 1.36 | 5.35E-06 | 1.89E-04 |
| metabolic process | GO:0008152 | 6420 | 953 | 705.27 | 1.35 | 2.63E-28 | 9.88E-26 |
| cellular component biogenesis | GO:0044085 | 1550 | 230 | 170.27 | 1.35 | 1.70E-06 | 6.66E-05 |
| cellular biosynthetic process | GO:0044249 | 2429 | 359 | 266.84 | 1.35 | 1.53E-09 | 9.16E-08 |
| primary metabolic process | GO:0044238 | 5319 | 786 | 584.32 | 1.35 | 9.87E-22 | 1.81E-19 |
| carbohydrate derivative metabolic process | GO:1901135 | 725 | 107 | 79.64 | 1.34 | 1.69E-03 | 2.66E-02 |
| lipid metabolic process | GO:0006629 | 794 | 117 | 87.22 | 1.34 | 9.91E-04 | 1.73E-02 |
| macromolecule localization | GO:0033036 | 1271 | 187 | 139.63 | 1.34 | 2.76E-05 | 8.33E-04 |
| macromolecule biosynthetic process | GO:0009059 | 1815 | 267 | 199.39 | 1.34 | 4.04E-07 | 1.76E-05 |
| response to stress | GO:0006950 | 1295 | 190 | 142.26 | 1.34 | 2.68E-05 | 8.15E-04 |
| macromolecule metabolic process | GO:0043170 | 4105 | 602 | 450.95 | 1.33 | 2.27E-15 | 2.19E-13 |
| organic substance metabolic process | GO:0071704 | 5633 | 825 | 618.81 | 1.33 | 8.48E-22 | 1.62E-19 |
| nitrogen compound transport | GO:0071705 | 915 | 134 | 100.52 | 1.33 | 5.56E-04 | 1.09E-02 |
| biological_process | GO:0008150 | 17206 | 2519 | 1890.16 | 1.33 | 1.10E-171 | 9.91E-168 |
| organonitrogen compound catabolic process | GO:1901565 | 776 | 113 | 85.25 | 1.33 | 1.96E-03 | 2.97E-02 |
| response to external stimulus | GO:0009605 | 772 | 112 | 84.81 | 1.32 | 2.33E-03 | 3.41E-02 |
| nitrogen compound metabolic process | GO:0006807 | 4830 | 700 | 530.60 | 1.32 | 6.93E-17 | 7.99E-15 |
| gene expression | GO:0010467 | 1439 | 207 | 158.08 | 1.31 | 4.43E-05 | 1.22E-03 |
| protein localization | GO:0008104 | 981 | 140 | 107.77 | 1.30 | 1.23E-03 | 2.11E-02 |
| small molecule metabolic process | GO:0044281 | 1052 | 150 | 115.57 | 1.30 | 8.75E-04 | 1.59E-02 |
| cellular macromolecule localization | GO:0070727 | 983 | 140 | 107.99 | 1.30 | 1.25E-03 | 2.14E-02 |
| organonitrogen compound metabolic process | GO:1901564 | 3602 | 496 | 395.70 | 1.25 | 2.15E-08 | 1.10E-06 |
| protein metabolic process | GO:0019538 | 2806 | 385 | 308.25 | 1.25 | 1.89E-06 | 7.28E-05 |

**Supplementary Table 6. Functional grouping of TFs implicated by motif enrichment (adult acinar ATAC-seq).**

Source: HOMER motif enrichment analysis performed on differentially accessible regions (DARs) from adult acinar ATAC-seq (z3′-DpE-/- vs z3′-DpE+/+). Criteria: DARs were defined by edgeR (FDR ≤ 0.05 and |log2FC| ≥ 1). Motifs were considered significantly enriched at HOMER q-value < 0.05 in Up or Down. TFs corresponding to differentially enriched motifs were grouped into functional categories based on curated literature evidence; unmatched TFs are labeled “Other”.

| Motif (HOMER) | Enriched in Down/Up DARs | ΔΔ% Enrichment over Background (Up - Down) | TF gene (human) | Functional category |
|---|---|---|---|---|
| Ptf1a(bHLH)/Panc1-Ptf1a-ChIP-Seq(GSE47459)/Homer | Down | -3.37 | PTF1A | Acinar Identity |
| BHLHA15(bHLH)/NIH3T3-BHLHB8.HA-ChIP-Seq(GSE119782)/Homer | Down | -1.59 | BHLHA15 | Acinar Identity |
| Nr5a2(NR)/Pancreas-LRH1-ChIP-Seq(GSE34295)/Homer | Down | -1.31 | NR5A2 | Acinar Identity |
| Gata4(Zf)/Heart-Gata4-ChIP-Seq(GSE35151)/Homer | Both | -0.82 | GATA4 | Acinar Identity |
| Nr5a2(NR)/mES-Nr5a2-ChIP-Seq(GSE19019)/Homer | Both | -0.49 | NR5A2 | Acinar Identity |
| E2A(bHLH)/proBcell-E2A-ChIP-Seq(GSE21978)/Homer | Down | -3.29 | TCF3 | Ptf1a Co-factors |
| HEB(bHLH)/mES-Heb-ChIP-Seq(GSE53233)/Homer | Down | -3.05 | TCF12 | Ptf1a Co-factors |
| TCF4(bHLH)/SHSY5Y-TCF4-ChIP-Seq(GSE96915)/Homer | Down | -1.17 | TCF4 | Ptf1a Co-factors |
| RBPJ:Ebox(?,bHLH)/Panc1-Rbpj1-ChIP-Seq(GSE47459)/Homer | Both | -0.42 | RBPJ | Ptf1a Co-factors |
| Ascl1(bHLH)/NeuralTubes-Ascl1-ChIP-Seq(GSE55840)/Homer | Down | -3.01 | ASCL1 | Endocrine/Progenitor Identity |
| Nkx6.1(Homeobox)/Islet-Nkx6.1-ChIP-Seq(GSE40975)/Homer | Down | -2.87 | NKX6-1 | Endocrine/Progenitor Identity |
| Ascl2(bHLH)/ESC-Ascl2-ChIP-Seq(GSE97712)/Homer | Down | -2.3 | ASCL2 | Endocrine/Progenitor Identity |
| NeuroD1(bHLH)/Islet-NeuroD1-ChIP-Seq(GSE30298)/Homer | Both | -2.24 | NEUROD1 | Endocrine/Progenitor Identity |
| Foxa2(Forkhead)/Liver-Foxa2-ChIP-Seq(GSE25694)/Homer | Both | -0.71 | FOXA2 | Endocrine/Progenitor Identity |
| FOXA1(Forkhead)/LNCAP-FOXA1-ChIP-Seq(GSE27824)/Homer | Both | -0.65 | FOXA1 | Endocrine/Progenitor Identity |
| FOXA1(Forkhead)/MCF7-FOXA1-ChIP-Seq(GSE26831)/Homer | Both | -0.46 | FOXA1 | Endocrine/Progenitor Identity |
| HNF1b(Homeobox)/PDAC-HNF1B-ChIP-Seq(GSE64557)/Homer | Both | 0.68 | HNF1B | Endocrine/Progenitor Identity |
| Fox:Ebox(Forkhead,bHLH)/Panc1-Foxa2-ChIP-Seq(GSE47459)/Homer | Up | 3.24 | FOXA2 | Endocrine/Progenitor Identity |
| JunD(bZIP)/K562-JunD-ChIP-Seq/Homer | Both | -0.34 | JUND | AP-1 complex |
| MafA(bZIP)/Islet-MafA-ChIP-Seq(GSE30298)/Homer | Down | -0.18 | MAFA | AP-1 complex |
| Atf2(bZIP)/3T3L1-Atf2-ChIP-Seq(GSE56872)/Homer | Both | 0.47 | ATF2 | AP-1 complex |
| Jun-AP1(bZIP)/K562-cJun-ChIP-Seq(GSE31477)/Homer | Up | 0.93 | JUN | AP-1 complex |
| Atf7(bZIP)/3T3L1-Atf7-ChIP-Seq(GSE56872)/Homer | Both | 1.19 | ATF7 | AP-1 complex |
| Fosl2(bZIP)/3T3L1-Fosl2-ChIP-Seq(GSE56872)/Homer | Up | 1.51 | FOSL2 | AP-1 complex |

| | | | | |
|---|---|---|---|---|
| Fra2(bZIP)/Striatum-Fra2-ChIP-Seq(GSE43429)/Homer | Up | 1.96 | FOSL2 | AP-1 complex |
| JunB(bZIP)/DendriticCells-Junb-ChIP-Seq(GSE36099)/Homer | Up | 2.13 | JUNB | AP-1 complex |
| Fos(bZIP)/TSC-Fos-ChIP-Seq(GSE110950)/Homer | Up | 2.32 | FOS | AP-1 complex |
| Atf1(bZIP)/K562-ATF1-ChIP-Seq(GSE31477)/Homer | Both | 2.45 | ATF1 | AP-1 complex |
| AP-1(bZIP)/ThioMac-PU.1-ChIP-Seq(GSE21512)/Homer | Up | 2.46 | JUN | AP-1 complex |
| Fra1(bZIP)/BT549-Fra1-ChIP-Seq(GSE46166)/Homer | Up | 2.47 | FOSL1 | AP-1 complex |
| BATF(bZIP)/Th17-BATF-ChIP-Seq(GSE39756)/Homer | Up | 2.55 | BATF | AP-1 complex |
| Atf4(bZIP)/MEF-Atf4-ChIP-Seq(GSE35681)/Homer | Up | 2.67 | ATF4 | AP-1 complex |
| Atf3(bZIP)/GBM-ATF3-ChIP-Seq(GSE33912)/Homer | Up | 2.81 | ATF3 | AP-1 complex |
| CTCF(Zf)/CD4+-CTCF-ChIP-Seq(Barski_et_al.)/Homer | Both | -8.58 | CTCF | association to pancreatic neoplasm |
| KLF10(Zf)/HEK293-KLF10.GFP-ChIP-Seq(GSE58341)/Homer | Both | -4.37 | KLF10 | association to pancreatic neoplasm |
| Snail1(Zf)/LS174T-SNAIL1.HA-ChIP-Seq(GSE127183)/Homer | Down | -3.18 | SNAI1 | association to pancreatic neoplasm |
| HIC1(Zf)/Treg-ZBTB29-ChIP-Seq(GSE99889)/Homer | Down | -1.31 | HIC1 | association to pancreatic neoplasm |
| Esrrb(NR)/mES-Esrrb-ChIP-Seq(GSE11431)/Homer | Down | -0.96 | ESRRB | association to pancreatic neoplasm |
| Bcl11a(Zf)/HSPC-BCL11A-ChIP-Seq(GSE104676)/Homer | Both | -0.85 | BCL11A | association to pancreatic neoplasm |
| E2F3(E2F)/MEF-E2F3-ChIP-Seq(GSE71376)/Homer | Down | -0.77 | E2F3 | association to pancreatic neoplasm |
| E2F1(E2F)/Hela-E2F1-ChIP-Seq(GSE22478)/Homer | Down | -0.73 | E2F1 | association to pancreatic neoplasm |
| Gata2(Zf)/K562-GATA2-ChIP-Seq(GSE18829)/Homer | Both | -0.63 | GATA2 | association to pancreatic neoplasm |
| Gata1(Zf)/K562-GATA1-ChIP-Seq(GSE18829)/Homer | Both | -0.58 | GATA1 | association to pancreatic neoplasm |
| ZFX(Zf)/mES-Zfx-ChIP-Seq(GSE11431)/Homer | Down | -0.57 | ZFX | association to pancreatic neoplasm |
| REST-NRSF(Zf)/Jurkat-NRSF-ChIP-Seq/Homer | Both | -0.39 | REST | association to pancreatic neoplasm |
| GATA3(Zf)/iTreg-Gata3-ChIP-Seq(GSE20898)/Homer | Both | -0.35 | GATA5 | association to pancreatic neoplasm |
| Gata6(Zf)/HUG1N-GATA6-ChIP-Seq(GSE51936)/Homer | Both | -0.21 | GATA6 | association to pancreatic neoplasm |
| YY1(Zf)/Promoter/Homer | Both | 0 | YY1 | association to pancreatic neoplasm |
| FOXM1(Forkhead)/MCF7-FOXM1-ChIP-Seq(GSE72977)/Homer | Both | 0 | FOXM1 | association to pancreatic neoplasm |
| Bach1(bZIP)/K562-Bach1-ChIP-Seq(GSE31477)/Homer | Up | 0.16 | BACH1 | association to pancreatic neoplasm |
| Mef2d(MADS)/Retina-Mef2d-ChIP-Seq(GSE61391)/Homer | Up | 0.18 | MEF2D | association to pancreatic neoplasm |
| Zfp281(Zf)/ES-Zfp281-ChIP-Seq(GSE81042)/Homer | Up | 0.22 | ZNF281 | association to pancreatic neoplasm |
| NFE2L2(bZIP)/HepG2-NFE2L2-ChIP-Seq(Encode)/Homer | Up | 0.29 | NFE2L2 | association to pancreatic neoplasm |
| VDR(NR),DR3/GM10855-VDR+vitD-ChIP-Seq(GSE22484)/Homer | Up | 0.31 | VDR | association to pancreatic neoplasm |
| Foxo3(Forkhead)/U2OS-Foxo3-ChIP-Seq(E-MTAB-2701)/Homer | Up | 0.45 | FOXO3 | association to pancreatic neoplasm |
| Six1(Homeobox)/Myoblast-Six1-ChIP-Chip(GSE20150)/Homer | Up | 0.51 | SIX1 | association to pancreatic neoplasm |
| Pbx3(Homeobox)/GM12878-PBX3-ChIP-Seq(GSE32465)/Homer | Up | 0.53 | PBX3 | association to pancreatic neoplasm |
| Cdx2(Homeobox)/mES-Cdx2-ChIP-Seq(GSE14586)/Homer | Up | 0.57 | CDX2 | association to pancreatic neoplasm |

| | | | | |
|---|---|---|---|---|
| Sp1(Zf)/Promoter/Homer | Up | 0.58 | SP1 | association to pancreatic neoplasm |
| RORa(NR)/Liver-Rora-ChIP-Seq(GSE101115)/Homer | Up | 0.64 | RORA | association to pancreatic neoplasm |
| Mef2a(MADS)/HL1-Mef2a.biotin-ChIP-Seq(GSE21529)/Homer | Up | 0.64 | MEF2A | association to pancreatic neoplasm |
| PGR(NR)/EndoStromal-PGR-ChIP-Seq(GSE69539)/Homer | Up | 0.71 | PGR | association to pancreatic neoplasm |
| FOXK1(Forkhead)/HEK293-FOXK1-ChIP-Seq(GSE51673)/Homer | Up | 0.73 | FOXK1 | association to pancreatic neoplasm |
| STAT1(Stat)/HelaS3-STAT1-ChIP-Seq(GSE12782)/Homer | Up | 0.76 | STAT1 | association to pancreatic neoplasm |
| Stat3(Stat)/mES-Stat3-ChIP-Seq(GSE11431)/Homer | Up | 0.77 | STAT3 | association to pancreatic neoplasm |
| c-Myc(bHLH)/mES-cMyc-ChIP-Seq(GSE11431)/Homer | Up | 0.82 | MYC | association to pancreatic neoplasm |
| WT1(Zf)/Kidney-WT1-ChIP-Seq(GSE90016)/Homer | Up | 0.95 | WT1 | association to pancreatic neoplasm |
| PPARa(NR),DR1/Liver-Ppara-ChIP-Seq(GSE47954)/Homer | Up | 1.1 | PPARA | association to pancreatic neoplasm |
| NRF(NRF)/Promoter/Homer | Up | 1.12 | NKRF | association to pancreatic neoplasm |
| bHLHE41(bHLH)/proB-Bhlhe41-ChIP-Seq(GSE93764)/Homer | Up | 1.37 | BHLHE41 | association to pancreatic neoplasm |
| Egr1(Zf)/K562-Egr1-ChIP-Seq(GSE32465)/Homer | Up | 1.42 | EGR1 | association to pancreatic neoplasm |
| IRF2(IRF)/Erythroblas-IRF2-ChIP-Seq(GSE36985)/Homer | Up | 1.69 | IRF2 | association to pancreatic neoplasm |
| Chop(bZIP)/MEF-Chop-ChIP-Seq(GSE35681)/Homer | Up | 1.7 | DDIT3 | association to pancreatic neoplasm |
| PPARE(NR),DR1/3T3L1-Pparg-ChIP-Seq(GSE13511)/Homer | Up | 1.76 | PPARG | association to pancreatic neoplasm |
| NFkB-p65(RHD)/GM12787-p65-ChIP-Seq(GSE19485)/Homer | Up | 1.85 | RELA | association to pancreatic neoplasm |
| IRF1(IRF)/PBMC-IRF1-ChIP-Seq(GSE43036)/Homer | Up | 2.06 | IRF1 | association to pancreatic neoplasm |
| RUNX(Runt)/HPC7-Runx1-ChIP-Seq(GSE22178)/Homer | Up | 2.44 | RUNX1 | association to pancreatic neoplasm |
| KLF5(Zf)/LoVo-KLF5-ChIP-Seq(GSE49402)/Homer | Up | 2.49 | KLF5 | association to pancreatic neoplasm |
| RUNX1(Runt)/Jurkat-RUNX1-ChIP-Seq(GSE29180)/Homer | Up | 2.54 | RUNX1 | association to pancreatic neoplasm |
| PRDM1(Zf)/Hela-PRDM1-ChIP-Seq(GSE31477)/Homer | Up | 2.66 | PRDM1 | association to pancreatic neoplasm |
| RUNX2(Runt)/PCa-RUNX2-ChIP-Seq(GSE33889)/Homer | Up | 2.68 | RUNX2 | association to pancreatic neoplasm |
| CLOCK(bHLH)/Liver-Clock-ChIP-Seq(GSE39860)/Homer | Up | 2.83 | CLOCK | association to pancreatic neoplasm |
| Foxo1(Forkhead)/RAW-Foxo1-ChIP-Seq(Fan_et_al.)/Homer | Up | 3.27 | FOXO1 | association to pancreatic neoplasm |
| PR(NR)/T47D-PR-ChIP-Seq(GSE31130)/Homer | Up | 3.76 | PGR | association to pancreatic neoplasm |
| MITF(bHLH)/MastCells-MITF-ChIP-Seq(GSE48085)/Homer | Both | 4.02 | MITF | association to pancreatic neoplasm |
| Tgif1(Homeobox)/mES-Tgif1-ChIP-Seq(GSE55404)/Homer | Up | 4.22 | TGIF1 | association to pancreatic neoplasm |
| IRF8(IRF)/BMDM-IRF8-ChIP-Seq(GSE77884)/Homer | Up | 4.63 | IRF8 | association to pancreatic neoplasm |
| GABPA(ETS)/Jurkat-GABPa-ChIP-Seq(GSE17954)/Homer | Up | 9.48 | GABPA | association to pancreatic neoplasm |
| ETV4(ETS)/HepG2-ETV4-ChIP-Seq(ENCODE)/Homer | Up | 10.06 | ETV4 | association to pancreatic neoplasm |
| Fli1(ETS)/CD8-FLI-ChIP-Seq(GSE20898)/Homer | Up | 10.91 | FLI1 | association to pancreatic neoplasm |

| | | | | |
|---|---|---|---|---|
| ETS1(ETS)/Jurkat-ETS1-ChIP-Seq(GSE17954)/Homer | Up | 11.03 | ETS1 | association to pancreatic neoplasm |
| ERG(ETS)/VCaP-ERG-ChIP-Seq(GSE14097)/Homer | Up | 11.83 | ERG | association to pancreatic neoplasm |
| EHF(ETS)/LoVo-EHF-ChIP-Seq(GSE49402)/Homer | Up | 12.46 | EHF | association to pancreatic neoplasm |
| BORIS(Zf)/K562-CTCFL-ChIP-Seq(GSE32465)/Homer | Both | -8.51 | CTCFL | Other |
| CRX(Homeobox)/Retina-Crx-ChIP-Seq(GSE20012)/Homer | Down | -3.39 | CRX | Other |
| Pitx1(Homeobox)/Chicken-Pitx1-ChIP-Seq(GSE38910)/Homer | Down | -3.24 | PITX1 | Other |
| GSC(Homeobox)/FrogEmbryos-GSC-ChIP-Seq(DRA000576)/Homer | Down | -2.86 | GSC | Other |
| MyoG(bHLH)/C2C12-MyoG-ChIP-Seq(GSE36024)/Homer | Down | -2.58 | MYOG | Other |
| Slug(Zf)/Mesoderm-Snai2-ChIP-Seq(GSE61475)/Homer | Down | -1.94 | SNAI2 | Other |
| Unknown-ESC-element(?)/mES-Nanog-ChIP-Seq(GSE11724)/Homer | Down | -1.92 | | Other |
| Ap4(bHLH)/AML-Tfap4-ChIP-Seq(GSE45738)/Homer | Down | -1.8 | TFAP4 | Other |
| Atoh1(bHLH)/Cerebellum-Atoh1-ChIP-Seq(GSE22111)/Homer | Down | -1.8 | ATOH1 | Other |
| Myf5(bHLH)/GM-Myf5-ChIP-Seq(GSE24852)/Homer | Down | -1.77 | MYF5 | Other |
| Barx1(Homeobox)/Stomach-Barx1.3xFlag-ChIP-Seq(GSE69483)/Homer | Down | -1.74 | BARX1 | Other |
| Otx2(Homeobox)/EpiLC-Otx2-ChIP-Seq(GSE56098)/Homer | Down | -1.68 | OTX2 | Other |
| Zac1(Zf)/Neuro2A-Plagl1-ChIP-Seq(GSE75942)/Homer | Down | -1.64 | PLAGL1 | Other |
| OCT:OCT(POU,Homeobox)/NPC-OCT6-ChIP-Seq(GSE43916)/Homer | Down | -1.54 | | Other |
| Tcf12(bHLH)/GM12878-Tcf12-ChIP-Seq(GSE32465)/Homer | Down | -1.51 | TCF12 | Other |
| LRF(Zf)/Erythroblasts-ZBTB7A-ChIP-Seq(GSE74977)/Homer | Down | -1.46 | ZBTB7A | Other |
| Tcf21(bHLH)/ArterySmoothMuscle-Tcf21-ChIP-Seq(GSE61369)/Homer | Down | -1.43 | TCF21 | Other |
| E2F4(E2F)/K562-E2F4-ChIP-Seq(GSE31477)/Homer | Down | -1.41 | E2F4 | Other |
| TRPS1(Zf)/MCF7-TRPS1-ChIP-Seq(GSE107013)/Homer | Both | -1.36 | TRPS1 | Other |
| Zic3(Zf)/mES-Zic3-ChIP-Seq(GSE37889)/Homer | Down | -1.29 | ZIC3 | Other |
| Olig2(bHLH)/Neuron-Olig2-ChIP-Seq(GSE30882)/Homer | Down | -1.24 | OLIG2 | Other |
| ZNF415(Zf)/HEK293-ZNF415.GFP-ChIP-Seq(GSE58341)/Homer | Down | -1.14 | ZNF415 | Other |
| MyoD(bHLH)/Myotube-MyoD-ChIP-Seq(GSE21614)/Homer | Both | -1.06 | MYOD1 | Other |
| NeuroG2(bHLH)/Fibroblast-NeuroG2-ChIP-Seq(GSE75910)/Homer | Down | -0.96 | NEUROG2 | Other |
| ZBTB18(Zf)/HEK293-ZBTB18.GFP-ChIP-Seq(GSE58341)/Homer | Down | -0.95 | ZBTB18 | Other |
| Erra(NR)/HepG2-Erra-ChIP-Seq(GSE31477)/Homer | Both | -0.87 | ESRRA | Other |
| E2F6(E2F)/Hela-E2F6-ChIP-Seq(GSE31477)/Homer | Down | -0.79 | E2F6 | Other |
| Foxf1(Forkhead)/Lung-Foxf1-ChIP-Seq(GSE77951)/Homer | Down | -0.75 | FOXF1 | Other |
| MYNN(Zf)/HEK293-MYNN.eGFP-ChIP-Seq(Encode)/Homer | Down | -0.67 | MYNN | Other |
| NF1-halfsite(CTF)/LNCaP-NF1-ChIP-Seq(Unpublished)/Homer | Down | -0.66 | | Other |
| FoxL2(Forkhead)/Ovary-FoxL2-ChIP-Seq(GSE60858)/Homer | Down | -0.63 | FOXL2 | Other |
| THRb(NR)/HepG2-THRb.Flag-ChIP-Seq(Encode)/Homer | Both | -0.53 | THRB | Other |
| E2F7(E2F)/Hela-E2F7-ChIP-Seq(GSE32673)/Homer | Down | -0.51 | E2F7 | Other |
| EBF1(EBF)/Near-E2A-ChIP-Seq(GSE21512)/Homer | Down | -0.5 | EBF1 | Other |
| ZNF711(Zf)/SHSY5Y-ZNF711-ChIP-Seq(GSE20673)/Homer | Down | -0.48 | ZNF711 | Other |

| | | | | |
|---|---|---|---|---|
| PBX2(Homeobox)/K562-PBX2-ChIP-Seq(Encode)/Homer | Down | -0.48 | PBX2 | Other |
| ZNF341(Zf)/EBV-ZNF341-ChIP-Seq(GSE113194)/Homer | Down | -0.44 | ZNF341 | Other |
| SF1(NR)/H295R-Nr5a1-ChIP-Seq(GSE44220)/Homer | Down | -0.42 | SF1 | Other |
| Pit1+1bp(Homeobox)/GCrat-Pit1-ChIP-Seq(GSE58009)/Homer | Down | -0.35 | | Other |
| E2F(E2F)/Hela-CellCycle-Expression/Homer | Down | -0.35 | | Other |
| HRE(HSF)/HepG2-HSF1-ChIP-Seq(GSE31477)/Homer | Down | -0.33 | | Other |
| Pax7(Paired,Homeobox),long/Myoblast-Pax7-ChIP-Seq(GSE25064)/Homer | Down | -0.23 | PAX7 | Other |
| CTCF-SatelliteElement(Zf?)/CD4+-CTCF-ChIP-Seq(Barski_et_al.)/Homer | Both | -0.2 | | Other |
| Zic2(Zf)/ESC-Zic2-ChIP-Seq(SRP197560)/Homer | Down | -0.17 | ZIC2 | Other |
| Foxa3(Forkhead)/Liver-Foxa3-ChIP-Seq(GSE77670)/Homer | Down | -0.13 | FOXA3 | Other |
| Hoxa13(Homeobox)/ChickenMSG-Hoxa13.Flag-ChIP-Seq(GSE86088)/Homer | Both | -0.09 | HOXA13 | Other |
| Zic(Zf)/Cerebellum-ZIC1.2-ChIP-Seq(GSE60731)/Homer | Down | -0.08 | | Other |
| ZNF768(Zf)/Rajj-ZNF768-ChIP-Seq(GSE111879)/Homer | Up | 0.13 | ZNF768 | Other |
| Rfx1(HTH)/NPC-H3K4me1-ChIP-Seq(GSE16256)/Homer | Up | 0.13 | RFX1 | Other |
| Six4(Homeobox)/MCF7-SIX4-ChIP-Seq(Encode)/Homer | Up | 0.16 | SIX4 | Other |
| LXRE(NR),DR4/RAW-LXRb.biotin-ChIP-Seq(GSE21512)/Homer | Up | 0.18 | | Other |
| GFX(?)/Promoter/Homer | Up | 0.19 | | Other |
| CRE(bZIP)/Promoter/Homer | Both | 0.2 | | Other |
| X-box(HTH)/NPC-H3K4me1-ChIP-Seq(GSE16256)/Homer | Up | 0.21 | | Other |
| T1ISRE(IRF)/ThioMac-Ifnb-Expression/Homer | Up | 0.22 | | Other |
| NF1:FOXA1(CTF,Forkhead)/LNCAP-FOXA1-ChIP-Seq(GSE27824)/Homer | Up | 0.23 | NFE2L2 | Other |
| COUP-TFII(NR)/Artia-Nr2f2-ChIP-Seq(GSE46497)/Homer | Both | 0.24 | NR2F2 | Other |
| RAR:RXR(NR),DR5/ES-RAR-ChIP-Seq(GSE56893)/Homer | Up | 0.25 | | Other |
| RORg(NR)/Liver-Rorc-ChIP-Seq(GSE101115)/Homer | Both | 0.28 | RORC | Other |
| Bach2(bZIP)/OCILy7-Bach2-ChIP-Seq(GSE44420)/Homer | Up | 0.3 | BACH2 | Other |
| E-box(bHLH)/Promoter/Homer | Both | 0.33 | | Other |
| NRF1(NRF)/MCF7-NRF1-ChIP-Seq(Unpublished)/Homer | Up | 0.36 | NRF1 | Other |
| c-Jun-CRE(bZIP)/K562-cJun-ChIP-Seq(GSE31477)/Homer | Both | 0.36 | | Other |
| GRE(NR),IR3/A549-GR-ChIP-Seq(GSE32465)/Homer | Up | 0.38 | | Other |
| ETS:E-box(ETS,bHLH)/HPC7-Scl-ChIP-Seq(GSE22178)/Homer | Up | 0.41 | | Other |
| OCT:OCT(POU,Homeobox)/NPC-Brn1-ChIP-Seq(GSE35496)/Homer | Up | 0.42 | | Other |
| PBX1(Homeobox)/MCF7-PBX1-ChIP-Seq(GSE28007)/Homer | Up | 0.49 | PBX1 | Other |
| IRF:BATF(IRF:bZIP)/pDC-Irf8-ChIP-Seq(GSE66899)/Homer | Up | 0.49 | | Other |
| NF1(CTF)/LNCAP-NF1-ChIP-Seq(Unpublished)/Homer | Up | 0.49 | NF1 | Other |
| Pknox1(Homeobox)/ES-Prep1-ChIP-Seq(GSE63282)/Homer | Up | 0.51 | PKNOX1 | Other |
| PSE(SNAPc)/K562-mStart-Seq/Homer | Up | 0.52 | | Other |
| RORgt(NR)/EL4-RORgt.Flag-ChIP-Seq(GSE56019)/Homer | Up | 0.54 | RORC | Other |
| ARE(NR)/LNCAP-AR-ChIP-Seq(GSE27824)/Homer | Up | 0.57 | | Other |
| Hoxa10(Homeobox)/ChickenMSG-Hoxa10.Flag-ChIP-Seq(GSE86088)/Homer | Up | 0.57 | HOXA10 | Other |
| FOXK2(Forkhead)/U2OS-FOXK2-ChIP-Seq(E-MTAB-2204)/Homer | Up | 0.57 | FOXK2 | Other |

| | | | | |
|---|---|---|---|---|
| DMRT1(DM)/Testis-DMRT1-ChIP-Seq(GSE64892)/Homer | Up | 0.58 | DMRT1 | Other |
| Hnf1(Homeobox)/Liver-Foxa2-Chip-Seq(GSE25694)/Homer | Both | 0.59 | | Other |
| ISRE(IRF)/ThioMac-LPS-Expression(GSE23622)/Homer | Up | 0.59 | | Other |
| THRa(NR)/C17.2-THRa-ChIP-Seq(GSE38347)/Homer | Up | 0.6 | THRA | Other |
| GFY(?)/Promoter/Homer | Up | 0.61 | GFY | Other |
| ZNF143\|STAF(Zf)/CUTLL-ZNF143-ChIP-Seq(GSE29600)/Homer | Up | 0.61 | | Other |
| Tlx?(NR)/NPC-H3K4me1-ChIP-Seq(GSE16256)/Homer | Up | 0.64 | | Other |
| NFkB-p65-Rel(RHD)/ThioMac-LPS-Expression(GSE23622)/Homer | Up | 0.73 | | Other |
| Ronin(THAP)/ES-Thap11-ChIP-Seq(GSE51522)/Homer | Up | 0.76 | THAP11 | Other |
| Mef2c(MADS)/GM12878-Mef2c-ChIP-Seq(GSE32465)/Homer | Up | 0.77 | MEF2C | Other |
| ETS:RUNX(ETS,Runt)/Jurkat-RUNX1-ChIP-Seq(GSE17954)/Homer | Up | 0.77 | | Other |
| Oct11(POU,Homeobox)/NCIH1048-POU2F3-ChIP-seq(GSE115123)/Homer | Up | 0.77 | POU2F3 | Other |
| CREB5(bZIP)/LNCaP-CREB5.V5-ChIP-Seq(GSE137775)/Homer | Both | 0.81 | CREB5 | Other |
| EKLF(Zf)/Erythrocyte-Klf1-ChIP-Seq(GSE20478)/Homer | Up | 0.82 | KLF1 | Other |
| GFY-Staf(?,Zf)/Promoter/Homer | Up | 0.88 | | Other |
| CDX4(Homeobox)/ZebrafishEmbryos-Cdx4.Myc-ChIP-Seq(GSE48254)/Homer | Up | 0.93 | CDX4 | Other |
| NFAT(RHD)/Jurkat-NFATC1-ChIP-Seq(Jolma_et_al.)/Homer | Up | 0.93 | | Other |
| TFE3(bHLH)/MEF-TFE3-ChIP-Seq(GSE75757)/Homer | Up | 0.96 | TFE3 | Other |
| Hoxd12(Homeobox)/ChickenMSG-Hoxd12.Flag-ChIP-Seq(GSE86088)/Homer | Up | 1.01 | HOXD12 | Other |
| Hoxa11(Homeobox)/ChickenMSG-Hoxa11.Flag-ChIP-Seq(GSE86088)/Homer | Up | 1.06 | HOXA11 | Other |
| GRE(NR),IR3/RAW264.7-GRE-ChIP-Seq(Unpublished)/Homer | Up | 1.07 | | Other |
| Egr2(Zf)/Thymocytes-Egr2-ChIP-Seq(GSE34254)/Homer | Up | 1.15 | EGR2 | Other |
| Maz(Zf)/HepG2-Maz-ChIP-Seq(GSE31477)/Homer | Up | 1.16 | MAZ | Other |
| STAT4(Stat)/CD4-Stat4-ChIP-Seq(GSE22104)/Homer | Up | 1.21 | STAT4 | Other |
| RXR(NR),DR1/3T3L1-RXR-ChIP-Seq(GSE13511)/Homer | Up | 1.23 | | Other |
| KLF3(Zf)/MEF-Klf3-ChIP-Seq(GSE44748)/Homer | Up | 1.25 | KLF3 | Other |
| SCL(bHLH)/HPC7-Scl-ChIP-Seq(GSE13511)/Homer | Up | 1.28 | TAL1 | Other |
| Mef2b(MADS)/HEK293-Mef2b.V5-ChIP-Seq(GSE67450)/Homer | Up | 1.44 | MEF2B | Other |
| Stat3+il21(Stat)/CD4-Stat3-ChIP-Seq(GSE19198)/Homer | Up | 1.45 | | Other |
| bZIP:IRF(bZIP,IRF)/Th17-BatF-ChIP-Seq(GSE39756)/Homer | Up | 1.52 | | Other |
| n-Myc(bHLH)/mES-nMyc-ChIP-Seq(GSE11431)/Homer | Up | 1.55 | MYCN | Other |
| NPAS2(bHLH)/Liver-NPAS2-ChIP-Seq(GSE39860)/Homer | Up | 1.87 | NPAS2 | Other |
| Hoxd10(Homeobox)/ChickenMSG-Hoxd10.Flag-ChIP-Seq(GSE86088)/Homer | Up | 1.9 | HOXD10 | Other |
| Npas4(bHLH)/Neuron-Npas4-ChIP-Seq(GSE127793)/Homer | Up | 1.94 | NPAS4 | Other |
| RUNX-AML(Runt)/CD4+-PolII-ChIP-Seq(Barski_et_al.)/Homer | Up | 2.01 | | Other |
| IRF4(IRF)/GM12878-IRF4-ChIP-Seq(GSE32465)/Homer | Up | 2.12 | IRF4 | Other |
| Meis1(Homeobox)/MastCells-Meis1-ChIP-Seq(GSE48085)/Homer | Up | 2.15 | MEIS1 | Other |
| KLF1(Zf)/HUDEP2-KLF1-CutnRun(GSE136251)/Homer | Up | 2.21 | KLF1 | Other |

| | | | | |
|---|---|---|---|---|
| Sp5(Zf)/mES-Sp5.Flag-ChIP-Seq(GSE72989)/Homer | Up | 2.3 | SP5 | Other |
| Max(bHLH)/K562-Max-ChIP-Seq(GSE31477)/Homer | Up | 2.52 | MAX | Other |
| Sp2(Zf)/HEK293-Sp2.eGFP-ChIP-Seq(Encode)/Homer | Up | 2.63 | SP2 | Other |
| KLF14(Zf)/HEK293-KLF14.GFP-ChIP-Seq(GSE58341)/Homer | Up | 2.68 | KLF14 | Other |
| USF1(bHLH)/GM12878-Usf1-ChIP-Seq(GSE32465)/Homer | Up | 2.7 | USF1 | Other |
| Usf2(bHLH)/C2C12-Usf2-ChIP-Seq(GSE36030)/Homer | Up | 2.72 | USF2 | Other |
| IRF3(IRF)/BMDM-Irf3-ChIP-Seq(GSE67343)/Homer | Up | 3.27 | IRF3 | Other |
| Hoxa9(Homeobox)/ChickenMSG-Hoxa9.Flag-ChIP-Seq(GSE86088)/Homer | Up | 3.31 | HOXA9 | Other |
| CEBP:AP1(bZIP)/ThioMac-CEBPb-ChIP-Seq(GSE21512)/Homer | Up | 3.34 | | Other |
| Tgif2(Homeobox)/mES-Tgif2-ChIP-Seq(GSE55404)/Homer | Up | 3.5 | TGIF2 | Other |
| NFY(CCAAT)/Promoter/Homer | Up | 3.57 | | Other |
| PU.1:IRF8(ETS:IRF)/pDC-Irf8-ChIP-Seq(GSE66899)/Homer | Up | 3.63 | | Other |
| ETS(ETS)/Promoter/Homer | Up | 4.2 | | Other |
| NFIL3(bZIP)/HepG2-NFIL3-ChIP-Seq(Encode)/Homer | Both | 4.25 | NFIL3 | Other |
| HLF(bZIP)/HSC-HLF.Flag-ChIP-Seq(GSE69817)/Homer | Both | 4.64 | HLF | Other |
| MNT(bHLH)/HepG2-MNT-ChIP-Seq(Encode)/Homer | Up | 5.03 | MNT | Other |
| ELF1(ETS)/Jurkat-ELF1-ChIP-Seq(SRA014231)/Homer | Up | 5.03 | ELF1 | Other |
| Elk4(ETS)/Hela-Elk4-ChIP-Seq(GSE31477)/Homer | Up | 5.08 | ELK4 | Other |
| Elk1(ETS)/Hela-Elk1-ChIP-Seq(GSE31477)/Homer | Up | 5.2 | ELK1 | Other |
| SPDEF(ETS)/VCaP-SPDEF-ChIP-Seq(SRA014231)/Homer | Up | 5.47 | SPDEF | Other |
| Ets1-distal(ETS)/CD4+-PolII-ChIP-Seq(Barski_et_al.)/Homer | Up | 5.59 | | Other |
| EWS:FLI1-fusion(ETS)/SK_N_MC-EWS:FLI1-ChIP-Seq(SRA014231)/Homer | Up | 5.79 | | Other |
| NPAS(bHLH)/Liver-NPAS-ChIP-Seq(GSE39860)/Homer | Up | 6.31 | NPAS2 | Other |
| SpiB(ETS)/OCILY3-SPIB-ChIP-Seq(GSE56857)/Homer | Up | 6.76 | SPIB | Other |
| CEBP(bZIP)/ThioMac-CEBPb-ChIP-Seq(GSE21512)/Homer | Up | 7.11 | | Other |
| BMAL1(bHLH)/Liver-Bmal1-ChIP-Seq(GSE39860)/Homer | Up | 7.89 | BMAL1 | Other |
| EWS:ERG-fusion(ETS)/CADO_ES1-EWS:ERG-ChIP-Seq(SRA014231)/Homer | Up | 8.26 | | Other |
| PU.1-IRF(ETS:IRF)/Bcell-PU.1-ChIP-Seq(GSE21512)/Homer | Up | 8.39 | | Other |
| ELF3(ETS)/PDAC-ELF3-ChIP-Seq(GSE64557)/Homer | Up | 8.59 | ELF3 | Other |
| PU.1(ETS)/ThioMac-PU.1-ChIP-Seq(GSE21512)/Homer | Up | 9.02 | SPI1 | Other |
| ELF5(ETS)/T47D-ELF5-ChIP-Seq(GSE30407)/Homer | Up | 9.18 | ELF5 | Other |
| Etv2(ETS)/ES-ER71-ChIP-Seq(GSE59402)/Homer | Up | 9.82 | ETV2 | Other |
| ETV1(ETS)/GIST48-ETV1-ChIP-Seq(GSE22441)/Homer | Up | 11.65 | ETV1 | Other |
| Elf4(ETS)/BMDM-Elf4-ChIP-Seq(GSE88699)/Homer | Up | 11.82 | ELF4 | Other |
